# Landscape of glioblastoma niches reveals the prognostic effects of tumor-infiltrating cells

**DOI:** 10.1101/2021.01.20.427411

**Authors:** Zixuan Xiao, Wei Zhang, Guanzhang Li, Wendong Li, Lin Li, Ting Sun, Yufei He, Guang Liu, Lu Wang, Xiaohan Han, Hao Wen, Yong Liu, Yifan Chen, Haoyu Wang, Jing Li, Yubo Fan, Jing Zhang

## Abstract

A comprehensive characterization of non-tumor cells in the niches of primary glioblastoma is not fully established yet. This study aims to present an overview of tumor-infiltrating non-malignant cells in the complex microenvironment of glioblastoma with detailed characterizations of their prognostic effects. We curate 540 gene signatures covering a total of 64 non-tumor cell types. Cell type-specific expression patterns are interrogated by normalized enrichment score (NES) across four large gene expression profiling cohorts of glioblastoma with a total number of 967 cases. The GBMs in each cohort are hierarchically clustered into negative or positive immune response classes with significantly different overall survival. Our results show that astrocytes, macrophages, monocytes, NKTs, preadipocytes, smooth muscle cells, and MSC are risk factors, while CD8 T cells, CD8+ T cells, and plasma cells are protective factors. Moreover, we find that the immune system and organogenesis are uniformly enriched in negative immune response clusters, in contrast to the enrichment of nervous system in positive immune response clusters. Mesenchymal differentiation is also observed in the negative immune response clusters. High enrichment status of macrophages in negative immune response clusters are independently validated by analyzing scRNA-seq data from eight high-grade gliomas, revealing that negative immune response samples comprised 46.63% to 55.12% of macrophages, whereas positive immune response samples comprised only 1.70% to 8.12%, with IHC staining of samples from six short-term and six long-term survivors of GBMs confirming the results.

**Simple Summary:** The landscape of infiltrating non-tumor cells in glioblastoma niches remains unclear. In this study, we explore the enrichment status of a total of 64 non-tumor cell types predicted by applying 540 gene signatures curated from literature and normalized enrichment score (NES) across four large gene expression profiling cohorts of glioblastoma with 967 cases. Based on non-tumor cell type-based enrichment status, GBMs in each cohort are classified into positive or negative immune response clusters, showing a statistically significant different overall survival. Astrocytes, macrophages, monocytes, NKTs, preadipocytes, smooth muscle cells, and MSC are identified as risk factors, as well as protector factors of CD8 T cells, CD8+ T cells, and plasma cells. Our results also find that immune system- and organogenesis-related GO terms are uniformly enriched in negative immune response clusters, whereas positive immune response clusters are enriched with GO terms concerning the nervous system. The mesenchymal differentiation is observed in the negative immune response clusters. Particularly, the high presence of macrophages in the negative immune response clusters is further validated using scRNA-seq analysis and IHC staining of GBMs from independent cohorts.

## 1. Introduction

Gliomas originating from astrocytes, oligodendroglia, and ependymal cells account for 70% of all brain tumors [1] and are categorized into four types: grade I pilocytic astrocytoma and grade II astrocytoma are low-grade gliomas, whereas grade III anaplastic astrocytoma and grade IV glioblastoma multiform (GBM) are malignant tumors [2]. The latter two types of tumors have poor prognosis with a median survival rate of 1 year after diagnosis and a 2-year survival rate of only 12.7% to 19.8% according to the SEER database (https://seer.cancer.gov/data/).

Glioblastoma shows significant heterogeneity, making prognosis and treatment challenging. Categorization of gliomas previously focused on histological features [3]; however, characterization methods have shifted toward high-resolution molecular profiling, including identification of isocitrate dehydrogenase (IDH) mutation, co-deletion of chromosomal arms, O6-methylguanine-DNA methyltransferase (*MGMT*) promoter methylation, and miR-181d expression [4]. Additionally, new stratifications have been proposed using gene expression profiles or specific gene mutations [5,6], methylation status [7,8], and the presence of neoantigens [9]. Numerous studies have focused on interpreting the RNA-seq profiles of gliomas in an attempt to elucidate their dynamics and mechanisms, with studies on secondary glioblastoma able to distinguish comprehensive transcriptome profiling in the malignant progression of human gliomas [10] and find critical clues of *MET*-related mutations [11] and oncogenic fusions [12]. The findings of these studies have markedly advanced the investigation of gliomas and facilitated prognostic and therapeutic developments.

The complexity of glioma tumor components and the immune microenvironment has attracted significant attention in recent years, with categorizations based on molecular profiling revealing tissue similarities between proneural, proliferative, and mesenchymal-type gliomas, respectively [5]. Certain immune components, such as tumor-associated macrophages (TAMs), have been identified as regulators of the proneural-to-mesenchymal transition [13] and contributors to immunosuppression [14], thus leading to poor prognosis. However, a comprehensive characterization of non-tumor cells in the niches of primary glioblastoma has not been fully established. Investigations into the tumor components and immune microenvironment would help unravel the cross-talk between the immune system and cancer cells and allow determination of therapeutic targets for the development of novel cancer treatments.

In this study, we generated a comprehensive description of the infiltrating non-tumor cell landscape using four large-scale gene expression data cohorts of glioblastoma. This description is presented in the form of enrichment status according to the stratification of patients into two clusters (positive immune response and negative immune response) with significant differences in overall survival (OS). Additionally, we investigated the risk levels associated with immune cell types and the enrichment of Gene Ontology (GO) terms. In particular, we confirmed enrichment of a negative prognostic factor (macrophages) in scRNAseq data of high grade gliomas and in samples from GBM patients exhibiting short-term survival by immunohistochemical (IHC) staining.

## 2. Methods

### 2.1. Gene Expression and Clinical Data

Four cohorts of gene expression profiles of GBM tumor tissues were collected from Samsung Medical Center [9,15], The Cancer Genome Atlas (TCGA; RNA sequences) [16], REMBRANDT (mRNA microarray) [17], and TCGA (mRNA microarray) [18], respectively. Samples that were not diagnosed as GBM or did not include complete gene expression or clinical data were removed, resulting in 75, 152, 181, and 559 samples in Cohorts 1, 2, 3, and 4, respectively. Tumor samples were obtained from 12 glioblastomas, including from six short-term-survival and six long-term-survival patients. All research protocols and ethics comply with the Declaration of Helsinki. Sample collection and data analyses were approved by the Beijing Tiantan Hospital institutional review board (KY 2020-093-02), and written informed consent was obtained from each participant.

### 2.2. Gene Signatures of Immune Cells

Gene signatures (*n* = 540) covering 64 cell types were collected from multiple sources [19-23]. The 64 cell types were further categorized into five groups: hematopoietic stem cells (HSCs) and hematopoietic cells (lymphoid and myeloid lineage), stromal cells, and others, as shown in Supplementary Data 1A and B.

### 2.3. Generating a Normalized Enrichment Score (NES) for Estimating Cell-Enrichment Status

An NES for the Mann–Whitney–Wilcoxon gene set test was adapted to evaluate the enrichment status of cells [24]. The NES was determined as follows:

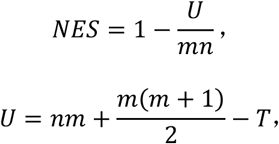

where m is the number of genes in a gene set, n is the number of genes outside the gene set, and T is the sum of the ranks of the genes in the gene set [9].

Given a gene signature, the gene expression data of a glioblastoma tumor sample were separated into two sections comprising genes expressed in the gene signature and the rest of the genes, respectively. The Wilcoxon rank-sum test was then applied to calculate the NES. In this study, the NES value was used to quantify the probability that the expression of a gene in the gene set was greater than the expression of a gene outside of that set.

### 2.4. Risk Level for Gene Signatures

Cox regression (proportional hazards regression) in the R was applied for every gene signature in each cohort. The protective factor was defined when the hazard ratio of a gene signature was <1, and the risk factor was defined when this was >1. Signatures with a p ≤ 0.05 were defined as significantly associated with survival (valid signatures), with only valid signatures used for further analysis. If all valid signatures of one cell type were either protective or risk factors, they were defined as consistent factors, otherwise, inconsistent factors.

### 2.5. Stratification of Glioblastoma Patients

Hierarchical clustering of GBMs were applied to z-score transformed NESs of these signatures using R. Euclidean distance and complete method were used for clustering, and heatmaps were drawn using the R: ‘pheatmap’. Kaplan–Meier survival analysis was performed using R: ‘survival’ and ‘survminer’.

### 2.6. GO Enrichment Analysis

Gene Set Enrichment Analysis (GSEA) [25] were performed upon negative and positive immune response clusters using a total of 6166 GO terms from the Molecular Signatures Database (MSigDB) [26], including cellular component (cc), molecular function (mf), and biological process (bp), followed by visualization through Cytoscape [27]. The results are shown in Supplementary Data 2A-D.

### 2.7. Identification of Non-Transformed Cells from ScRNA-seq Data

The single-cell gene expression data used in this analysis were accessed from the Gene Expression Omnibus (accession: GSE103224) [28]. For scRNA-seq data, genes expressed in less than or equal to 10 cells were eliminated, followed by a moving average method [29] to determine chromosome expression patterns. The number of original molecules per cell was converted to log_2_(cpm + 1). The moving average used 100 gene lengths as the window, and the value for the gene in the center of the window was considered the average expression of the window. We used the Seurat package (v.3.0) [30,31] to analyze the screened data according to standard procedures. Amplification of chromosome 7 and loss of chromosome 10 were used to differentiate malignant (transformed) cells from non-malignant (non-transformed) cells [32].

### 2.8. Determination of Non-Transformed Cell Types

scibet [33] was used to predict the identities of the non-transformed cells in the scRNA-seq data. The trained model ‘30 major human cell types’ (http://scibet.cancer-pku.cn/download_references.html), including 30 major human cell types from 42 scRNA-seq datasets, served as the reference for cell type identification.

### 2.9. Stratification of Single-cell Gene Expression Samples

To determine whether a sample in the scRNA-seq data was positive or negative immune response, Spearman correlation analysis was applied between the sample in the scRNA-seq cohort and the samples in the four gene expression profiling cohorts, respectively. Only positive correlations were retained, and the mean value of the correlation coefficients in each cohort was calculated. The fold change for a sample in the scRNA-seq data was calculated as the mean correlation coefficient of the sample in the scRNA-seq data involving samples in the positive immune response clusters divided by the mean correlation coefficients of the sample in the scRNA-seq data involving samples in the negative immune response clusters. The fold changes in the correlation coefficients calculated for the four cohorts were multiplied to determine the total fold change. A total fold change >1 indicated that the Spearman correlation coefficient was higher in the positive immune response clusters, and thus the sample in the scRNA-seq data was determined as positive immune response; otherwise, it was designated as negative immune response (Supplementary Data 3).

### 2.10. IHC Staining for Macrophage Markers

Tumor samples used for IHC staining were obtained from 12 GBMs, including six short-term-survival and six long-term-survival patients. The surgically removed tumor tissues were stored in formalin immediately after excision and embedded in paraffin within 3 days. IHC staining and image capture were performed as previously described [11]. The primary antibody for the detection of macrophage marker MS4A4A was obtained from Sigma-Aldrich (HPA029323; St. Louis, MO, USA), with staining was performed according to manufacturer instructions. The proportion of positive cells was counted using ImageJ software (v.1.52; National Institutes of Health, Bethesda, MD, USA). Clinical information and IHC staining results are summarized in Supplementary Data 4.

### 2.11. Statistical Analysis

P-values for NES distributions in negative immune response and positive immune response clusters were calculated using Student’s *t* test, and those for IHC staining percentages were generated from the Wilcoxon test. All analyses were conducted in R. P-values ≤ 0.05 were determined as statistical significance.

## 3. Results

### 3.1. Stratification of Glioblastomas Based on Cell Type-specific Enrichment Status

To determine the enrichment status of each cell type, we applied the NES algorithm using a total of 540 gene signatures covering 64 cell types (Supplementary Data 1A). The workflow for stratifying samples is shown in Figure 1A. The overview of the NES distribution landscape revealed that the enrichment status calculated from different gene signatures exhibited similar and stable trends for CD8 naїve T cells, common lymphoid progenitors (CLPs), epithelial cells, hepatocytes, HSCs, keratinocytes, lymphoid endothelial cells, neurons, natural killer T cells (NKTs), osteoblasts, sebocytes, and γΔT cells (Figures 1B and S1A).

**Figure 1.**
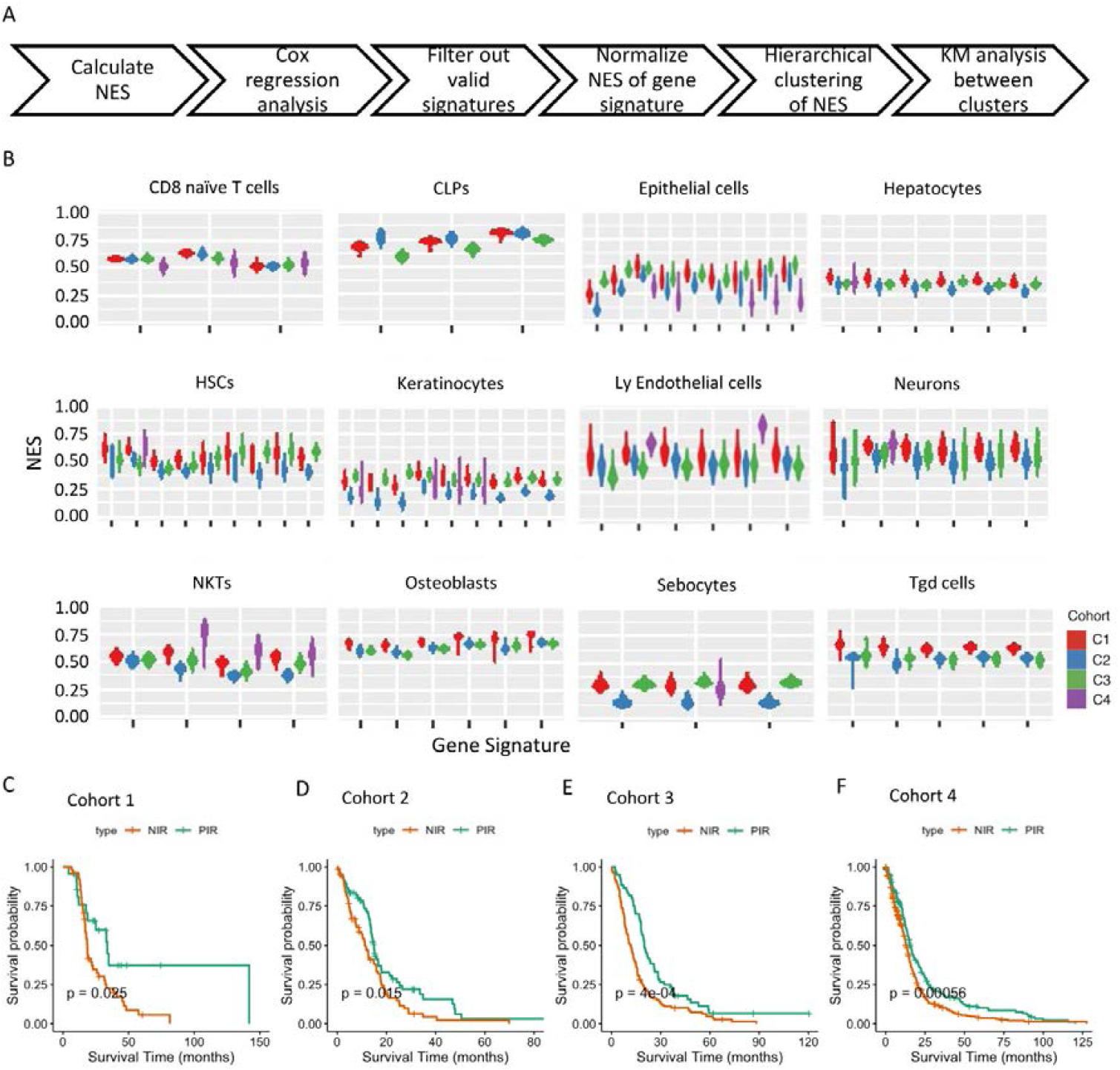
NES-based stratification of patient survival. (A) Workflow of NES-based stratification and validation of survival time. (B) NES distribution of four gene expression profiling cohorts of tumor tissues from GBM patients. Cell types and cohorts are noted. (C–F) Kaplan–Meier survival curves of the NIR and PIR clusters in the four cohorts (PIR, orange; NIR, green). NES, normalized enrichment score; NIR, negative immune response; PIR, positive immune response.

Based on the enrichment status of the gene signatures correlated with OS (valid signatures), unsupervised hierarchical clustering stratified samples into two significantly different prognostic clusters among the four cohorts (p=0.025, p=0.015, p=0.0004 and p=0.00056 for cohort 1-4, respectively) (Figures 1C–F and S1B–E) (Table 1). Clusters with patients exhibiting long- and short-term OS were designated as positive and negative immune response, respectively. We discovered that negative immune response samples were characterized by enrichment of “stromal cells”, such as skeletal muscle cells and mesenchymal stem cells (MSCs), whereas CD8 T cells were universally enriched in positive immune response samples.

**Table 1.** Hierarchical clustering results for the four cohorts.

| Cohort | Sample | Signature | Cell type | PIR | NIR | *p* | Data source |
| --- | --- | --- | --- | --- | --- | --- | --- |
| 1 | 75 | 31 | 18 | 22 | 53 | 0.02499 | Samsung Medical Center |
| 2 | 152 | 51 | 24 | 67 | 85 | 0.01462 | TCGA (RNA-seq) |
| 3 | 181 | 57 | 24 | 60 | 121 | 0.0004 | REMBRANDT (mRNA microarray) |
| 4 | 559 | 138 | 46 | 198 | 361 | 0.00056 | TCGA (mRNA microarray) |

NIR, negative immune response; PIR, positive immune response; TCGA, The Cancer Genome Atlas.

### 3.2. The Predicted Risk and Protective Landscape of Non-tumor Cells in The Glioblastoma Microenvironment

To understand the prognostic effect of different cell types, we estimated associations between the enrichment status of gene signatures and OS through Cox regression analysis across four gene expression profiling cohorts. In each cohort, statistically significant gene signatures with a hazard ratio >1 or <1 were defined as risk or protective factors, respectively. We found that risk effects consistently agreed with statistically significant gene signatures for given cell types, including activated dendritic cells (aDCs), astrocytes, class-switched memory (CSM) B cells, epithelial cells, fibroblasts, macrophages, M2 macrophages, monocytes, MSCs, NKTs, plasmacytoid (p)DCs, and preadipocytes. By contrast, CD8 naїve T cells, CD8 T cells, endothelial cells, eosinophils, megakaryocyte–erythroid progenitor cells, plasma cells, and regulatory T cells (Tregs) were consistently estimated as being protective. Additionally, basophils, B cells, CD8 central memory T cells, mast cells, multi-potent progenitor cells, memory B cells, naїve B cell, and T helper 1 (Th1) cells were predicted as being protective according to a majority of gene signatures across the four cohorts, whereas CD4 central memory T cells, chondrocytes, mesangial cells, pericytes, and smooth muscle cells were predicted as a risk by most of the gene signatures. Interestingly, the risk and protective effects of CD8 effector memory T cells, DCs, myocytes, NK cells, and skeletal muscle cells were inconsistent according to the different gene signatures (Figure 2A).

**Figure 2.**
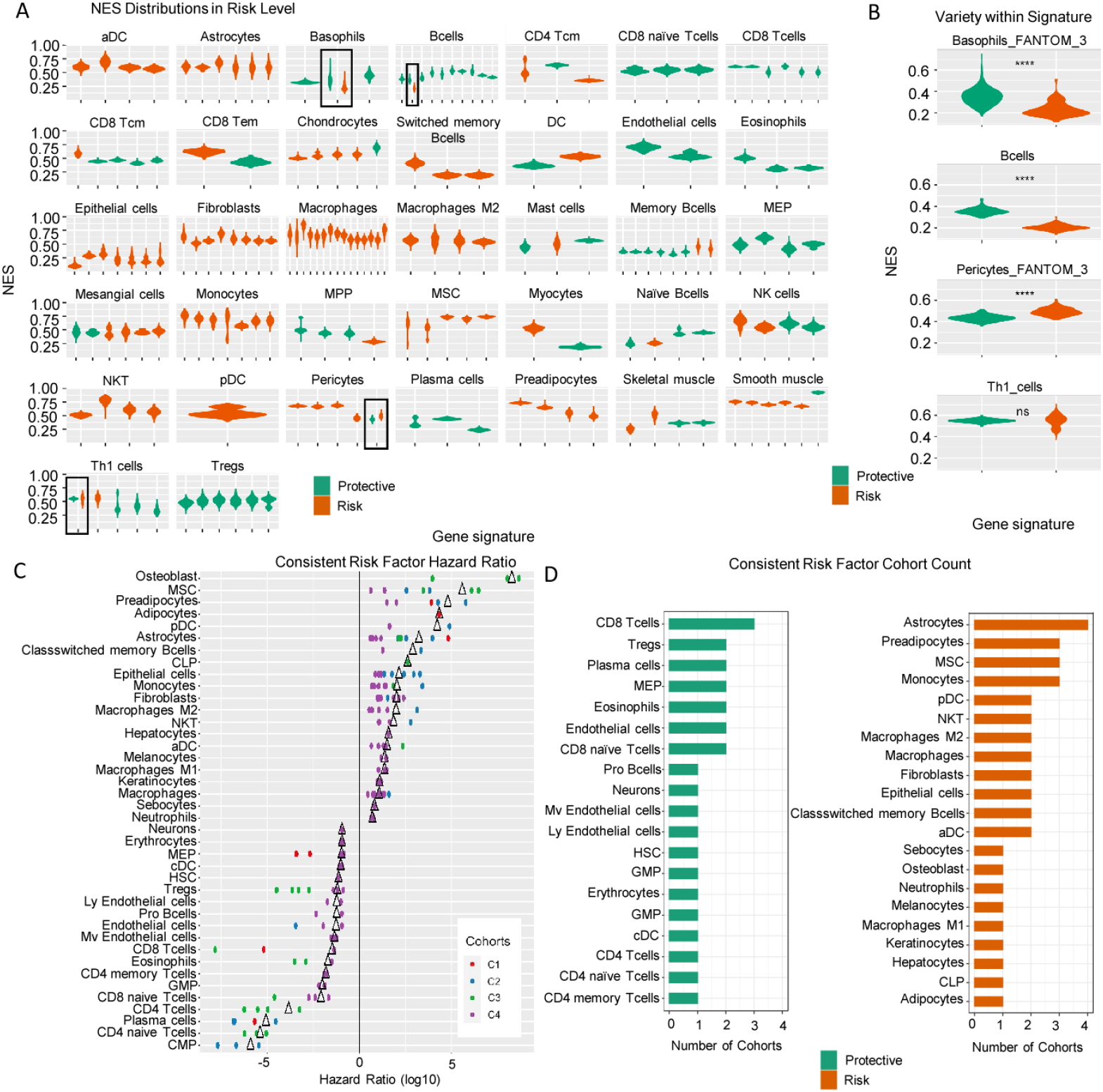
Risk levels according to calculated NESs. (A) NES distribution of valid signatures as denoted by risk levels (risk factors, orange; protective factors, green). (B) Variety of NES distribution within signatures. ns, p > 0.05; *p ≤ 0.05; **p ≤ 0.01; ***p ≤ 0.001; ****p ≤ 0.0001. (C) Hazard ratio of consistent risk factors (>1, risk factor; <1, protective factor; Δ, mean value). (D) Group count of consistent risk factors. NES, normalized enrichment score; NIR, negative immune response; ns, not significant; PIR, positive immune response.

Notably, we identified inconsistencies in some estimated risk or protective effects predicted by the gene signatures across the four cohorts. The prognostic effects of enrichment status estimated from one gene signature for basophils, B cells, pericytes, and Th1 cells were inconsistent among the four cohorts (Figure 2A and B); however, basophils, B cells, and pericytes were more likely to manifest an enrichment-dependent effect on survival time, with basophils and B cells being protective when highly enriched and pericytes presenting a risk when highly enriched.

The risk level landscape across statistically significant signatures showed consistency in risk level for certain cell types. Figure 2C shows the hazard ratios for cell types demonstrating consistent agreement in their prognostic effects across all corresponding signatures in at least two cohorts. Osteoblasts, MSCs, preadipocytes, adipocytes, pDCs, CSM B cells, and CLPs were consistent risk factors with relatively high hazard ratios in at least two cohorts. Conversely, common myeloid progenitors, CD4 naїve T cells, plasma cells, and CD4 T cells showed hazard ratios <1, suggesting potentially strong protective effects (Figure 2C). Figure 2D shows the group count of consistent risk levels. Astrocytes, preadipocytes, MSCs, monocytes, pDCs, NKTs, macrophages, M2 macrophages, fibroblasts, epithelial cells, CSM B cells, and aDCs were consistent risk factors appearing in at least two cohorts, with astrocytes being significantly negatively correlated with OS in all four cohorts. CD8 T cells, Tregs, plasma cells, MEPs, eosinophils, endothelial cells, and CD8 naїve T cells were also consistent risk factors, with CD8 T cells most frequently identified in three cohorts; however, for risk factors identified in only two cohorts (i.e., Tregs), more evidence is needed to support these findings.

### 3.3. Identification of Immune Dysregulation in the negative immune response Cluster

We then performed GSEA for the four cohorts. Enrichment map analysis of dysregulated GO terms revealed that those related to the immune system, metabolism, and organogenesis were highly enriched in all four cohorts (Figures 3A, S2A-C, and Supplementary Data 2). Specifically, GO terms related to the immune system (defense response, cytokines, myeloid lineage, and lymphoid lineage cell regulation) were enriched in negative immune response clusters, suggesting uniform dysregulation of the immune response in negative immune response clusters. Interferon (IFN)-related GO terms were significantly enriched in the negative immune response group (Figure 3B), consistent with constitutive type I IFNs (IFN-α and IFN-β) facilitating glioma-related immune escape [34], unfavorable prognosis, chemotherapy resistance, and more aggressive immune reponse [35].

**Figure 3.**
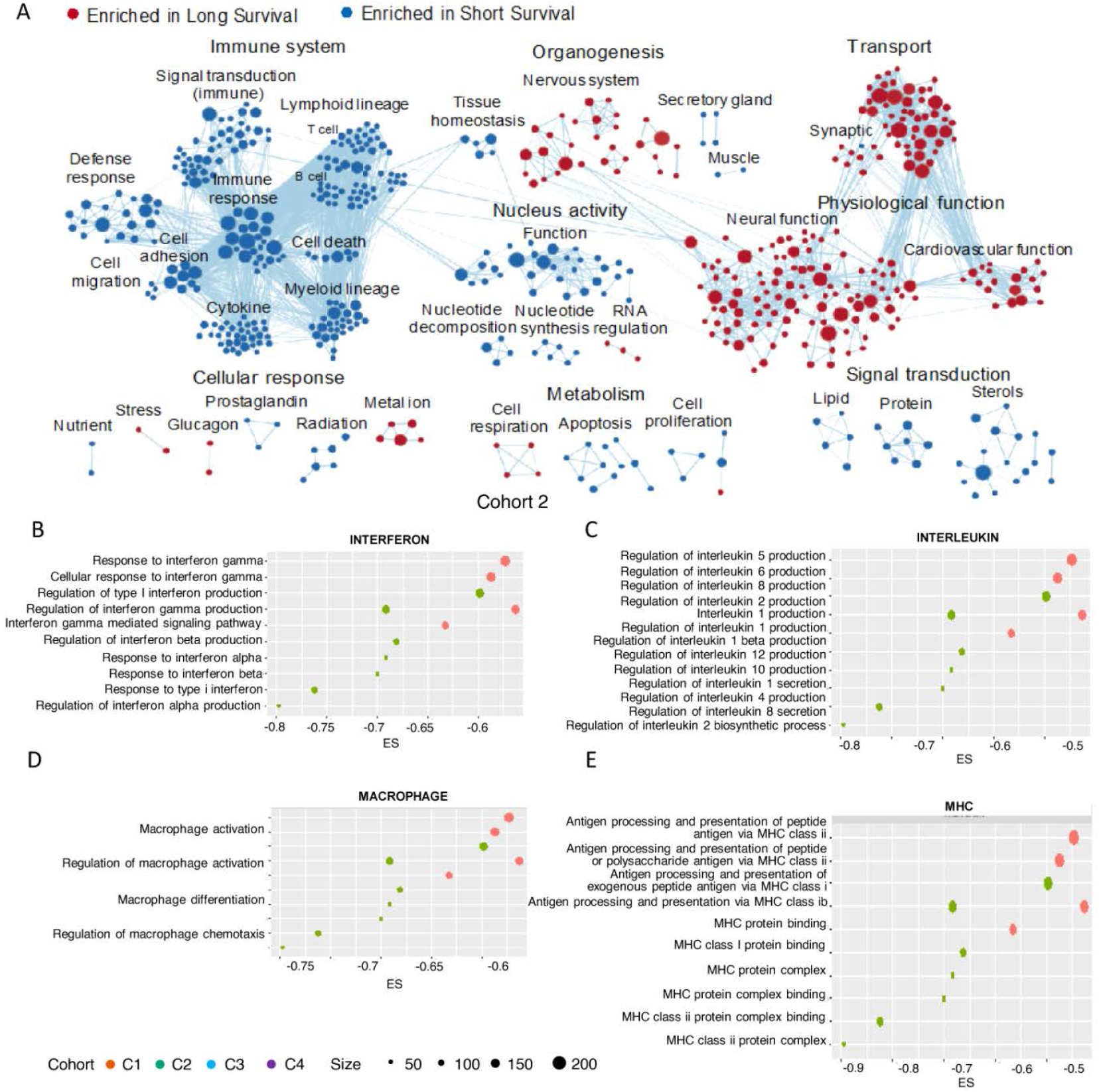
GO enrichment in clusters. Enrichment map of GO terms (selected according to p < 0.05) aggregated by functions for cohorts (A) 2. Enrichments scores for GO terms (selected according to p < 0.05) associated with the immune-related (B) interferons, (C) interleukins, (D) macrophages, and (E) MHCs. GO, Gene Ontology; MHC, major histocompatibility complex.

Activities associated with several interleukins (ILs), including IL-6, IL-8, and IL-10, were enriched in negative immune response clusters (Figure 3C), with IL-8 expression negatively correlated with GBMs survival and positively correlated with the expression of genes associated with the glioblastoma-initiating cell phenotype, as well as the possibility of GBM recurrence [36]. Additionally, IL-1β contributes to cancer cell stemness, invasiveness, and drug resistance in glioblastoma [37,38].

Moreover, we identified macrophage activation, differentiation, and chemotaxis as enriched activities in negative immune response clusters (Figure 3D), consistent with identification of macrophages as risk factors. Downregulation of major histocompatibility complex (MHC)-I and -II molecules is associated with glioma migration and invasion [39], with their altered expression associated with the negative immune response cluster (Figure 3E).

The majority of nervous system-associated GO terms (nervous system organogenesis in G1, nervous system organogenesis, neural function and synaptic in G2, and nervous system organogenesis in G4) were enriched in the positive immune response cluster (Figure 3A, S2B and S2C), demonstrating that regulation of the nervous system was a shared feature in the positive immune response cluster. This agrees with the proneural subtype of gliomas categorized by molecular profiling, in that this subtype usually demonstrated tissue similarity with adult and fetal brain and biological processes related to neurogenesis [5]. Additionally, this glioma subtype is regarded as less malignant relative to other subtypes (e.g., proliferative and mesenchymal) [5].

### 3.4. *Mesenchymal Differentiation Characterized in the* negative immune response *Cluster*

Gliomas of the mesenchymal subtype are defined by high expression of chitinase 3-like 1 and MET5, as well as a high frequency of neurofibromatosis type 1 (*NF1*) mutation/deletion and low levels of *NF1* mRNA [40]. The negative immune response clusters defined by cell-enrichment analysis shared an obvious similarity with this glioma subtype. Among the cell signatures applied to cell-enrichment analysis, 14 were categorized as stromal cells, including cells relevant to angiogenesis, muscle and bone development, and other components of connective tissue. Nine of 14 stromal cell types exhibited a significantly higher NES value in the negative immune response cluster than in the positive immune response cluster in at least three cohorts (Figure 4A–I). For the remaining five cell types, negative immune response clusters with skeletal muscle cells and endothelial cells showed higher NESs in three cohorts but distributed between two different signatures (Figure S2D and E). Lymphoid endothelial cells showed higher negative immune response-enrichment in one cohort, with no significant differences observed in other cohorts, and chondrocytes and adipocytes were more highly enriched in negative immune response clusters for two of the four cohorts. These results supported tissue similarities between negative immune response clusters and the mesenchymal subtype.

**Figure 4.**
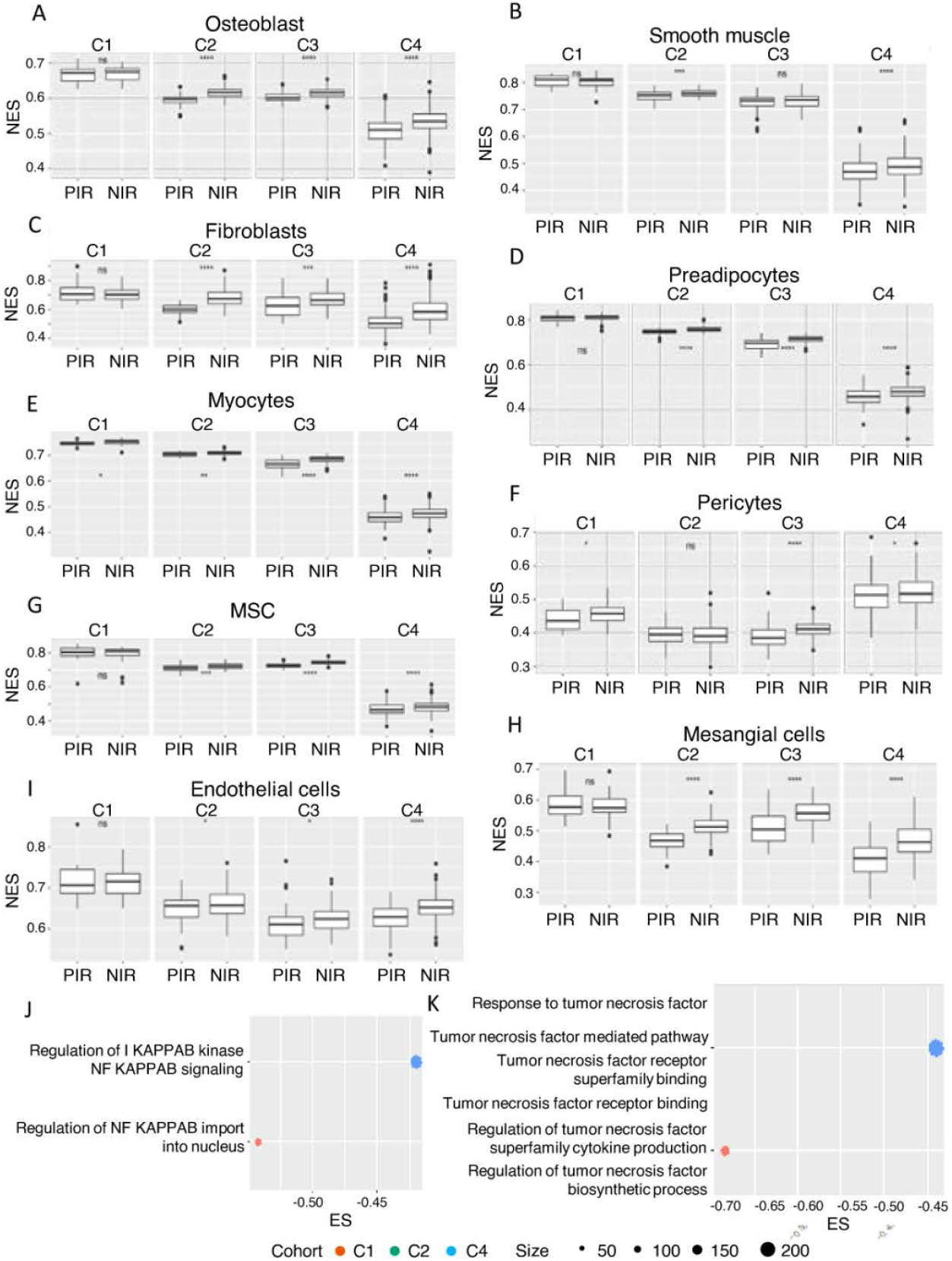
NES distributions in NIR and PIR clusters. (A) Osteoblasts, (B) smooth muscle cells, (C) fibroblasts, (D) preadipocytes, (E) myocytes, (F) pericytes, (G) MSCs, (H) mesangial cells, and (I) endothelial cells. ns, p > 0.05; *p ≤ 0.05; **p ≤ 0.01; ***p ≤ 0.001; ****p ≤ 0.0001. Enrichment scores for GO terms associated with the mesenchymal differentiation-related cytokines (selected according to a p < 0.05) (J) NF-κB and (K) TNF-α. GO, Gene Ontology; MSC, mesenchymal stem cell; NES, normalized enrichment score; NF-κB, nuclear factor-kappaB; NIR, negative immune response; PIR, positive immune response; TNF-α, tumor necrosis factor-α.

Furthermore, we identified aspects related to mesenchymal differentiation in negative immune response clusters, with enrichment of activities related to tumor necrosis factor (TNF)-α and nuclear factor-kappaB (NF-κB) identified from three cohorts and all four cohorts (Figure 4J and K), respectively. Previous studies of glioma sphere cultures indicated that TNF-α promotes mouse embryonic stem cell differentiation accompanied by increased resistance to radiotherapy in an NF-κB-dependent manner [13]. Macrophages are also an important source of TNF-α secretion.

### 3.5. ScRNA-seq and IHC Confirmation of The Negative Prognostic Effects of TAMs

To validate our findings, we collected scRNA-seq data for cell-component analysis. We classified all eight samples with available scRNA-seq data into negative or positive immune response clusters by calculating NES-based Spearman similarity between single-cell samples and bulk tumor samples (Supplementary Data 3). The results identified samples PJ016, PJ017, PJ032, and PJ048 as negative immune response and PJ018, PJ025, PJ032, and PJ035 as positive immune response.

We applied Seurat and copy number variation (CNV) analyses to distinguish non-transformed cells from malignant transformed glioma cells in the scRNA-seq data. All HGGs, except PJ016, harbored clear amplification of chromosome 7 and loss of chromosome 10 (Figure S3A–H), consistent with transformed tissues demonstrating large-scale copy number alterations and aneuploidies [41,42], as well as glioblastoma often being accompanied with amplification of chromosome 7 and loss of chromosome 10 [39]. PJ016 was found apparent loss of chromosomes 13 and 19, revealing that the cell population had indeed undergone transformation.

The identities of non-transformed cells in the glioma microenvironment were then determined using Scibet [33] (Figure 5A–H). We found no immune cells in PJ016 or PJ048 (Table 2), possibly due to the heterogeneity of different sampling areas. Those with a high percentage of macrophages (PJ017 and PJ032; 46.63% and 55.12%, respectively) belonged to the negative immune response cluster (Table 2), whereas samples with fewer macrophages (PJ018, PJ025, and PJ035; 2.28%, 1.70%, and 8.12%, respectively) overlapped with the positive immune response cluster (Table 2), confirming macrophage enrichment as a risk factor.

**Table 2.** Summary of scRNA-seq analysis.

| Patient | Age | Sex | Diagnosis | Macrophage | Microglial | All | Macrophage:All (%) | Macrophage:Microglia | Cluster |
| --- | --- | --- | --- | --- | --- | --- | --- | --- | --- |
| PJ016 | 49 | F | Glioblastoma, WHO grade IV | 0 | 0 | 3085 | 0 | — | NIR |
| PJ017 | 62 | M | Glioblastoma, WHO grade IV | 588 | 6 | 1261 | 46.63 | 98 | NIR |
| PJ018 | 65 | M | Glioblastoma, WHO grade IV | 50 | 13 | 2197 | 2.28 | 3.85 | PIR |
| PJ025 | 74 | M | Glioblastoma, WHO grade IV | 101 | 33 | 5924 | 1.70 | 3.06 | PIR |
| PJ030 | 56 | F | Anaplastic astrocytoma, WHO grade III | 258 | 108 | 3097 | 8.33 | 2.39 | PIR |
| PJ032 | 63 | F | Glioblastoma, recurrent | 759 | 22 | 1377 | 55.12 | 34.5 | NIR |
| PJ035 | 50 | M | Glioblastoma, recurrent | 306 | 232 | 3768 | 8.12 | 1.32 | PIR |
| PJ048 | 59 | M | Glioblastoma, WHO grade IV | 0 | 0 | 3084 | 0 | — | NIR |

NIR, negative immune response; PIR, positive immune response; scRNA, single-cell RNA; WHO, World Health Organization.

**Figure 5.**
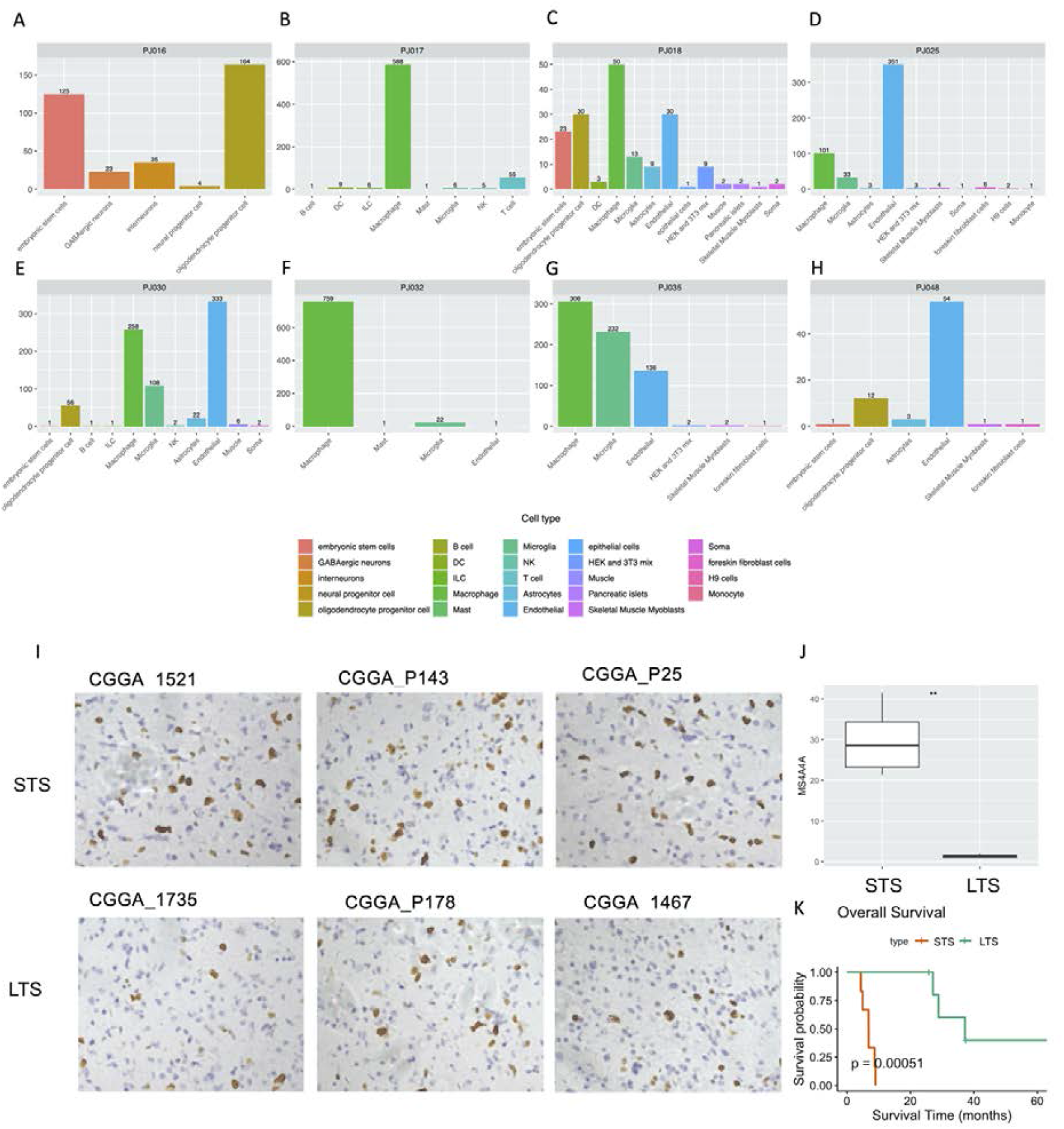
Cell type analysis using scRNA-seq data. (A–H) Cell type counts in scRNA-seq samples. (I) IHC staining of macrophages (NIR samples, upper; PIR samples, bottom). (J) Percentage of macrophages in NIR and PIR samples (according to staining for MS4A4A; scale bar: 100 μm). (K) Kaplan–Meier survival curves of NIR and PIR samples. IHC, immunohistochemical; LTS, long-term survival; NIR, negative immune response; PIR, positive immune response; scRNA, single-cell RNA; STS, short-term survival.

Moreover, we confirmed the negative prognosis associated with macrophages IHC staining for the macrophage marker MS4A4A in 12 glioblastoma samples, including six from short-term-survival and six from long-term-survival patients (Figures 5K and Supplementary Data 4). The short-term-survival samples showed a significantly higher percentage of MS4A4A-positive cells relative to the six long-term-survival samples (p = 0.00051) (Figure 5I and J).

## 4. Discussion

In this study, we generated a landscape of glioblastoma niches using four gene expression profiling cohorts of tumor tissues from GBMs based on the NES method. The patients in each cohort were divided into two categories (positive or negative immune response) according to hierarchical clustering analysis of cell type-based enrichment status and showing a significantly different survival (p < 0.05). The analysis revealed risk factors, including astrocytes, macrophages, monocytes, NKTs, preadipocytes, smooth muscle cells, and MSC, as well as protective factors, CD8 T cells, CD8+ T cells, and plasma cells. Additionally, GSEA demonstrated that immune system- and organogenesis-related GO terms were uniformly enriched in negative immune response clusters, whereas positive immune response clusters were enriched in the nervous system. Moreover, significant signs of mesenchymal differentiation were observed in the negative immune response clusters, and validation using scRNA-seq analysis and IHC staining showed correlations between the presence of macrophages and negative immune response.

Potential mechanisms associated with specific cell types manifested consistent risk levels. Some cell types exhibited identical risk levels across the four cohorts and all gene expression signatures. Specifically, astrocytes were frequently observed as a consistent risk factor with a high hazard ratio. As an important component of the blood–brain barrier and the tripartite synaptic neural network, the normal physiological role of astrocytes involves promoting mutual communication with neurons. However, astrocytes can also develop into tumor cells and form astrocytomas. Given the heterogeneity of gliomas, the high frequency of astrocytes as a risk factor is explainable. Moreover, evidence suggests that tumor-reactive astrocytes can interact with glioma tumor cells and promote the development, invasion, and survival of gliomas by releasing different cytokines or regulating the entry and exit of calcium and hydrogen ions in cell channels [43].

NKTs were also a consistent risk factor. miR-92a was reported to induce immune tolerance of NKTs to glioma cells [44]. Co-culture of glioma cells and NKTs showed miR-92a expressing in glioma cells played a key role in inducing the elevated expression of IL-6 and IL-10 in NKTs [44]. In the present study, we found IL-6- and IL-10-related GO terms in the negative immune response cluster. Compared with NKTs cultured alone, the expression of antitumor molecules, including perforin, Fas ligand, and IFN-γ, was significantly reduced in NKTs co-cultured with glioma cells [44]. Moreover, IL-6+IL-10+ NKTs exhibit a weak ability to induce apoptosis in glioma cells but have an immunosuppressive effect on CD8+ T cell activity [44].

CD8 T cells play defensive roles against cancer cells, consistent with the risk levels generated in the present analysis. Serologic analysis of antigens using recombinant cDNA expression cloning (SEREX) identified several tumor-associated antigens capable of generating a specific response in a variety of human cancers, including malignant glioma [45,46]. Tumor-related antigens can be recognized by cytotoxic CD8+ T cells in the context of tumors expressing MHC-I [47,48], suggesting that a T cell-dependent immune response might improve the outcome of glioma patients through an antigen-mediated immune response. This was supported by a clinical study of newly diagnosed glioblastoma patients that reported significantly attenuated CD8+ T cell infiltration in samples from long-survival patients (>403 days) relative to that in samples from short-survival patients (<95 days) [49]. These findings agreed with those of the present study showing that CD8+ T cells were categorized as a protective factor.

Some cell types exhibited inconsistent risk levels. In these cases, it is likely that other conditions caused a shift in risk levels (e.g., age, co-existence with other cells, or a combination of other clinical symptoms). Different signatures of the same cell type might display different risk levels, suggesting the impact of cell status. To further investigate this concept, a specific gene in each gene signature should be investigated. Other conditions, such as the presence of neoantigens [9], IDH mutation(s) [5,50], and *MGMT* methylation [8], can also provide insight into conditions causing a shift in risk levels. Furthermore, the data used in this study were from primary gliomas; therefore, comparisons between recurrent and primary glioma samples would provide additional information concerning dynamics in the glioma microenvironment.

Myeloid lineage cells, such as monocytes and macrophages, were consistent risk factors in agreement with previously reported results [51]. These cells (i.e., TAMs) account for up to 40% of the total number of solid tumor cells [52]. Numerous studies report that the frequency of TAM detection is usually higher in tumors with a mesenchymal subtype and/or recurrent tumors [53]. Glioma stem cells are recently shown to release periostin, which accumulates in the surrounding environment of blood vessels and acts as an inducer of TAM chemotaxis through signaling via the integrin receptor αvβ3 [54]. Transforming growth factor (TGF)-β released by TAMs induces matrix metalloprotein-9 expression in glioblastoma stem cells, thereby increasing their invasiveness [55]. Furthermore, the supernatant from glioma stem cells (GSCs) inhibits the phagocytic activity of TAMs and induces IL-10 and TGF-β secretion [56].

Ontogeny analysis revealed that macrophages in human GBM can be divided into either blood-derived or tissue-resident variants (i.e., microglia) [57]. These two ontogenies were also found in other types of cancer and displayed different prognostic effects. In mouse mammary carcinoma, a distinction was made between monocyte-derived TAMs and resident mammary tissue macrophages; it was found that only the former contributes to the suppression of antitumor cytotoxic T cell responses [58,59]. Normal naїve microglial cells can reduce the ability of human stem cells to acquire a spheroid morphology, thereby adversely affecting GSCs and inhibiting the growth of gliomas. However, another study suggested that microglial cells or monocytes derived from gliomas lack such antitumor potential [60]. scRNA-seq analysis of human gliomas showed that blood-derived TAMs upregulate immunosuppressive cytokines and demonstrate an altered metabolism relative to microglial TAMs and that the gene signature of blood-derived TAMs but not microglial TAMs correlates with significantly inferior survival in low-grade glioma [56]. Signatures of microglial TAMs were not included among the curated markers used for tumor tissue analysis; however, scRNA-seq analysis showed that negative immune response samples comprised a significantly higher macrophage:microglia ratio than positive immune response samples (98 vs. 34.5, respectively) (Table 2).

This study has some limitations. The stratification in this study was based on hierarchical clustering, an advantage of which is that the batch difference in each cohort is considered. However, the trade-off is that all stratifications need to be conducted in a given cohort, meaning that individual samples cannot be stratified in this way unless adopting a new categorization standard for stratification of single-cell sequencing samples. Therefore, further investigations should consider designing a supervised machine learning method for stratification. Under such circumstances, the focus should be to initially filter features for classification. In the present study, Cox regression analysis showed that not all signatures exert significant effects on survival or prognosis. Furthermore, prior to application of supervised machine learning, determination of label values will need to be undertaken.

## 5. Conclusions

In conclusion, we present a comprehensive characterization of non-tumor cells in the niches of primary glioblastoma by integrating four large cohorts of GBM gene expression data and 540 gene signatures covering 64 non-tumor cells types. We find that non-tumor cell type enrichment status are useful for stratifying glioblastomas into different prognostic groups (positive or negative immune response clusters). The negative immune response clusters are uniformly enriched with immune system- and organogenesis-related GO terms, whereas positive immune response clusters are enriched with the nervous system. The mesenchymal differentiation is also observed in the negative immune response clusters. Moreover, risk analysis using cell components to determine glioma niches help interpret the impact of cell type on cancer prognosis. Astrocytes, macrophages, monocytes, NKTs, preadipocytes, smooth muscle cells, and MSC are found as risk factors, and CD8 T cells, CD8+ T cells, and plasma cells are protective factors. Particularly, the high presence of macrophages in the negative immune response clusters is validated using scRNA-seq analysis and IHC staining of GBMs from independent cohorts. Future investigations should focus on cell types with variable risk levels in order to elucidate the potential mechanisms involved in shifts in prognostic effects. Other stratification methods should be established and evaluated for categorizing samples individually rather than as groups.

## Author Contributions

Conceptualization, Wei Zhang, Yubo Fan and Jing Zhang; Data curation, Lin Li, Ting Sun, Yufei He, Guang Liu, Lu Wang, Xiaohan Han, Hao Wen, Yong Liu, Yifan Chen, Haoyu Wang and Jing Li; Formal analysis, Zixuan Xiao, Guanzhang Li and Wendong Li; Funding acquisition, Wei Zhang, Yubo Fan and Jing Zhang; Investigation, Zixuan Xiao, Guanzhang Li and Jing Zhang; Supervision, Wei Zhang, Yubo Fan and Jing Zhang; Validation, Wei Zhang, Guanzhang Li and Wendong Li; Visualization, Zixuan Xiao and Wendong Li; Writing – original draft, Zixuan Xiao, Wei Zhang, Yubo Fan and Jing Zhang; Writing – review & editing, Zixuan Xiao, Wei Zhang, Guanzhang Li, Wendong Li, Lin Li, Ting Sun, Yufei He, Guang Liu, Lu Wang, Xiaohan Han, Hao Wen, Yong Liu, Yifan Chen, Haoyu Wang, Jing Li, Yubo Fan and Jing Zhang.

## Institutional Review Board Statement

The study was conducted according to the guidelines of the Declaration of Helsinki, and approved by the Beijing Tiantan Hospital institutional review board (KY 2020-093-02).

## Informed Consent Statement

**I**nformed consent was obtained from each patient involved in the study.

## Data Availability Statement

The data that support the findings of this study are openly available. The availability of download URL and clinical information for the four datasets were indicated in the original researches including the Samsung Medical Center^9,15^, The Cancer Genome Atlas (TCGA; RNA sequences)^16^, REMBRANDT (mRNA microarray)^17^, and TCGA (mRNA microarray)^18^. The code for calculating clustering and survival analysis, cox regression analysis, NES score, and NES distribution, were deposited at github (https://github.com/zhangjbig/xzx).

