## Supplementary Materials for "Landscape of glioblastoma niches reveals the prognostic effects of tumor-infiltrating cells"

### Description of supplementary materials

#### Supplementary Figures:

Figure S1. NES distribution and NES heatmap of gene signatures generated from hierarchical clustering.

Figure S2. NES distribution in NIR and PIR of gene signatures for Skeletal muscle and Endothelial cells in four cohorts.

Figure S3. Separation of malignant (transformed) cells from non-malignant (non-transformed) cells based on amplification of chromosome 7 and loss of chromosome 10 for eight high grade gliomas.

#### Supplementary Data

Supplementary Data 1A. Summary and categorization of 64 cell types used in this analysis.

Supplementary Data 1B. A full list of 540 gene signatures for a total of 64 cells types.

Supplementary Data 2A. The grouping of GO terms in Cohort 1.

Supplementary Data 2B. The grouping of GO terms in Cohort 2.

Supplementary Data 2C. The grouping of GO terms in Cohort 3.

Supplementary Data 2D. The grouping of GO terms in Cohort 4.

Supplementary Data 3. Classification of all eight samples with scRNA-seq data available into NIR or PIR clusters.

Supplementary Data 4. Clinical information and IHC staning results for 12 patients.

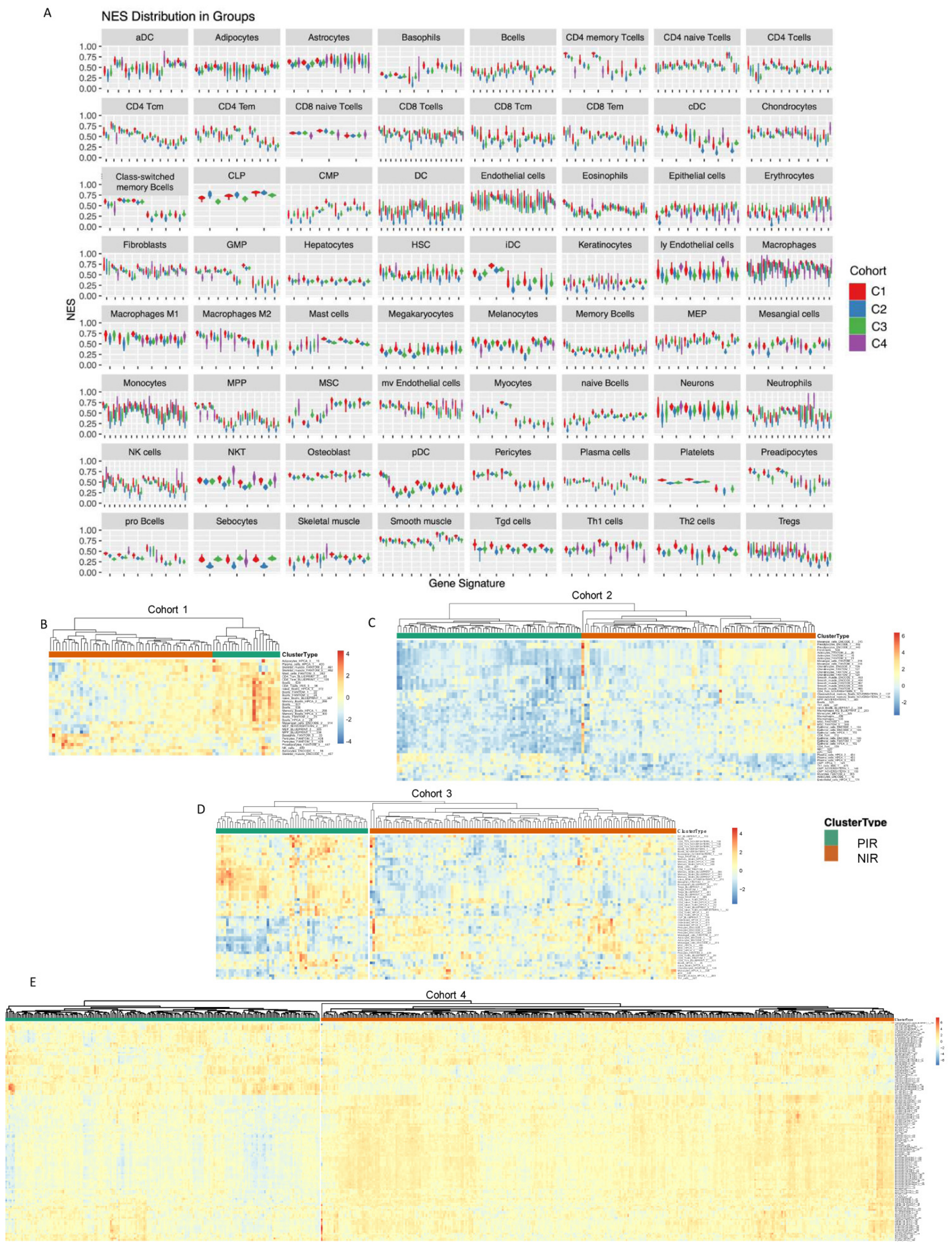


Figure S1. (A) NES distribution of four gene expression profiling cohorts of tumor tissues from GBM patients, marked in cell types and cohorts. NES heatmap of gene signatures generated from hierarchical clustering (B) Cohort 1. (C) Cohort 2. (D) Cohort 3. (E) Cohort 4. (Green: samples of PIR, orange: samples of NIR).

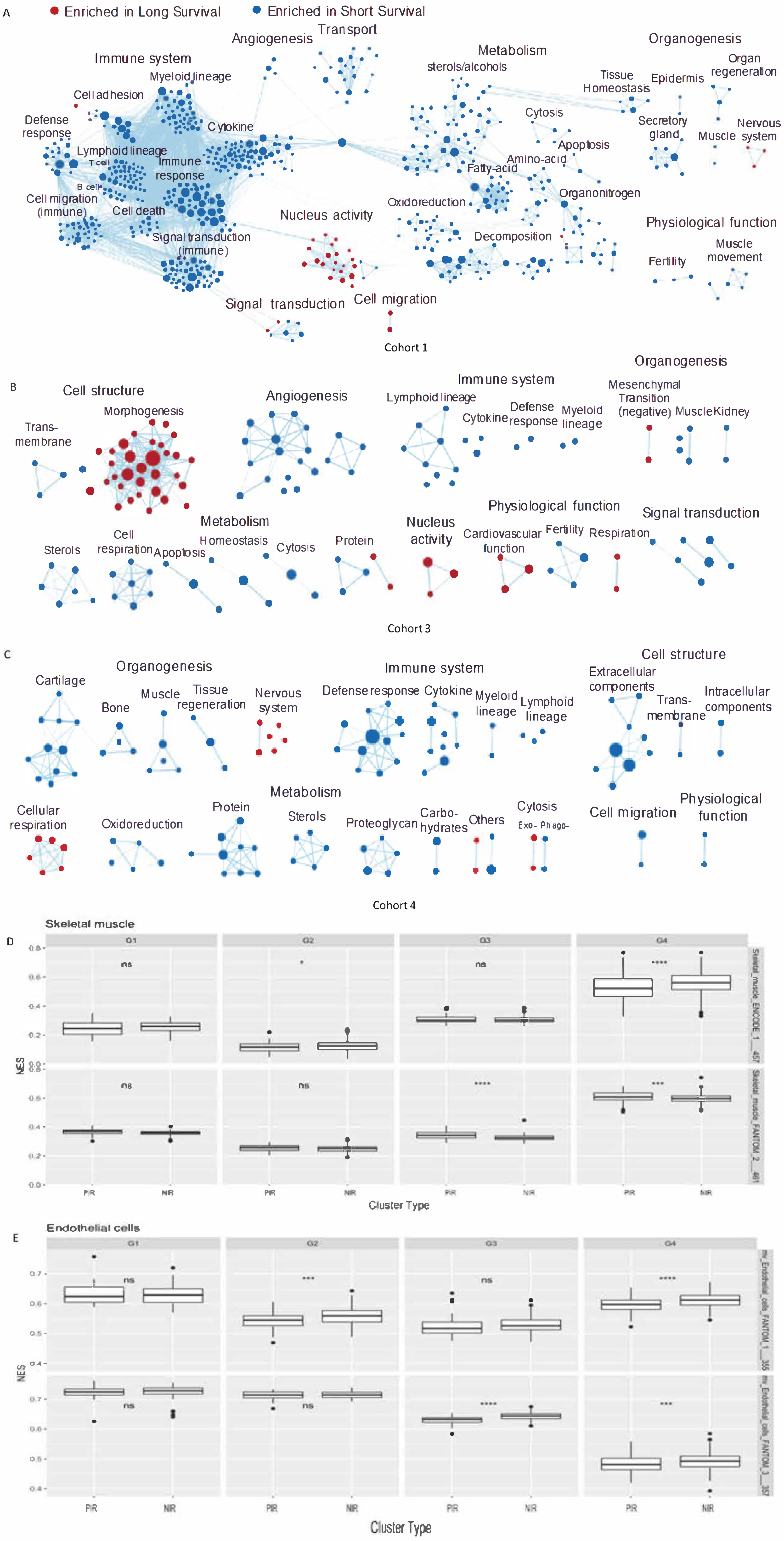


Figure S2. GO enrichment in clusters and NES distribution in NIR and PIR of gene signatures for Skeletal muscle and Endothelial cells in four cohorts. Enrichment map of GO terms (selected according to p < 0.05) aggregated by functions for cohorts (A) 1, (B) 3, and (C) 4. GO, Genen Ontology. (D) skeletal muscle and (E) endothelial cells. (ns: p > 0.05; *: p < 0.05; **: p < 0.01; ***: p < 0.001; ****: p < 0001)

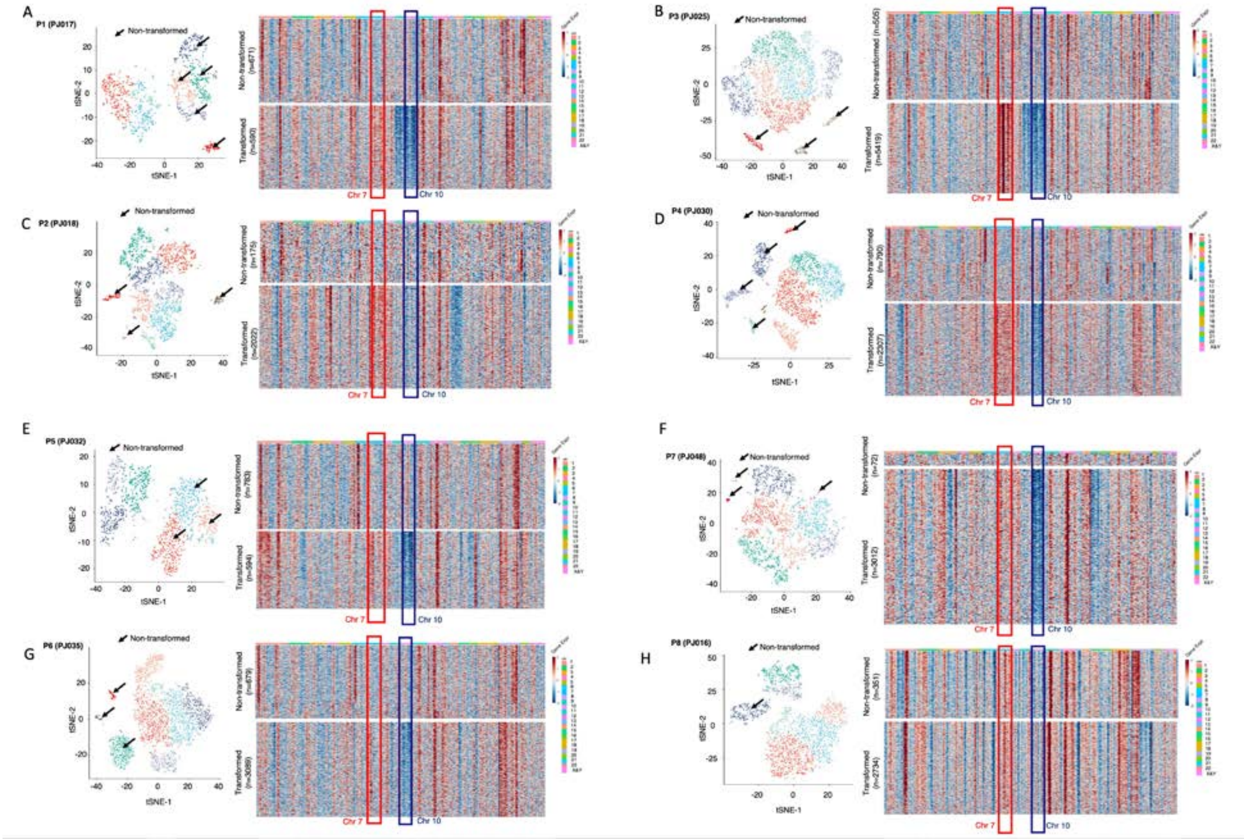


Figure S3. Separation of malignant (transformed) cells from non-malignant (non-transformed) cells based on amplification of chromosome 7 and loss of chromosome 10 for eight high grade gliomas including (A) PJ017, (B) PJ025, (C) PJ018, (D) PJ030, (E) PJ032, (F) PJ048, (G) PJ035 and (H) PJ016.

**Supplementary Data 1A. Summary and categorization of 64 cell types used in this analysis**

| Full name | Cell types | Number of Signtures | Subgroup |
|---|---|---|---|
| Epithelial cells | Epithelial cells | 9 | |
| Keratinocytes | Keratinocytes | 9 | |
| Melanocytes | Melanocytes | 7 | |
| Astrocytes | Astrocytes | 6 | |
| Neurons | Neurons | 6 | |
| Hepatocytes | Hepatocytes | 6 | |
| Sebocytes | Sebocytes | 3 | Others |
| Erythrocytes | Erythrocytes | 12 | |
| Multipotent progenitors | MPP | 12 | |
| Common myeloid progenitors | CMP | 9 | |
| Granulocyte-macrophage progenitors | GMP | 9 | |
| Megakaryocyte–erythroid progenitors | MEP | 9 | |
| Hematopoietic stem cells | HSC | 9 | |
| Megakaryocytes | Megakaryocytes | 6 | |
| Common lymphoid progenitors | CLP | 3 | |
| Platelets | Platelets | 3 | Hematopoietic stem cells and hematopoietic cells |
| CD8+ T-cells | CD8+ T-cells | 18 | Lymphoid |
| NK cells | NK cells | 18 | lineage |
| CD4+ naive T-cells | CD4+ naive T-cells | 12 | |
| B-cells | B-cells | 13 | |
| CD4+ T-cells | CD4+ T-cells | 11 | |
| CD8+ effector memory T-cells | CD8+ Tem | 10 | |
| Regulatory T-cells | Tregs | 11 | |
| Plasma cells | Plasma cells | 9 | |
| CD4+ central memory T-cells | CD4+ Tcm | 11 | |
| CD4+ effector memory T-cells | CD4+ Tem | 11 | |
| Memory B-cells | Memory B-cells | 10 | |
| CD8+ central memory T-cells | CD8+ Tcm | 10 | |
| naive B-cells | naive B-cells | 9 | |
| CD4+ memory T-cells | CD4+ memory T-cells | 6 | |
| pro B-cells | pro B-cells | 6 | |
| Class-switched memory B-cells | Class-switched memory B-cells | 6 | |
| Type 2 T-helper cells | Th2 cells | 5 | |
| Type 1 T-helper cells | Th1 cells | 5 | |
| CD8+ naive T-cells | CD8+ naive T-cells | 3 | |
| Natural killer T-cells | NKT | 4 | |
| Gamma delta T-cells | Tgd cells | 5 | |
| Monocytes | Monocytes | 16 | Myeloid |
| Macrophages | Macrophages | 16 | lineage |
| Dendritic cells | DC | 14 | |
| Neutrophils | Neutrophils | 15 | |
| Eosinophils | Eosinophils | 11 | |
| Macrophages M1 | Macrophages M1 | 6 | |
| Macrophages M2 | Macrophages M2 | 6 | |
| Activated dendritic cells | aDC | 8 | |
| Basophils | Basophils | 6 | |
| Conventional dendritic cells | cDC | 6 | |
| Plasmacytoid dendritic cells | pDC | 8 | |
| Immature dendritic cells | iDC | 5 | |
| Mast cells | Mast cells | 5 | |
| Endothelial cells | Endothelial cells | 13 | |
| Smooth muscle cells | Smooth muscle | 9 | |
| Fibroblasts | Fibroblasts | 10 | |
| Chondrocytes | Chondrocytes | 9 | |
| Adipocytes | Adipocytes | 9 | |
| Microvascular endothelial cells | mv Endothelial cells | 9 | |
| Myocytes | Myocytes | 6 | |
| Lymphatic endothelial cells | ly Endothelial cells | 6 | |
| Mesenchymal stem cells | MSC | 6 | |
| Osteoblasts | Osteoblast | 6 | |
| Preadipocytes | Preadipocytes | 6 | |
| Skeletal muscle cells | Skeletal muscle | 6 | |
| Pericytes | Pericytes | 6 | |
| Mesangial cells | Mesangial cells | 6 | Stromal cells |
| **SUM** | **64** | **540** | **5** |

**Supplementary Data 1B. A full list of 540 gene signatures for a total of 64 cells types.**

| | | |
|---|---|---|
| aDC_HPCA_1___1 | Aran_et_al_2017 | C1QA, C1QB, CD80, IL12B, CCL13, CCL17, CCL19, CCL22 |
| aDC_HPCA_2___2 | Aran_et_al_2017 | C1QA, C1QB, CD80, FPR3, HLA-DQA1, IL12B, CCL13, CCL17, CCL19, CCL22 |
| aDC_HPCA_3___3 | Aran_et_al_2017 | C1QA, C1QB, CD80, FPR3, HLA-DQA1, IL12B, CCL13, CCL17, CCL19, CCL22 |
| aDC_IRIS_1___4 | Aran_et_al_2017 | CD80, IL3RA, IL12B, CXCL9, PTGIR, CCL8, CCL13, CCL17, CCL19, CCL23, SLAMF1, SIGLEC1, TRAF1, TNFRSF4, TXN, SOCS3, HS3ST3B1, CD209, LILRA5 |
| aDC_IRIS_2___5 | Aran_et_al_2017 | ACHE, ADPRH, ALOX15B, ANXA5, XIAP, ARF3, RHOG, ATP1B3, BLVRA, C1QA, C1QB, C3AR1, CASP5, CD80, CD86, CCR5, CMKLR1, EIF5, ENO1, ETV3, FCER1G, FCER2, FPR3, GMFB, GNG5, GRB2, RAPGEF1, HCK, HLA-DQA1, HRH2, DNAJA1, IL2RA, IL3RA, IL9, IL10, IL10RA, IL12B, IMPDH1, IRF4, KCNMB1, LAIR1, LOR, RAB8A, CXCL9, MTF1, NFE2L2, NFKB1, NFKBIB, NRAS, OSM, P2RX7, PAK2, PGK1, PITPNA, MAP2K1, PTGIR, RAB5A, RELA, CLIP1, S100A10, CCL1, CCL4, CCL7, CCL8, CCL17, CCL18, CCL19, CCL22, CCL23, CCL24, SLAMF1, SLC1A2, SLC6A12, SIGLEC1, SRC, STAT2, TCF21, DYNLT1, TPI1, TRAF1, TNFRSF4, TXN, VRK2, XPNPEP1, ZBTB17, CUL1, RRP1, SCARF1, ALDH1A2, TAX1BP1, SOCS3, SPAG9, TMSB10, MAP3K13, TRIP4, H6PD, WTAP, ARHGEF11, TMCC2, HS3ST3B1, ABI1, BCL2L11, ARPC4, ARFRP1, RAMP3, LILRB2, BCKDK, CXCL13, EXOC5, MTHFD2, SEC24A, NEU3, LILRB1, CD300C, LILRB5, RAB35, OGFR, HPS5, TREX1, TFEC, RAB21, KDM6B, FAM175B, MGRN1, TDRD7, ABCA6, DNPEP, ACOT9, NUP62, BCL2L13, FBXL4, SIGLEC7, CYTH4, SIGLEC9, TOR1B, GPR132, IL19, CD209, ANKFY1, AZIN1, RIN2, TBC1D13, GPN2, C1orf27, ARL8B, ZNF654, TBC1D22B, MFN1, SCYL2, CHFR, KCNK13, C5orf15, RPGRIP1, CAMK1G, DENND1A, GPR107, HAMP, MIIP, NSUN3, MMP25, UBE2Z, TNIP2, OPA3, TMX1, SLCO5A1, NETO2, MED25, SLC25A28, ADPGK, MAGT1, DOT1L, LILRA5, TMEM41B |
| aDC_IRIS_3___6 | Aran_et_al_2017 | ACHE, ALOX15B, RHOG, ATP1B3, BLVRA, C1QB, CD80, CD86, EIF5, ENO1, ETV3, GNG5, RAPGEF1, HCK, IL2RA, IL3RA, IL9, IL10, IL10RA, IL12B, IRF4, RAB8A, CXCL9, MTF1, NFE2L2, NFKB1, P2RX7, PGK1, MAP2K1, PTGIR, RELA, CLIP1, CCL7, CCL8, CCL17, CCL19, CCL23, SLAMF1, SLC1A2, SLC6A12, SIGLEC1, SRC, STAT2, TRAF1, TNFRSF4, TXN, ZBTB17, ELL, CUL1, ALDH1A2, SOCS3, SPAG9, TMSB10, MAP3K13, WTAP, N4BP1, ARHGEF11, HS3ST3B1, ABI1, ARPC4, MTHFD2, SEC24A, NEU3, OGFR, TFEC, FAM175B, TDRD7, ACOT9, NUP62, ABTB2, TRPC4AP, FBXL4, CYTH4, TOR1B, SNX11, GPR132, IL19, CD209, CHMP5, AZIN1, C1orf27, ARL8B, ZNF654, MFN1, NECAP2, CHFR, C5orf15, CAMK1G, DENND1A, GPR107, MIIP, NSUN3, UBE2Z, TNIP2, SLCO5A1, ADPGK, DOT1L, LILRA5 |
| Adipocytes_ENCODE_1___7 | Aran_et_al_2017 | ADH1B, DLAT, GPD1, LBP, PLIN1, PPP1R1A, PPP2R1B, PTGER3, CILP, ADIPOQ, COL5A3, PNPLA2 |
| Adipocytes_ENCODE_2___8 | Aran_et_al_2017 | ADH1B, ATP5G3, DLAT, GPD1, HADHA, LBP, PLIN1, PPP1R1A, PPP2R1B, PTGER3, CILP, ADIPOQ, COL5A3, PNPLA2 |
| Adipocytes_ENCODE_3___9 | Aran_et_al_2017 | ADH1B, ATP5G3, DBI, DLAT, GPD1, LBP, PLIN1, PPP2R1B, PTGER3, CILP, ADIPOQ, COL5A3, PNPLA2 |
| Adipocytes_FANTOM_1___10 | Aran_et_al_2017 | ADH1B, ATP1A2, GPD1, HP, LBP, PLIN1, TF, ADIPOQ |
| Adipocytes_FANTOM_2___11 | Aran_et_al_2017 | ADH1B, ATP1A2, GPD1, HP, LBP, PLIN1, TF, ADIPOQ |
| Adipocytes_FANTOM_3___12 | Aran_et_al_2017 | ADH1B, ATP1A2, C6, GPD1, HP, LBP, PLIN1, TF, ADIPOQ |
| Adipocytes_HPCA_1___13 | Aran_et_al_2017 | ADH1B, ATP5G3, DLAT, GPD1, HADHA, LBP, PLIN1, PPP1R1A, PPP2R1B, PTGER3, CILP, ADIPOQ, COL5A3, PNPLA2 |
| Adipocytes_HPCA_2___14 | Aran_et_al_2017 | ADH1B, DLAT, GPD1, LBP, PLIN1, PPP1R1A, PPP2R1B, PTGER3, CILP, ADIPOQ, COL5A3, PNPLA2 |
| Adipocytes_HPCA_3___15 | Aran_et_al_2017 | ADH1B, SLC25A6, ATP5G3, DBI, DLAT, GPD1, LBP, PLIN1, PPP1R1A, PPP2R1B, PTGER3, CILP, ADIPOQ, COL5A3, ECHDC1, PNPLA2 |
| Astrocytes_ENCODE_1___16 | Aran_et_al_2017 | CNN1, COL11A1, DNA2, FABP7, GPC4, CENPI, FZD2, GLI2, GNG3, GPX7, TNC, LOXL1, LRP4, NNAT, POU3F3, PTN, PTX3, RBP1, SDC2, TFAP2C, TRO, XRCC2, ADAM12, DPF3, ST8SIA2, ZNF239, DCHS1, KIF20B, REC8, PNMA2, CAND2, FBXL7, SYT11, DAPK2, ANKRD1, NT5DC3, PNMAL1, ASAP3, DOK5, GABRQ, AGPAT4, NCAPG, FSD1, KIF18A, MXD3, PSRC1, OLFML2A |
| Astrocytes_ENCODE_2___17 | Aran_et_al_2017 | CBR3, CLCN2, CNN1, COL11A1, FABP7, GPC4, FZD2, GRM7, HIST1H1D, TNC, KIFC1, LOXL1, MCM3, POU3F3, PTN, SCN5A, SDC2, STAC, TFAP2C, TRO, XRCC2, |

| | | |
|---|---|---|
| | | ADAM12, DPF3, ST8SIA2, ZNF239, DCHS1, RECQL4, REC8, TRIOBP, CAND2, FBXL7, DAPK2, RIBC2, FAM64A, PNMAL1, ASAP3, DOK5, GABRQ, AGPAT4, NT5DC2, TMEM135, FSD1, ACSS3, ATF7IP2, TRIM45, MXD3, GINS4, PSRC1, CBX2, H2AFV, OLFML2A |
| Astrocytes_ENCODE_3___18 | Aran_et_al_2017 | CNN1, COL11A1, FABP7, FZD2, GNG3, TNC, LRP4, POU3F3, PTN, ADAM12, DPF3, DAPK2, ANKRD1, PNMAL1, ASAP3, DOK5, GABRQ, AGPAT4, PSRC1, OLFML2A |
| Astrocytes_FANTOM_1___19 | Aran_et_al_2017 | ACTA2, ACTG2, APLP1, BGN, BST1, C1S, SERPINH1, CDH6, CDH11, CHN1, CLU, CNN1, COL1A1, COL1A2, COL3A1, COL5A1, COL5A2, COL6A1, COL6A2, COL6A3, COL8A2, COL13A1, CRABP2, VCAN, DCN, DIO2, DPYSL3, DUSP2, FAP, FBLN2, FBN1, FGFR1, FGG, FLNC, FN1, FZD2, GATA6, GBP2, GEM, GNG11, CFH, HOXB2, CFI, IGFBP2, IGFBP3, IGFBP5, IGFBP6, ITGA7, AFF3, LAMA4, LIF, LOX, LOXL1, LUM, MAP1A, MATN2, MDK, MEST, MFAP4, MGP, MMP2, MN1, MYLK, NAP1L3, CLDN11, OXTR, PALM, PAX6, PCOLCE, PDGFRA, PDGFRB, SERPINF1, PLN, PPP1R3C, PTGIS, PTN, PTPRN, RARB, RARRES1, RARRES2, RGS4, CCL2, CXCL12, SFRP4, SLC14A1, SPARC, TAGLN, NR2F1, TFPI, TGM2, THY1, TIMP1, TIMP2, TNS1, TPM1, TPM2, TNFSF4, UCHL1, VCAM1, WNT5A, ADAM12, MFAP5, FZD7, RGS5, PDE5A, ADAM19, CACNA1H, DIRAS3, CRLF1, DHRS3, ADAMTS2, SEMA3E, HEPH, MRC2, GFPT2, HS3ST3A1, EDIL3, LRRC17, BTN3A3, MYL9, SPON2, POSTN, NES, GLIPR1, PADI2, FILIP1L, SYNPO, NID2, NFASC, SULF1, MXRA5, MOXD1, PCOLCE2, GREM1, ANKRD1, SRPX2, EFEMP2, CPA4, COPZ2, SLC22A17, DACT1, TRPV2, SCARA3, MXRA8, SYTL2, PNMAL1, MEG3, OLFML3, RCN3, FKBP10, CLSTN2, XYLT1, C9orf16, GLT8D2, SLC12A8, CREB3L1, CHRDL1, LYPD1, ZCCHC24 |
| Astrocytes_FANTOM_2___20 | Aran_et_al_2017 | ACTA2, ACTG2, BGN, CNN1, COL1A1, COL1A2, COL3A1, COL5A1, COL6A1, COL6A2, COL8A2, VCAN, DIO2, DPYSL3, FBLN2, FGG, FLNC, FN1, CFH, HOXB2, CFI, IGFBP2, IGFBP3, IGFBP5, LAMA4, LOXL1, LUM, MEST, MFAP4, MGP, MYLK, OXTR, PCOLCE, PDGFRA, PDGFRB, PLN, PPP1R3C, PTGIS, PTN, PTPRN, RARRES1, RARRES2, RGS4, CCL2, CXCL12, SFRP4, SLC14A1, SPARC, TAGLN, TGM2, THY1, TPM1, TPM2, TNFSF4, VCAM1, ADAM12, MFAP5, FZD7, RGS5, ADAM19, CACNA1H, DIRAS3, CRLF1, DHRS3, ADAMTS2, SEMA3E, GFPT2, LRRC17, MYL9, POSTN, GLIPR1, PADI2, NID2, SULF1, MXRA5, GREM1, ANKRD1, EFEMP2, CPA4, COPZ2, DACT1, MXRA8, OLFML3, RCN3, XYLT1, SLC12A8 |
| Astrocytes_FANTOM_3___21 | Aran_et_al_2017 | ACTA2, ACTG2, APLP1, BGN, C1S, CDH11, CLU, CNN1, COL1A1, COL1A2, COL3A1, COL5A1, COL5A2, COL6A1, COL6A2, COL6A3, COL8A2, COL13A1, VCAN, DCN, DIO2, DPYSL3, DUSP2, FAP, FBLN2, FBN1, FGG, FLNC, FN1, CFH, HOXB2, CFI, IGFBP2, IGFBP3, IGFBP5, IGFBP6, LAMA4, LOX, LOXL1, LUM, MAP1A, MDK, MEST, MFAP4, MGP, MYLK, NAP1L3, CLDN11, OXTR, PCOLCE, PDGFRA, PDGFRB, PLN, PPP1R3C, PTGIS, PTN, PTPRN, RARRES1, RARRES2, RGS4, CCL2, CXCL12, SFRP4, SLC14A1, SPARC, TAGLN, NR2F1, TGM2, THY1, TPM1, TPM2, TNFSF4, VCAM1, ADAM12, MFAP5, FZD7, RGS5, PDE5A, ADAM19, CACNA1H, DIRAS3, CRLF1, DHRS3, ADAMTS2, SEMA3E, MRC2, GFPT2, HS3ST3A1, EDIL3, LRRC17, BTN3A3, MYL9, SPON2, POSTN, NES, GLIPR1, PADI2, FILIP1L, NID2, NFASC, SULF1, MXRA5, PCOLCE2, GREM1, ANKRD1, SRPX2, EFEMP2, CPA4, COPZ2, DACT1, SCARA3, MXRA8, OLFML3, RCN3, FKBP10, CLSTN2, XYLT1, GLT8D2, SLC12A8, CREB3L1, LYPD1 |
| Bcells_FANTOM_1___22 | Aran_et_al_2017 | BLK, CD19, MS4A1, CD22, CD37, CD79A, PNOC, SNX2, MBD4, STAG3, PWP1, SP140, GGA2, STAP1, FCRL2, SMC6 |
| Bcells_FANTOM_2___23 | Aran_et_al_2017 | BLK, CD19, CD22, CD37, CD79A, PNOC, MBD4, STAG3, SP140, GGA2, FCRL2, SMC6 |
| Bcells_FANTOM_3___24 | Aran_et_al_2017 | BLK, BTK, CD19, MS4A1, CD22, CD37, CD79A, CSNK1G3, PHKB, PNOC, SNX2, TRAF3, MBD4, DEPDC5, STAG3, PWP1, SP140, GGA2, STAP1, FCRL2, SMC6 |
| Bcells_HPCA_1___25 | Aran_et_al_2017 | BLK, BTK, CD19, MS4A1, CD22, CD37, CD79A, CSNK1G3, CD180, PHKB, PNOC, POU2F1, PRKCB, SNX2, TRAF3, UBE2G1, PRDM2, MBD4, SLC24A1, DEPDC5, BCL2L11, STAG3, PRDM4, PWP1, SP140, RRAS2, GGA2, SIPA1L3, STAP1, P2RY10, CDC40, MIOS, AFTPH, DEF8, ARHGAP17, MFN1, WDR11, FCRL2, SMC6, C12orf49, PIKFYVE, JMJD1C, MCM9, EGOT |
| Bcells_HPCA_2___26 | Aran_et_al_2017 | CD19, MS4A1, CD22, CD37, CD79A, SNX2, PRDM4, GGA2, FCRL2, SMC6 |
| Bcells_HPCA_3___27 | Aran_et_al_2017 | BLK, CD19, HLA-DOA, SPIB, TNFRSF13B, STAP1, VPREB3, FCRL2 |
| Bcells_NOVERSHTERN_1___28 | Aran_et_al_2017 | ACTN2, TNFRSF17, BLK, CXCR5, CD19, MS4A1, CD22, CD37, CD53, CD72, CD79A, CD79B, CNR1, CNR2, DAXX, DNASE1, AFF2, GDI2, GPR18, HLA-DOA, HTR3A, IFNW1, IL17A, INPP5B, KCNN3, CD180, MGAT5, PAX5, PGR, PNOC, POU2F2, SPIB, SYPL1, TERT, SEC62, SLC30A4, TCL1A, RNGTT, MBD4, S1PR2, RECQL5, LY86, MYOT, TCL1B, STAG3, LSM6, SP140, IKZF3, CNOT1, SIPA1L3, KIAA1033, TNFRSF13B, STAP1, TCL6, VPREB3, ITSN2, HDAC7, ARHGAP17, QRSL1, ATF7IP, UTP6, AICDA, |

| | | |
|---|---|---|
| | | DCLRE1C, FCRL2, PIKFYVE |
| Bcells_NOVERSHTERN_2___29 | Aran_et_al_2017 | TNFRSF17, BLK, CXCR5, BMP8B, CD19, MS4A1, CD37, CD53, CD72, CD79A, CD79B, CHAD, CCR6, COL19A1, CR1, CSNK1G3, GPR25, HLA-DOA, HLA-DPB1, HSPA4, HTR3A, LY9, CD180, MGAT5, CIITA, MAP3K9, MMP17, PAX5, RPS11, RPS16, SNX2, SPIB, TROVE2, ZNF37A, ZNF154, ZNF202, ZNF208, TCL1A, AP3B1, S1PR4, BAIAP3, LY86, TCL1B, KIAA0430, DEPDC5, CEPT1, SP140, IKZF3, GGA2, NUP160, KIAA1033, TNFRSF13B, ZZZ3, STAP1, PLA2G2D, ZNF638, P2RY10, VPREB3, KCNIP2, ANKMY1, TLR7, POLR3K, QRSL1, GPRC5D, NSUN5, WDR11, ZNF701, UTP6, C5orf15, DCLRE1C, FCRL2, RIC3, SMC6, MRM1, CEACAM21, ZNF688 |
| Bcells_NOVERSHTERN_3___30 | Aran_et_al_2017 | ACTN2, TNFRSF17, BLK, CXCR5, CD19, MS4A1, CD22, CD37, CD53, CD72, CD79A, CD79B, CNR1, AFF2, GDI2, GPR18, HLA-DOA, HTR3A, IFNA2, IFNW1, IL17A, INPP5B, KCNN3, PAX5, PNOC, POU2F2, SPIB, SYPL1, TERT, SLC30A4, TCL1A, RNGTT, MBD4, SLC13A2, S1PR2, RECQL5, LY86, MYOT, TCL1B, STAG3, LSM6, SP140, IKZF3, CNOT1, STAP1, TCL6, VPREB3, ITSN2, QRSL1, ATF7IP, UTP6, AICDA, DCLRE1C, FCRL2, PIKFYVE |
| Basophils_FANTOM_1___31 | Aran_et_al_2017 | ALAS2, ARR3, ART3, C8A, CD72, DEFA5, EFNA2, GPR3, ONECUT1, HP, CXCR2, KCNA5, KCNJ13, LECT2, LEP, SMCP, MPO, MYH1, NPY5R, OMG, PRKACG, PRL, TGM3, TCL1A, AKAP4, C6orf10, FRS3, LAMB4, CLCA4, AHCTF1, BMP10, AMOTL2, ARHGAP17, TEX12, C11orf16, OTUD7B, SLC17A6, CASS4, LRTM1, HPSE2, FBRS, CXorf36, SLCO5A1, ATXN3L, GATC |
| Basophils_FANTOM_2___32 | Aran_et_al_2017 | ACTN2, ADCY2, ADH1A, ADPRH, ADRA1A, AGTR2, ALDOB, APOF, ARR3, ART1, ART3, BRDT, C8A, C9, CETP, CEACAM7, COL19A1, CPA1, CRHR2, CTRL, CYP2A7, CYP17A1, CYP19A1, CYP21A2, DAZL, DCT, DSP, EFNA2, EPHA3, F9, F13B, FCN2, GABRA1, GABRG3, GNG3, GPR3, GRB2, GRIK4, GRIN2A, GRM4, GRM7, GUCA1B, ONECUT1, HP, HTN3, HTR1D, HTR1E, HTR6, IBSP, IFNAR1, IFNW1, CXCR2, IL12RB1, INHA, IRF4, KCNA4, KCNC3, KCND3, KCNJ1, KCNJ13, KRT19, KRT33B, KRT83, KRT84, LECT2, LPO, MAGEB4, MCF2, SMCP, TRPM1, MOBP, MUC7, MYF6, MYOC, MYOD1, NKX6-1, NPY5R, NTRK1, NTRK2, PAX7, PDHA2, PLP1, PMP2, POU1F1, PPEF2, PRKACG, PRL, MAPK12, SCN7A, CCL21, SEMG2, SI, SLC5A4, SLC5A5, SLC6A1, SLC6A11, SLC10A2, SLC13A1, SSX5, TAC1, TLL2, CLEC3B, TNNT2, TNNT3, TNP2, USH2A, VPREB1, ZNF16, ZNF214, SLC30A4, FGF23, TCL1A, HIST1H2BL, UNC5C, ABCB11, ADAM7, AKAP4, ALDH1A2, PHOX2B, WASL, HTR3B, FHL5, NR1I3, NR2E3, SSX3, CACNG3, TACC3, RPP38, C6orf10, FRS3, SMR3B, ZPBP, KERA, INSL6, CYP4F8, RBPJL, CLCA4, FSTL4, CRB1, SPO11, AIPL1, SLC24A2, AHCTF1, CLEC4E, FBXO22, INVS, RBMXL2, ADAMDEC1, BMP10, PNMA3, PDZRN4, CALY, TAS2R14, PDE11A, MRPS7, MYO15A, TFDP3, SPTBN5, MS4A4A, AMOTL2, SLCO1C1, GPR85, DDX4, MAGEL2, DNAJC28, ACSM5, PDPR, HJURP, DNAH3, IL26, TEX12, C11orf16, MEPE, SLC17A6, CASS4, CD177, MYOZ1, HPSE2, NEUROD6, ELSPBP1, NPVF, FBRS, MRPS15, TRIM48, ADM2, TRPM3, ZKSCAN3, SLCO5A1, SPACA1, CCDC70, BRIP1, MYLK3, ATXN3L, ZXDB, KRT24, KHDRBS2, GATC |
| Basophils_FANTOM_3___33 | Aran_et_al_2017 | SCGB2A2, NPY5R, SSX5, NR2E3, SSX3, PCDHA2, C11orf16, TMEM212 |
| Basophils_NOVERSHTERN_1___34 | Aran_et_al_2017 | CLK1, DEFA4, GPR183, ERN1, FCN1, FGR, GRK6, GTF3C1, GZMH, GZMB, HRH2, KLRD1, LAIR2, MMP8, NKG7, PF4V1, PI4KB, PLCB2, RFX2, RGS1, RNASE2, S100A12, XCL1, VIM, VAPA, CD101, GSTO1, BCL2L11, NMUR1, GNLY, LILRA1, LILRA2, JMJD6, PADI4, TBK1, WRAP53, RETN, CYSLTR2, DENND1A, NAA16, TUBB1, POLDIP3 |
| Basophils_NOVERSHTERN_2___35 | Aran_et_al_2017 | CEACAM8, FCN1, PI4KB, NCR1, BCL2L11, NMUR1, NXT1, DERL2, WRAP53, DENND1A, POLDIP3 |
| Basophils_NOVERSHTERN_3___36 | Aran_et_al_2017 | FCN1, NCR1, BCL2L11, NMUR1, NXT1, WRAP53, DENND1A, POLDIP3 |
| CD4_memory_Tcells_FANTOM_1___37 | Aran_et_al_2017 | AAMP, ACTL6A, ADSL, AKT2, ANXA7, ARL2, ATF1, BTF3, CCNC, CD2, CD5, CD6, CD28, CDK9, CETN3, CCR4, COX7C, CSNK1A1, CSNK2A2, CTBP1, CTLA4, DAD1, DDX3X, DR1, GPR183, EIF4G2, ERH, ESD, FNTA, GABPA, GLUD1, HDAC1, HINT1, HMOX2, HNRNPH3, HNRNPU, DNAJB1, EIF3E, ITK, JAK3, KIF22, KTN1, LDHB, SH2D1A, SMAD2, MAGOH, NCL, NDUFS5, PFN1, PLP2, POLD2, PPID, PPP1CB, PPP1CC, PPP2R5D, PPP6C, PTPN11, RAD21, RANBP1, RBM3, RGS1, RPL4, RPL5, RPL8, RPL13, RPS3, RPS6, RPS19, SLAMF1, SOD1, SP3, SRP9, SURF2, THOP1, TPP2, TPT1, TSN, TSSC1, UBE2D2, UBE2D3, UBE2N, UQCRC2, CNBP, DAP3, FXR1, UXT, PKP4, API5, PRPF18, DENR, STK16, RUVBL1, SSNA1, PABPC4, SUCLG1, GALR2, DPM1, NAE1, EIF2B5, FUBP3, BUB3, WDR46, ATG5, PSMF1, BAG3, PREPL, TTC37, SAFB2, CEP57, MATR3, RANBP9, ARPC4, BCAS2, DNAJA2, MAEA, PPIH, PFDN6, HMGN4, EIF3M, ANP32B, TBL3, USP39, PTGES3, CLPX, PAPOLA, SUB1, DBF4, C11orf58, HNRNPA0, COPS5, CPSF6, NUPL2, PDCD10, CBX3, U2AF2, RRP1B, MRPS27, SMC5, METAP1, RPL13A, CD2AP, EID1, ATXN10, RPL36, AHCTF1, ZZZ3, PPA2, |

| | | |
|---|---|---|
| | | SERP1, MMADHC, PRPF19, MRPS18B, METTL5, ICOS, TRA2A, UBIAD1, GPR132, UBQLN2, GLOD4, FCF1, COPS4, RSL24D1, CDC40, EIF3L, RWDD1, TRMT112, CMPK1, ETAA1, GATAD2A, MRPL20, SMU1, IMP3, LIN7C, CDV3, CDKN2AIP, AMBRA1, OSGEP, INTS8, KBTBD4, EXOC2, PCID2, ZC3H15, UNC45A, NKRF, PCNP, THAP11, ZDHHC6, MRPL11, ACD, MRPL44, MRPS34, DDX50, SPAG16, CBLL1, RPF1, THOC7, GRPEL1, ISCA1, DOHH, C12orf29, ATPIF1 |
| CD4_memory_Tcells_FANTOM_2___38 | Aran_et_al_2017 | CD6, CD28, GPR183, EIF3E, ITK, PKP4, PREPL, DNAJA2, PTGES3, CD2AP, ICOS |
| CD4_memory_Tcells_FANTOM_3___39 | Aran_et_al_2017 | ADSL, SLC25A6, ANXA7, ATF1, BTF3, CD5, CD6, CD28, CCR4, COX7C, CSNK2A2, CTBP1, DAD1, DDX3X, DR1, GPR183, EEF2, EIF4G2, ESD, FNTA, GLUD1, HDAC1, HINT1, HNRNPH3, DNAJB1, EIF3E, ITK, KARS, KIF22, KTN1, LDHB, SH2D1A, SMAD2, NDUFS5, PFN1, PLP2, PPP1CB, PPP1CC, PPP2R5D, PPP6C, RAD21, RBM3, RNF6, RPL5, RPL8, RPS3, RPS6, RPS19, SLAMF1, SOD1, SP3, SRP9, SURF2, TSN, UBE2D2, UBE2D3, CNBP, DAP3, FXR1, UXT, PKP4, PRPF18, DENR, RUVBL1, SSNA1, PABPC4, NAE1, FUBP3, BUB3, WDR46, ATG5, PSMF1, BAG3, PREPL, TTC37, CEP57, MATR3, THRAP3, ARPC4, BCAS2, DNAJA2, HMGN4, EIF3M, ANP32B, FARS2, USP39, PTGES3, PAPOLA, SUB1, C11orf58, HNRNPA0, CBX3, RBM34, RRP1B, SMC5, METAP1, RPL13A, CD2AP, EID1, RPL36, ZZZ3, TINF2, SERP1, PRPF19, ICOS, TRA2A, UBIAD1, GPR132, UBQLN2, GLOD4, FCF1, RSL24D1, CDC40, EIF3L, RWDD1, TRMT112, CMPK1, ETAA1, GATAD2A, SLC25A38, IMP3, CDV3, CDKN2AIP, AMBRA1, EXOC2, PCID2, UNC45A, NKRF, PCNP, THAP11, ZDHHC6, MRPL11, MRPS34, DDX50, CBLL1, RPF1, THOC7, ISCA1, C12orf29 |
| CD4_memory_Tcells_IRIS_1___40 | Aran_et_al_2017 | CD3G, CD28, CD40LG, CCR4, CTLA4, GPR15, LIMS1, RBL2, DLEC1, CXCR6, PDCD10, TRAT1, ARHGAP15 |
| CD4_memory_Tcells_IRIS_2___41 | Aran_et_al_2017 | CD28, CD40LG, CCR4, CTLA4, GPR15, GZMA, GZMK, HMGB2, PDCD1, RBL2, CD226, GPR171, TRAT1, UBASH3A, ARHGAP15 |
| CD4_memory_Tcells_IRIS_3___42 | Aran_et_al_2017 | ADSL, CD28, CD40LG, CCR4, CTLA4, GZMA, GZMK, HMGB2, LIMS1, MEN1, PTPN4, ZNF236, DLEC1, CD96, HMGN4, SEC23IP, ICOS, TRAT1, AURKAIP1, ARHGAP15, RNF34 |
| CD4_naive_Tcells_FANTOM_1___43 | Aran_et_al_2017 | CD3E, CD6, CD7, CCR7, DSC1, JAK3, PLCL1, PLCG1, ZAP70, CABIN1, PLXDC1, GPSM3, ANKRD55, LIMD2, CHMP7 |
| CD4_naive_Tcells_FANTOM_2___44 | Aran_et_al_2017 | APBB1, CD3E, CD6, CD7, CD27, TNFSF8, CCR7, DSC1, GRK6, GZMM, PRMT2, IDUA, INSL3, ITK, JAK3, NUMA1, PHF1, PLCL1, PLCG1, RXRB, SELPLG, ZAP70, ZNF76, CUBN, PIP4K2B, NCK2, CDK10, TNK1, ACAP1, TSPAN32, ARFRP1, 44083, MSL3, WDR6, MLXIP, LEPROTL1, CABIN1, SH2B1, IPCEF1, KLHL3, SIT1, TRAT1, CRLF3, RAPGEF6, FAM193B, SIRPG, PACS1, PLXDC1, RNPEPL1, CREBZF, GPSM3, ANKRD55, NDFIP1, LIMD2, OBSCN, CHMP7 |
| CD4_naive_Tcells_FANTOM_3___45 | Aran_et_al_2017 | CD27, CLC, CTSW, DNAJB1, RBL2, HAUS3, ANKRD55, ZNF394, CHMP7 |
| CD4_naive_Tcells_HPCA_1___46 | Aran_et_al_2017 | CD6, CCR7, DSC1, GLG1, LY9, MAK, PLCL1, RBL2, RBMS1, ATXN7, CUL1, MTRF1, BMS1, CEPT1, AAK1, SORCS3, CLUAP1, IPCEF1, ICOS, COQ6, PHF20L1, POP5, RAPGEF6, NUDT9, UBASH3A, GIN1, SETD5, KDM3A, PLXDC1, NAA16, ANKRD55, TRAF3IP3, NDFIP1, CHMP7, TMEM30B |
| CD4_naive_Tcells_HPCA_2___47 | Aran_et_al_2017 | CCR7, FKTN, IL16, KRT2, NPAT, RBMS1, VPS52, MTRF1, CEPT1, SORCS3, UTP20, PHF20L1, POP5, RAPGEF6, RNF216, TUG1, PLXDC1, ANKRD55, TRAF3IP3, CHMP7 |
| CD4_naive_Tcells_HPCA_3___48 | Aran_et_al_2017 | CD6, CCR7, FKTN, IL16, KRT2, NPAT, PLCL1, RBMS1, VPS52, MTRF1, CEPT1, AAK1, SORCS3, UTP20, PHF20L1, POP5, RAPGEF6, GIN1, TUG1, PLXDC1, NOL9, ANKRD55, TRAF3IP3, CHMP7 |
| CD4_naive_Tcells_IRIS_1___49 | Aran_et_al_2017 | CD3G, CD4, CD5, CD7, HMOX2, NPAT, RPA3, RPLP2, RPL14, SNPH, TBC1D5, RAB3GAP1, MLH3, SIRPG, PARP11, MKL1, GIMAP6 |
| CD4_naive_Tcells_IRIS_2___50 | Aran_et_al_2017 | CD4, CD5, CD7, CD40LG, INPP4A, NPAT, TRAF1, TBC1D5, SIRPG, PARP11, MKL1, GIMAP6 |
| CD4_naive_Tcells_IRIS_3___51 | Aran_et_al_2017 | CD2, CD3G, CD247, CD4, CD5, CD7, CD27, CD40LG, CCR7, HMOX2, PRMT2, INPP4A, NPAT, RPA3, RPL38, RPLP2, RPS6, TRAF1, RPL14, ZNF264, SNPH, TBC1D5, ZBTB40, PRMT3, USP16, NUP50, RAB3GAP1, ZNF609, RPRD2, LEPROTL1, MLH3, TRAT1, FCF1, SIRPG, PARP11, PLXDC1, MKL1, COPS7B, DDX31, KRI1, DDX50, ACBD4, SLTM, RPAP2, WDR82, ZNF780B, GIMAP6 |
| CD4_naive_Tcells_NOVERSHTERN_1___52 | Aran_et_al_2017 | CASP8, CD3E, CD4, CD27, CD28, CD40LG, CDK1, CCR7, CTLA4, HMOX2, LAIR2, PLCL1, RBL2, SUPV3L1, TPP2, GRAP2, SNPH, ZBTB40, ZNF263, USP16, CD226, CA5B, ZNF609, FNBP4, NUDCD3, LEPROTL1, STAP1, SIT1, ICOS, TRAT1, PHF20L1, TEX264, SIRPG, POLR3E, PLXDC1, MKL1, DPEP2, COPS7B, GIMAP6 |

| | | |
|---|---|---|
| CD4_naive_Tcells_NOVERSHTERN_2___53 | Aran_et_al_2017 | APBB1, PRMT2, TATDN2, DIDO1, REV1, RNF216, PLXDC1, NDFIP1, CHMP7 |
| CD4_naive_Tcells_NOVERSHTERN_3___54 | Aran_et_al_2017 | CD3E, CD6, CD7, CCR7, DSC1, JAK3, PLCL1, PLCG1, ZAP70, CABIN1, PLXDC1, GPSM3, ANKRD55, LIMD2, CHMP7 |
| CD4_Tcells_BLUEPRINT_1___55 | Aran_et_al_2017 | BAD, CD2, CD3G, CD5, CD28, CD40LG, CCR4, CCR7, CTLA4, GOLGA4, HMOX2, MGAT2, NFE2L2, PLCL1, PPP2CA, SON, SUPV3L1, TRAF1, TSPYL1, FBXO21, FNBP4, NUDCD3, LEPROTL1, ICOS, TRAT1, FOXP3, SIRPG, POLR3E, USP36, DDX31, RIC3, SLTM, HIPK1, ATXN7L1 |
| CD4_Tcells_BLUEPRINT_2___56 | Aran_et_al_2017 | BAD, CD2, CD3E, CD3G, CD5, CD6, CD28, CD40LG, CCR4, CCR7, CTLA4, DNAH6, GOLGA4, HMOX2, HNRNPU, DNAJB1, MBNL1, MGAT2, NFE2L2, PLCL1, PPP2CA, SFPQ, SON, SUPV3L1, TPP2, TRAF1, TSPYL1, UBP1, ZAP70, OFD1, HERC1, RBM19, DLEC1, CCR9, FBXO21, ZCCHC11, FNBP4, NUDCD3, PHF3, LEPROTL1, LSM14A, TOR1AIP1, ICOS, IL21R, TRAT1, FOXP3, WBP11, USP47, THAP1, SIRPG, POLR3E, MKL1, USP36, RBM25, DDX31, RSRC2, ZXDC, HAUS3, RIC3, C14orf169, SLTM, ARID5B, TOE1, TUBGCP5, HIPK1, ATXN7L1, SNX19 |
| CD4_Tcells_BLUEPRINT_3___57 | Aran_et_al_2017 | BAD, CD2, CD3G, CD5, CD28, CD40LG, CCR4, CTLA4, GOLGA4, HMOX2, MGAT2, NFE2L2, PLCL1, PPP2CA, SUPV3L1, TSPYL1, NUDCD3, LEPROTL1, ICOS, TRAT1, FOXP3, WBP11, SIRPG, POLR3E, USP36, DDX31, HAUS3, SLTM, HIPK1 |
| CD4_Tcells_FANTOM_1___58 | Aran_et_al_2017 | APBB1, CD5, CD6, CD28, CD40LG, CCR7, CTLA4, ITK, PLCL1, PLCG1, PSD, SSTR3, TTN, CUBN, SNPH, SPEG, MSL3, FNBP4, KLHL3, ICOS, RAPGEF6, PLXDC1, NOL9, ANKRD55, OBSCN, CHMP7 |
| CD4_Tcells_FANTOM_2___59 | Aran_et_al_2017 | APBB1, CD28, CTLA4, ITK, PLCL1, SNPH, PPWD1, PHF3, ICOS, TRAT1, RAPGEF6, NOL9, CHMP7 |
| CD4_Tcells_FANTOM_3___60 | Aran_et_al_2017 | KRIT1, CCNT2, CD2, CD4, CD28, TNFSF8, CCR7, CCR8, DDX5, DSC1, GPR183, EZH1, FCN1, GP5, PRMT2, ITIH4, ITK, LY9, PABPC3, POU6F1, PPM1B, RBL2, RXRG, ATXN7, TPT1, NR2C1, CUBN, NCK2, CDC14A, FUBP1, UBA3, SGSM2, HMGN4, CCR9, MSL3, DIDO1, CA5B, AAK1, MORC2, SACM1L, USP33, LEPROTL1, MTO1, IPCEF1, ZBTB11, SIT1, PNMA3, UBQLN2, TRAT1, CRLF3, RAPGEF6, UBASH3A, LAX1, TUG1, PRPF38B, ASXL2, GIMAP4, SIRPG, THUMPD1, ARHGAP15, ZC3HAV1, PLXDC1, ANKRD55, ALG13, CBLL1, TRMT2B, TRIM46, FBXO11, TRAF3IP3, ZNF611, OBSCN, NFATC2IP, CHMP7, TMEM123, ZFC3H1, GIMAP6, CCR2 |
| CD4_Tcells_HPCA_1___61 | Aran_et_al_2017 | ABCD2, BAD, CD2, CD3D, CD3E, CD3G, CD5, CD7, CD27, CD28, CD40LG, CCR4, CTLA4, GOLGA4, HMOX2, NFE2L2, PLCL1, PPP2CA, SON, SUPV3L1, TNFRSF4, ARHGEF1, TOMM20, CD96, FBXO21, NUDCD3, LEPROTL1, ICOS, GPR171, TRAT1, FOXP3, SIRPG, POLR3E, USP36, ZNF335, RIC3, HIPK1 |
| CD4_Tcells_HPCA_2___62 | Aran_et_al_2017 | BAD, CD2, CD3D, CD3E, CD3G, CD5, CD6, CD7, CD27, CD28, CD40LG, CCR4, CTLA4, HMOX2, PLCL1, PPP2CA, SON, SUPV3L1, TSPYL1, ARHGEF1, CD96, NUDCD3, LEPROTL1, ICOS, GPR171, FOXP3, SIRPG, ARHGAP15, HAUS3, HIPK1 |
| CD4_Tcells_HPCA_3___63 | Aran_et_al_2017 | ABCD2, BAD, CD2, CD3D, CD3E, CD3G, CD5, CD7, CD27, CD28, CD40LG, CCR4, CTLA4, GOLGA4, HMOX2, MGAT2, PLCL1, PPP2CA, SON, SUPV3L1, TRAF1, TNFRSF4, ARHGEF1, CD96, NUDCD3, LEPROTL1, ICOS, GPR171, TRAT1, FOXP3, ZNHIT6, SIRPG, HAUS3, RIC3, HIPK1 |
| CD4_Tcm_BLUEPRINT_1___64 | Aran_et_al_2017 | ADSL, AK1, CD247, CD40LG, CSNK1D, CTLA4, DNMT1, GOLGB1, ITIH4, MAN2C1, NFRKB, PLCG1, POLR2A, PSMD2, RAD9A, SPTAN1, SUPT6H, NR2C1, TSC1, USP4, DGCR14, DHX16, RAE1, BAG3, SGSM2, HUWE1, RBM5, NXF1, SORCS3, MYO16, EDC4, MTO1, ZNF638, CNIH4, ICOS, FAM193B, FBXL8, EXOC1, KDM3A, USP36, CORO7, ANKRD55, CBLL1 |
| CD4_Tcm_BLUEPRINT_2___65 | Aran_et_al_2017 | AK1, CD2, CD247, CD40LG, CCR7, CTLA4, DNMT1, GOLGB1, ITIH4, NFRKB, PLCG1, SPTAN1, TPR, NR2C1, TSC1, DHX16, TRADD, BAG3, SGSM2, RBM5, MSL3, AAK1, SORCS3, MYO16, EDC4, CNIH4, ICOS, FAM193B, FBXL8, EXOC1, KDM3A, ARHGAP15, YLPM1, USP36, ANKRD55, WDR59, CBLL1, TRAF3IP3, 44084 |
| CD4_Tcm_BLUEPRINT_3___66 | Aran_et_al_2017 | ADSL, AK1, CD2, CD247, CD40LG, CCR7, CSNK1D, CTLA4, DNMT1, GOLGB1, ITIH4, LY9, MAN2C1, NAP1L4, NFRKB, NPAT, PLCL1, PLCG1, POLR2A, PSMD2, RAD9A, SLAMF1, SPTAN1, SUPT6H, TPR, NR2C1, TSC1, USP4, ZNF200, DGCR14, DHX16, RAE1, TRADD, EIF2B5, USP10, ITGB1BP1, BAG3, ACAP1, SGSM2, HUWE1, RBM5, NME6, NXF1, USP39, FASTK, MSL3, DIDO1, WDR6, CA5B, AAK1, TCF25, SORCS3, MYO16, MCF2L2, GGA3, SMG5, EDC4, MTO1, KBTBD2, IPCEF1, ZNF638, CNIH4, ICOS, UBQLN2, COL5A3, ANAPC5, UBASH3A, FAM193B, PIGG, OLAH, FBXL8, EXOC1, KDM3A, ARHGAP15, PCDHGA9, YLPM1, INPP5E, DDX24, USP36, CORO7, ANKRD55, WDR59, CBLL1, NUP85, TRAF3IP3, 44084, CAMSAP1 |

| | | |
|---|---|---|
| CD4_Tcm_HPCA_1___67 | Aran_et_al_2017 | BMPR1A, CD4, CD5, CD6, CD28, TNFSF8, CD40LG, CD48, CTLA4, DAB1, SARDH, DVL1, ERN1, GP5, GPR15, IDUA, ITK, KRT1, POU6F1, PSD, RPL38, SCNN1D, TCF20, TPO, TNFRSF4, XPC, NCK2, CDC14A, TRADD, SNPH, DLEC1, SPEG, HNRNPUL1, MORC2, NCDN, RRS1, GMEB2, ICOS, PNMA3, GLTSCR2, TRAT1, UBASH3A, FXYD7, FBXL8, KBTBD4, ARHGAP15, ZC3HAV1, SLC4A5, DNAI2, ANKRD55, TRIM46, OBSCN, ARID5B, TRMT61A, ZXDB, TMEM30B |
| CD4_Tcm_HPCA_2___68 | Aran_et_al_2017 | BMPR1A, CD4, CD5, CD6, CD28, TNFSF8, CD40LG, CD48, CCR4, CRY2, CTLA4, DAB1, SARDH, DVL1, ERN1, GP5, GPR15, DNAJB1, IDUA, ITK, KRT1, LTA, POU6F1, PSD, PURA, RPL38, RPS16, RPS21, SCNN1D, STK11, TCF20, TFAP4, TPO, TNFRSF4, XPC, ZRSR2, NCK2, CDC14A, TRADD, ACAP1, SNPH, JOSD1, DLEC1, SPEG, IKZF1, 44083, HNRNPUL1, MORC2, TAB2, NCDN, RRS1, GMEB2, ICOS, PNMA3, GLTSCR2, TRAT1, UBASH3A, FXYD7, SNTG2, TOMM7, FBXL8, SIRPG, CDKN2AIP, KBTBD4, ARHGAP15, ZC3HAV1, THAP11, NDRG3, SLC4A5, DNAI2, ANKRD55, TRIM46, OBSCN, ARID5B, TRMT61A, ZXDB, TMEM30B |
| CD4_Tcm_HPCA_3___69 | Aran_et_al_2017 | CD5, CD6, CD28, TNFSF8, CD40LG, CD48, CTLA4, DAB1, ERN1, GP5, GPR15, ITK, KRT1, POU6F1, PSD, RPL38, TPO, TNFRSF4, NCK2, CDC14A, TRADD, SNPH, DLEC1, MORC2, RRS1, ICOS, GLTSCR2, TRAT1, UBASH3A, FXYD7, FBXL8, ARHGAP15, DNAI2, ANKRD55, OBSCN, TRMT61A, TMEM30B |
| CD4_Tcm_NOVERSHTERN_1___70 | Aran_et_al_2017 | CD40LG, CCR4, CCR8, CTLA4, DAB1, ERN1, GPR15, DNAJB1, KRT1, POU6F1, TPO, CDC14A, TRADD, DLEC1, ICOS, TRAT1, FXYD7, FBXL8, SIRPG, SLC4A5, ANKRD55, OBSCN |
| CD4_Tcm_NOVERSHTERN_2___71 | Aran_et_al_2017 | CCR4, CCR8, CTLA4, DAB1, ERN1, GPR15, DNAJB1, KRT1, POU6F1, TPO, TRADD, DLEC1, ICOS, TRAT1, FXYD7, FBXL8, SIRPG, ANKRD55, OBSCN |
| CD4_Tcm_NOVERSHTERN_3___72 | Aran_et_al_2017 | APBB1, CD5, CD6, CD28, TNFSF8, CD40LG, CD48, CCR4, CCR6, CCR7, CCR8, CTLA4, DAB1, SARDH, DVL1, ERN1, GP5, GPR15, GPR25, DNAJB1, IL2RA, ITK, KRT1, PLCG1, POU6F1, PSD, RPL38, TPO, TNFRSF4, NCK2, CDC14A, TRADD, KALRN, SOCS3, ZFYVE9, PLCH2, SNPH, DLEC1, 44083, MORC2, RRS1, ICOS, PNMA3, GLTSCR2, COL5A3, TRAT1, UBASH3A, FXYD7, SNTG2, FBXL8, SIRPG, ARHGAP15, PLXDC1, DNAI2, ANKRD55, OBSCN, ZXDB, TMEM30B |
| CD4_Tem_BLUEPRINT_1___73 | Aran_et_al_2017 | RPN2, SLAMF1, SPTAN1, TRADD, MYO16, MCF2L2, ESYT1, TRAPPC2L, ARHGAP15, RIC8A |
| CD4_Tem_BLUEPRINT_2___74 | Aran_et_al_2017 | TNFSF8, GPR25, SELPLG, SMARCC2, LZTR1, TRADD, CCR9, SIT1, FBXL8, RNPEPL1 |
| CD4_Tem_BLUEPRINT_3___75 | Aran_et_al_2017 | CD2, CTLA4, DNAJB1, RGS1, CLIP1, SLAMF1, SPTAN1, DYNLT1, COLQ, CDC14A, TRADD, RANBP9, CCR9, MYO16, MCF2L2, ICOS, TRAT1, FBXL8, SMAP1, SAMSN1, HAUS3, 44084 |
| CD4_Tem_HPCA_1___76 | Aran_et_al_2017 | ARAF, CAPZB, CCNT1, CD2, CD3E, CD4, CD5, CD40LG, CD52, COPB1, CTLA4, DYNC1H1, E4F1, EMD, GOLGB1, GPI, CXCR3, GRM3, DNAJB1, IDE, IK, IL10RA, ITIH4, ITK, MARK3, NDUFB2, NFKB1, NFRKB, PABPC3, PGK1, PLCL1, PSMD9, RABGGTA, RBL1, RGS1, RPN2, RRM1, S100A11, SELPLG, SLAMF1, SOS1, SPTAN1, SSR2, TAF10, TBCC, TCF20, DYNLT1, TRAF1, TUFM, XPC, BRPF1, DGCR14, RBM10, COLQ, MAPKAPK5, CDC14A, MKNK1, EIF3A, EIF3G, TRADD, RIPK1, TAF1B, USP10, MTMR6, ZFYVE9, BAG3, APBA3, RNF7, SEC24C, N4BP1, MAML1, RANBP9, SLC35B1, BCAS2, SF3A1, SLC9A6, CGRRF1, DCTN6, GIPC1, 44083, CCR9, JTB, COPS6, SF3B2, CDC37, MORC2, TCF25, CNOT1, MYO16, MCF2L2, JMJD6, RRS1, ESYT1, KIAA0368, CABIN1, COG4, AHCTF1, IBTK, GORASP2, GIT1, C19orf53, ICOS, UBIAD1, COL5A3, TRAT1, GOLGA7, RWDD1, TRAPPC4, VPS54, SRRT, TRAPPC2L, DDX56, STX17, FBXL8, THUMPD1, NECAP2, ARHGAP15, ZC3HAV1, RNPEPL1, USP36, ELAC2, RIC8A, SMAP1, GPSM3, SAMSN1, NSD1, NOL6, PVRIG, TNIP2, HAUS3, FYCO1, CDC73, FBXO31, MUS81, ZNF394, SPSB3, SLC38A10, ASB6, 44084 |
| CD4_Tem_HPCA_2___77 | Aran_et_al_2017 | AIRE, CD5, CD6, CD28, TNFSF8, CD40LG, CD48, CCR4, CCR6, CCR8, CTLA4, ERN1, GPR15, GZMK, DNAJB1, IL5RA, ITGB7, KLRB1, KRT1, LTA, LTK, OSM, PDCD1, RPL38, RPS21, SPTAN1, TPO, TNFRSF4, TKTL1, HIST1H3A, NCK2, CDC14A, TRADD, DLEC1, STUB1, HMGN4, CCR9, MCF2L2, NCDN, RRS1, ESYT1, GALNT8, GPR171, GLTSCR2, TRAT1, UBASH3A, FXYD7, FBXL8, KBTBD4, ARHGAP15, SAMSN1, DNAI2, CCR2 |
| CD4_Tem_HPCA_3___78 | Aran_et_al_2017 | RPN2, SLAMF1, SPTAN1, TRADD, MYO16, MCF2L2, ESYT1, TRAPPC2L, ARHGAP15, RIC8A |
| CD4_Tem_NOVERSHTERN_1___79 | Aran_et_al_2017 | AIRE, CD5, CD6, CD28, TNFSF8, CD40LG, CD48, CCR4, CCR6, CCR8, CTLA4, ERN1, GPR15, GZMK, DNAJB1, IL5RA, ITGB7, KLRB1, KRT1, LTA, LTK, OSM, PDCD1, RPL38, RPS21, TPO, TNFRSF4, HIST1H3A, NCK2, CDC14A, TRADD, DLEC1, CCR9, MCF2L2, RRS1, GALNT8, GPR171, GLTSCR2, TRAT1, FXYD7, FBXL8, ARHGAP15, SAMSN1, CCR2 |

| | | |
|---|---|---|
| CD4_Tem_NOVERSHTERN_2___80 | Aran_et_al_2017 | AIRE, CD40LG, CD48, CCR6, CCR8, ERN1, KLRB1, LTK, TPO, CCR9, MCF2L2, GALNT8, GLTSCR2, TRAT1, CCR2 |
| CD4_Tem_NOVERSHTERN_3___81 | Aran_et_al_2017 | AIRE, CD40LG, CD48, CCR6, CCR8, ERN1, KLRB1, LTK, TPO, CCR9, MCF2L2, GALNT8, GLTSCR2, TRAT1, CCR2 |
| CD8_naive_Tcells_HPCA_1___82 | Aran_et_al_2017 | AMBN, CD8A, CD8B, CCR8, COX4I1, EEF1D, GPR15, HTR1B, SMCP, MYL1, NDUFA1, NDUFA4, NDUFS5, NKTR, OMG, PSG11, SKI, SON, TPP2, SHFM1, HIST1H3A, HIST1H4F, RNMT, BUD31, GPR52, ZNHIT3, RNF7, JOSD1, SLC17A4, BCAS2, RPP38, CGRRF1, RRH, SMR3B, JMJD6, RRP8, CA14, NGDN, C19orf53, SETD2, IL21R, MED31, SS18L2, LUZP4, CDK5RAP1, FXYD7, GDAP2, CCDC87, CWF19L1, DDX24, USP36, PCIF1, MS4A5, HAUS3, LIN28A, BTNL8, SLC35E1, MOGAT2, TNKS2, KIAA1109, GJB4, HIGD2A |
| CD8_naive_Tcells_HPCA_2___83 | Aran_et_al_2017 | CD8A, CD8B, EEF1D, GPR15, MYL1, NDUFA4, NDUFS5, PSG11, SKI, SON, HIST1H3A, BUD31, GPR52, ZNHIT3, CGRRF1, RRP8, NGDN, C19orf53, SETD2, MED31, SS18L2, CDK5RAP1, DDX24, PCIF1, MS4A5, HAUS3, LIN28A, TNKS2 |
| CD8_naive_Tcells_HPCA_3___84 | Aran_et_al_2017 | CD8A, CD8B, EEF1D, GPR15, KRT1, MYL1, NDUFA4, NDUFS5, PSG11, RFX2, SKI, HIST1H3A, CILP, GPR52, SLC17A4, JMJD6, CA14, NGDN, C19orf53, SETD2, IL21R, MED31, FXYD7, DDX24, MS4A5, HAUS3, LIN28A, TNKS2, GJB4 |
| CD8_Tcells_BLUEPRINT_1___85 | Aran_et_al_2017 | CD8A, CD8B, CD27, CCR7, DSC1, PRMT2, IL16, MMP19, NFKB1, NPAT, PCNT, PFN2, PURA, RING1, MTRF1, TSPAN32, CD96, CEPT1, MSL3, DIDO1, AAK1, RBM34, CLUAP1, CBY1, POP5, RAPGEF6, YLPM1, CRTAM, CIAPIN1, TRAF3IP3 |
| CD8_Tcells_BLUEPRINT_2___86 | Aran_et_al_2017 | CA6, CD7, CD8A, CD8B, CD27, CCR7, CTSW, DSC1, FKTN, PRMT2, IL16, MMP19, NDUFS2, NFKB1, NPAT, PCNT, PFN2, PURA, RING1, S100B, ZNF200, MYOM1, MTRF1, TSPAN32, CD96, CEPT1, SDCCAG3, MSL3, DIDO1, BTN2A1, COG2, AAK1, RBM34, CLUAP1, CBY1, UTP20, UBQLN2, POP5, RAPGEF6, DPP8, CCDC25, POLR3E, NKRF, YLPM1, CRTAM, CIAPIN1, PLXDC1, GGNBP2, WDR82, TRAF3IP3, NDFIP1, TMEM41B |
| CD8_Tcells_BLUEPRINT_3___87 | Aran_et_al_2017 | CA6, CD7, CD8A, CD8B, CD27, CCR7, CTSW, DSC1, PCNT, PFN2, MYOM1, ARHGEF1, MTRF1, MSL3, CLUAP1, POP5, CRTAM, CIAPIN1, NAA16, TRAF3IP3, NDFIP1 |
| CD8_Tcells_FANTOM_1___88 | Aran_et_al_2017 | CA6, CD7, CD8A, CD8B, CD27, CCR7, CTSW, DSC1, FKTN, PRMT2, IL16, MMP19, NDUFS2, NFKB1, NPAT, PCNT, PFN2, PURA, RING1, S100B, ZNF200, MYOM1, MTRF1, TSPAN32, CD96, CEPT1, SDCCAG3, MSL3, DIDO1, BTN2A1, COG2, AAK1, RBM34, CLUAP1, CBY1, UTP20, UBQLN2, POP5, RAPGEF6, DPP8, CCDC25, POLR3E, NKRF, YLPM1, CRTAM, CIAPIN1, PLXDC1, GGNBP2, WDR82, TRAF3IP3, NDFIP1, TMEM41B |
| CD8_Tcells_FANTOM_2___89 | Aran_et_al_2017 | CASP8, CD8A, CD8B, GZMK, PTGDR, SLC1A7, TSPAN32, KLRG1, NPRL2, GIMAP4, CRTAM, ZNF611 |
| CD8_Tcells_FANTOM_3___90 | Aran_et_al_2017 | APBB1, CA6, CD8A, CD8B, DHX15, DSC1, GZMM, HNRNPL, PRMT2, KRT2, LY9, PCNT, PLCG1, PRL, PSD, RASA2, RBL2, RPL37A, S100B, SFPQ, SSTR3, TTN, ZNF154, PRPF4B, MED17, CD96, HNRNPA0, FNBP4, LSM14A, KLHL3, ZBTB11, SHANK1, ZNF639, USP47, CRTAM, ZC3HAV1, PLXDC1, GJC2, GGNBP2, NDFIP1 |
| CD8_Tcells_HPCA_1___91 | Aran_et_al_2017 | CD8A, CD8B, GZMK, IRF3, LY9, TBCC, TSPAN32, KLRG1, DPP8, SDAD1, FTO |
| CD8_Tcells_HPCA_2___92 | Aran_et_al_2017 | CD3D, CD8A, CD8B, DSC1, GZMK, IRF3, LY9, PCNT, TBCC, RNF113A, TSPAN32, KLRG1, CD160, COG2, COPZ1, MKRN2, DPP8, SDAD1, UBE2Q1, C7orf26, FTO, EML3 |
| CD8_Tcells_HPCA_3___93 | Aran_et_al_2017 | CD8A, CD8B, CD27, CTSW, CX3CR1, EEF1D, GZMH, GZMK, DNAJB1, KLRB1, LAIR2, LY9, PTGDR, PTPN4, RBL2, FBXW4, TSPAN32, KLRG1, CD160, RWDD3, IPCEF1, TOMM7, SIRPG, NAA16, FAM134C |
| CD8_Tcells_IRIS_1___94 | Aran_et_al_2017 | CD8A, CD8B, DSC1, PCNT, PLCG1, PSD, RBL2, SSTR3, CD96, PLXDC1 |
| CD8_Tcells_IRIS_2___95 | Aran_et_al_2017 | CD8A, CD8B, CD27, DSC1, MMP19, MYOM1, MTRF1, CRTAM, CIAPIN1 |
| CD8_Tcells_IRIS_3___96 | Aran_et_al_2017 | CD8A, CD8B, CD27, DSC1, MMP19, NFKB1, PCNT, RING1, MYOM1, MTRF1, COG2, CBY1, CCDC53, NKRF, CRTAM, CIAPIN1, GGNBP2 |
| CD8_Tcells_NOVERSHTERN_1___97 | Aran_et_al_2017 | CA6, CD8A, CD8B, DHX15, DSC1, PRMT2, KRT2, LY9, PCNT, PLCG1, PRL, PSD, RASA2, RBL2, RPL37A, S100B, SSTR3, TTN, ZNF154, CD96, HNRNPA0, FNBP4, ZBTB11, SHANK1, USP47, CRTAM, ZC3HAV1, PLXDC1, GGNBP2, NDFIP1 |
| CD8_Tcells_NOVERSHTERN_2___98 | Aran_et_al_2017 | CD8A, CD8B, GZMK, IRF3, LY9, TBCC, TSPAN32, KLRG1, DPP8, SDAD1, FTO |

| | | |
|---|---|---|
| CD8_Tcells_NOVERSHTERN_3___99 | Aran_et_al_2017 | APBB1, CA6, CD8A, CD8B, DHX15, DSC1, GZMM, HNRNPL, PRMT2, KRT2, LY9, PCNT, PLCG1, PRL, PSD, RASA2, RBL2, RPL37A, S100B, SFPQ, SSTR3, TTN, ZNF154, PRPF4B, MED17, CD96, HNRNPA0, FNBP4, LSM14A, KLHL3, ZBTB11, SHANK1, ZNF639, USP47, CRTAM, ZC3HAV1, PLXDC1, GJC2, GGNBP2, NDFIP1 |
| CD8_Tcm_BLUEPRINT_1___100 | Aran_et_al_2017 | CASP8, CD27, TNFSF8, GZMK, NCK1, GPR171, CRTAM, PARP11, TMEM30B |
| CD8_Tcm_BLUEPRINT_2___101 | Aran_et_al_2017 | ADCYAP1R1, ABCD2, ALK, CASP8, CD8A, CD8B, CD27, CD28, TNFSF8, CD48, ESR2, CXCR3, GZMK, LAG3, SH2D1A, MMP11, NCK1, SLC6A7, SYN2, TPP2, DUSP11, HELZ, CD96, ZMYND11, SEC24A, ZC3H13, LEPROTL1, ADAT1, CHST5, SIT1, GPR171, HAO2, GIMAP4, PCDHA10, CRTAM, PARP11, SMAP1, VPS33A, TMEM30B, GIMAP6 |
| CD8_Tcm_BLUEPRINT_3___102 | Aran_et_al_2017 | ADCYAP1R1, CASP8, CD8B, CD27, TNFSF8, ESR2, GZMK, SH2D1A, NCK1, CD96, ZMYND11, SIT1, GPR171, CRTAM, PARP11, TMEM30B |
| CD8_Tcm_HPCA_1___103 | Aran_et_al_2017 | CASP8, CD8A, CD8B, CD27, GZMK, DNAJB1, LAG3, SH2D1A, RASA2, RGS1, DYNLT1, TPP2, CDC14A, CGRRF1, DCTN6, PPWD1, ADAT1, CRTAM, PARP11, USP36, HAUS3, NAA16, TMEM30B |
| CD8_Tcm_HPCA_2___104 | Aran_et_al_2017 | CD8A, CD8B, CTSW, GZMK, IL2RB, SH2D1A, GPR171, CRTAM, USP36, TMEM30B |
| CD8_Tcm_HPCA_3___105 | Aran_et_al_2017 | BMPR1A, C21orf2, CASP8, CD3E, CD8A, CD8B, CD27, CD28, CTSW, GZMH, GZMA, GZMK, DNAJB1, IL2RB, INPP4A, KIF2A, KLRD1, KRT1, LAG3, SH2D1A, PTPN4, RASA2, RGS1, RPS6KB1, ATXN7, DYNLT1, TPP2, ZAP70, DUSP11, CDC14A, TRADD, LRIG2, MED6, KLRG1, CD96, CGRRF1, DCTN6, ZBTB1, TNRC6B, PPWD1, ADAT1, GPR171, ATF7IP, CRTAM, PARP11, CYP20A1, USP36, HAUS3, NAA16, TNKS2, ISCA1, SFXN1, TMEM30B |
| CD8_Tcm_NOVERSHTERN_1___106 | Aran_et_al_2017 | ABCD2, CD3E, CD7, CD8A, CD8B, CXCR3, GZMK, LAG3, SH2D1A, PDCD1, PTPRCAP, S100B, XCL1, SGCD, TBCC, HIST1H4F, KLRG1, STUB1, HMGN4, AKAP3, GNLY, TRAF3IP1, GPR171, ELP3, GIMAP4, CRTAM, C21orf59, WDR18, INTS5, GIMAP6 |
| CD8_Tcm_NOVERSHTERN_2___107 | Aran_et_al_2017 | ABCD2, CD3E, CD7, CD8A, CD8B, CXCR3, GZMK, LAG3, SH2D1A, PDCD1, PTPRCAP, S100B, XCL1, SGCD, HIST1H4F, KLRG1, STUB1, AKAP3, GNLY, TRAF3IP1, GPR171, ELP3, CRTAM, INTS5 |
| CD8_Tcm_NOVERSHTERN_3___108 | Aran_et_al_2017 | CD8B, CXCR3, GZMK, PTPRCAP, SGCD, HIST1H4F, STUB1, ELP3, CRTAM, INTS5 |
| CD8_Tem_BLUEPRINT_1___109 | Aran_et_al_2017 | DHX8, GZMH, GZMK, LAG3, ZAP70, COLQ, RGS9, CXCR6, PVRIG, PYHIN1 |
| CD8_Tem_BLUEPRINT_2___110 | Aran_et_al_2017 | ABCF1, ACADVL, RHOG, SLC25A20, CAPZB, CD2, CD3D, CD3G, CD7, CD8A, CD8B, CCR5, COPB1, CTSW, CX3CR1, DAXX, DHX8, DIAPH1, DMWD, DYNC1H1, EMD, GOLGA1, GTF3C1, GYG1, GZMH, GZMA, GZMK, GZMM, HMOX2, DNAJB1, IDH3B, IFNG, IL10RA, ITGAL, KLRB1, KLRD1, LAG3, LAIR2, MARK3, MT2A, MYO1F, NDUFB1, NDUFB2, NDUFS6, NKG7, PABPC3, PCNT, PHKG2, PPP2R5C, PSMC5, PSME1, PTGDR, PTPN4, PTPRA, PZP, RGS1, RPN2, CCL4, SLAMF1, SNTB2, SSR2, STX4, TAF10, DYNLT1, TUFM, WAS, ZAP70, RNF113A, LZTR1, COLQ, DHX16, GPR65, MKNK1, B4GALT3, RIPK1, RGS9, IL18RAP, ARHGEF1, CIAO1, SEC24C, N4BP1, CROCC, MAML1, KIAA0196, SEC16A, PREB, KLRG1, SLC35B1, SF3A1, GNLY, CXCR6, SF3B2, WWP2, BTN2A1, CD160, PUF60, HMGXB3, SNRNP200, JMJD6, TMEM184B, RNF167, ERAL1, CYTH4, MRPL22, ABT1, UBN1, GPR171, TIMM22, NRBP1, COL5A3, RWDD1, TRAPPC4, SRRT, STX18, TRNAU1AP, IMP3, ABCF3, CHST12, UBE2Q1, AMBRA1, MAP7D1, EXOC2, WSB2, BIN3, CRTAM, MRPS22, RIC8A, NOL6, AHNAK, PVRIG, TNIP2, HAUS3, FYCO1, C14orf169, ZMYM1, MUS81, DEFB126, ARPC5L, ZNF394, RHOT2, SPSB3, PYHIN1, NCR3, CCDC85C |
| CD8_Tem_BLUEPRINT_3___111 | Aran_et_al_2017 | SLC25A20, CALM1, CD2, CD3D, CD3G, CD8A, CD8B, CTSW, DMWD, GYG1, GZMH, GZMA, GZMB, GZMK, GZMM, IFNG, IL2RB, ITGAL, KLRB1, KLRD1, LAG3, SH2D1A, MT2A, MYO1F, NKG7, PPP1CA, PRF1, MAPK13, PTGDR, PTPN4, PTPRA, PZP, CCL4, SLAMF1, SNTB2, STX4, COLQ, GPR65, RGS9, IL18RAP, CROCC, KLRG1, GNLY, NPRL2, CXCR6, CD160, CD300A, GPR171, TBX21, COL5A3, BIN2, SASH3, UBE2Q1, MAP7D1, CRTAM, CTDSP1, RIC8A, AHNAK, PVRIG, FYCO1, ZMYM1, CCDC130, ZNF394, PYHIN1 |
| CD8_Tem_HPCA_1___112 | Aran_et_al_2017 | ABCD2, FASLG, ATR, BMPR1A, C8G, C21orf2, CACNB1, CAPN2, CASP8, CD2, CD3D, CD3G, CD8A, CD8B, CCR5, CSNK1G2, CX3CR1, DUSP8, E4F1, FLT4, GIPR, GOLGA4, CXCR3, GTF3C1, GZMH, GZMA, GZMB, GZMK, GZMM, HLA-A, HLCS, IFNG, KLRD1, KIF22, LAG3, LTK, SH2D1A, MAN2C1, MEN1, MAP3K10, MSH3, NKG7, |

| | | |
|---|---|---|
| | | PDCD1, PLCG1, PMS1, POLG, PPP2R5C, PRF1, PRKG2, PTGDR, PTPN4, PTPRC, PTPRCAP, PURA, RBL2, RFX1, S100B, XCL1, FBXW4, SLC1A7, TBCC, TCOF1, USP1, ZAP70, ZNF79, ZNF142, ARHGEF5, GPR68, LZTR1, COLQ, GPR65, IKBKAP, CDK10, SLC25A12, RIPK1, RGS9, DDX18, PSTPIP1, ARHGEF1, GPR52, OTOF, GRAP2, CTR9, CROCC, SART3, USP34, URB2, ZBTB39, DLEC1, ACTR1B, ARFRP1, KLRG1, CD96, STUB1, NMUR1, HMGN4, CXCR6, SRCAP, CD160, CAPN10, CEP250, IKZF3, AAK1, ZBTB1, SCAP, MDN1, CSTF2T, ADAT1, MTO1, FBXO3, TRMT2A, RPUSD2, VPS4A, GPKOW, SIT1, ASTE1, ANAPC2, GPR171, PNMA3, TBX21, COL5A3, ZNF639, LIPT1, PRMT7, KLHDC4, ALKBH4, CNNM2, DPP8, C2orf42, USP47, ELP3, KLHL11, IMP3, GIMAP4, CDKN2AIP, URGCP, LTB4R2, INPP5E, PRDM8, TULP4, PLXDC1, CHD8, CREBZF, ZNF335, RNF25, PAPOLG, KRI1, AHNAK, C7orf26, PVRIG, C1orf35, FYCO1, GCC1, CORO7, GSDMD, MUS81, WDR82, SLC38A1, SPSB3, ZNF276, PWWP2A, OSBPL7, MRFAP1L1, DNAJC24, ZNF428, PYHIN1, ZNF549, GIMAP6 |
| CD8_Tem_HPCA_2___113 | Aran_et_al_2017 | ABCD2, FASLG, ATR, BMPR1A, C8G, C21orf2, CACNB1, CAPN2, CASP8, CD2, CD3D, CD3E, CD3G, CD8A, CD8B, CCR5, CTBP1, CX3CR1, DUSP8, E4F1, ELK4, ERN1, FLT4, GIPR, GOLGA4, CXCR3, GTF3C1, GZMH, GZMA, GZMB, GZMK, GZMM, HLA-A, HLCS, IFNG, IL12RB1, IRF3, ITGAL, KLRB1, KLRD1, LAG3, LTK, SH2D1A, MAN2C1, MEN1, MAP3K10, NKG7, PDCD1, PLCG1, POLG, PPP2R5C, PRF1, PTGDR, PTPN4, PTPRC, PTPRCAP, PURA, RBL2, RFX1, SBF1, XCL1, FBXW4, SLAMF1, SLC1A7, TBCC, TCOF1, UBTF, ZAP70, ZNF142, ARHGEF5, GPR68, LZTR1, COLQ, GPR65, IKBKAP, CDC14A, RIPK1, RGS9, IL18RAP, PSTPIP1, ARHGEF1, GPR52, OTOF, GRAP2, NCR1, CTR9, CROCC, URB2, ZBTB39, SGSM2, DLEC1, ACTR1B, ARFRP1, KLRG1, CD96, STUB1, NMUR1, HMGN4, STK25, GNLY, CXCR6, DIDO1, CD160, CAPN10, CEP250, IKZF3, AAK1, SACM1L, MDN1, CSTF2T, ADAT1, TRMT2A, RPUSD2, VPS4A, SIT1, ASTE1, ANAPC2, GPR171, PNMA3, TBX21, COL5A3, ZNF639, KLHDC4, TTC22, KLHL11, PANK4, GIMAP4, CDKN2AIP, RNF126, TRMU, EXOC2, LTB4R2, OTUD7B, PRDM8, TULP4, PLXDC1, CREBZF, HIVEP3, ZNF335, KRI1, PVRIG, FYCO1, GCC1, CORO7, FBXO31, GSDMD, ZNF696, MUS81, WDR82, TRAF3IP3, SLC38A1, KIAA1109, RHOT2, ANGEL2, SPSB3, ZNF276, OSBPL7, MRFAP1L1, DNAJC24, ZNF428, PYHIN1, NCR3 |
| CD8_Tem_HPCA_3___114 | Aran_et_al_2017 | ABCD2, BMPR1A, C8G, CACNB1, CASP8, CD8A, DUSP8, GZMH, IFNG, LAG3, LTK, SH2D1A, MSH3, PDCD1, PPP2R5C, PURA, S100B, XCL1, FBXW4, TBCC, TCOF1, ARHGEF5, CDK10, SLC25A12, KLRG1, HMGN4, CXCR6, CEP250, IKZF3, ADAT1, RPUSD2, ASTE1, GPR171, COL5A3, GIMAP4, PVRIG, FYCO1, GCC1, MRFAP1L1, ZNF428, GIMAP6 |
| CD8_Tem_NOVERSHTERN_1___115 | Aran_et_al_2017 | FASLG, C8G, CACNB1, CD8A, CD8B, CX3CR1, GZMH, GZMB, GZMK, IFNG, LAG3, LTK, SH2D1A, PDCD1, PRF1, PTGDR, RBL2, SLC1A7, ZAP70, KLRG1, NMUR1, CXCR6, TBX21, COL5A3, PYHIN1 |
| CD8_Tem_NOVERSHTERN_2___116 | Aran_et_al_2017 | FASLG, C8G, CD8A, CD8B, CX3CR1, GZMH, GZMB, GZMK, IFNG, PDCD1, PRF1, PTGDR, SLC1A7, ZAP70, KLRG1, NMUR1, CXCR6, TBX21, COL5A3, PYHIN1 |
| CD8_Tem_NOVERSHTERN_3___117 | Aran_et_al_2017 | FASLG, C8G, CACNB1, CD8A, CD8B, CX3CR1, GZMH, GZMB, GZMK, IFNG, SH2D1A, PDCD1, PRF1, PTGDR, SLC1A7, ZAP70, KLRG1, NMUR1, CXCR6, TBX21, COL5A3, PYHIN1 |
| cDC_HPCA_1___118 | Aran_et_al_2017 | ALCAM, ANXA1, CD1C, CD1E, DBI, FCER1A, ITGAX, PITPNA, SSR1, RAB7A, CLEC10A, TCTN3, CCDC88A, SLAMF8 |
| cDC_HPCA_2___119 | Aran_et_al_2017 | ALCAM, CD1C, CD1E, DBI, FCER1A, ITGAX, SSR1, RAB7A, ACTR3, CLEC10A, CCDC88A, SLAMF8 |
| cDC_HPCA_3___120 | Aran_et_al_2017 | CD1C, CD86, FCER1A, FLT3, S100A10, CD163, CLEC10A, FGL2, CD93, WDFY3, CLEC4A |
| cDC_NOVERSHTERN_1___121 | Aran_et_al_2017 | CD1A, CD1B, CD1C, CD1E, CD80, DNASE1L3, FCER1A, GFRA2, CCL17, CCL24, RRP1B, CD209, KCNK13 |
| cDC_NOVERSHTERN_2___122 | Aran_et_al_2017 | CD1A, CD1B, CD1C, CD1E, CD80, DNASE1L3, FCER1A, GFRA2, CCL17, CCL24, CD209, KCNK13 |
| cDC_NOVERSHTERN_3___123 | Aran_et_al_2017 | ALOX15, CD1A, CD1B, CD1C, CD1E, CD80, CD86, CRH, DNASE1L3, FCER1A, GFRA2, CCL13, CCL17, CCL23, CCL24, ALDH1A2, CLEC10A, RRP1B, CD209, KCNK13 |
| Chondrocytes_ENCODE_1___124 | Aran_et_al_2017 | ACAN, ARL1, COMP, ERG, ISLR, OGN, PGK1, PRELP, COL14A1, RGS11, TBX4, MORF4L2, LAPTM4A, SNUPN, MPHOSPH6, SCRG1, FBXW11, REM1, ZNF471, PODNL1 |
| Chondrocytes_ENCODE_2___125 | Aran_et_al_2017 | ACAN, ARL1, COMP, ERG, GLG1, ISLR, OGN, PGK1, PRELP, COL14A1, PTP4A2, RGS11, TBX4, GOSR2, MORF4L2, LAPTM4A, SNUPN, MPHOSPH6, SCRG1, FBXW11, REM1, ZNF471, PODNL1 |

| | | |
|---|---|---|
| Chondrocytes_ENCODE_3___126 | Aran_et_al_2017 | ACAN, CCNB1, COL10A1, COMP, CSNK1A1, ERG, FOXD2, LPAR4, ISLR, MYOC, NFATC4, NKX3-1, OMD, PRELP, STAT2, TNXB, COL14A1, PTP4A2, GDF5, FZD9, CILP, TNFSF11, WASL, TBX4, TMEM59, MORF4L2, LAPTM4A, SNUPN, MPHOSPH6, PRG4, ANGPTL7, SLC38A3, DSTN, WWP2, SCRG1, IQSEC2, FBXW11, HSPB7, REM1, COL5A3, CRTAC1, NPLOC4, ZNF471, PODNL1, NDFIP1, DYNLRB1, LRRC15 |
| Chondrocytes_FANTOM_1___127 | Aran_et_al_2017 | ADRA1D, ACAN, AK1, ABCD1, APOC3, ARCN1, ARL1, ARNT, CACNA1C, CACNB1, CAMLG, RUNX2, RUNX1, COL10A1, COMP, COPA, COPB1, CYP19A1, DMWD, DPT, DVL1, ELN, ENO1, ERF, ETF1, FGF7, FOXC2, FSHB, GOLGA4, MCHR1, GRIA3, HAS1, HDLBP, HOXD3, IBSP, IDUA, IFNB1, IGF1, ISLR, KRT10, LEP, LTBR, SMAD5, MIF, MLN, MLLT1, NFATC4, NFKBIL1, OCRL, OMD, OGN, P4HB, PCDHGC3, PDE3A, PFN2, PGM1, PRELP, PRL, PYY, PTH1R, PTPN11, RAB3A, RGR, ROS1, S100A6, SGCD, SNTB2, SSX1, TACR3, TF, CLEC3B, TNNT3, TUB, COL14A1, ZFPL1, ZBTB16, CDK2AP1, GDF5, CILP, PRKRA, MYH13, WISP1, SPAG9, MAP3K6, OTOF, CABP1, TMEM59, BAG3, NCOR2, TTC37, LAPTM4A, CUL7, PJA2, NAALADL1, RANBP9, ABCC9, ABI2, PRG4, ANGPTL7, CALCOCO2, YAP1, CDIPT, TM9SF1, PRDX4, TRIM3, CORIN, ZMYND11, NCKAP1, CCL27, YIF1A, COPS8, SPIN1, KDELR1, SERINC3, OS9, TMED10, EMILIN1, MAP4K5, NXPH3, SCRG1, FNDC3A, TBC1D9B, ERC1, MAST2, SNX13, PHLDB1, GANAB, SCFD1, POFUT2, KIAA0368, MKRN2, EID1, BCL2L13, TMEM59L, SNED1, GORASP2, SLC13A4, RNF11, HSPB7, TUBG2, IL17B, KLF15, CNIH4, GMPPA, SEC61A1, POMT2, NRBF2, COL5A3, TMED7, CELA2B, NGRN, CMPK1, MIOS, MIER2, SLC41A3, WDR41, KLHL26, OLAH, SPATA7, GLT8D1, IRGC, IFT46, OTUD7B, ZNF471, RIC8A, ZFAND3, PKNOX2, NEUROG2, ACBD3, TTC23, SLC26A10, C11orf95, YIPF2, FTO, PODNL1, HDAC11, TCEAL4, ADM2, SVEP1, UBXN6, TM2D1, TSSK1B, IL17RC, TBC1D16, LRRC15, PRRC1, ZNF358, 44084, SLC5A12, NPHP4, PPIL6, DPY19L4, CTRB2 |
| Chondrocytes_FANTOM_2___128 | Aran_et_al_2017 | ADRA1D, ACAN, AK1, APOC3, ARNT, CACNA1C, CAMLG, RUNX2, RUNX1, COL10A1, COMP, COPA, CYP19A1, DPT, DVL1, ELN, ERF, FGF7, GRIA3, HAS1, HDLBP, IBSP, IFNB1, ISLR, MIF, MLLT1, NFATC4, NFKBIL1, OMD, OGN, P4HB, PCDHGC3, PRELP, PYY, PTH1R, CLEC3B, TUB, COL14A1, ZFPL1, ZBTB16, GDF5, CILP, PRKRA, WISP1, MAP3K6, CABP1, BAG3, TTC37, LAPTM4A, CUL7, PJA2, ABCC9, PRG4, ANGPTL7, CALCOCO2, CDIPT, TM9SF1, CORIN, ZMYND11, NCKAP1, YIF1A, COPS8, SPIN1, KDELR1, TMED10, EMILIN1, MAP4K5, NXPH3, SCRG1, PHLDB1, GANAB, MKRN2, EID1, BCL2L13, TMEM59L, SNED1, HSPB7, TUBG2, IL17B, POMT2, COL5A3, TMED7, WDR41, KLHL26, OLAH, SPATA7, GLT8D1, IFT46, ZNF471, RIC8A, ZFAND3, PKNOX2, TTC23, C11orf95, YIPF2, FTO, PODNL1, TCEAL4, ADM2, SVEP1, UBXN6, IL17RC, LRRC15, PRRC1, ZNF358, 44084, NPHP4 |
| Chondrocytes_FANTOM_3___129 | Aran_et_al_2017 | ADRA1D, ACAN, AK1, ARL1, CACNA1C, CAMLG, RUNX1, COL10A1, COMP, CYP19A1, DPT, DVL1, ELN, ENO1, ERF, FGF7, FSHB, GOLGA4, MCHR1, GRIA3, HAS1, HDLBP, IBSP, IDUA, IGF1, ISLR, KRT10, LEP, MLLT1, NFATC4, NFKBIL1, OMD, OGN, P4HB, PDE3A, PGM1, PRELP, PYY, SGCD, SNTB2, TF, CLEC3B, TNNT3, COL14A1, ZFPL1, ZBTB16, CDK2AP1, GDF5, CILP, WISP1, MAP3K6, BAG3, TTC37, LAPTM4A, CUL7, PJA2, ABCC9, PRG4, ANGPTL7, CALCOCO2, YAP1, CDIPT, CORIN, NCKAP1, COPS8, KDELR1, SERINC3, OS9, TMED10, EMILIN1, MAP4K5, NXPH3, SCRG1, FNDC3A, SNX13, PHLDB1, GANAB, MKRN2, EID1, BCL2L13, TMEM59L, SNED1, RNF11, HSPB7, TUBG2, KLF15, SEC61A1, POMT2, NRBF2, COL5A3, WDR41, KLHL26, OLAH, SPATA7, GLT8D1, PKNOX2, C11orf95, YIPF2, PODNL1, HDAC11, TCEAL4, UBXN6, IL17RC, LRRC15, PRRC1, ZNF358, 44084, NPHP4 |
| Chondrocytes_HPCA_1___130 | Aran_et_al_2017 | ACAN, COMP, CSNK1A1, ERG, ISLR, NFATC4, PRELP, COL14A1, TBX4, MORF4L2, MPHOSPH6, DSTN, SCRG1, HSPB7, REM1, PODNL1 |
| Chondrocytes_HPCA_2___131 | Aran_et_al_2017 | ACAN, COMP, ERG, ISLR, LBP, PRELP, COL14A1, TBX4, MPHOSPH6, SCRG1, HSPB7, REM1 |
| Chondrocytes_HPCA_3___132 | Aran_et_al_2017 | ACAN, COMP, ERG, ISLR, PRELP, COL14A1, RGS11, TBX4, MPHOSPH6, SCRG1, REM1 |
| Classswitched_memory_Bcells_BLUEPRINT_1___133 | Aran_et_al_2017 | TNFRSF17, BLK, CR1, EPS15, RAPGEF1, PTPN6, RAD17, TRAF3, UBE2G1, UBE2I, BAIAP3, DEPDC5 |
| Classswitched_memory_Bcells_BLUEPRINT_2___134 | Aran_et_al_2017 | TNFRSF17, BLK, CR1, EPS15, RAPGEF1, NDUFA9, PTPN6, RAD17, SLC12A3, TAF6, TRAF3, UBE2G1, UBE2I, UBE2N, TRRAP, BAIAP3, DEPDC5, SCRN1, ABI1, CPSF4, SUB1, SP140, TNFRSF13B, ADAMDEC1, AFTPH, PRDM10, NARFL, PIKFYVE |
| Classswitched_memory_Bcells_BLUEPRINT_3___13 | Aran_et_al_2017 | TNFRSF17, BLK, CR1, EPS15, RAPGEF1, NDUFA9, PTPN6, RAD17, TRAF3, UBE2G1, UBE2I, BAIAP3, DEPDC5, ABI1, SEC24A, TNFRSF13B, AFTPH, NGLY1, PRDM10, |

| | | |
|---|---|---|
| 5 | | PIKFYVE |
| Classswitched_memory_Bcells_NOVERSHTERN_1___136 | Aran_et_al_2017 | TNFRSF17, BLK, CXCR5, MS4A1, CD80, COL19A1, GPR25, HLA-DQB2, PAX5, SPIB, BAIAP3, TNFRSF13B, SNED1, ZBTB32, FCRL2, KHDRBS2 |
| Classswitched_memory_Bcells_NOVERSHTERN_2___137 | Aran_et_al_2017 | BLK, CXCR5, MS4A1, CD80, COL19A1, GPR25, HLA-DQB2, PAX5, SPIB, BAIAP3, TNFRSF13B, SNED1, ZBTB32, FCRL2 |
| Classswitched_memory_Bcells_NOVERSHTERN_3___138 | Aran_et_al_2017 | BLK, CXCR5, MS4A1, CD80, COL19A1, GPR25, HLA-DQB2, PAX5, SPIB, BAIAP3, TNFRSF13B, SNED1, ZBTB32, FCRL2 |
| CLP_BLUEPRINT_1___139 | Aran_et_al_2017 | COX6C, DNTT, IGLL1, PLP2, PSMA6, VPREB1, AIMP1, TOMM20, GPN3, HIVEP3, MYL12B |
| CLP_BLUEPRINT_2___140 | Aran_et_al_2017 | COX6C, DNTT, H3F3B, IDH3A, IGLL1, OXA1L, PLP2, PSMA6, SNRPD1, VPREB1, AIMP1, ADNP, C19orf53, GPN3, C11orf57, HIVEP3, FAM76A |
| CLP_BLUEPRINT_3___141 | Aran_et_al_2017 | CALM1, COX6C, DNTT, GAPDH, H3F3B, IDH3A, IGBP1, IGLL1, OXA1L, PLP2, PSMA6, PSMB3, RFC4, RPL8, SNRPD1, VPREB1, AIMP1, ATP6V1G1, TOMM20, METAP2, DSTN, ADNP, C19orf53, ASCC1, GPN3, C11orf57, WDR33, NGLY1, HIVEP3, MYL12B, FAM76A |
| CMP_BLUEPRINT_1___142 | Aran_et_al_2017 | AZU1, MS4A3, CPA3, CRHBP, CRYGD, CTSG, ELANE, MS4A2, IGLL1, MPO, SERPINB10, PRG2, PRTN3, RNASE2, RNASE3, STAR, EPX, NAALADL1 |
| CMP_BLUEPRINT_2___143 | Aran_et_al_2017 | AZU1, MS4A3, CLC, CPA3, CRHBP, CTSG, ELANE, MS4A2, FLT3, HDC, MPO, SERPINB10, PRG2, PRTN3, RNASE2, RNASE3, EPX, NAALADL1, HPGDS |
| CMP_BLUEPRINT_3___144 | Aran_et_al_2017 | AZU1, MS4A3, CLC, CPA3, CRHBP, CTSG, ELANE, FLT3, MPO, PRG2, PRTN3, RNASE2, RNASE3, EPX, HPGDS |
| CMP_HPCA_1___145 | Aran_et_al_2017 | ALOX15, ANXA3, AZU1, BYSL, C1QB, CACNA1E, CAMK2A, RUNX1, CCNC, CD1E, MS4A3, CD63, CDSN, CLC, CLIC1, CNGA3, CPA3, CPB1, CPN1, CRYGA, CSF2RB, CTSG, DAB1, DEFA6, EDN3, EYA3, MS4A2, FMO2, AFF2, G6PC, GALR1, GDF9, GNAT1, GPD1, GRIA4, GRM2, GUCA2B, H2AFX, HDC, HSF4, IDUA, IL5RA, IL13, IMPG1, PDX1, ITIH3, ITIH4, KCNJ10, KCNN1, LCT, MEA1, MKI67, MPL, MPO, MYF5, MYH11, NDUFB4, OXT, P2RY4, PAX1, PAX3, PGR, SERPINB10, SERPINI2, PRG2, PRKG2, PYY, PTPRS, RCVRN, RHO, RORB, SCN1A, CCL18, CCL23, SIM1, SLC5A2, SLC18A2, SNAPC4, STAR, TNR, TP73, TRPC3, TYR, COL14A1, UNG, VPREB1, ZNF174, MADCAM1, CLPP, GDF5, HIST1H2BO, PLA2G6, BFSP2, IRS4, LIPF, BRSK2, SLC13A2, CCNB2, LONP1, NTN1, FHL5, FRMPD4, DLEC1, NAALADL1, CHAF1A, GLYAT, TRAIP, CACNG3, TUBA1B, PRG3, TAB1, ERLIN1, SPAG5, RASL10A, NEU3, RUVBL2, ACTL7A, PTPRT, SNW1, FAIM2, POLA2, IRF2BP1, DAZAP1, DNAI1, FOXB1, SNX5, NDOR1, GNMT, CNTN6, HPGDS, MYLPF, SMARCAL1, UBQLN3, RASL12, TOLLIP, MED18, COMMD4, CRTAC1, KLHL11, DHX32, SPATA7, CENPN, ZMAT5, C21orf62, CYSLTR2, VN1R1, SLC4A5, CADM3, PRODH2, DNASE2B, ALX4, XPNPEP3, DPEP3, GPR135, PDIA2, CYP3A43, LRRC19, C22orf46, LIN28A, ZBBX, RERGL, L2HGDH, TSGA10, LRRC3, NRIP2, COG7, TIMM50, TBC1D16, ZNF428, RAB40A, FAM76A, ATXN7L1, ANKRD34C |
| CMP_HPCA_2___146 | Aran_et_al_2017 | ADSS, MS4A3, MAPK14, TOR1A, FANCG, MS4A2, LPAR4, GPR27, LTC4S, LYL1, MPO, MYO9A, PGM1, POLE, MAP2K5, TRIM27, RNASE2, RREB1, SPN, STAR, TFCP2, TOP2B, TPSAB1, WHSC1, XPO1, ZNF221, HIST1H2BL, HIST1H2BO, HIST1H3C, HIST1H4C, NSMAF, JRK, ZMYM4, TGM5, SCAMP1, HDAC6, ZNF197, ERLIN1, TAF6L, USP19, EHMT2, HMGXB3, SMC5, MGA, KIAA1033, ZNF629, TNPO3, ZKSCAN5, ZZZ3, SPAG8, REV1, PHF7, VPS54, EXD3, SETD5, RHOT1, ZNF701, CENPJ, KLHL9, ZNF471, BAHCC1, HRH4, FAM111A, NARFL, S100PBP, MTHFSD, ZNF747, CCDC121, ZNF768, ATP8B4, ZKSCAN3, TSGA10, CDT1, USP48, ARHGAP33, ZNF780B, PIKFYVE, ZKSCAN4, ZNF324B |
| CMP_HPCA_3___147 | Aran_et_al_2017 | MS4A3, MS4A2, GPR27, LTC4S, LYL1, MPO, STAR, TPSAB1, ZNF221, HIST1H2BO, HIST1H3C, NAT6, CENPJ, BAHCC1 |
| CMP_NOVERSHTERN_1___148 | Aran_et_al_2017 | MS4A3, CD33, TOR1A, LPAR4, GPR27, MPO, TFCP2, TPSAB1, HIST1H4C, ZMYM4, TGM5, HDAC6, ZNF197, ERLIN1, TNPO3, NAT6, ZZZ3, SPAG8, FBXL4, ZNF701, CENPJ, ZNF471, S100PBP, ZKSCAN3, TSGA10, CDT1, ATP8B3, ZNF324B |
| CMP_NOVERSHTERN_2___149 | Aran_et_al_2017 | TOR1A, LPAR4, GPR27, MPO, ZMYM4, TGM5, HDAC6, ZNF197, ERLIN1, TNPO3, SPAG8, ZNF701, ZNF471, ZKSCAN3 |
| CMP_NOVERSHTERN_3___150 | Aran_et_al_2017 | MS4A2, LPAR4, GPR27, LTC4S, LYL1, MPO, TRIM27, STAR, TPSAB1, ZNF221, HIST1H2BO, HIST1H3C, ZNF197, ZNF629, SPAG8, PHF7, CENPJ, BAHCC1, ATP8B4, |

| | | |
|---|---|---|
| | | ZKSCAN3 |
| DC_BLUEPRINT_1___151 | Aran_et_al_2017 | ALOX15, CD1A, CD1B, CD1E, CCL13, CCL17, ALDH1A2, CD209 |
| DC_BLUEPRINT_2___152 | Aran_et_al_2017 | ALOX15, CD1A, CD1B, CD1E, HLA-DQA1, CCL13, CCL17, ALDH1A2, CD209 |
| DC_BLUEPRINT_3___153 | Aran_et_al_2017 | ALOX15, CD1A, CD1B, CD1E, FPR3, CCL13, CCL17, CD209 |
| DC_FANTOM_1___154 | Aran_et_al_2017 | C1QA, C1QB, CD1A, CD1B, CD1E, CD9, FPR3, CCL13, CCL17, CCL22, CLEC10A, TFEC, TREM2, SLAMF8 |
| DC_FANTOM_2___155 | Aran_et_al_2017 | ACHE, ALOX15B, CD1A, CD1B, CD1E, CD80, CD86, CCR7, DPYS, ETV3, GRIN1, GRSF1, HCRTR2, IL12B, IRF4, KCNC3, KCNN1, LOR, MCF2, RAB8A, NFKB1, PLD2, PRRG2, PTGIR, RNF2, CCL13, CCL17, CCL18, CCL22, CCL23, SLAMF1, SIGLEC1, TRAF1, TNFRSF4, TXN, VAV2, SLC30A4, CUL1, MAP3K6, TMSB10, MAP3K13, CEP350, BCL2L11, MPHOSPH6, SPINT2, HPS5, NXPH3, TDRD7, TMEM131, SUZ12, BCL2L13, FBXL4, SNX11, IL21R, TBC1D13, ARL8B, NECAP2, CAMK1G, CCDC81, SAMSN1, UBE2Z, PTGES2, SLCO5A1 |
| DC_FANTOM_3___156 | Aran_et_al_2017 | ALCAM, C1QA, C1QB, CD1A, CD1B, CD1C, CD1E, F13A1, FCER2, FPR3, TACSTD2, CCL13, CCL17, CCL22, CLEC10A, SPINT2, STAB1, CD209, TREM2, SLAMF8, MS4A6A |
| DC_HPCA_1___157 | Aran_et_al_2017 | ALOX15, CD1B, CD1E, CD80, CCL13, CCL17, CCL18, CCL19, CCL22, CD209, SLCO5A1 |
| DC_HPCA_2___158 | Aran_et_al_2017 | ALOX15, CD1B, CD1E, CD80, CCL13, CCL17, CCL18, CCL19, CCL22, CD209, SLCO5A1 |
| DC_HPCA_3___159 | Aran_et_al_2017 | ALOX15, CD1B, CD1E, CCL13, CCL17, CCL18, CCL19, SLCO5A1 |
| DC_IRIS_1___160 | Aran_et_al_2017 | CD1B, CD1E, CD86, FCER2, CCL17, ALDH1A2, RRP1B, KCNK13 |
| DC_IRIS_2___161 | Aran_et_al_2017 | ALOX15, C1QB, CD1A, CD1B, CD1E, CD80, CD86, DNASE1L3, F13A1, FCER2, FPR3, GUCA1A, HK3, HLA-DQA1, CCL8, CCL13, CCL17, CCL18, CCL22, CCL23, CCL24, ALDH1A2, HS3ST2, CLEC10A, SPINT2, FGL2, CD209, NAGPA, MS4A4A, KCNK13, MS4A6A |
| DC_IRIS_3___162 | Aran_et_al_2017 | ALOX15, C1QB, CD1A, CD1B, CD1E, CD86, FPR3, HLA-DQA1, CCL13, CCL17, CCL18, CCL22, CCL23, ALDH1A2, HS3ST2, CLEC10A, CD209 |
| Endothelial_cells_BLUEPRINT_1___163 | Aran_et_al_2017 | ACVRL1, TIE1, VWF, HYAL2, ARHGEF15, ROBO4, MMRN2, FAM124B |
| Endothelial_cells_BLUEPRINT_2___164 | Aran_et_al_2017 | ACVRL1, TIE1, VWF, HYAL2, ARHGEF15, ROBO4, MMRN2, FAM124B |
| Endothelial_cells_BLUEPRINT_3___165 | Aran_et_al_2017 | ACVRL1, ADSS, AP2A2, ANGPT2, ANXA2, ANXA3, RHOC, BMX, PTTG1IP, CANX, CAV1, CCT6A, CD9, CDC27, AP2S1, CLTA, DAD1, DDX10, EIF4G2, ERG, FOXC2, FLT1, FLT4, GJA4, GNB1, GOT2, GPR4, HSPA4, HTR2B, KDR, TNPO1, MNAT1, MYL6, NEDD8, NNAT, NOTCH4, PNP, PDE3A, PIK3C2A, PLS3, PPP2R2A, PSMB7, PSMD1, PSMD2, PSMD10, RALA, RANGAP1, RARS, RCN2, S100A6, MAPK12, SELE, SH3GL1, SLC16A1, SNTB2, SSBP1, TARBP2, TEK, TIE1, TJP1, CLDN5, TPD52L2, HSP90B1, UFD1L, VWF, FXR1, SCARF1, DYNLL1, HYAL2, CDC123, EIF2B2, MTMR2, BCL10, TNFSF18, ITGB1BP1, TAOK2, EI24, GDF3, FEZ2, PPM1F, SAE1, KIF20A, ACTR1A, PDIA6, TXNDC9, PCGF3, TIMM17A, CARM1, SEMA6B, IPO7, PITRM1, ARPC1A, IGF2BP3, YKT6, LYVE1, COPS6, PWP1, CDC37, FAM107A, ECD, TUSC2, ARHGEF15, MMRN1, NCBP2, CD93, ATF6, TTLL5, STAB1, ARL2BP, MTCH1, TMEM184B, CLEC1A, BFAR, PCDH12, SPTBN5, EMCN, BTBD1, SOX18, ROBO4, TMED9, DEF8, RASIP1, TMEM39B, LRRC59, NPLOC4, CISD1, KLHL9, ANO2, MRPL17, SPATS2, CXorf36, MMRN2, FAM124B, EDC3, MYCT1, NETO2, PLVAP, DCTN5, G6PC3, MYL12B, KANK3 |
| Endothelial_cells_ENCODE_1___166 | Aran_et_al_2017 | ACVRL1, ANGPT2, ANXA2, RHOC, BMX, PTTG1IP, CAV1, CLTA, DAD1, ERG, FOXC2, GPR4, KDR, MYL6, PIK3C2A, PLS3, PSMD10, RALA, MAPK12, SLC16A1, TEK, TIE1, CLDN5, VWF, HYAL2, FEZ2, ACTR1A, TXNDC9, PCGF3, COPS6, PWP1, FAM107A, ECD, ARHGEF15, MMRN1, CD93, TTLL5, MTCH1, CLEC1A, PCDH12, EMCN, SOX18, ROBO4, RASIP1, LRRC59, MRPL17, CXorf36, MMRN2, MYCT1, PLVAP, KANK3 |
| Endothelial_cells_ENCODE_2___167 | Aran_et_al_2017 | ACVRL1, ANGPT2, ANXA2, RHOC, BMX, PTTG1IP, CAV1, CLTA, ERG, GPR4, KDR, PLS3, PSMD10, RALA, MAPK12, SLC16A1, TIE1, CLDN5, VWF, HYAL2, ACTR1A, PCGF3, COPS6, PWP1, FAM107A, ARHGEF15, MMRN1, CD93, TTLL5, MTCH1, CLEC1A, PCDH12, EMCN, SOX18, ROBO4, RASIP1, MRPL17, CXorf36, MMRN2, MYCT1, PLVAP, KANK3 |

| | | |
|---|---|---|
| Endothelial_cells_ENCODE_3___168 | Aran_et_al_2017 | ACVRL1, ANGPT2, RHOC, BMX, CLTA, GPR4, KDR, PLS3, PSMD10, RALA, MAPK12, TIE1, VWF, HYAL2, ACTR1A, ARHGEF15, CLEC1A, EMCN, SOX18, ROBO4, RASIP1, CXorf36, MMRN2, MYCT1, KANK3 |
| Endothelial_cells_FANTOM_1___169 | Aran_et_al_2017 | ACVRL1, ANGPT2, KDR, TIE1, HYAL2, ARHGEF15, EMCN, ROBO4, MMRN2, MYCT1 |
| Endothelial_cells_FANTOM_2___170 | Aran_et_al_2017 | ACVRL1, ANGPT2, BMX, KDR, TIE1, HYAL2, ARHGEF15, EMCN, ROBO4, MMRN2, MYCT1 |
| Endothelial_cells_FANTOM_3___171 | Aran_et_al_2017 | ACVRL1, ANGPT2, ANXA3, ART4, BMX, CAV1, ERG, FLT1, GJA4, GPR4, HTR2B, KDR, KRT19, TACSTD2, PDE3A, RALA, MAPK12, SELE, SNTB2, TIE1, CLDN5, VWF, HYAL2, EIF2B2, TNFSF18, GDF3, IGF2BP3, ARHGEF15, MMRN1, CD93, CLEC1A, EMCN, SOX18, ROBO4, RASIP1, CXorf36, MMRN2, FAM124B, MYCT1, KANK3 |
| Endothelial_cells_HPCA_1___172 | Aran_et_al_2017 | ANGPT2, BMX, KDR, TIE1, CLDN5, VWF, MMRN1, CD93, STAB1, EMCN, ROBO4, GIMAP4 |
| Endothelial_cells_HPCA_2___173 | Aran_et_al_2017 | ANGPT2, BMX, FLT4, KDR, TIE1, VWF, SEMA6B, MMRN1 |
| Endothelial_cells_HPCA_3___174 | Aran_et_al_2017 | ACVRL1, CDC27, AP2S1, ERCC1, FBL, FDPS, FOXC2, FLT4, GJA4, GPR4, HTR1B, LYL1, NOTCH4, NOVA2, PNP, PPP2R2A, PSMC5, RALA, RELA, RPS14, MAPK12, TIE1, CLDN5, VWF, CLPP, SCARF1, HYAL2, EIF2B2, MTMR2, BCL10, TNFSF18, BUB3, TAOK2, DLGAP5, ACTR1A, LYPLA1, STK25, SEMA6C, SEMA6B, PTTG2, WDR4, LYVE1, STRAP, TUSC2, ARHGEF15, CD93, TTLL5, N4BP3, STAB1, EDC4, PRKD2, PRPF19, GIT1, CLEC1A, ROBO4, TMEM39B, CEP55, NPLOC4, CISD1, METTL3, TUT1, SPATS2, FAM65A, MMRN2, FAM124B, MYCT1, MTG1, KANK3, GIMAP6 |
| Eosinophils_BLUEPRINT_1___175 | Aran_et_al_2017 | C3AR1, CLC, CCR3, DRP2, IL5RA, KCNA5, KIF5A, TACSTD2, NPY2R, PLXNB3, RGS13, TNFSF11, ADAM18, CUX2, CDH19, RASL12, HRH4, NYX |
| Eosinophils_BLUEPRINT_2___176 | Aran_et_al_2017 | CCR3, DRP2, GIPR, TACSTD2, RGS13, SYCP1, PHLDA2, TNFSF11, NXPH3, LMTK2, AP4E1, FBXO40, HRH4, TRIM48, MMRN2 |
| Eosinophils_BLUEPRINT_3___177 | Aran_et_al_2017 | AGTR2, ASPA, ATOH1, BMX, C3AR1, MS4A3, CLC, CCR1, CCR3, DRP2, FCER1A, FMO3, GAST, GIPR, HDC, IL5RA, KCNA5, KIF5A, LECT2, TACSTD2, MOG, NPY2R, PLXNB3, PRL, RGS13, STATH, TNFSF11, ADAM21, ADAM18, HS3ST2, ZNF197, POLQ, NXPH3, LMTK2, CUX2, CDH19, PURG, RASL12, FBXO40, MYO3A, FEV, LRRC36, MMP26, HRH4, NYX, TRIM48 |
| Eosinophils_FANTOM_1___178 | Aran_et_al_2017 | ADORA3, ALOX15, CA4, ENTPD2, CEBPE, CEACAM8, CLC, CCR3, DEFA4, GIPR, HIC1, IL5RA, LTF, MNT, RARA, SLC19A1, KSR1, PGLYRP1, CD101, P2RY14, KCNK7, OLIG2, PTTG2, EPN2, SETD1B, KDM6B, ABTB2, SIGLEC8, GMIP, CYSLTR2, MKL1, HRH4, DPEP2, MMP25, MBOAT7, CORO7, ARHGAP33, TRPM6 |
| Eosinophils_FANTOM_2___179 | Aran_et_al_2017 | ADORA3, ALOX15, CA4, ENTPD2, CEBPE, CLC, CCR3, DEFA4, GIPR, IL5RA, LTF, MNT, RARA, SLC19A1, KSR1, PGLYRP1, KCNK7, OLIG2, PTTG2, EPN2, SETD1B, KDM6B, ABTB2, SIGLEC8, GMIP, CYSLTR2, MKL1, HRH4, DPEP2, MMP25, MBOAT7, CORO7, ARHGAP33, TRPM6 |
| Eosinophils_FANTOM_3___180 | Aran_et_al_2017 | ADORA3, ALOX15, BPI, CA4, CAMP, ENTPD2, CEBPE, CEACAM8, CLC, CCR3, CSF2RB, DEFA4, GIPR, HIC1, IL5RA, LTF, MNT, RARA, CCL23, SLC19A1, KSR1, PGLYRP1, CD101, P2RY14, KCNK7, OLIG2, PTTG2, EPN2, SETD1B, KDM6B, SRRM2, ABTB2, SIGLEC8, GMIP, CYSLTR2, MKL1, HRH4, DPEP2, DPEP3, MMP25, MBOAT7, CORO7, ARHGAP33, TRPM6 |
| Eosinophils_NOVERSHTERN_1___181 | Aran_et_al_2017 | ALOX15, CEBPE, CLC, CCR3, DEFA4, IL5RA, RARA, KCNK7, EPN2, KDM6B, ABTB2, SIGLEC8, HRH4, DPEP2, MMP25, TRPM6 |
| Eosinophils_NOVERSHTERN_2___182 | Aran_et_al_2017 | ALOX15, CEBPE, CLC, CCR3, DEFA4, IL5RA, KSR1, ABTB2, SIGLEC8, HRH4, DPEP2, MMP25, TRPM6 |
| Eosinophils_NOVERSHTERN_3___183 | Aran_et_al_2017 | ENTPD2, CEBPE, CLC, CCR3, GIPR, HIC1, IL5RA, LTF, KSR1, OLIG2, SIGLEC8, HRH4, DPEP2, CORO7, ARHGAP33 |
| Epithelial_cells_ENCODE_1___184 | Aran_et_al_2017 | SFN, TACSTD2, PRSS8, AP1M2, B3GNT3, CBLC, HES2, RAB25, S100A14 |
| Epithelial_cells_ENCODE_2___185 | Aran_et_al_2017 | FLNB, SFN, GRB7, LAD1, LAMA5, PRSS8, SEMA3F, SOX15, TUFT1, SLC10A3, IER3, SH2D3A, AP1M2, CNKSR1, B3GNT3, RHOD, HES2, TMEM40, TBC1D2, RAB25, S100A14 |
| Epithelial_cells_ENCODE_3___186 | Aran_et_al_2017 | BIK, AP1S1, DFNA5, DSG3, HBEGF, EFNB1, NR2F6, EVPL, EXT2, F3, FLNB, GJB3, GJB5, SFN, GRB7, IRF6, JUP, LAD1, LAMA5, LIMK2, TACSTD2, MST1R, PI3, PLAGL2, PRRG2, PRSS8, RELB, SEMA3F, SHC1, SLC12A4, SOX15, SPINT1, ST14, STXBP2, TAPBP, TUFT1, CORO2A, BTG2, SLC10A3, AXIN1, TNK1, TNFSF9, ADAM15, |

| | | |
|---|---|---|
| | | TNFRSF10B, IER3, RPS6KA4, PDLIM1, CELSR1, FGFBP1, SH2D3A, AP1M2, CNKSR1, B3GNT3, CDC42EP2, FST, PPP1R13L, ARHGEF18, ETHE1, TNFRSF21, TFCP2L1, RHOD, GLTP, A4GALT, RHOF, HES2, SSH3, TMEM40, TBC1D2, DOK4, FGD6, TMPRSS4, RAB25, SMAGP, S100A14, XYLT2, PORCN, ZFYVE21, CHAC1, C1orf116, ZBED2, RHBDF2, PIP4K2C, FBXL18, LRRC8E, EPPK1 |
| Epithelial_cells_FANTOM_1___187 | Aran_et_al_2017 | CLDN4, DSC2, SFN, ITGB4, ITGB6, KRT7, LAD1, LAMA3, LAMB3, SPINT1, THBD, SH2D3A, AP1M2, MPZL2, G0S2, LSR, RAB25 |
| Epithelial_cells_FANTOM_2___188 | Aran_et_al_2017 | ADM, CLDN4, DSC2, EFNA1, EGFR, EVPL, SFN, ITGB4, ITGB6, KRT7, KRT17, LAD1, LAMA3, LAMB3, LLGL2, TACSTD2, PCDH1, PDGFB, PPL, PRSS8, S100A2, SCNN1A, SPINT1, ST14, THBD, HMGA2, PTGES, RAB3D, SH2D3A, AP1M2, TSPAN1, MPZL2, PPP1R13L, RAP1GAP2, DAPP1, G0S2, ANGPTL4, LSR, SLC35F2, EPS8L1, FERMT1, RAB25, C1orf116, RAB11FIP1, ALS2CL |
| Epithelial_cells_FANTOM_3___189 | Aran_et_al_2017 | CLDN4, DSC2, ELF3, SFN, ITGB4, ITGB6, KRT6A, KRT7, LAD1, LAMA3, LAMB3, SLPI, ST14, TGFA, SCEL, PTGES, SH2D3A, AP1M2, MPZL2, EHF, G0S2, LSR, S100A14 |
| Epithelial_cells_HPCA_1___190 | Aran_et_al_2017 | F3, SFN, TACSTD2, PRSS8, SLPI, AP1M2, CBLC, RAB25, S100A14 |
| Epithelial_cells_HPCA_2___191 | Aran_et_al_2017 | F3, SFN, PRSS8, SLPI, STXBP2, AP1M2, CBLC, RAB25, S100A14 |
| Epithelial_cells_HPCA_3___192 | Aran_et_al_2017 | NQO1, F3, FLNB, SFN, TACSTD2, PRSS8, SDC4, SLPI, STXBP2, RASSF7, AP1M2, B3GNT3, SH3BP1, CBLC, APOBEC3C, RIPK4, HES2, RAB25, S100A14 |
| Erythrocytes_BLUEPRINT_1___193 | Aran_et_al_2017 | ALAS2, CA1, EPB42, GYPE, HBD, HMBS, MYL4, RHAG, EPX, KLF1, XPO7, AHSP |
| Erythrocytes_BLUEPRINT_2___194 | Aran_et_al_2017 | ALAS2, CA1, EPB42, GYPE, HBD, HMBS, MYL4, RHAG, EPX, KLF1, XPO7, AHSP |
| Erythrocytes_BLUEPRINT_3___195 | Aran_et_al_2017 | ALAS2, CA1, EPB42, GYPE, HBD, HMBS, MYL4, RHAG, EPX, KLF1, XPO7, AHSP |
| Erythrocytes_FANTOM_1___196 | Aran_et_al_2017 | ALAS2, CA1, EPB42, GYPA, GYPE, KRT1, MYL4, SLC4A1, AHSP |
| Erythrocytes_FANTOM_2___197 | Aran_et_al_2017 | ALAS2, EPB42, GYPA, GYPB, GYPE, HBB, HBD, MYL4, PKLR, PRG2, RHAG, RHCE, RHD, SLC4A1, KLF1, AHSP, TSPO2 |
| Erythrocytes_FANTOM_3___198 | Aran_et_al_2017 | ALAS2, ART4, EPB42, GYPA, GYPB, GYPE, HBB, HBD, HMBS, MYL4, PKLR, RHAG, RHD, KLF1, AHSP |
| Erythrocytes_HPCA_1___199 | Aran_et_al_2017 | CENPA, GATA1, RHAG, SPTA1, AURKA, KLF1, GLRX5, AHSP |
| Erythrocytes_HPCA_2___200 | Aran_et_al_2017 | ALAS2, CA1, EPB42, GYPE, HBD, HMBS, MYL4, RHAG, EPX, KLF1, XPO7, AHSP |
| Erythrocytes_HPCA_3___201 | Aran_et_al_2017 | ALAS2, CA1, EPB42, GYPE, HBD, HMBS, MYL4, RHAG, EPX, KLF1, XPO7, AHSP |
| Erythrocytes_NOVERSHTERN_1___202 | Aran_et_al_2017 | APLNR, ALAS2, ATP1B2, BRCA1, BUB1B, CDC20, CDKN3, CENPF, CHEK1, DES, EPB42, FEN1, GYPA, GYPB, GYPE, HBB, HMBS, HMGB2, KEL, KIF22, EPCAM, MCM4, MYL4, NEK2, OAT, PCNA, PKLR, POLE2, PRG2, RHD, RRM1, RRM2, STIL, SLC2A4, SLC4A1, AURKA, TOP2A, TUBG1, ST7, HIST1H4C, RAD54L, CCNB2, MINPP1, DLGAP5, MELK, GINS1, TROAP, TRIM10, RCL1, PRMT3, UBAC1, KLF1, KIF2C, ZWINT, OIP5, PAXIP1, KIF4A, HTRA2, GMNN, NUSAP1, GLRX5, AHSP, DTL, FANCI, HJURP, MCM10, C1orf112, CENPN, KIF15, SPC25, FKBPL, GINS3, CENPO, CDCA3, TSPO2 |
| Erythrocytes_NOVERSHTERN_2___203 | Aran_et_al_2017 | ALAS2, BRCA1, CENPF, DES, EPB42, FEN1, GYPA, GYPB, GYPE, HBB, HMBS, HMGB2, KIF22, MYL4, NEK2, PCNA, PKLR, POLE2, RHD, RRM1, TOP2A, TUBG1, CCNB2, MINPP1, DLGAP5, MELK, GINS1, TROAP, PRMT3, UBAC1, KLF1, GMNN, NUSAP1, AHSP, DTL, FANCI, MCM10, CENPN, FKBPL, CDCA3 |
| Erythrocytes_NOVERSHTERN_3___204 | Aran_et_al_2017 | ALAS2, AMHR2, BIRC5, ART4, ATP1B2, BRCA1, BUB1, CA1, CAST, CCNA2, CCNB1, CDK1, CDC20, CDKN3, CENPE, CENPF, CSE1L, DES, EPB42, EPRS, FEN1, GATA1, GYPA, GYPB, GYPE, HBB, HBD, HMBS, HMGB2, HPS1, KEL, KIF22, EPCAM, MYL4, NEK2, PCNA, PKLR, PNMT, POLE2, PRG2, RAD51, RFC4, RHAG, RHD, RPA3, RRM1, RRM2, STIL, SLC2A4, SLC16A1, SPTA1, TAL1, TARS, TOP2A, TUBG1, UMPS, ST7, CHAF1B, GFI1B, HIST1H4C, DPM2, EIF2S2, CCNB2, PTTG1, MINPP1, RGS6, DLGAP5, MELK, GINS1, DCLRE1A, RBX1, TROAP, SMC4, RCL1, PRMT3, UBAC1, SMC2, KLF1, DBF4, METAP2, KIF2C, ZWINT, WBP4, RACGAP1, GMNN, NUSAP1, AHSP, DTL, NCAPG2, CDCA8, FANCI, HJURP, MCM10, CENPN, PBK, SPC25, FKBPL, NUP37, CDCA3 |
| Fibroblasts_ENCODE_1___205 | Aran_et_al_2017 | ADH5, ARF4, ARL1, ASPA, ATP2A2, BAD, BMPR1A, CACNA1C, AP2M1, CSNK1A1, CSNK1G3, DCTD, DPT, ECT2, ELN, ETF1, FKTN, FGF7, GARS, GOLGA4, GRIA1, |

| | | |
|---|---|---|
| | | GRIA3, HIF1A, HLCS, HTR2A, HTR2B, ITIH3, IPO5, KRT19, LGALS1, SMAD5, MARS, MYH1, MYH2, 44076, NFATC4, OCRL, PPIB, PRKG1, PTGIR, RAD23B, RCN2, PRPH2, RYK, ATXN2, SGCD, SGCG, SHMT2, SIM1, SNTB2, TBX5, CLEC3B, TPD52L2, TSPYL1, SLC35A2, VCL, WNT2, ZFPL1, B4GALT2, MPZL1, ZMYM4, ZFYVE9, GGPS1, SCAMP1, BAG2, PRDX6, RNF14, SCRN1, RNF41, TFG, YAP1, SPTLC1, CORIN, FRS2, SPIN1, TMED10, KDELR2, DSTN, EMILIN1, MAP4K5, XPOT, RRAS2, RAB11FIP2, RAB3GAP1, TRIM32, GANAB, DNAJC13, ICMT, LMOD1, MYOF, TBL2, SEC22A, NPTN, HSPB7, SEC61A1, TNFRSF12A, NGRN, MBTPS2, AMOTL2, GPR85, TMED9, ASPN, ST7L, SLC35A5, KIF26B, GPATCH2, IMPACT, POMGNT1, CAND1, ACTR10, SAR1A, THAP10, CCDC90B, MAGEF1, ACBD3, C7orf25, AHNAK, FTO, ZC3H14, PODNL1, TCEAL4, SVEP1, TTC26, SLC25A32, ADAMTS12, TM2D1, ANKRD40, MYL12B, LRRC42, HSPB6, 44084, TOR1AIP2, TXLNA, RNASEH1, DPY19L4, SNX19 |
| Fibroblasts_ENCODE_2___206 | Aran_et_al_2017 | ARF4, ATP2A2, BMPR1A, CACNA1C, CSNK1A1, CSNK1G3, DPT, ELN, FKTN, FGF7, GRIA1, GRIA3, HIF1A, HTR2B, ITIH3, IPO5, LGALS1, MYH2, 44076, NFATC4, PRKG1, RAD23B, PRPH2, RYK, SGCD, SGCG, SIM1, TBX5, SLC35A2, WNT2, ZFPL1, SCAMP1, BAG2, PRDX6, SCRN1, RNF41, YAP1, SPTLC1, CORIN, FRS2, SPIN1, KDELR2, MAP4K5, XPOT, RRAS2, RAB11FIP2, TRIM32, MYOF, SEC22A, HSPB7, MBTPS2, AMOTL2, GPR85, ASPN, SLC35A5, KIF26B, GPATCH2, CAND1, THAP10, CCDC90B, ACBD3, AHNAK, PODNL1, SVEP1, TM2D1, MYL12B, LRRC42, 44084, TXLNA, RNASEH1, DPY19L4 |
| Fibroblasts_ENCODE_3___207 | Aran_et_al_2017 | BMPR1A, DPT, ELN, FKTN, FGF7, GRIA3, HTR2B, ISLR, ITIH3, KRT19, MYH2, PRKG1, SGCD, SGCG, SIM1, SNTB2, TBX5, WNT2, BAG2, YAP1, CORIN, TRIM32, LMOD1, MYOF, TNFRSF12A, AMOTL2, ASPN, KIF26B, THAP10, HSPB6, DPY19L4 |
| Fibroblasts_FANTOM_1___208 | Aran_et_al_2017 | ADD1, ADH1B, ALDH9A1, ANXA11, ARF4, ARHGAP6, C7, CACNA1C, CAMLG, CIRBP, CSF1, CYBA, DPT, DUT, FGF7, FMO2, FTL, GARS, GOLGA1, GSTM5, HEXA, HIC1, HPS1, HTR2B, IDUA, ISLR, JAK3, IPO5, LTBR, LTC4S, SMAD5, MGMT, MGST3, MMP17, MMP19, NFATC4, NFE2L2, P4HB, PDE4A, PFDN5, PGK1, PIK3R2, PRKG2, MASP1, PTGIR, RASA2, RNF4, RNH1, ROM1, RPL37A, SH3BP2, SNAPC2, TADA2A, TBX5, TCF21, TFDP1, CLEC3B, TNXB, VIM, ZNF32, DEK, NDST2, CGGBP1, WISP1, S1PR2, RPL23, HAND2, BAG2, LAPTM4A, HDAC5, MPHOSPH10, TFG, PRDX4, MTHFD2, SEC24A, KDELR1, KDELR2, EMILIN1, LSM6, XPOT, ZBTB1, FAIM2, GANAB, CSTF2T, ABCA6, SNED1, TOR1AIP1, CECR5, RASL12, ZNF771, AMOTL2, HDAC7, UBE2D4, ASPN, TXNL4B, SLC35A5, SHQ1, ADI1, DDX19A, ZNF444, FBXL8, ZNF446, EXOC1, SPATA7, TMEM165, PCDHGA11, GPR137, CASS4, ATP13A1, KIAA1614, EDA2R, PAPPA2, MOSPD3, C11orf95, C7orf25, ATG9A, TBC1D17, SLC35E1, SVEP1, SLC25A32, C6orf62, WDR73, MFSD5, ZNF358, EML3, DPY19L4 |
| Fibroblasts_FANTOM_2___209 | Aran_et_al_2017 | CAMLG, FTL, GSTM5, ISLR, JAK3, MMP19, NFATC4, PRKG2, PTGIR, S1PR2, HAND2, ZNF771, GPR137, KIAA1614, SPAG16, SVEP1, C6orf120 |
| Fibroblasts_FANTOM_3___210 | Aran_et_al_2017 | ADH1B, ARHGAP6, C7, DPT, FGF7, FMO2, FTL, HIC1, HTR2B, ISLR, JAK3, MGMT, MIF, MMP17, MMP19, POLR2E, PRKG2, MASP1, PTGIR, ROM1, MRPL12, TBX5, TCF21, TNXB, COL14A1, GDF5, WISP1, S1PR2, HAND2, PRDX4, KDELR1, EMILIN1, ABCA6, ASPN, C19orf24, ADI1, GPR137, KIAA1614, PODNL1, ADM2, SVEP1, ZNF358, SIX5 |
| Fibroblasts_HPCA_1___211 | Aran_et_al_2017 | ADH1B, C7, CACNA1C, CAMLG, FTL, HEXA, IDUA, ISLR, JAK3, IPO5, MGST3, NFATC4, NFE2L2, PRKG2, MASP1, PTGIR, RNH1, TBX5, TCF21, TFDP1, HAND2, BAG2, PRDX4, MTHFD2, XPOT, CSTF2T, ZNF771, ADI1, FBXL8, PCDHGA11, GPR137, KIAA1614, PAPPA2, C7orf25 |
| Fibroblasts_HPCA_2___212 | Aran_et_al_2017 | ADH1B, ANXA11, ARF4, C7, CACNA1C, CAMLG, FTL, GOLGA1, HEXA, IDUA, ISLR, JAK3, IPO5, LTBR, SMAD5, MGST3, NFATC4, NFE2L2, PFDN5, PRKG2, MASP1, PTGIR, RNH1, TADA2A, TBX5, TCF21, TFDP1, VIM, CGGBP1, HAND2, BAG2, HDAC5, PRDX4, MTHFD2, KDELR1, TMEM115, XPOT, CSTF2T, DNPEP, MKRN2, TOR1AIP1, ZCCHC4, RASL12, ZNF771, ADI1, FBXL8, PCDHGA11, GPR137, CASS4, ATP13A1, CYP20A1, ZNF471, KIAA1614, PAPPA2, MOSPD3, C7orf25, UBE3B, TEX261, ZNF358, EML3, C6orf120 |
| Fibroblasts_HPCA_3___213 | Aran_et_al_2017 | ADD1, ADH1B, ALDH9A1, ANXA11, ARF4, ARHGAP6, C7, CACNA1C, CAMLG, CIRBP, CSF1, CYBA, DPT, DUT, FGF7, FMO2, FTL, GARS, GOLGA1, GSTM5, HEXA, HIC1, HPS1, HTR2B, IDUA, ISLR, JAK3, IPO5, LTBR, LTC4S, SMAD5, MGMT, MGST3, MMP17, MMP19, NFATC4, NFE2L2, P4HB, PDE4A, PFDN5, PGK1, PIK3R2, PRKG2, MASP1, PTGIR, RASA2, RNF4, RNH1, ROM1, RPL37A, SH3BP2, SNAPC2, TADA2A, TBX5, TCF21, TFDP1, CLEC3B, TNXB, VIM, ZNF32, DEK, NDST2, CGGBP1, WISP1, |

| | | |
|---|---|---|
| | | S1PR2, RPL23, HAND2, BAG2, LAPTM4A, HDAC5, MPHOSPH10, TFG, PRDX4, MTHFD2, SEC24A, KDELR1, KDELR2, EMILIN1, LSM6, XPOT, ZBTB1, FAIM2, GANAB, CSTF2T, ABCA6, SNED1, TOR1AIP1, CECR5, RASL12, ZNF771, AMOTL2, HDAC7, UBE2D4, ASPN, TXNL4B, SLC35A5, SHQ1, ADI1, DDX19A, ZNF444, FBXL8, ZNF446, EXOC1, SPATA7, TMEM165, PCDHGA11, GPR137, CASS4, ATP13A1, KIAA1614, EDA2R, PAPPA2, MOSPD3, C11orf95, C7orf25, ATG9A, TBC1D17, SLC35E1, SVEP1, SLC25A32, C6orf62, WDR73, MFSD5, ZNF358, EML3, DPY19L4 |
| GMP_BLUEPRINT_1___214 | Aran_et_al_2017 | ARHGAP6, ATP5J, CD5L, MS4A3, CDH9, CPA3, CRHBP, CRYGD, CTSG, DNTT, FLT3, GABPA, LPAR4, H2AFZ, HDC, HMGB2, HNRNPA1, ITGA9, KCNJ14, LDHB, MPO, MTIF2, HNRNPM, NPM1, SERPINB10, PNN, PRG2, PRKG2, PSMA4, RAG2, RNASE2, RNASE3, RPL5, RYR3, SELP, STIL, SRP9, SSB, STXBP3, TEC, TOP2B, TRH, UBB, UMPS, VPREB1, XPO1, DEK, NCOA4, TTF2, UBA3, UBA2, SMNDC1, ARPP21, MRPL3, PARK7, CBX3, SMC5, GTPBP4, HPGDS, GNL2, CLEC1B, NOL7, UFC1, VPS54, GAR1, MRPL20, TTC27, WDR12, KIF17, MAP7D3, CXorf21 |
| GMP_BLUEPRINT_2___215 | Aran_et_al_2017 | APLNR, ARHGAP6, ATP5J, CD5L, MS4A3, CDH9, CPA3, CRHBP, CRYGD, CTSG, DHX9, DNTT, EIF4E, ELANE, EWSR1, FOXI1, FLT3, GABPA, GLRA2, LPAR4, GUCY2D, H2AFZ, HDC, HMGB2, HNRNPA1, HNRNPD, ITGA9, KARS, KCNJ14, KPNA2, LDHB, MC4R, MPO, MTIF2, HNRNPM, NDUFC2, NPM1, SERPINB10, PNN, PRG2, PRKG2, PSMA4, RAG2, RNASE2, RNASE3, RPL5, RPL15, RYR3, SELP, STIL, SLN, SRP9, SSB, STXBP3, TEC, TOP2B, TRH, UBB, UMPS, VPREB1, XPO1, DEK, NCOA4, ANP32A, TAF15, TTF2, UBA3, COX7A2L, DDX21, UBA2, SMNDC1, ARPP21, MRPL3, PARK7, CBX3, SMC5, METAP1, GTPBP4, CLDN17, HPGDS, GNL2, CLEC1B, RWDD1, NOL7, NOP16, UFC1, VPS54, LUC7L3, GAR1, COMMD8, MRPL20, TTC27, WDR12, KIF17, MAP7D3, CXorf21 |
| GMP_BLUEPRINT_3___216 | Aran_et_al_2017 | ATP5J, CD5L, MS4A3, CDH9, CPA3, CRHBP, CRYGD, CTSG, DNTT, FOXI1, FLT3, LPAR4, H2AFZ, HDC, HMGB2, HNRNPA1, LDHB, MPO, MTIF2, HNRNPM, NPM1, SERPINB10, PNN, PRG2, PSMA4, RAG2, RNASE2, RNASE3, RPL5, RPL15, RYR3, STIL, SRP9, SSB, TEC, TOP2B, TRH, UBB, UMPS, VPREB1, DEK, TTF2, UBA3, UBA2, ARPP21, MRPL3, PARK7, CBX3, SMC5, GTPBP4, GNL2, NOL7, UFC1, VPS54, GAR1, MRPL20, TTC27, WDR12, KIF17 |
| GMP_HPCA_1___217 | Aran_et_al_2017 | ALOX12, CD33, CEACAM4, CTSG, DNTT, ERCC3, GPR3, GSTM5, H3F3B, HNRNPA2B1, INSL3, KARS, KCNJ14, LPO, MPO, NCL, OMD, PARK2, SERPINI2, POU2F1, PRTN3, PEX2, SLC5A5, SUPV3L1, SUV39H1, TRH, DNAJC7, VPREB1, ZNF35, ZNF207, LUZP1, SLC25A14, TRIP11, CNOT8, NCOR1, PUM1, BMS1, SMG7, ARPP21, CLP1, GLMN, LAMB4, GPATCH8, WDR43, RPRD2, GTPBP4, ZNF593, CDK5RAP1, SETD4, POLE3, WDR60, RNF220, ENOSF1, PRPF40A, DDX27, IQCC, NGLY1, ZC3H15, C21orf59, KIF17, RBM25, MRPL9, B3GNT4, MUL1, TCTN2, TTC26, CHD9, CEP63, LAS1L, RBM4B, MYOZ3, IQSEC3 |
| GMP_HPCA_2___218 | Aran_et_al_2017 | CEACAM8, CLC, CSF1R, DEFA4, FEN1, LPAR4, MEFV, CXCL9, MPO, CFP, PRG2, RNASE2, CLEC1B, ZNF710 |
| GMP_HPCA_3___219 | Aran_et_al_2017 | CCNT2, CD33, CHD4, AP2M1, DDOST, DHX9, DNMT1, ENO1, ERCC3, FLT3, GAS8, GGCX, GPR3, H3F3B, HNRNPA1, HNRNPA2B1, HNRNPH3, IK, IL3RA, IMPDH2, INSL3, KARS, KCNJ14, IPO5, LPO, MPO, MTIF2, NCL, OMD, PAFAH1B2, SERPINI2, POU2F1, PRTN3, RFC1, SLC5A5, SUPV3L1, SUV39H1, TCP1, NR2C1, TRH, DNAJC7, UBE2G2, ZNF35, ZNF207, LUZP1, ARID1A, TTF2, EIF3A, HDAC3, SLC24A1, CNOT8, RBM39, NCOR1, BMS1, ZBTB39, SMG7, RBM19, THRAP3, HMGXB4, HNRNPR, CALCOCO2, DNAJA2, ARPP21, ERP29, CLP1, SF3B2, DUSP12, GPN1, LAMB4, WDR43, RPRD2, PPIP5K2, GTPBP4, ATXN10, COG4, APPL1, DNAJC2, GNL2, GOLGA7, ANAPC5, LUC7L3, TXNL4B, MED9, WDR60, HEATR1, RNF220, SLC25A36, WDR41, ENOSF1, DDX27, NGLY1, ZC3H15, NIT2, KIF17, USP36, RBM25, MRPL9, B3GNT4, MUL1, MAP7D3, CSPP1, TCTN2, CPSF7, LAS1L, HPS4, G6PC3, IRGQ, PRRC1, HNRNPA3, IQSEC3 |
| GMP_NOVERSHTERN_1___220 | Aran_et_al_2017 | CLC, CPA3, CRHBP, DNTT, FLT3, MPO, PRG2, RNASE2 |
| GMP_NOVERSHTERN_2___221 | Aran_et_al_2017 | CLC, CPA3, CRHBP, DNTT, FLT3, MPO, PRG2, RNASE2 |
| GMP_NOVERSHTERN_3___222 | Aran_et_al_2017 | CLC, CPA3, CRHBP, DNTT, FLT3, MPO, PRG2, RNASE2 |
| Hepatocytes_FANTOM_1___223 | Aran_et_al_2017 | ABAT, ACADL, ADH1A, ADH6, AFM, AGT, AGXT, AHSG, ALB, ALDOB, AMBP, ANPEP, AOX1, APCS, APOA1, APOA2, APOC1, APOC3, APOE, APOH, ARSE, ASGR1, SERPINC1, BAAT, BHMT, SERPING1, C1R, C1S, C2, C4BPA, C4BPB, C5, C8A, C8B, C8G, C9, SERPINA6, CD14, CDO1, AKR1C4, ABCC2, CPB2, CPN2, CPS1, CYP2C19, CYP2C8, CYP2C9, CYP2E1, CYP3A4, CYP3A5, CYP7A1, DAO, DIO1, DPYS, EHHADH, ENPEP, EPHX2, F2, F5, F9, F13B, FABP1, FGA, FGB, FGG, FGL1, GAS2, GATM, GC, |

| | | |
|---|---|---|
| | | GHR, GLDC, GRB14, GYS2, HABP2, SERPIND1, HGD, HHEX, HLF, HPD, HPN, HPX, HRG, IGFBP1, ITIH2, ITIH3, KCNJ8, KNG1, LECT2, LGALS4, LIPC, MAN1A1, MAOB, MAT1A, MBL2, MT1G, MT1M, MTTP, MYLK, NNMT, ORM1, OTC, PAH, SERPINA5, PCK1, ABCB1, ABCB4, SERPINA1, SERPINA4, SERPINF2, FXYD1, PON1, PPP1R3C, PROX1, PXMP2, RARRES2, RBP4, SALL1, SDC2, SLC2A2, SLC6A1, SLC10A1, SLC22A1, SPINK1, SPP1, AKR1D1, SULT2A1, TAT, SERPINA7, TDO2, TFPI, TFR2, TM4SF4, TTR, UGT2B4, UGT2B15, VIL1, VTN, ACOX2, KMO, PLA2G4C, HSD17B6, SKAP1, VNN1, NAT8, TM4SF5, PAPSS2, RGN, NR1I3, NR1H4, SLC17A2, ABCA8, UBD, SLCO1B1, SLC17A3, IQGAP2, FERMT2, SLC7A9, ANXA10, ATF5, CUX2, QPRT, BHMT2, MYRIP, PCOLCE2, GNMT, ANGPTL3, SLCO1B3, TMEM176B, SNX10, A1CF, SERPINA10, HSD17B11, HAO2, PIPOX, HAO1, FAM134B, ACSM5, SLC38A4, SLC47A1, TMEM176A, SLC30A10, APOM, GBA3, PBLD, ABCG5, ALDH8A1, TRPM8, ECHDC3, UGT2A3, AGMAT, INHBE, RUNDC3B |
| Hepatocytes_FANTOM_2___224 | Aran_et_al_2017 | ABAT, ADH1A, ADH1C, ADH6, AFM, AGT, AGXT, AHSG, ALB, ALDH1A1, ALDOB, AMBP, ANPEP, AOX1, APCS, APOA1, APOA2, APOC1, APOC3, APOE, APOH, ARSE, ASGR1, SERPINC1, BAAT, BHMT, SERPING1, C1R, C1S, C4BPA, C4BPB, C5, C8A, C8B, C8G, C9, SERPINA6, CDO1, AKR1C4, CPB2, CPS1, CRYM, CYP2C8, CYP2C9, CYP2E1, CYP3A4, CYP3A5, CYP7A1, DIO1, DPYS, EHHADH, ENPEP, F2, F5, F9, F13B, FABP1, FGA, FGB, FGG, FGL1, GAS2, GATM, GC, GHR, GRB14, GSTA1, GYS2, HABP2, SERPIND1, HGD, HHEX, HPD, HPN, HPX, HRG, IGFBP1, ITIH2, ITIH3, KNG1, LECT2, LGALS4, LIPC, MAOB, MAT1A, MBL2, MT1G, MT1M, MTTP, MYLK, NNMT, ORM1, OTC, PAH, SERPINA5, PCK1, ABCB1, ABCB4, SERPINA1, SERPINA4, SERPINF2, FXYD1, PON1, RARRES2, RBP4, SALL1, SDC2, SEPP1, SLC2A2, SLC6A1, SLC10A1, SLC22A1, SPINK1, SPP1, AKR1D1, SULT2A1, TAT, SERPINA7, TDO2, TM4SF4, TTR, UGT2B4, UGT2B15, VIL1, VTN, ACOX2, PLA2G4C, HSD17B6, SKAP1, VNN1, NAT8, PAPSS2, NR1I3, NR1H4, ABCA8, UBD, SLCO1B1, SLC17A3, IQGAP2, ANXA10, QPRT, BHMT2, MYRIP, PCOLCE2, GNMT, ANGPTL3, SLCO1B3, TMEM176B, SNX10, A1CF, PIPOX, HAO1, ACSM5, SLC38A4, TMEM176A, SLC30A10, APOM, PLSCR4, GBA3, ABCG5, ALDH8A1, TRPM8, UGT2A3, AGMAT, EFHD1, INHBE, RUNDC3B |
| Hepatocytes_FANTOM_3___225 | Aran_et_al_2017 | ACADL, ADH1A, ADH6, AFM, AGT, AGXT, AHSG, ALB, ALDOB, AMBP, ANPEP, AOX1, APCS, APOA1, APOA2, APOC1, APOC3, APOE, APOH, ARSE, ASGR1, SERPINC1, BAAT, BHMT, SERPING1, C1R, C1S, C2, C4BPA, C4BPB, C5, C8A, C8G, C9, SERPINA6, CDO1, AKR1C4, ABCC2, CPB2, CPS1, CYP2C8, CYP2C9, CYP2E1, CYP3A4, CYP3A5, DIO1, DPYS, EHHADH, ENPEP, F2, F5, F9, F13B, FABP1, FGA, FGB, FGG, FGL1, GAS2, GATM, GC, GHR, GLDC, GRB14, GYS2, HABP2, SERPIND1, HGD, HHEX, HLF, HPD, HPN, HPX, HRG, IGFBP1, ITIH2, ITIH3, KNG1, LGALS4, LIPC, MAOB, MAT1A, MBL2, MT1G, MT1M, MTTP, MYLK, NNMT, ORM1, OTC, PAH, SERPINA5, PCK1, ABCB1, ABCB4, SERPINA1, SERPINA4, SERPINF2, FXYD1, PON1, PXMP2, RARRES2, RBP4, SALL1, SDC2, SLC2A2, SLC6A1, SLC10A1, SLC22A1, SPINK1, SPP1, AKR1D1, SULT2A1, TAT, SERPINA7, TDO2, TFPI, TM4SF4, TTR, UGT2B4, UGT2B15, VIL1, VTN, ACOX2, KMO, PLA2G4C, HSD17B6, SKAP1, VNN1, NAT8, PAPSS2, NR1I3, NR1H4, ABCA8, SLCO1B1, SLC17A3, IQGAP2, ANXA10, BHMT2, MYRIP, PCOLCE2, GNMT, ANGPTL3, TMEM176B, A1CF, SERPINA10, HAO2, PIPOX, HAO1, ACSM5, SLC38A4, TMEM176A, APOM, GBA3, ABCG5, ALDH8A1, TRPM8, UGT2A3, AGMAT, INHBE, RUNDC3B |
| Hepatocytes_HPCA_1___226 | Aran_et_al_2017 | AADAC, ACADS, ACAT1, ADH1A, ADH1B, ADH1C, AFM, AHSG, ALDH2, ALDH3A2, AMBP, APOF, APCS, APOA1, APOA2, APOC3, APOE, ARG1, SERPINC1, BCHE, CEACAM1, BHMT, SERPING1, C1R, C4BPA, C5, C6, C8A, C8B, C8G, SERPINA6, CDO1, CEBPA, AKR1C4, CPB2, CPN2, CRP, CYB5A, CYP2A6, CYP2A7, CYP2C8, CYP2C9, CYP2J2, CYP3A5, DDC, DEFB1, DIO1, DPYS, EHHADH, ELF3, EPHX1, EPHX2, F2, F5, F7, F9, F10, F11, F13B, FABP1, FCN2, FGA, FGB, FGG, FGL1, G6PC, GC, GCGR, GCH1, GCKR, CXCL2, GSTA1, HAL, SERPIND1, CFH, HGFAC, HMGCS2, HP, HPX, HRG, HYAL1, IGFALS, ITIH3, KHK, KLKB1, KNG1, LBP, LCAT, LGALS4, MAN1A1, ALDH6A1, MT1F, MT1G, MT1X, ORM1, PCSK6, PCCA, PCK1, SERPINF1, SERPINA1, PLA2G2A, SERPINF2, PON1, PZP, CCL16, SLC2A2, SLC6A1, SLC7A2, SLC22A1, SPINK1, TAT, SERPINA7, UGT2B4, APOL1, HSD17B6, ABCB11, PROZ, KYNU, SELENBP1, TM4SF5, NR1H4, SLC17A4, SLC17A2, SLCO1B1, SLC22A7, GADD45G, SLC38A3, SDS, SLC27A5, SLC27A2, QPRT, HAAO, SHC2, CCDC69, ANGPTL3, SLCO1B3, NPC1L1, OSGIN1, A1CF, MLXIPL, SERPINA10, UPB1, HAO1, ACSM5, PID1, SLC38A4, C14orf105, SLC47A1, C4orf19, SLC30A10, IL17RB, OGDHL, SEMA4G, CYP4F12, AGMAT, ATF7IP2, MIA2, SEC14L4 |
| Hepatocytes_HPCA_2___227 | Aran_et_al_2017 | AADAC, ACAT1, ADH1A, ADH1C, AFM, AGT, AHSG, ALDH2, AMBP, APOF, APCS, APOA1, APOA2, APOB, APOC3, APOE, APOH, ASGR2, BAAT, BDH1, CFB, BHMT, |

| | | |
|---|---|---|
| | | SERPING1, C2, C4BPA, C4BPB, C5, C6, C8A, C8B, C8G, CAT, SERPINA6, ENTPD5, CEBPA, AKR1C4, CPB2, CPN2, CTSO, CYP2A6, CYP2B6, CYP2C8, CYP2C9, CYP2J2, CYP3A5, DAO, DDC, DIO1, DPYS, EHHADH, EPHX1, EPHX2, F2, F5, F7, F9, F10, F11, F13B, FABP1, FCN2, FGA, FGL1, NR5A2, G6PC, GC, GCGR, GCH1, GCKR, GHR, GYS2, HABP2, HAL, SERPIND1, CFH, HGFAC, HMGCS2, HRG, HSD11B1, HSD17B2, HYAL1, IGFALS, INHBC, INSR, ITIH2, ITIH3, KHK, KLKB1, KNG1, LBP, LCAT, LECT2, LGALS4, LIPC, MAN1A1, MAT1A, ALDH6A1, MT1F, MT1G, PC, PCCA, PCK1, ENPP1, SERPINA1, SERPINF2, PON1, MASP1, PZP, RET, RNASE4, CCL16, SEL1L, SLC1A1, SLC2A2, SLC10A1, SLC22A1, SPINK1, SPP2, SULT2A1, TAT, SERPINA7, TFR2, THPO, TMPRSS2, TTPA, UGT2B4, DDO, APOL1, FCN3, HSD17B6, KCNK5, ABCB11, CREG1, PROZ, TM4SF5, LPIN2, NR1I3, NR1H4, SLC17A4, SLC25A13, SEC23A, UBD, SLCO1B1, SLC26A1, SLC22A7, SLC27A5, SLC27A2, SLC7A9, SEPHS2, SLC35D1, HAAO, BHMT2, GNMT, ANGPTL3, SLCO1B3, NPC1L1, A1CF, MLXIPL, SERPINA10, DCXR, UPB1, HAO1, ACSM5, PID1, SLC38A4, SLC47A1, C4orf19, SLC35C1, SLC30A10, IL17RB, RAB20, OGDHL, CYP4F11, GREM2, ALDH8A1, CYP4F12, COLEC11, GRTP1, AGMAT, ATF7IP2, COL18A1, CFHR5, INHBE, MIA2, SEC14L4 |
| Hepatocytes_HPCA_3___228 | Aran_et_al_2017 | AADAC, ACAT1, ADH1A, ADH1B, ADH1C, AFM, AHSG, ALDH2, ALDH3A2, AMBP, APOF, APCS, APOA1, APOA2, APOC3, APOE, ARG1, SERPINC1, CEACAM1, BHMT, SERPING1, C1R, C4BPA, C5, C6, C8A, C8B, C8G, SERPINA6, CDO1, CEBPA, AKR1C4, CPB2, CPN2, CRP, CYB5A, CYP2A6, CYP2A7, CYP2C8, CYP2C9, CYP2J2, CYP3A5, DDC, DEFB1, DIO1, DPYS, EHHADH, ELF3, EPHX1, EPHX2, F2, F5, F7, F9, F10, F11, F13B, FABP1, FCN2, FGA, FGB, FGG, FGL1, G6PC, GC, GCGR, GCH1, GCKR, CXCL2, GSTA1, HAL, SERPIND1, CFH, HGFAC, HMGCS2, HP, HPX, HRG, HYAL1, IGFALS, ITIH3, KHK, KLKB1, KNG1, LBP, LCAT, LGALS4, MAN1A1, MT1F, MT1G, MT1X, ORM1, PCSK6, PCK1, SERPINA1, PLA2G2A, SERPINF2, PON1, PZP, CCL16, SLC2A2, SLC6A1, SLC7A2, SLC22A1, SPINK1, TAT, SERPINA7, UGT2B4, APOL1, HSD17B6, ABCB11, PROZ, KYNU, TM4SF5, NR1H4, SLC17A4, SLC17A2, SLCO1B1, SLC22A7, GADD45G, SLC38A3, SLC27A5, SLC27A2, HAAO, SHC2, CCDC69, ANGPTL3, SLCO1B3, NPC1L1, OSGIN1, A1CF, MLXIPL, SERPINA10, UPB1, HAO1, ACSM5, SLC38A4, C14orf105, SLC47A1, C4orf19, SLC30A10, IL17RB, OGDHL, CYP4F12, AGMAT, MIA2, SEC14L4 |
| HSC_BLUEPRINT_1___229 | Aran_et_al_2017 | CD34, CRHBP, ERG, GSTM5, PLS3, MMRN1, LAPTM4B, FAM124B |
| HSC_BLUEPRINT_2___230 | Aran_et_al_2017 | CD34, CRHBP, ERG, FLT3, GSTM5, PLS3, SLC4A1, SCARF1, ZMYM3, MMRN1, XPO7, EXD2, LAPTM4B, KLHL9, LSM2, CCDC121, FAM124B, ATP8B4, TCEAL4, MYCT1 |
| HSC_BLUEPRINT_3___231 | Aran_et_al_2017 | ALAS2, CA1, CD34, CRHBP, ERG, FLT3, GSTM5, GYPA, GYPE, PLS3, SLC4A1, SCARF1, MMRN1, AHSP, EXD2, LAPTM4B, KLHL9, FAM124B, ATP8B4, TCEAL4 |
| HSC_FANTOM_1___232 | Aran_et_al_2017 | CRHBP, CRYGD, ELN, ERG, KCNJ13, MPL, PDE6G, PLA2G1B, SYPL1, ZNF3, RNF7, LECT1, CTNNA3, PHF20, EMCN, MYCT1 |
| HSC_FANTOM_2___233 | Aran_et_al_2017 | CRHBP, ELN, GSTM5, PLA2G1B, SYPL1, LECT1, CTNNA3, EMCN |
| HSC_FANTOM_3___234 | Aran_et_al_2017 | CRHBP, ELN, GSTM5, SYPL1, CLEC3B, LECT1, CTNNA3, EMCN |
| HSC_NOVERSHTERN_1___235 | Aran_et_al_2017 | CRHBP, ELN, GSTM5, PDE6G, SYPL1, LECT1, CTNNA3, EMCN, VN1R1 |
| HSC_NOVERSHTERN_2___236 | Aran_et_al_2017 | CRHBP, ELN, GSTM5, SYPL1, CLEC3B, LECT1, CTNNA3, EMCN |
| HSC_NOVERSHTERN_3___237 | Aran_et_al_2017 | CRHBP, CRYGD, ELN, PDE6G, SYPL1, LECT1, CTNNA3, PHF20, EMCN |
| iDC_HPCA_1___238 | Aran_et_al_2017 | ALOX15, F13A1, FCER2, IL3RA, CCL13, CCL17, CCL18, CCL23, CCL24, ALDH1A2, CD209 |
| iDC_HPCA_2___239 | Aran_et_al_2017 | ALOX15, CD86, FCER2, IL3RA, CCL13, CCL17, CCL24, ALDH1A2, CLEC10A, SPINT2 |
| iDC_HPCA_3___240 | Aran_et_al_2017 | ALOX15, F13A1, CCL13, CCL17, CCL18, CCL23, CCL24, CD209 |
| Keratinocytes_ENCODE_1___241 | Aran_et_al_2017 | CALML3, CSTA, DSG3, F3, GJB5, SFN, IL1A, JAG2, KRT5, KRT6B, LAD1, PI3, SERPINB5, SOX15, SULT2B1, LY6D, FGFBP1, AP1M2, FST, HES2, TMEM40, RAB25, S100A14, MMP28, ZBED2, KRT6C |
| Keratinocytes_ENCODE_2___242 | Aran_et_al_2017 | CALML3, CSTA, DSG3, GJB5, SFN, IL1A, KRT6B, PI3, SOX15, FGFBP1, AP1M2, FST, HES2, TMEM40, RAB25, S100A14, ZBED2, KRT6C |
| Keratinocytes_ENCODE_3___243 | Aran_et_al_2017 | DSG3, GJB5, SFN, IL1A, KRT6B, SOX15, FGFBP1, AP1M2, TMEM40, S100A14 |

| | | |
|---|---|---|
| Keratinocytes_FANTOM_1___244 | Aran_et_al_2017 | ADAM8, AIM1, ALDH1A3, CALML3, CDH3, COL17A1, CSTA, CYP27B1, DSC3, DSG3, EPHA1, EREG, F3, FAT2, FGFR3, GJB3, GJB5, GNA15, SFN, CXCL3, IL1A, IL1B, ITGA6, ITGB4, ITGB6, JAG2, KCNJ15, KRT5, KRT6A, KRT6B, KRT14, KRT16, KRT17, LAD1, LAMA3, LAMB3, LAMC2, MMP9, NGFR, SERPINB2, PGF, SERPINB5, PKP1, FXYD3, PPL, PRSS8, PTHLH, S100A2, S100A8, S100A9, SDC1, SORL1, SOX15, SPRR1B, ST14, TGFA, XDH, TP63, ARTN, CLCA2, PLCH2, KIAA0040, FGFBP1, SH2D3A, AP1M2, TSPAN1, MPZL2, NDRG1, FST, ST6GALNAC2, PPP1R13L, PKP3, KLK8, LPAR3, CBLC, TRIM29, KLK5, G0S2, DEF6, IRX4, ANGPTL4, LSR, GPR87, HES2, ESRP1, EPN3, TMEM40, FERMT1, SLC2A9, RAB25, S100A14, DLK2, C1orf116, MMP28, ZBED2, ELMO3, TNS4, ALS2CL |
| Keratinocytes_FANTOM_2___245 | Aran_et_al_2017 | ADAM8, COL17A1, CSTA, CYP27B1, DSC3, DSG3, EREG, FAT2, GJB3, GJB5, GNA15, SFN, IL1A, IL1B, ITGB4, KRT5, KRT6A, KRT6B, KRT14, KRT16, KRT17, LAD1, LAMA3, LAMB3, MMP9, SERPINB5, PKP1, FXYD3, PRSS8, S100A2, S100A8, S100A9, SOX15, ST14, TP63, ARTN, CLCA2, PLCH2, FGFBP1, SH2D3A, AP1M2, MPZL2, FST, PPP1R13L, PKP3, KLK8, CBLC, TRIM29, KLK5, G0S2, ANGPTL4, LSR, GPR87, ESRP1, EPN3, FERMT1, SLC2A9, RAB25, S100A14, C1orf116, ZBED2, ELMO3, TNS4 |
| Keratinocytes_FANTOM_3___246 | Aran_et_al_2017 | ADAM8, CALML3, COL17A1, CSTA, CYP27B1, DSC3, DSG3, EREG, FAT2, GJB3, GJB5, GNA15, SFN, IL1A, IL1B, ITGB4, KRT5, KRT6A, KRT6B, KRT14, KRT16, KRT17, LAD1, LAMA3, LAMB3, LAMC2, MMP9, SERPINB5, PKP1, FXYD3, PRSS8, S100A2, S100A8, S100A9, SOX15, ST14, TP63, ARTN, CLCA2, PLCH2, FGFBP1, SH2D3A, AP1M2, MPZL2, FST, PPP1R13L, PKP3, KLK8, CBLC, TRIM29, KLK5, G0S2, ANGPTL4, LSR, GPR87, ESRP1, EPN3, FERMT1, SLC2A9, RAB25, S100A14, C1orf116, ZBED2, ELMO3, TNS4 |
| Keratinocytes_HPCA_1___247 | Aran_et_al_2017 | DSG3, IL1A, KRT5, KRT14, SERPINB5, PTK6, SULT2B1, LY6D, CLCA2, FGFBP1, PRMT5, PKP3, FLRT3, TP53AIP1, DLK2, MMP28, GJB4, KRT6C |
| Keratinocytes_HPCA_2___248 | Aran_et_al_2017 | BDKRB2, CALML3, CSTA, DSC3, DSG3, F3, GJB3, GJB5, SFN, IL1A, IRF6, JAG2, KRT5, KRT6B, KRT14, LAD1, PGF, PI3, SERPINB5, PKP1, PLA2G4A, PRSS8, PTK6, SLC6A11, SLC12A4, SOX15, SULT2B1, CORO2A, LY6D, CLCA2, FGFBP1, AP1M2, CNKSR1, B3GNT3, PRMT5, FST, DUSP14, LPAR3, FLRT3, TFCP2L1, HES2, TMEM40, FGD6, LTB4R2, RAB25, S100A14, DLK2, C1orf116, MMP28, ZBED2, ZNF750, GJB4, KRT6C |
| Keratinocytes_HPCA_3___249 | Aran_et_al_2017 | BDKRB2, CALML3, CSTA, DSC3, DSG3, F3, GJB3, GJB5, SFN, IL1A, IRF6, JAG2, KRT5, KRT6B, KRT14, LAD1, MST1R, PI3, SERPINB5, PKP1, PTK6, SOX15, SULT2B1, CORO2A, LY6D, CLCA2, FGFBP1, AP1M2, CNKSR1, B3GNT3, PRMT5, FST, LPAR3, TFCP2L1, HES2, TMEM40, FGD6, LTB4R2, RAB25, S100A14, C1orf116, MMP28, ZBED2, ZNF750, GJB4, KRT6C |
| ly_Endothelial_cells_FANTOM_1___250 | Aran_et_al_2017 | FLT4, HYAL2, CLEC1A, SOX18, ROBO4, CXorf36, MYCT1, KANK3 |
| ly_Endothelial_cells_FANTOM_2___251 | Aran_et_al_2017 | ACVRL1, ANGPT2, RHOC, BMX, CETP, ERG, FLT4, FUS, GPR4, KDR, MYL2, NOTCH4, RALA, MAPK12, TEK, TIE1, CLDN5, HYAL2, TNFSF18, KALRN, FEZ2, SEMA6B, LYVE1, ARHGEF15, MMRN1, CD93, N4BP3, CLEC1A, EMCN, SOX18, ROBO4, RASIP1, TMEM39B, POMGNT1, MRPL17, CXorf36, MMRN2, MYCT1, NETO2, PLVAP, KANK3 |
| ly_Endothelial_cells_FANTOM_3___252 | Aran_et_al_2017 | ANGPT2, FLT4, MAPK12, TEK, HYAL2, KALRN, FEZ2, CLEC1A, SOX18, ROBO4, CXorf36, MYCT1, KANK3 |
| ly_Endothelial_cells_HPCA_1___253 | Aran_et_al_2017 | GJA4, SELE, TIE1, CLDN5, VWF, HYAL2, TNFSF18, ARHGEF15, CLEC1A, ROBO4, MMRN2, KANK3 |
| ly_Endothelial_cells_HPCA_2___254 | Aran_et_al_2017 | GJA4, SELE, TIE1, CLDN5, VWF, TNFSF18, ROBO4, MMRN2, KANK3 |
| ly_Endothelial_cells_HPCA_3___255 | Aran_et_al_2017 | FLT4, GJA4, SELE, TIE1, CLDN5, TPM3, VWF, HYAL2, TNFSF18, ARHGEF15, CLEC1A, ROBO4, MMRN2, KANK3 |
| Macrophages_M1_BLUEPRINT_1___256 | Aran_et_al_2017 | ACP2, ABCD1, C1QA, FDX1, CCL22, CD163, SCAMP2, ADAMDEC1, ARL8B, HAMP |
| Macrophages_M1_BLUEPRINT_2___257 | Aran_et_al_2017 | ACP2, ABCD1, FDX1, CCL8, CCL22, CD163, ADAMDEC1, TREM2, HAMP |
| Macrophages_M1_BLUEPRINT_3___258 | Aran_et_al_2017 | ACP2, ADRA2B, ALCAM, ABCD1, ATOX1, ATP6V0C, ATP6V1E1, BLVRA, C1QA, CD48, CD63, CLCN7, TPP1, CLTC, CCR1, CMKLR1, SLC31A1, COX5B, FCER1G, FDX1, FOLR2, FPR3, FTL, HEXB, HK3, IL10, IL12B, ITGAE, LAIR1, CXCL9, MMP19, NARS, NDUFS2, P2RX7, PDCL, MAPK13, PTGIR, PTPRA, RELA, CCL7, CCL8, CCL19, CCL22, SRC, STX4, TCEB1, TFRC, AGPS, MARCO, SNX3, CD84, USP14, ITGB1BP1, ATP6V1F, TRIP4, CD163, CIAO1, WTAP, ARHGEF11, ABI1, SCAMP2, ACTR2, BCAP31, ZMPSTE24, BCKDK, EXOC5, STIP1, UQCR11, SDS, LILRB4, OGFR, TFEC, FKBP15, DNAJC13, TDRD7, STX12, IL17RA, ABTB2, FAM32A, SIGLEC7, SIGLEC9, ADAMDEC1, |

| | | |
|---|---|---|
| | | CECR5, SLC25A24, NRBP1, MS4A4A, TREM2, OTUD4, PQLC2, HAUS2, ARL8B, NECAP2, WDR11, ZC3H15, CCDC47, UTP3, MRS2, HAMP, MRPL40, VPS33A, CORO7, LIMD2, TMX1, DOT1L, ADO, ADCK2 |
| Macrophages_M1_FANTOM_1___259 | Aran_et_al_2017 | ACP2, ADRA2B, ALCAM, TSPO, C3AR1, DAGLA, CALR, CHIT1, CYBB, CYC1, CYP19A1, DLAT, FCER1G, GP1BA, GPD1, IFNAR1, IL10, KCNJ5, KIFC3, MT2A, MYBPH, MYH11, MYO7A, P2RX7, PRDX1, RAB3IL1, RNH1, MRPL12, CCL1, CCL7, CCL8, CCL24, SRC, VIM, RRP1, MARCO, S1PR2, AP1M2, ACTR3, LILRB1, AFG3L2, SDS, LILRB4, EMILIN1, VSIG4, HSPB7, COQ2, ADAMDEC1, CECR5, WSB2, SLAMF8, DNASE2B, CLPB, MFSD7, ADCK2 |
| Macrophages_M1_FANTOM_2___260 | Aran_et_al_2017 | ACP2, ADCY3, ADRA2B, ALCAM, TSPO, C1QA, C1QB, C3AR1, DAGLA, CD63, CHIT1, CMKLR1, SLC31A1, CSF1, CSF1R, CYBB, CYC1, CYP19A1, FANCE, FCER1G, FDX1, FPR3, FTL, GP1BA, GPD1, HEXB, IL10, KCNJ1, KCNJ5, KIFC3, LAMP1, MMP19, MSR1, MT2A, MYBPH, MYO7A, P2RX7, PRDX1, RAB3IL1, MRPL12, CCL1, CCL7, CCL8, CCL18, CCL19, CCL24, SLC6A12, SPR, SRC, RRP1, MARCO, PKD2L1, S1PR2, CD163, LONP1, AP1M2, IGSF6, LILRB1, SDS, LILRB4, EMILIN1, VSIG4, TFEC, PHLDB1, CYFIP1, FKBP15, NCAPH, MYOF, HSPB7, ADAMDEC1, GLRX2, NDUFAF1, SPG21, MS4A4A, ATP6V1D, ATP6V1H, TREM2, PQLC2, TMEM70, PLEKHB2, TMEM33, SLAMF8, HAMP, DNASE2B, MYOZ1, LONRF3, CLPB, MFSD7, ADCK2 |
| Macrophages_M1_FANTOM_3___261 | Aran_et_al_2017 | ACP2, ADCY3, ADRA2B, ALCAM, ABCD1, ANXA2, ATP6V1A, C1QA, C1QB, C3AR1, DAGLA, CD80, CD63, CHIT1, CMKLR1, SLC31A1, CSF1, CSF1R, CYBB, CYC1, CYP19A1, FANCE, FDX1, FPR2, FPR3, GPD1, HEXB, KCNJ1, KCNJ5, KIFC3, MMP19, MSR1, MT2A, MYBPH, P2RX7, MAPK13, S100A11, CCL1, CCL7, CCL8, CCL18, CCL19, CCL22, CCL24, SLC1A2, SLC6A12, SLC11A1, SIGLEC1, SRC, TIE1, MARCO, HYAL2, CD163, LONP1, IGSF6, LILRB1, CD300C, SDS, LILRB4, EMILIN1, VSIG4, PHLDB1, NCAPH, CLEC4E, MYOF, HSPB7, ADAMDEC1, GLRX2, MS4A4A, ATP6V1H, TREM2, TMEM70, TMEM33, KCNK13, SLAMF8, HAMP, DNASE2B, MYOZ1, MFSD7, ADO, ADCK2, TBC1D16 |
| Macrophages_M2_BLUEPRINT_1___262 | Aran_et_al_2017 | ACP2, ADCY3, ABCD1, ALK, ARSB, ATP2A2, ATP6V1C1, ATP6V0A1, TSPO, CAMP, CANX, CD63, CD81, CLCN7, TPP1, SLC31A1, FDX1, FGR, FTL, GLB1, HADHB, NCKAP1L, HEXA, HEXB, HPS1, IFNAR1, ITGAX, KCNJ5, LAIR1, LAMP1, MSR1, MYO9B, P2RX7, SDCBP, SNX1, SNX2, STX4, MARCO, CDS2, PABPC4, ATP6V0D1, PICK1, ARHGEF11, HS3ST2, PDCD6IP, SCAMP2, COL4A3BP, HSPH1, OS9, SDS, LILRB4, VSIG4, GABARAP, TFEC, WDFY3, TBC1D9B, ZC3H3, CYFIP1, PLEKHM2, FKBP15, SMG5, UNC50, GGA1, SNX5, SLC39A1, ADAMDEC1, COMMD9, SLC25A24, SPG21, MS4A4A, ANKFY1, BTBD1, STX18, RIN2, PQLC2, TMEM70, ACSM5, AGGF1, SLC38A7, VPS53, NOP10, IARS2, CCDC88A, VPS35, TMEM184C, EXOC1, SLAMF8, C16orf62, POGK, HAMP, DNASE2B, MTMR14, GORASP1, C10orf76, LONRF3, UBXN6 |
| Macrophages_M2_BLUEPRINT_2___263 | Aran_et_al_2017 | CLCN7, FGR, GLB1, HEXA, HEXB, HS3ST2, FKBP15, PQLC2, TMEM70, SLC38A7 |
| Macrophages_M2_BLUEPRINT_3___264 | Aran_et_al_2017 | ACP2, ALK, ARSB, ATP6V0A1, CD63, CLCN7, TPP1, SLC31A1, FGR, GLB1, HADHB, NCKAP1L, HEXA, HEXB, IFNAR1, KCNJ5, MYO9B, P2RX7, SDCBP, MARCO, ATP6V0D1, HS3ST2, PDCD6IP, COL4A3BP, CYFIP1, PLEKHM2, FKBP15, SMG5, SLC39A1, COMMD9, MS4A4A, STX18, PQLC2, TMEM70, SLC38A7, VPS53, HAMP, LONRF3 |
| Macrophages_M2_HPCA_1___265 | Aran_et_al_2017 | ADRA2B, AP1B1, ALDH9A1, ANXA11, AQP8, CD52, ELK1, FH, FLT1, GPD1, NCKAP1L, HEXB, KCNJ1, LAMP1, MMP19, MSR1, NDUFB1, NPR1, PDE1B, PEX19, S100A6, SLC6A7, SLC6A12, SNAPC2, SNX1, TAF10, UCP3, UGP2, USF2, XPNPEP2, AKR7A2, SNX3, TNFSF14, NFS1, GSTO1, KIAA0196, HS3ST2, BCAP31, CEPT1, BAIAP2, SLC9A6, VTI1B, ARFGEF2, HSPH1, LILRA2, EFR3A, FKBP15, NCAPH, ZCCHC4, TMED5, MYO15A, NAGPA, ZNF219, MS4A4A, ANGPT4, ATP6V1D, TREM2, KCTD5, PQLC2, AGGF1, MFN1, IARS2, CCDC88A, KCNK13, TMEM9B, POGK, MYOZ1, IPPK, CARD14, ALG9, MRM1, DHX57, SLC25A46, OSBPL11, CCDC85C |
| Macrophages_M2_HPCA_2___266 | Aran_et_al_2017 | ADRA2B, DNASE1L3, FDX1, GPD1, GUCA1A, KCNJ1, MSR1, PDE1B, UCP3, HS3ST2, SDS, MS4A4A, TREM2, DNASE2B, MYOZ1 |
| Macrophages_M2_HPCA_3___267 | Aran_et_al_2017 | ADRA2B, CD52, GPD1, MSR1, NPR1, UCP3, HS3ST2, MYO15A |
| Macrophages_BLUEPRINT_1___268 | Aran_et_al_2017 | ACP2, ATOX1, ATP6V0C, ATP6V1E1, C1QA, CD9, CD48, CLCN7, CCR1, CRYBB1, CYBB, FCER1G, FDX1, FOLR2, FPR3, FTL, HEXA, HEXB, KCNJ5, LAIR1, M6PR, MDH1, NUBP1, P2RX7, RAC1, CCL8, CCL22, STX4, TCEB1, TYROBP, VAMP8, MARCO, CD84, ATP6V0E1, ATP6V1F, CD163, CIAO1, LY86, ARHGEF11, HS3ST2, ARPC4, |

| | | |
|---|---|---|
| | | ATP6AP2, BAIAP2, PRDX3, ERP29, LILRB5, SDS, LILRB4, VSIG4, FKBP15, ZZZ3, SIGLEC7, SIGLEC9, ADAMDEC1, ORMDL2, COMMD9, SPG21, MS4A4A, UBE2D4, TRAPPC2L, TREM2, TMEM70, ARL8B, TMEM126B, SLAMF8, C12orf4, HAMP, GUF1, YIF1B |
| Macrophages_BLUEPRINT_2___269 | Aran_et_al_2017 | ACP2, ATOX1, ATP6V0C, ATP6V1E1, C1QA, CD9, CD48, CD63, CLCN7, TPP1, CCR1, CMKLR1, SLC31A1, COX5B, COX7B, COX8A, CRYBB1, CYBA, CYBB, FCER1G, FDX1, FOLR2, FPR3, FTL, GLB1, HEXA, HEXB, KCNJ5, LAIR1, LAMP1, NUBP1, NDUFB3, NDUFS3, NDUFS6, P2RX7, PCMT1, MAPK13, PSME1, RB1, CCL8, CCL22, SDHD, SUMO3, SNX2, TCEB1, TYROBP, UQCRC2, VAMP8, MARCO, SNX3, CD84, ATP6V0E1, ITGB1BP1, S1PR2, ATP6V1F, CD163, COX5A, CIAO1, LY86, ARHGEF11, HS3ST2, SCAMP2, ARPC4, ATP6AP2, BAIAP2, PRDX3, ERP29, UQCR11, LILRB5, SDS, LILRB4, VSIG4, STAB1, FKBP15, DNAJC13, CLEC5A, ZZZ3, SIGLEC7, SIGLEC9, ADAMDEC1, ORMDL2, COMMD9, UQCR10, SPG21, MS4A4A, UBE2D4, TRAPPC2L, TREM2, KCTD5, PQLC2, COMMD8, TMEM70, ARL8B, SLC38A7, NOP10, WDR11, TMEM126B, TMEM9B, SLAMF8, C12orf4, MRS2, HAMP, DNASE2B, GUF1, MS4A6A, MRPL40, PPCS, PMFBP1, YIF1B, ADCK2, HIGD2A |
| Macrophages_BLUEPRINT_3___270 | Aran_et_al_2017 | ACP2, ATOX1, ATP6V0C, ATP6V1E1, C1QA, CD9, CD48, CD63, CLCN7, CCR1, COX8A, CRYBB1, CYBB, FCER1G, FDX1, FOLR2, FPR3, FTL, HEXA, HEXB, KCNJ5, LAIR1, LAMP1, NDUFB3, NDUFS6, P2RX7, MAPK13, CCL8, CCL22, SNX2, TCEB1, TYROBP, UQCRC2, VAMP8, MARCO, SNX3, CD84, ATP6V0E1, ITGB1BP1, ATP6V1F, CD163, COX5A, CIAO1, LY86, ARHGEF11, HS3ST2, SCAMP2, ARPC4, ATP6AP2, ERP29, UQCR11, LILRB5, SDS, LILRB4, VSIG4, STAB1, FKBP15, DNAJC13, CLEC5A, ZZZ3, SIGLEC7, SIGLEC9, ADAMDEC1, ORMDL2, COMMD9, SPG21, MS4A4A, UBE2D4, TREM2, KCTD5, TMEM70, ARL8B, NOP10, WDR11, TMEM126B, SLAMF8, C12orf4, HAMP, GUF1, MS4A6A, MRPL40, PPCS, PMFBP1, YIF1B, ADCK2 |
| Macrophages_FANTOM_1___271 | Aran_et_al_2017 | ACP2, CHIT1, CSF1, CYP19A1, FDX1, HK3, MSR1, CCL22, SLC6A12, CD84, SDS, VSIG4, CLEC5A, ADAMDEC1, HAMP, DNASE2B, MYOZ1 |
| Macrophages_FANTOM_2___272 | Aran_et_al_2017 | ACP2, ADCY3, ALCAM, ABCD1, BPI, CHIT1, CYP19A1, HK3, KCNJ1, KCNMB1, CXCL9, MMP8, MSR1, CCL1, CCL7, CCL22, SLC1A2, SLC6A12, CD84, CD163, SDS, LILRB4, VSIG4, FKBP15, NCAPH, CLEC5A, ADAMDEC1, ATP6V1H, PQLC2, SLAMF8, HAMP, DNASE2B, MYOZ1, LONRF3, MFSD7 |
| Macrophages_FANTOM_3___273 | Aran_et_al_2017 | ACP2, ADCY3, ALCAM, ABCD1, ATP6V1A, ATP6V0A1, BPI, CHIT1, SLC31A1, CSF1, CYBB, CYP19A1, FDX1, FGR, HK3, ITGAX, KCNJ1, KCNMB1, CXCL9, MMP8, MMP19, MSR1, P2RX7, MAPK13, CCL1, CCL7, CCL22, SLC1A2, SLC6A12, SLC11A1, STX4, NUMB, MARCO, CD164, CD84, CD163, CIR1, ARHGEF11, BCAP31, ATP6AP2, SDS, LILRB4, VSIG4, FKBP15, NCAPH, CLEC5A, ADAMDEC1, ATP6V1H, PQLC2, CCDC88A, PCDHB11, SLAMF8, HAMP, DNASE2B, MYOZ1, LONRF3, MFSD7 |
| Macrophages_HPCA_1___274 | Aran_et_al_2017 | ACADVL, ACP2, ARSB, ATP6V1A, ATP6V0C, ATP6V1C1, CD63, CETN2, CCR1, SLC31A1, COX5B, COX15, CYBB, DBI, ECHS1, FDX1, GRB2, HADHB, HEXA, HEXB, HK3, HMGCL, ITGAX, KIFC3, TNPO1, LAIR1, LAMP1, MGST3, MSR1, NARS, NDUFA8, NDUFB6, NDUFS8, PRDX1, PEX14, MAPK13, PSMD10, PTPN12, PEX19, QDPR, RAB1A, RAB5C, RALA, RENBP, CLIP1, CCL7, CCL18, SDHB, SRC, STX4, TCEB1, MLX, TRAF3, NSMAF, AGPS, MARCO, SNX4, ATP6V1F, LONP1, GSTO1, BAG3, CIR1, FEZ2, PDCD6IP, SNUPN, BCAP31, STAM2, IGSF6, ZMPSTE24, TMEM147, VTI1B, TGOLN2, SPIN1, LILRB4, EMILIN1, VSIG4, EFR3A, FKBP15, CLEC5A, IBTK, NPTN, ATP2C1, SIGLEC9, ADAMDEC1, CNIH4, GLRX2, DERA, NDUFAF1, HSD17B12, ZDHHC3, TNFRSF12A, MS4A4A, ATP6V1D, ATP6V1H, TMBIM4, STYXL1, BTBD1, NUDT9, TMEM33, NOP10, TMEM127, ACTR10, KCMF1, TULP4, C12orf4, RTN4, MKL2, HAMP, DNASE2B, KLHL12, NSUN3, ELOVL1, SLC30A5, MAPKAP1, LONRF3, TCEAL4, CHD9, TM2D1, MFSD7, G6PC3, TBC1D16, 44084, ZDHHC24 |
| Macrophages_HPCA_2___275 | Aran_et_al_2017 | ACP2, ARSB, ATP6V1A, ATP6V0C, ATP6V1C1, CD63, CETN2, CCR1, SLC31A1, COX5B, COX15, CYBB, DBI, ECHS1, FDX1, HEXB, LAIR1, LAMP1, MGST3, MSR1, NDUFA8, NDUFS8, PRDX1, MAPK13, PSMD10, PTPN12, PEX19, RAB1A, RALA, RENBP, CCL7, SDHB, STX4, TCEB1, MLX, TRAF3, NSMAF, AGPS, MARCO, ATP6V1F, GSTO1, BCAP31, IGSF6, TMEM147, VTI1B, LILRB4, EMILIN1, VSIG4, EFR3A, CLEC5A, NPTN, ATP2C1, SIGLEC9, ADAMDEC1, CNIH4, GLRX2, DERA, NDUFAF1, ZDHHC3, TNFRSF12A, MS4A4A, ATP6V1D, ATP6V1H, NUDT9, TMEM33, C12orf4, RTN4, HAMP, DNASE2B, ELOVL1, SLC30A5, MAPKAP1, LONRF3, MFSD7, G6PC3, TBC1D16, 44084, ZDHHC24 |
| Macrophages_HPCA_3___276 | Aran_et_al_2017 | ACADVL, ACP2, ARSB, ATP6V1A, ATP6V0C, ATP6V1C1, CD63, CETN2, CCR1, SLC31A1, COX5B, COX15, CYBB, DBI, ECHS1, FDX1, GRB2, HADHB, HCCS, HEXA, HEXB, HK3, HMGCL, ITGAX, TNPO1, LAIR1, LAMP1, MGST3, MSR1, NDUFA8, NDUFS8, PRDX1, MAPK13, PSMD10, PTPN12, PEX19, QDPR, RAB1A, RALA, RENBP, CLIP1, |

| | | |
|---|---|---|
| | | CCL7, SDHB, SRC, STX4, TCEB1, MLX, TRAF3, UQCRC2, USF2, NSMAF, AGPS, MARCO, SNX4, ATP6V1F, GSTO1, PDCD6IP, BCAP31, STAM2, IGSF6, ZMPSTE24, CEPT1, TMEM147, VTI1B, TGOLN2, SPIN1, LILRB4, TMEM115, EMILIN1, VSIG4, EFR3A, FKBP15, CLEC5A, NPTN, ATP2C1, TRAPPC3, SIGLEC9, ADAMDEC1, CNIH4, SLC25A24, GLRX2, DERA, NDUFAF1, ZDHHC3, TNFRSF12A, MS4A4A, ATP6V1D, ATP6V1H, BTBD1, NUDT9, TMEM33, TMEM127, ACTR10, KCMF1, TULP4, C12orf4, RTN4, MKL2, HAMP, DNASE2B, NSUN3, ELOVL1, SLC30A5, C7orf25, MAPKAP1, PANK3, LONRF3, TM2D1, MFSD7, SETD3, G6PC3, TBC1D16, 44084, ZDHHC24 |
| Macrophages_IRIS_1___277 | Aran_et_al_2017 | ACP2, ADCY3, ABCD1, ATP6V1A, CHIT1, CLCN7, TPP1, SLC31A1, COX5B, CYP19A1, FDX1, HEXA, HK3, KCNJ1, M6PR, MSR1, SLC6A12, CD164, CD84, LONP1, ATP6AP2, TGOLN2, SDS, LILRB4, VSIG4, FKBP15, CLEC5A, ADAMDEC1, ATP6V1H, STX18, PQLC2, DNASE2B, MYOZ1, SLC30A5, LONRF3, MFSD7, LDHAL6B |
| Macrophages_IRIS_2___278 | Aran_et_al_2017 | ACP2, ADCY3, ALCAM, ABCD1, ARSB, ATOX1, ATP6V1A, ATP6V0C, ATP6V1E1, ATP6V0A1, BPI, TSPO, CD63, CHIT1, CLCN7, TPP1, SLC31A1, COX5B, CSF1, CYBB, CYP19A1, FDX1, HTT, HEXA, HK3, ITGAX, KCNJ1, LAMP1, M6PR, MMP19, MSR1, MTHFR, P2RX7, MAPK13, PTPRA, RABGGTA, SLC1A2, SLC6A12, STX4, TCEB1, NUMB, CD164, CD84, LONP1, CPNE6, CIR1, TTLL4, ARHGEF11, BCAP31, ATP6AP2, TGOLN2, SDS, LILRB4, VSIG4, FKBP15, CLEC5A, IL17RA, ADAMDEC1, NRBP1, SH3GLB1, ATP6V1H, STX18, PQLC2, TMEM33, SLC38A7, VPS53, CCDC88A, SLAMF8, HAMP, DNASE2B, MYOZ1, MTMR14, SLC30A5, MUL1, C12orf49, LONRF3, MFSD7, LDHAL6B, ZDHHC24 |
| Macrophages_IRIS_3___279 | Aran_et_al_2017 | ACP2, CHIT1, CSF1, CYP19A1, FDX1, HK3, MSR1, CCL22, SLC6A12, CD84, SDS, VSIG4, CLEC5A, ADAMDEC1, HAMP, DNASE2B, MYOZ1 |
| Mast_cells_FANTOM_1___280 | Aran_et_al_2017 | AMHR2, ANXA1, ANXA11, ATP6V1C1, BMPR1A, BTK, C3AR1, C8G, CASP10, CD22, CD33, SIGLEC6, CMA1, CPA3, CTSG, DIAPH1, DR1, MS4A2, GATA1, HDC, IL5, IL5RA, ITGA2B, KCNJ5, KRT1, LCP2, LTC4S, LYL1, MTR, NTRK1, OSBP, P2RX1, PAK2, PDE4A, PIK3R2, POLR2A, PPP3R1, PRG2, PRKAR1A, PTGDR, PTGER3, RAD23B, RENBP, RGS13, RXRB, SLC18A2, SNRNP70, SOS2, SYPL1, TADA2A, MAP3K7, TAL1, TEC, TPSAB1, RNF103, ZNF212, STAM, NDST2, AGPS, PABPC4, RGS11, CD84, CDC16, USP10, ZMYM4, WDR46, ZNF264, CTR9, DEPDC5, LRIG2, THRAP3, SNUPN, KLRG1, BAIAP2, HSPH1, SERINC3, IFT27, ATXN2L, U2AF2, ZBTB1, BAHD1, TRIM32, FAM120A, DNAJC13, ESYT1, ARHGEF12, ATXN10, UPF2, SPAG8, STAP1, HIBCH, SIGLEC7, SNX5, SIGLEC8, HPGDS, REM1, MRPS28, CPSF1, BET1L, ZBTB7A, NBAS, LAX1, DNAJC28, AGGF1, BTBD7, TTC17, CENPJ, ZNF471, CTDSP1, HRH4, ACBD3, DCLRE1B, ZNF426, MBOAT7, ZMYM1, FBXO11, CHD9, ZKSCAN3, C6orf25, FBXO38, MAGT1, USP48, OSBPL9, ZNF549, TIPRL |
| Mast_cells_FANTOM_2___281 | Aran_et_al_2017 | AMHR2, ANXA1, ANXA11, ATP6V1C1, BMPR1A, BTK, C3AR1, C8G, CASP10, CD22, SIGLEC6, CPA3, CTSG, DIAPH1, DR1, MS4A2, GATA1, HDC, IL5, ITGA2B, KCNJ5, KRT1, LCP2, LTC4S, LYL1, MTR, NTRK1, OSBP, P2RX1, PAK2, PIK3R2, POLR2A, PPP3R1, PRG2, PRKAR1A, PTGER3, RENBP, RGS13, RXRB, SLC18A2, SNRNP70, SYPL1, TADA2A, MAP3K7, TAL1, TEC, TPSAB1, RNF103, STAM, NDST2, AGPS, PABPC4, RGS11, CD84, CDC16, USP10, WDR46, ZNF264, CTR9, DEPDC5, SNUPN, KLRG1, BAIAP2, HSPH1, SERINC3, IFT27, ATXN2L, U2AF2, ZBTB1, FAM120A, DNAJC13, ESYT1, ARHGEF12, ATXN10, UPF2, STAP1, SIGLEC7, SNX5, SIGLEC8, HPGDS, REM1, CPSF1, BET1L, NBAS, LAX1, BTBD7, CENPJ, CTDSP1, HRH4, ACBD3, DCLRE1B, MBOAT7, CHD9, ZKSCAN3, C6orf25, USP48, OSBPL9, ZNF549, TIPRL |
| Mast_cells_FANTOM_3___282 | Aran_et_al_2017 | AMHR2, ANXA1, ANXA11, BMPR1A, BTK, C3AR1, C8G, CASP10, CD22, SIGLEC6, CPA3, CTSG, DIAPH1, MS4A2, GATA1, HDC, IL5, ITGA2B, KRT1, LTC4S, LYL1, NTRK1, OSBP, P2RX1, PAK2, PIK3R2, PRG2, PTGER3, RENBP, RGS13, RXRB, SLC18A2, SNRNP70, TADA2A, TAL1, TPSAB1, NDST2, AGPS, PABPC4, RGS11, USP10, WDR46, CTR9, DEPDC5, KLRG1, HSPH1, IFT27, ATXN2L, U2AF2, FAM120A, UPF2, STAP1, SNX5, SIGLEC8, HPGDS, REM1, CPSF1, BET1L, NBAS, LAX1, HRH4, ACBD3, DCLRE1B, MBOAT7, ZKSCAN3, C6orf25, OSBPL9, TIPRL |
| Megakaryocytes_BLUEPRINT_1___283 | Aran_et_al_2017 | ARHGAP6, GP1BA, HTR2A, MPL, PF4V1, SELP, GP6, CLEC1B, RUFY1 |
| Megakaryocytes_BLUEPRINT_2___284 | Aran_et_al_2017 | ANXA3, ARHGAP6, GP1BA, MPL, PF4V1, SELP, NCKAP1, CLEC1B, TUBB1 |
| Megakaryocytes_BLUEPRINT_3___285 | Aran_et_al_2017 | ANXA3, ARHGAP6, GP1BA, MPL, SELP, CLEC1B, LRP2BP, TUBB1 |
| Megakaryocytes_NOVERSHTERN_1___286 | Aran_et_al_2017 | GP1BA, HBD, PF4V1, SELP, VWF, KALRN, RGS6, GP6, CLEC1B |
| Megakaryocytes_NOVERSHTERN_2___287 | Aran_et_al_2017 | GP1BA, HBD, PF4V1, SELP, VWF, KALRN, RGS6, GP6, CLEC1B, TUBB1 |

| | | |
|---|---|---|
| Megakaryocytes_NOVERSHTERN_3___288 | Aran_et_al_2017 | GP1BA, PF4V1, SELP, VWF, KALRN, RGS6, GP6, CLEC1B |
| Melanocytes_ENCODE_1___289 | Aran_et_al_2017 | CA8, MLANA, GJA3, OCA2, S100B, SOX10, GPR137B, TYR, KCNAB2, PIR, FARP2, KCNE4, QPCT, SLC45A2, TBC1D16 |
| Melanocytes_ENCODE_2___290 | Aran_et_al_2017 | CA8, CDH3, MLANA, GJA3, MMP17, OCA2, CLEC11A, SOX10, GPR137B, TYR, TYRP1, KCNAB2, PIR, TNFRSF14, FARP2, QPCT, SLC45A2, WIPI1, MCOLN1, WFDC1, APH1B, UAP1L1, TBC1D16 |
| Melanocytes_ENCODE_3___291 | Aran_et_al_2017 | CA8, CBR3, NOV, OCA2, SOX10, TYR, PIR, FARP2, QPCT, SLC45A2, UAP1L1, TBC1D16 |
| Melanocytes_FANTOM_1___292 | Aran_et_al_2017 | DCT, MLANA, TRPM1, OCA2, SOX10, TYR, TYRP1, SLC16A6, PLXNC1, SLC45A2 |
| Melanocytes_FANTOM_2___293 | Aran_et_al_2017 | ACACB, ACP5, ASPA, BCL2A1, CEACAM1, CAPN3, RUNX3, CDK2, LYST, CLCN5, ABCC2, CYP27A1, DAB2, DCT, EDNRB, ERBB3, ESR2, MLANA, IFI6, GCNT2, GJB1, GK, GMPR, GNAL, HLA-DRA, HLA-DRB1, HLA-F, IFI16, IFI27, IFI35, SP110, IFIT2, IFIT1, IFIT3, IRF4, ISG20, ITGB8, ITPKB, KCNJ13, KIT, L1CAM, MBP, MITF, TRPM1, MMP8, MX1, MX2, MYO5A, NOV, GPR143, OAS1, OAS2, OCA2, PAEP, PAX3, PDE3A, PDE3B, PDE4B, PLP1, PLSCR1, PROS1, RAB27A, SGK1, SLC1A4, SOX10, STAT1, TAP1, TCN1, TLR1, TYR, TYRP1, UBA7, WARS, ALX1, KCNAB2, GAS7, PIR, OASL, TNFRSF14, INPP4B, GYG2, KYNU, SLC16A6, ITM2A, SOX13, AATK, ISG15, GREB1, ZFYVE16, RNF144A, ZEB2, FARP2, XYLB, PLXNC1, HMG20B, TESK2, GPNMB, IFI44, IQGAP2, IFI44L, LZTS1, NLGN1, ATP10B, ANKRD28, MCF2L, NEDD4L, TDRD7, DAAM2, DDX58, CA14, SLC7A11, PRKD3, IFIT5, QPCT, SLC39A6, GAPDHS, FOXD3, CDH19, OSTM1, SLC43A3, ATP6V0A4, CDON, PI15, LAP3, SLC45A2, LEF1, HERC5, PLA1A, TRPV2, C21orf91, PLEKHA5, XAF1, BNC2, SAMD9, HERC6, MOCOS, WIPI1, TMEM140, MCOLN3, DDX60, DOCK10, BIN3, CYSLTR2, SEMA6A, AVPI1, RTP4, IFIH1, RAB17, GNPTAB, TMPRSS5, ARHGAP24, RSAD2, TBC1D16, ASB9, PRUNE2, PHACTR1, ZNF749 |
| Melanocytes_FANTOM_3___294 | Aran_et_al_2017 | ABL2, ACACB, ACP5, APOD, ASPA, BCL2A1, CEACAM1, CAPN3, RUNX3, CBR3, CD36, CDK2, LYST, CLCN5, CLCN7, ABCC2, CTSK, CYP27A1, DAB2, DCT, NQO1, EDNRB, ERBB3, ESR2, ETV5, ACSL3, MLANA, IFI6, GCNT2, GJB1, GK, GMPR, GNAL, HLA-DPA1, HLA-DRA, HLA-DRB1, HLA-F, IFI16, IFI27, IFI35, SP110, IFIT2, IFIT1, IFIT3, IRF4, IRF7, ISG20, ITGB8, ITPKB, KCNJ13, KIT, L1CAM, LGALS3, MBP, MITF, TRPM1, MME, MMP8, MX1, MX2, MYO5A, NOV, GPR143, OAS1, OAS2, OCA2, PAEP, PAX3, PDE3A, PDE3B, PDE4B, PDE4D, PDK4, PLP1, PLSCR1, PRKCE, PROS1, RAB27A, SGK1, SLC1A4, SLC12A2, SOX10, SP100, STAT1, STAT5A, STX3, TAP1, TBX2, TCN1, TLR1, TYR, TYRP1, UBA7, VGF, WARS, ALX1, SORBS2, PPFIBP2, KCNAB2, IFITM1, GAS7, PIR, OASL, TNFRSF14, INPP4B, GYG2, KYNU, SLC16A6, UBE2L6, ITM2A, SOX13, AATK, ISG15, GREB1, ZFYVE16, RNF144A, TRIM14, ZEB2, FARP2, XYLB, NAMPT, PLXNC1, HMG20B, TESK2, GPNMB, IFI44, IQGAP2, IFI44L, SLC27A3, CIT, LZTS1, IRAK3, NLGN1, PDZRN3, ATP10B, ANKRD28, MCF2L, NEDD4L, TDRD7, DAAM2, ZFYVE26, DDX58, CA14, SLC7A11, PRKD3, IFIT5, QPCT, SLC39A6, SAMHD1, GAPDHS, FOXD3, CDH19, TRIB2, OSTM1, SLC43A3, ATP6V0A4, CDON, PI15, LAP3, SLC45A2, LEF1, HERC5, PLA1A, TRPV2, C21orf91, PLEKHA5, SPATA6, EGLN1, XAF1, BNC2, SAMD9, RPP25, HERC6, MOCOS, WIPI1, TMEM140, MCOLN3, DDX60, DOCK10, BIN3, CYSLTR2, SEMA6A, AVPI1, BCAN, RTP4, IFIH1, POPDC3, RAB17, GNPTAB, APOL6, TMPRSS5, APOLD1, ARHGAP24, RSAD2, TBC1D16, ASB9, PYHIN1, PRUNE2, PHACTR1, ZNF749 |
| Memory_Bcells_BLUEPRINT_1___295 | Aran_et_al_2017 | ACRV1, ADCY2, AQP8, ART1, TNFRSF17, BLK, C4BPA, S100G, CASQ2, CD1C, CD19, MS4A1, CD22, SIGLEC6, CD37, CD72, CD79A, CD79B, CETP, CHRM2, CHRNA2, CCR6, CNR2, COX6A2, CPA2, CPB1, CSN1S1, NCAN, CYLC2, CYP2A7, CYP2C19, DCC, DPP6, DSP, FMO1, FSHR, GABRA4, GAD2, GK2, GNAT2, GNRHR, GPX5, GRIN2B, GRM6, HCRTR2, HLA-DPB1, HSD3B2, HTN3, INHBC, KCNA5, KCNJ10, KIF5A, KRT2, LECT2, CD180, MBL2, MC4R, MEFV, MEP1B, MAP3K9, NPY5R, NTRK3, OTC, PAX5, SERPINA4, PLIN1, PNOC, PNLIPRP1, POU4F2, PRKCB, PTH1R, PRPH2, SELP, SLC12A3, SLC17A1, SLN, SPIB, SYN2, SYPL1, TSHB, WNT2, ZIC3, NPHS2, UNC5C, ADAM20, PROZ, BAIAP3, KALRN, KRT75, CER1, TMPRSS11D, LY86, KIAA0125, AP1M2, SSX3, GLYAT, DSCR4, COLEC10, RRH, CCR9, CD3EAP, ADAM30, SP140, GGA2, HECW1, CRB1, TNFRSF13B, CHST5, AIPL1, SLC24A2, STAP1, CLDN17, TSPAN13, VPREB3, FSCN3, TAS2R14, WNT16, SLCO1C1, DDX4, ULK4, QRSL1, GPRC5D, SLC30A10, SLC17A7, PGLYRP4, VN1R1, HRH4, SLC5A7, CHP2, MS4A5, ZNF747, FCRL2, RIC3, TCTN2, TRPM3, MOGAT2, ADAMTS12, OBSCN, MYOZ3, ZNF548, KHDRBS2, SHISA6, FMO6P, ANKRD34C |
| Memory_Bcells_BLUEPRINT_2___296 | Aran_et_al_2017 | TNFRSF17, BLK, CAPN3, CD19, MS4A1, CD22, CD37, CD72, CD79A, CD79B, CCR6, CNR2, CPA2, CPB1, CSN1S1, DPP6, DSP, FMO1, GK2, GRM6, HNRNPL, INHBC, |

| | | |
|---|---|---|
| | | KCNA5, CD180, MC4R, PAX5, PNOC, POU4F2, PRKCB, SLC17A1, SPIB, SYPL1, TSHB, ZIC3, ADAM21, LY86, SCRN1, KIAA0125, RRH, CCR9, CD3EAP, SP140, GGA2, TNFRSF13B, AIPL1, STAP1, VPREB3, WNT16, SLCO1C1, DDX4, ULK4, QRSL1, SLC30A10, PGLYRP4, HRH4, SLC5A7, MS4A5, FCRL2, TCTN2, TRPM3, MOGAT2, OBSCN, ZNF548, PIKFYVE, SHISA6, FMO6P, ANKRD34C |
| Memory_Bcells_BLUEPRINT_3___297 | Aran_et_al_2017 | ART1, BLK, S100G, CASQ2, CD1C, CD19, MS4A1, CD22, SIGLEC6, CD37, CD72, CD79A, CD79B, CHRM2, CCR6, CNR2, CPB1, CSN1S1, CYP2A7, FMO1, GABRA4, GAD2, GK2, GNAT2, GRM6, INHBC, KCNJ10, KRT2, CD180, MEFV, MAP3K9, NPY5R, PAX5, PNOC, PNLIPRP1, POU4F2, PRKCB, SELP, SPIB, TSHB, KALRN, CER1, LY86, KIAA0125, SSX3, RRH, CD3EAP, SP140, GGA2, HECW1, TNFRSF13B, CHST5, AIPL1, SLC24A2, STAP1, TSPAN13, VPREB3, TAS2R14, SLCO1C1, DDX4, QRSL1, PGLYRP4, VN1R1, SLC5A7, ZNF747, FCRL2, TRPM3, MOGAT2, ZNF548, SHISA6 |
| Memory_Bcells_HPCA_1___298 | Aran_et_al_2017 | BLK, CD19, MS4A1, CD22, CD79A, CD79B, GRM6, INHBC, PNOC, SPIB, TSHB, KIAA0125, CD3EAP, TNFRSF13B, TSPAN13, VPREB3, QRSL1, FCRL2, MOGAT2 |
| Memory_Bcells_HPCA_2___299 | Aran_et_al_2017 | TNFRSF17, BLK, CD19, MS4A1, CD22, CD37, CD72, CD79A, CD79B, CSN1S1, DSP, FMO1, GK2, GRM6, INHBC, CD180, PNOC, POU4F2, PRKCB, SPIB, TSHB, LY86, KIAA0125, RRH, CCR9, CD3EAP, SP140, GGA2, TNFRSF13B, STAP1, VPREB3, QRSL1, HRH4, SLC5A7, FCRL2, ZNF548, SHISA6, FMO6P, ANKRD34C |
| Memory_Bcells_HPCA_3___300 | Aran_et_al_2017 | BLK, CD19, MS4A1, CD22, CD79A, CD79B, INHBC, PNOC, SPIB, KIAA0125, TSPAN13, VPREB3, QRSL1, FCRL2 |
| Memory_Bcells_IRIS_1___301 | Aran_et_al_2017 | BLK, CD1C, CD79B, ODC1, SPIB, MBD4, LY86, SP140, TNFRSF13B, FSCN2, ZBTB32, NT5C, WNT16, TRMT61A |
| Memory_Bcells_IRIS_2___302 | Aran_et_al_2017 | BLK, CXCR5, CD19, MS4A1, CD22, CCR6, HTR3A, MGAT5, SPIB, RNGTT, MBD4, LY86, CXCL13, SP140, TNFRSF13B, ZBTB32, MIOS, AICDA, FCRL2 |
| Memory_Bcells_IRIS_3___303 | Aran_et_al_2017 | BLK, CD1C, CD79B, ODC1, SPIB, MBD4, SP140, TNFRSF13B, FSCN2, ZBTB32, NT5C, WNT16 |
| MEP_BLUEPRINT_1___304 | Aran_et_al_2017 | AHCY, CDK4, CPA3, CRHBP, ERG, GSTM5, HDC, EPCAM, PRG2, RYR3, MINPP1, PAICS, HPGDS, LAPTM4B |
| MEP_BLUEPRINT_2___305 | Aran_et_al_2017 | AHCY, CDK4, CPA3, CRHBP, GSTM5, HDC, EPCAM, POLE2, HPGDS, LAPTM4B, TCEAL4 |
| MEP_BLUEPRINT_3___306 | Aran_et_al_2017 | BUB1B, CPA3, CRHBP, ERG, HDC, POLE2, RYR3, STIL, KIAA0101, SNX5, HPGDS, MRPL15, C11orf95, FAM124B |
| MEP_HPCA_1___307 | Aran_et_al_2017 | FXN, GATA1, HBD, KEL, MYL4, PCCB, PNMT, PRG2, RHAG, SURF2, TPSAB1, UNG, AIMP2, CHAF1B, RUVBL1, NAT6, ERAL1, HPGDS, MRPS2, CTNNBL1, EXOSC5, ACD, DCTPP1, MTG1 |
| MEP_HPCA_2___308 | Aran_et_al_2017 | FCER1A, FMO1, FXN, KRT1, NTRK1, PCCB, PNMT, UNG, IFRD2, AIMP2, CHAF1B, RUVBL1, MRPL28, TMED1, FBXO7, ERAL1, HPGDS, MECR, MRPS2, DDX41, CTNNBL1, C21orf59, EXOSC5, ACD, DCTPP1, NOL12, MTG1 |
| MEP_HPCA_3___309 | Aran_et_al_2017 | FXN, PCCB, PNMT, TMED1, HPGDS, MRPS2, CTNNBL1, ACD, NOL12 |
| MEP_NOVERSHTERN_1___310 | Aran_et_al_2017 | ATIC, BCS1L, CCT6A, DDX1, DDX10, FXN, HNRNPAB, ITGA2B, PCCB, POLE2, RYR3, IFRD2, RUVBL1, PSMG1, LDB1, VAPA, TXNL1, KEAP1, URB1, LRPPRC, TRAP1, PRMT3, NUDC, FASTKD2, PDCD11, MYO16, PPRC1, WDR43, MLC1, SERBP1, ERAL1, GNL3, NUFIP1, RPUSD2, RRP15, MRPS2, MRTO4, NOP16, MKS1, PAK1IP1, CDKN2AIP, TSR1, WDR12, CTNNBL1, BCCIP, GUF1, CARS2, FN3KRP, OGFOD2, MRM1, NOL10, NAA15, ZKSCAN3, MED25, TIMM50 |
| MEP_NOVERSHTERN_2___311 | Aran_et_al_2017 | CA1, FMO1, FXN, HDC, NTRK1, PNMT, PRG2, HPGDS, MRPS2, COQ3, CTNNBL1, ACD |
| MEP_NOVERSHTERN_3___312 | Aran_et_al_2017 | FCER1A, FMO1, FXN, KRT1, NTRK1, PCCB, PNMT, UNG, IFRD2, AIMP2, CHAF1B, RUVBL1, MRPL28, TMED1, FBXO7, ERAL1, HPGDS, MECR, MRPS2, DDX41, CTNNBL1, C21orf59, EXOSC5, ACD, DCTPP1, NOL12, MTG1 |
| Mesangial_cells_ENCODE_1___313 | Aran_et_al_2017 | CDH6, CDKN1C, EDN2, FOXF1, FOXD1, HOXA11, HOXD1, LHX1, MICB, TLL2, DOC2B, PADI2, RHOF, KIRREL, CLSTN2, CRISPLD2 |
| Mesangial_cells_ENCODE_2___314 | Aran_et_al_2017 | CDKN1C, EDN2, FOXD1, HOXA11, HOXD1, LHX1, MICB, TLL2, DOC2B, ARL4C, KIRREL, CLSTN2 |
| Mesangial_cells_ENCODE_3___315 | Aran_et_al_2017 | CDH6, CDH16, CDKN1C, COL4A1, ARID3A, EDN2, EPHB2, FOXD1, GPI, HOXA3, HOXA11, HOXB2, HOXB3, HOXC10, HOXD1, HOXD11, ILK, INPP5A, ITGA3, ITGB3, LAMA5, LHX1, MICB, NME3, NTHL1, PAX2, PDCD2, PFDN1, PLEC, PVR, HNF1B, NR2F2, TLL2, UBA1, UCP2, UROD, WNT7B, PXDN, DOC2B, RNASET2, FGF18, CLDN6, |

| | | |
|---|---|---|
| | | DHRS3, HS3ST3A1, REC8, ARL4C, PROCR, PNPLA6, RBPMS, PLA2G16, MGAT4B, CSDC2, CPA4, ISYNA1, SMOX, RHOF, KIRREL, LAPTM4B, RBM38, CORO1B, CLSTN2, PCYOX1L, GALNT14, PDZD7, BICC1, ORAI2, THAP7, CRISPLD2, APOBEC3F, ASPHD1, HSPA12A |
| Mesangial_cells_FANTOM_1___316 | Aran_et_al_2017 | BGN, BST2, CD70, CNN1, COL1A1, COL3A1, COL4A1, COL5A1, COL6A3, CLDN4, VCAN, CTGF, DCN, DPYSL3, ELF3, SLC29A1, EPHB2, F2RL2, FGB, FOXC1, FOXD1, FLNC, FN1, GATA6, GLI2, SFN, HOXA5, HOXA7, HOXA10, HOXA11, HOXB2, HOXB3, HOXB6, HOXB9, HOXD11, IGFBP2, IGFBP4, IGFBP5, ITGB3, KCNJ15, LAD1, LAMA5, LHX1, LIF, SMAD6, MITF, MMP7, MYLK, PDGFB, PDGFRB, ABCB1, PLAT, PODXL, PPARG, PTGER2, PTX3, RAB3B, RGS4, TAGLN, HNF1B, TFPI, TGFB2, TGM2, UCP2, VCAM1, WNT7B, PAX8, HMGA2, DYSF, DOC2B, CLDN6, CLDN1, DIRAS3, IL32, SPOCK2, HS3ST3A1, REC8, LRRC17, DLC1, MYL9, POSTN, PPP1R13L, MMP24, NID2, RAP1GAP2, NUP210, GREM1, ANKRD1, LMCD1, LSR, RHOF, MXRA8, RCN3, ALPK3, HKDC1, NUAK2, CRISPLD2, CREB3L1, LYPD1, OLFML2A |
| Mesangial_cells_FANTOM_2___317 | Aran_et_al_2017 | CD70, F2RL2, FGB, FOXC2, HOXA3, HOXA7, HOXA10, HOXA11, HOXB8, HOXB9, HOXD10, HOXD11, PAX2, ABCB1, PODXL, RAB3B, SALL1, HNF1B, UCP2, PAX8, DOC2B, CLDN6, REC8, B3GALT5, IFITM2, NUP210, RHOF, C14orf105, CDHR1 |
| Mesangial_cells_FANTOM_3___318 | Aran_et_al_2017 | ACTA2, ADORA1, AEBP1, ARHGAP4, ATP7B, BGN, BMP4, CD70, CDH6, CNN1, COL1A1, COL1A2, COL3A1, COL5A1, COL6A1, COL6A2, COL6A3, CLDN4, CLDN3, CRIP1, CYBA, DCN, DNM1, DPYSL3, EDN2, ELF3, ENPEP, SLC29A1, F2RL2, FGB, FOXC1, FOXD1, FOXC2, FN1, FZD2, GATA6, GLI2, CCR10, GRB14, HOXA3, HOXA4, HOXA6, HOXA7, HOXA10, HOXA11, HOXB6, HOXB8, HOXB9, HOXC10, HOXD9, HOXD10, HOXD11, HPGD, HSPA2, IGFBP4, ITGB3, KRT19, LFNG, LHX1, SMAD6, MEIS2, MGP, MITF, MMP7, NCAM1, NOV, NPPB, OXTR, PAPPA, PAX2, PCOLCE, PDGFB, PDGFRB, ABCB1, PLAT, PODXL, PPARG, PTGER2, PTPRJ, RAB3B, RGS4, RRAD, SALL1, SDC2, SIX1, SLC22A3, SPP1, SST, TAGLN, TBL1X, TBXAS1, HNF1B, NR2F1, TGFB1I1, TGFB2, TRPC4, UCP2, WT1, PAX8, DYSF, DOC2B, ADAM19, TNFSF10, PROM1, CACNA1H, CLIC3, CLDN6, CLDN1, DIRAS3, IL32, PDLIM7, RPH3AL, ADAMTS3, GAL3ST1, SPOCK2, KBTBD11, HS3ST3A1, REC8, ARL4C, MSLN, B3GALT5, DLC1, MYL9, IFITM2, POSTN, SIX2, PLK2, MMP24, RBPMS, PLA2G16, NID2, DENND3, PLXND1, SULF1, NUP210, QPRT, OLFML2B, GREM1, CPNE7, HIPK2, TRHDE, EFEMP2, ZNF580, CPA4, SHC3, FXYD6, RHOF, HCFC1R1, C14orf105, LRRC20, KIRREL, GPRC5C, OLFML3, SDR39U1, RCN3, ALPK3, SYT13, GALNT14, DNAJC22, BICC1, HKDC1, RAB11FIP1, COL18A1, LBH, CRISPLD2, CREB3L1, CDHR1, ATP6V0E2, CADM4 |
| Monocytes_BLUEPRINT_1___319 | Aran_et_al_2017 | ASGR2, FCN1, CFP, RNASE2, CD300C, LILRA1, TLR7, MS4A6A, LILRA5, CCR2 |
| Monocytes_BLUEPRINT_2___320 | Aran_et_al_2017 | AIF1, ASGR2, CYBB, FCAR, FCN1, KCNMB1, MEFV, MNDA, CFP, S100A12, CD163, CD101, CLEC5A, FBXL5, TREM1, RETN, CCR2 |
| Monocytes_BLUEPRINT_3___321 | Aran_et_al_2017 | ASGR2, CSF3R, F13A1, FCN1, CFP, RNASE2, S100A12, LILRB2, CD93, PADI4, P2RY13, MS4A6A |
| Monocytes_FANTOM_1___322 | Aran_et_al_2017 | AIF1, APAF1, RHOA, RHOG, ARNT, ASGR2, C3AR1, CASP5, TNFRSF8, CD33, CSF1R, CSF3R, CYBB, FCAR, FCN1, FPR2, HCK, HK3, HRH2, HSPA6, KCNMB1, MEFV, MAP3K3, MNDA, MYO1F, CFP, PHKG2, PTGIR, RARA, S100A12, TYROBP, UBE2D1, UPK3A, VASP, BEST1, LST1, NUP214, IQGAP1, ATP6V0D1, CD163, CD101, EIF4E2, CALCOCO2, LILRB2, TGOLN2, CAMKK2, LILRB1, CD300C, FGL2, LILRA1, LILRB3, LILRA2, TREX1, GABARAP, RPH3A, ACAP2, CLEC5A, IL17RA, FBXL5, CLEC4E, VENTX, COMMD9, PILRA, METTL9, TLR7, TLR8, P2RY13, TREM1, RIN2, RHOT1, TMEM127, TMEM9B, RETN, RPGRIP1, DENND1A, DPEP2, MS4A6A, OSBPL11, LILRA5, CCR2 |
| Monocytes_FANTOM_2___323 | Aran_et_al_2017 | ABCB7, AIF1, ANXA1, ARF5, ASGR2, BPI, BTK, TSPO, C3AR1, CAPN2, CAPN3, CAPNS1, CAST, CD4, CD33, CEACAM4, MAPK14, CSF1R, CSF3R, CTBP2, CX3CR1, CYBB, DHX8, F13A1, FCAR, FCER1G, FCN1, FGR, FOLR2, FPR2, HADHA, HCK, HK3, AGFG1, HSPA6, CXCR2, IL10RA, IMPDH1, KCNMB1, LTBR, LYL1, MAN2C1, MARK3, MAP3K3, MAP3K11, MNDA, MNT, MYO1F, NUBP1, NCF4, CFP, PLP2, PPP1CB, PRKACA, PTEN, PTGIR, RARA, RNASE2, S100A6, S100A10, S100A12, SH3BP2, SLC11A1, NEK4, CLEC3B, TPD52L2, UBE2D1, USP4, BEST1, WAS, LST1, NUP214, NCOA4, NDST2, PIAS1, MARCO, TMEM11, MTMR3, BTAF1, DOK2, PSTPIP1, CD163, CD101, QKI, LY86, EIF4E2, H2AFY, PPM1F, SNX17, GIT2, USP15, COL4A3BP, CALCOCO2, RGS19, LILRB2, RTN3, CEPT1, CLEC10A, CAMKK2, LILRB1, CD300C, LILRA1, LILRB3, LILRA2, OGFR, WWP2, AKAP13, TREX1, GABARAP, RPH3A, CD93, SPEN, SETX, KDM6B, STAB1, KLHL18, ANKS1A, UBR2, SIN3B, DNAJC13, KIAA1033, SUN1, SIK3, MED13L, SEC11A, ACAP2, PADI4, STX12, IL17RA, PGLS, FAM32A, FBXL5, CLEC4E, PTPN18, SIGLEC9, COQ2, VENTX, CECR5, NKIRAS2, COMMD9, STRN4, PILRA, CLEC4A, METTL9, TLR7, ZDHHC3, TLR8, TMBIM4, P2RY13, TREM1, RIN2, RBM41, RHOT1, NPLOC4, WDR11, NSFL1C, RETN, DENND1A, PCTP, DPEP2, |

| | | |
|---|---|---|
| | | MS4A6A, MTMR14, ATG3, AHNAK, TSEN34, MBOAT7, CARS2, DOK3, PANK2, FBXO11, SPG11, CXorf21, DHX57, SLC38A10, UBXN2B, HIPK1, YTHDF3, LILRA5, CCR2 |
| Monocytes_FANTOM_3___324 | Aran_et_al_2017 | ABCB7, AIF1, ANXA1, ARF5, ASGR2, BPI, BTK, TSPO, C3AR1, CAPN2, CAPN3, CAPNS1, CAST, CD4, CD33, CEACAM4, MAPK14, CSF1R, CSF3R, CTBP2, CX3CR1, CYBB, DHX8, F13A1, FCAR, FCER1G, FCN1, FGR, FPR2, HCK, HK3, AGFG1, HSPA6, IL10RA, IMPDH1, KCNMB1, LTBR, LYL1, MAN2C1, MARK3, MAP3K3, MAP3K11, MNDA, MNT, MYO1F, CFP, PLP2, PPP1CB, PRKACA, PTEN, RARA, RNASE2, S100A6, S100A10, S100A12, SH3BP2, CLEC3B, TPD52L2, UBE2D1, USP4, BEST1, WAS, LST1, NUP214, NCOA4, CUL5, NDST2, PIAS1, MARCO, TMEM11, MTMR3, BTAF1, DOK2, PSTPIP1, ZMYM4, CD163, CD101, LY86, EIF4E2, H2AFY, PPM1F, SNX17, GIT2, USP15, CALCOCO2, RGS19, LILRB2, RTN3, CLEC10A, CAMKK2, LILRB1, CD300C, LILRA1, LILRB3, LILRA2, OGFR, WWP2, AKAP13, TREX1, GABARAP, RPH3A, CD93, SPEN, KDM6B, STAB1, KLHL18, ANKS1A, SIN3B, DNAJC13, KIAA1033, SUN1, SIK3, SEC11A, ACAP2, PADI4, STX12, PGLS, FAM32A, FBXL5, PTPN18, SIGLEC9, COQ2, VENTX, CECR5, NKIRAS2, COMMD9, STRN4, PILRA, CLEC4A, METTL9, TLR7, TLR8, P2RY13, TREM1, RIN2, RBM41, RHOT1, NPLOC4, WDR11, NSFL1C, RETN, DENND1A, PCTP, DPEP2, MS4A6A, MTMR14, ATG3, AHNAK, TSEN34, MBOAT7, CARS2, DOK3, PANK2, FBXO11, SPG11, USP48, DHX57, SLC38A10, PIKFYVE, HIPK1, JMJD1C, LILRA5, CCR2 |
| Monocytes_HPCA_1___325 | Aran_et_al_2017 | CSNK1A1, FCN1, FGR, HUS1, NUBP1, PNP, CFP, SH3BP2, SRC, LRRFIP1, LILRB2, EIF1B, FNDC3A, FKBP15, CLEC5A, RPS6KC1, CD244, ARL8B, RETN |
| Monocytes_HPCA_2___326 | Aran_et_al_2017 | CASP5, CD48, CSNK1A1, FCN1, FGR, FPR2, MEFV, PNP, CFP, LILRA1, FKBP15, CLEC5A, SIGLEC9, ARL8B, RETN |
| Monocytes_HPCA_3___327 | Aran_et_al_2017 | AP1G1, ASGR2, BNIP2, TSPO, CASP5, CD1E, TNFRSF8, CD33, CDK9, CSF1R, DDX3X, DHX8, DLG4, GPR183, ETF1, ETV3, EWSR1, FCAR, FCN1, FOLR2, FOLR3, FPR2, GALNT3, HIF1A, HNRNPU, AGFG1, KCNC3, KCNMB1, MEFV, MMP17, MTF1, MTHFR, OSM, PDE6H, CFP, PGGT1B, PLD2, PLEK, POU2F2, PPM1A, PRKACA, MAPK6, MAP2K1, PTGIR, RAB5A, RELA, CLIP1, S100A10, S100A12, SH3BP2, SLC11A1, STX5, SUPT6H, PHLDA2, UBE2D1, UPK3A, PTP4A2, ELL, NDST2, DNAH17, PTCH2, RIOK3, KSR1, BCL10, SOCS3, USP8, DDX21, PLAA, CD101, QKI, WTAP, AATK, TTLL4, BCL2L11, MPHOSPH6, AKAP8, LILRB2, APBB3, CLEC10A, GNA13, MAP3K2, LILRB1, CD300C, FGL2, LILRA1, OGFR, AKAP13, TREX1, CD93, IQSEC2, STAB1, JMJD6, CBX6, CLEC5A, PGLS, GGA1, FBXL5, CLEC4E, SERP1, GNMT, GPR162, VENTX, RABGEF1, TBK1, STRN4, CLEC4A, ADIPOR1, METTL9, RLIM, DCTN4, CLEC1A, ZBTB7A, CDC40, AZIN1, P2RY13, TREM1, MIOS, TMEM104, VNN3, SAR1A, CYSLTR2, DENND1A, SAMSN1, MS4A6A, MTMR14, ZNF668, GRPEL1, REEP4, ZNF787, ZFC3H1, YTHDF3, LILRA5, ZNF710, TMEM110 |
| Monocytes_IRIS_1___328 | Aran_et_al_2017 | ASGR2, TNFRSF8, FCAR, FCN1, FOLR2, HIC1, MEFV, CFP, S100A12, SLC11A1, FBXL5, VENTX, RABGEF1, TREM1, VNN3, SAMSN1, MS4A6A, LILRA5 |
| Monocytes_IRIS_2___329 | Aran_et_al_2017 | ASGR2, FCAR, FCN1, S100A12, UPK3A, VENTX, TREM1, VNN3, MS4A6A, LILRA5 |
| Monocytes_IRIS_3___330 | Aran_et_al_2017 | ASGR2, CASP5, TNFRSF8, FCAR, FCN1, FOLR2, HIC1, HIF1A, MEFV, MMP17, OSM, CFP, PTGIR, S100A12, SLC11A1, UPK3A, RIOK3, SOCS3, SEMA6B, GNA13, FGL2, STAB1, PADI4, CLEC5A, FBXL5, VENTX, RABGEF1, METTL9, P2RY13, TREM1, VNN3, DENND1A, SAMSN1, MS4A6A, LILRA5 |
| Monocytes_NOVERSHTERN_1___331 | Aran_et_al_2017 | AIF1, ANXA1, ASGR2, BPI, BTK, TSPO, C3AR1, CAPN3, CAST, CD4, CD33, CEACAM4, MAPK14, CSF3R, CTBP2, CX3CR1, CYBB, F13A1, FCAR, FCER1A, FCN1, FGR, FPR2, HCK, HK3, HSPA6, CXCR2, KCNMB1, LTBR, LYL1, MNDA, MYO1F, NCF4, CFP, PLP2, PTEN, RNASE2, S100A12, NEK4, CLEC3B, UBE2D1, LST1, NUP214, MARCO, DOK2, PSTPIP1, CD163, CD101, LY86, PPM1F, GIT2, USP15, RGS19, LILRB2, RTN3, CLEC10A, LILRB1, CD300C, LILRA1, LILRB3, LILRA2, CD93, STAB1, ANKS1A, ACAP2, PADI4, PGLS, FBXL5, PTPN18, COQ2, COMMD9, PILRA, CLEC4A, METTL9, TLR7, TLR8, MS4A4A, P2RY13, TREM1, RHOT1, RETN, PCTP, DPEP2, MS4A6A, ATG3, AHNAK, TSEN34, CARS2, UBXN2B, LILRA5, CCR2 |
| Monocytes_NOVERSHTERN_2___332 | Aran_et_al_2017 | ASGR2, FCAR, FCN1, FOLR2, MEFV, CFP, S100A12, SLC11A1, UPK3A, FBXL5, VENTX, TREM1, VNN3, MS4A6A |
| Monocytes_NOVERSHTERN_3___333 | Aran_et_al_2017 | ASGR2, FCAR, FCN1, FOLR2, MEFV, CFP, S100A12, SLC11A1, UPK3A, VENTX, TREM1, VNN3, MS4A6A |
| MPP_BLUEPRINT_1___334 | Aran_et_al_2017 | ADCY3, CRHBP, ERG, CAPRIN1, P2RX1, SERPINI2, TEC, TIE1, UMPS, MLF2, ZNF282, NSMAF, USP6, MED7, H2AFY, PPM1F, PARP2, MRPS31, CALCOCO2, TMEM147, SF3A3, CSTF2T, LSM5, ATXN10, HIBCH, ATP2C1, ZNF219, TAOK3, PTRH2, TBC1D13, TMEM70, ATP5SL, WDR60, DPPA4, AGK, MKL2, BAHCC1, ALG8, CENPO, SPAG16, UBA5, SFXN3, FAM136A, G6PC3, HNRNPA3, DPY19L4 |

| | | |
|---|---|---|
| MPP_BLUEPRINT_2___335 | Aran_et_al_2017 | ABO, AIF1, AMD1, ATP5J, AVP, BTF3, ERCC8, CLNS1A, CPA3, CRHBP, CRYGD, DNTT, DUT, FANCA, HINT1, IGLL1, EIF3E, ITGA9, LRCH4, MPL, MPO, NACA, NFYA, NPM1, NUP98, PIGF, PRTN3, PSMA2, PSMA6, RAG2, RBBP7, RFC2, RNASE2, RNASE3, RPS3, RPS24, SMARCC2, SNRPF, TEC, TERT, TRH, TSSC1, TTF1, ZNF32, ZNF35, ZNF134, CD164, GALR2, CDC123, MED7, PMPCB, TATDN2, KIAA0125, EIF1B, PPIH, ANP32B, ERLIN1, AKAP13, ESYT1, HAUS5, NAT6, ATP2C1, INVS, LSM1, BZW2, GLTSCR2, RSL24D1, ZNF639, REV1, TEX10, SRBD1, NHP2, PCID2, NSFL1C, TMEM9B, TFB2M, GGCT, MAP7D3, MTDH, IMP4 |
| MPP_BLUEPRINT_3___336 | Aran_et_al_2017 | ABO, AMD1, ATP5J, AVP, BTF3, ERCC8, CLNS1A, CRHBP, CRYGD, DUT, FANCA, HINT1, IGBP1, IGLL1, IMPDH2, EIF3E, ITGA9, LRCH4, MPL, MPO, NACA, NUP98, PIGF, PSMA2, PSMA6, RBBP7, RFC2, RNASE2, RPS3, RPS24, SNRPF, TERT, ZNF32, ZNF134, GALR2, MED7, TATDN2, KIAA0125, NUBP2, PPIH, AKAP13, ESYT1, HAUS5, INVS, LSM1, BZW2, RSL24D1, ZNF639, TEX10, SRBD1, NHP2, PCID2, NSFL1C, TMEM9B, TFB2M, ZDHHC6, GGCT, MAP7D3, CSPP1, FAM136A, MTDH, IMP4 |
| MPP_ENCODE_1___337 | Aran_et_al_2017 | AVP, AZU1, CRHBP, DNTT, FLT3, IGLL1, ITGA9, MPL, MPO, NUP98, RNASE2, SPN, CD164, KIAA0125, LSM1, VPREB3 |
| MPP_ENCODE_2___338 | Aran_et_al_2017 | AVP, CRHBP, FLT3, IGLL1, MPL, MPO, KIAA0125, VPREB3 |
| MPP_ENCODE_3___339 | Aran_et_al_2017 | AVP, CRHBP, FLT3, IGLL1, MPL, MPO, KIAA0125, VPREB3 |
| MPP_FANTOM_1___340 | Aran_et_al_2017 | ABO, AVP, CD37, CD52, CPA3, CRHBP, CSF2RB, CSF3R, CTSW, FCER1A, FLT3, GATA1, HBB, HBD, HDC, HLA-DOA, IGLL1, ITGA2B, ITGA9, ITGAL, JAK3, CIITA, MPL, MPO, NKG7, OSM, P2RX1, PIK3CG, PLCB2, PRKCB, PRTN3, PTPN6, PTPN7, PTPRC, PTPRCAP, SELPLG, SPN, GFI1B, S1PR4, BAIAP3, ACAP1, KIAA0125, TSPAN32, IKZF1, KLF1, MLC1, CD244, DPPA4, ARHGAP15, DPEP2, DOK3, TRAF3IP3, AGAP2 |
| MPP_FANTOM_2___341 | Aran_et_al_2017 | ABO, AVP, FMNL1, CD37, CD52, CPA3, CRHBP, CSF2RB, CSF3R, CTSW, FCER1A, FLT3, GATA1, HBB, HBD, HDC, HLA-DOA, IGLL1, IRF5, ITGA2B, ITGA9, ITGAL, JAK3, CIITA, MPL, MPO, NKG7, OSM, P2RX1, PIK3CG, PLCB2, PRKCB, PRTN3, PTPN6, PTPN7, PTPRC, PTPRCAP, SELPLG, SPN, LST1, GFI1B, S1PR4, BAIAP3, ACAP1, KIAA0125, TSPAN32, IKZF1, KLF1, CD300A, MLC1, CD244, SASH3, DPPA4, ARHGAP15, DPEP2, ATP8B4, DOK3, TRAF3IP3, AGAP2 |
| MPP_FANTOM_3___342 | Aran_et_al_2017 | AVP, CD37, CD53, CPA3, CRHBP, CSF3R, CTSW, FCER1A, HBD, NCKAP1L, HLA-DOA, IGLL1, ITGA2B, LAIR1, MPO, MYO1F, NCF4, OSM, P2RX1, PTPN7, PTPRC, PTPRCAP, SPI1, SPN, GFI1B, S1PR4, KIAA0125, TSPAN32, IKZF1, MLC1, SASH3, AGAP2 |
| MPP_NOVERSHTERN_1___343 | Aran_et_al_2017 | AVP, CRHBP, FLT3, IGLL1, MPL, MPO, NUP98, KIAA0125, ESYT1, VPREB3 |
| MPP_NOVERSHTERN_2___344 | Aran_et_al_2017 | AVP, CRHBP, FLT3, IGLL1, MPL, MPO, KIAA0125, VPREB3 |
| MPP_NOVERSHTERN_3___345 | Aran_et_al_2017 | AVP, CRHBP, FLT3, IGLL1, MPL, MPO, KIAA0125, VPREB3 |
| MSC_FANTOM_1___346 | Aran_et_al_2017 | HTR7, MMP17, PLA2G5, CDKL5, PKD2L1, ADAMTS12, ZNRF4, CTRB2 |
| MSC_FANTOM_2___347 | Aran_et_al_2017 | HTR7, MMP17, PRB3, CCL21, CDKL5, TNP2, PKD2L1, ZNF408, ADAMTS12, ZNRF4, CTRB2 |
| MSC_FANTOM_3___348 | Aran_et_al_2017 | COL10A1, DVL1, HAS1, HTR7, MMP17, NPAS1, PLA2G5, PRB3, CDKL5, TNP2, WISP1, PKD2L1, TRIM3, COPS8, ADAMTS12, TSSK1B, ZNRF4, CTRB2 |
| MSC_HPCA_1___349 | Aran_et_al_2017 | DCTD, DDOST, EEF1D, KIF22, LAMP1, NDUFA8, PARN, POLR2G, SLC35A2, HAND2, EIF4E2, CUL7, SCAMP3, STUB1, TMEM147, YIF1A, RER1, EMILIN1, SNF8, CABIN1, MKRN2, SPCS1, TRMT112, ZNF446, PRX, MRPS11, CUEDC2, SPAG16, MRPL24, TBC1D17, NHEJ1, PODNL1, SF3B5, THAP3, SLC38A10, SIX5 |
| MSC_HPCA_2___350 | Aran_et_al_2017 | DCTD, EEF1D, KIF22, POLR2G, HAND2, EIF4E2, CUL7, SCAMP3, YIF1A, RER1, EMILIN1, CABIN1, TRMT112, PRX, MRPL24, NHEJ1, PODNL1, SF3B5, THAP3 |
| MSC_HPCA_3___351 | Aran_et_al_2017 | ATP6V1C1, CYC1, DCTD, DDOST, EEF1D, ERCC1, GSTM5, HIC1, IMPDH1, KIF22, LAMP1, MDH2, NDUFA8, NDUFB4, NFATC4, PARN, POLR2G, PTPN11, RPL8, RPN1, SLC35A2, AAAS, TMEM11, HAND2, EIF4E2, BAG3, RNF7, SNX17, CUL7, SCAMP3, RNF41, STUB1, TMEM147, TIMM44, VTI1B, YIF1A, KDELR1, LMAN2, RER1, EMILIN1, SNF8, COPZ1, HMGXB3, CLUAP1, IQSEC2, PMPCA, CABIN1, MKRN2, LMOD1, TMEM184B, TCTN3, TIMM10, SPCS1, MRPS18B, GMPPA, STOML2, CYHR1, ARL6IP4, ZNF771, MRPS17, TRMT112, CUTA, TMEM161A, COMMD4, CPSF3L, OGFOD1, LRRC59, ZNF446, PRX, MRPS11, CUEDC2, C7orf26, TMEM223, SPAG16, MRPL24, TBC1D17, NHEJ1, PODNL1, INTS5, SF3B5, MFSD5, THAP3, SLC38A10, SIX5, GTF2H5 |

| | | |
|---|---|---|
| mv_Endothelial_cells_ENCODE_1___352 | Aran_et_al_2017 | ACVRL1, ANGPT2, SLC25A6, RHOC, TSPO, CAV2, CCNG1, CD34, CETP, COX4I1, CSNK2A2, DYNC1LI2, ECHS1, FDPS, FLT4, GJA4, GOLGA3, GOT2, HADHA, HCFC1, HCRTR1, HOXD3, ILF2, ITGA9, KDR, LGALS1, NOTCH4, NOVA2, PLXNB3, MAPK3, PRPSAP1, PSMB7, PSMC5, PSMD8, RALA, RANGAP1, MAPK12, SELE, SH3GL1, SMARCD1, SMARCE1, SUMO3, SNAPC4, SNTB2, TARBP2, TBX1, TEK, TIE1, CLDN5, TPM3, UBE2E1, UFD1L, VWF, YWHAE, ZNF205, CLPP, ZNF282, SCARF1, HYAL2, MBTPS1, PABPC4, RGS11, BANF1, EIF2B2, MTMR2, BCL10, TNFSF18, DYRK1B, COX7A2L, SLC24A1, GTF3C5, VAMP3, TAOK2, EI24, PPM1F, DHX38, KEAP1, MED16, NUBP2, ACTR1A, TXNDC9, RAMP3, PCGF3, LYPLA1, STK25, SEMA6B, DCTN2, HTATIP2, GIPC1, LYVE1, RNPS1, COPS6, RAB35, TMEM115, STRAP, ATXN2L, LYPLA2, TUSC2, RRAS2, ARHGEF15, RALY, TTLL5, N4BP3, STAB1, ZC3H7B, SUN1, ABCB9, KCTD2, DNPEP, EDC4, MTCH1, MTCH2, PRKD2, GPKOW, CECR5, MRPS28, GIT1, MRPL15, CNIH4, F11R, MRPS16, CLEC1A, BFAR, GMPR2, PCDH12, PIAS4, FAM96B, HSPB11, EMCN, SOX18, ROBO4, NDUFB11, DEF8, CPSF3L, EXD2, CHST12, LMBR1L, UBAP2, CISD1, METTL3, INPP5E, CIAPIN1, FN3K, MMS19, RNF25, WDR13, TUT1, MRPL9, SPATS2, DDA1, DCTPP1, FAM65A, CXorf36, GEMIN7, MMRN2, FAM124B, EDC3, MYCT1, GFOD2, URM1, FRMD8, MTG1, FTSJ3, SAMD14, ZDHHC24, KANK3 |
| mv_Endothelial_cells_ENCODE_2___353 | Aran_et_al_2017 | ACVRL1, ANGPT2, SLC25A6, RHOC, ATP1B3, BYSL, TSPO, CAV2, CD34, CDC27, CETP, CLIC1, COPA, COX4I1, CSNK2A2, DYNC1LI2, FDPS, FLT4, GDI2, GOT2, GYG1, HCFC1, HCRTR1, HOXD3, ILF2, KDR, LAMP1, LGALS1, MTHFR, MYL6, NAP1L4, NDUFC2, NOTCH4, NOVA2, PLD2, PLXNB3, POLR2F, MAPK3, PRPSAP1, PSMB7, PSMC5, ABCD4, RALA, RANGAP1, RPL4, RPN2, MAPK12, SELE, SH3GL1, SMARCD1, SMARCE1, SNAPC4, SNTB2, SSBP1, TARBP2, TBX1, TEK, TIAL1, TIE1, CLDN5, TPM3, UFD1L, YWHAE, ZFPL1, CLPP, ZNF282, SCARF1, HYAL2, MBTPS1, PABPC4, EIF2B2, MTMR2, TNFSF18, DYRK1B, COX7A2L, SLC24A1, TAOK2, TXNL1, EIF4E2, DHX38, KEAP1, ACTR1A, TXNDC9, DCAF7, RAMP3, PCGF3, LYPLA1, STK25, KAT5, DCTN2, HTATIP2, GIPC1, FRS3, LYVE1, COPS6, ATF7, RAB35, TMEM115, STRAP, TUSC2, RRAS2, ARHGEF15, TTLL5, N4BP3, ZC3H7B, EXOC7, SIN3B, ABCB9, KCTD2, EDC4, STX12, MTCH1, MTCH2, PGLS, PRKD2, TBC1D10B, FBXO22, GPKOW, AHDC1, EIF3K, CECR5, MRPS28, GIT1, MRPL15, CNIH4, UBIAD1, F11R, MRPS16, UBXN1, THAP4, TACO1, CLEC1A, BFAR, GMPR2, PCDH12, PIAS4, FAM96B, HSPB11, EMCN, INPP5K, SOX18, ROBO4, NDUFB11, GPN2, DEF8, CPSF3L, C19orf24, ZWILCH, TMEM39B, EXD2, CHST12, LMBR1L, UBAP2, CISD1, METTL3, INPP5E, CIAPIN1, ENOPH1, CCDC90B, FN3K, MOSPD3, WDR13, ELOVL1, SPATS2, FAM65A, MUL1, CXorf36, GEMIN7, GSDMD, MMRN2, FAM124B, EDC3, MYCT1, URM1, FRMD8, FTSJ3, TOR1AIP2, SAMD14, SENP5, ZDHHC24, KANK3, TIPRL |
| mv_Endothelial_cells_ENCODE_3___354 | Aran_et_al_2017 | ACVRL1, ANGPT2, SLC25A6, RHOC, CAV2, CD34, CETP, COX4I1, CSNK2A2, DYNC1LI2, FDPS, FLT4, GJA4, GOLGA3, GOT2, HADHA, HCFC1, HCRTR1, HOXD3, ILF2, ITGA9, KDR, LGALS1, NOTCH4, NOVA2, PLXNB3, MAPK3, PRPSAP1, PSMB7, PSMC5, RALA, RANGAP1, MAPK12, SELE, SH3GL1, SMARCD1, SMARCE1, SUMO3, SNAPC4, SNTB2, TARBP2, TBX1, TEK, TIE1, CLDN5, TPM3, UFD1L, VWF, ZNF205, CLPP, ZNF282, HYAL2, RGS11, EIF2B2, BCL10, TNFSF18, DYRK1B, SLC24A1, GTF3C5, TAOK2, EI24, PPM1F, KEAP1, MED16, RAMP3, PCGF3, LYPLA1, STK25, SEMA6B, HTATIP2, GIPC1, LYVE1, COPS6, RAB35, TMEM115, STRAP, ATXN2L, LYPLA2, TUSC2, RRAS2, ARHGEF15, RALY, TTLL5, N4BP3, STAB1, ZC3H7B, ABCB9, KCTD2, EDC4, MTCH1, MTCH2, PRKD2, CECR5, MRPS28, GIT1, MRPL15, F11R, MRPS16, CLEC1A, BFAR, GMPR2, PCDH12, HSPB11, EMCN, SOX18, ROBO4, NDUFB11, CPSF3L, CHST12, LMBR1L, CISD1, METTL3, CIAPIN1, FN3K, WDR13, TUT1, MRPL9, SPATS2, DDA1, DCTPP1, FAM65A, CXorf36, GEMIN7, MMRN2, FAM124B, EDC3, MYCT1, GFOD2, URM1, FRMD8, MTG1, FTSJ3, SAMD14, KANK3 |
| mv_Endothelial_cells_FANTOM_1___355 | Aran_et_al_2017 | ACVRL1, ANGPT2, ART4, ARVCF, BMX, CALM1, CASP10, CAV1, CD9, CD34, ENTPD1, CDK9, CETP, CSF2RB, CTNNA1, ACE, ELK4, ERG, ERH, FLOT2, FLT1, FLT4, GABPB1, HDAC1, HOXD3, HTR1B, IL3RA, KDR, LYL1, MAP3K3, MGAT5, NCK1, NOTCH4, NOVA2, PNP, PDCL, PIK3CG, PLCG1, PRPSAP1, RALA, RALB, SELE, TAL1, TBX1, TEK, TIE1, CLDN5, UFD1L, VWF, ZNF22, SCARF1, HYAL2, BCL10, HERC1, ATP6V0E1, TNFSF18, LRRFIP1, TAOK2, PPM1F, RAMP3, SEMA6C, SEMA6B, HTATIP2, MYL12A, LYVE1, TFEC, ARHGEF15, MMRN1, CD93, ATF6, N4BP3, STAB1, TDRD7, TSPAN13, MAT2B, CLEC1A, PCDH12, SPTBN5, EMCN, SOX18, ROBO4, RASIP1, TMEM39B, GIMAP4, LSG1, NECAP2, LMBR1L, ANO2, KIF17, TMEM109, CXorf36, MMRN2, FAM124B, MYCT1, RNF34, GFOD2, PLVAP, CEACAM21, KANK3, GIMAP6 |
| mv_Endothelial_cells_FANTOM_2___356 | Aran_et_al_2017 | ACVRL1, ANGPT2, ARVCF, BMX, CASP10, CAV1, CD9, CD34, ENTPD1, CETP, CSF2RB, ACE, ERG, FLT1, FLT4, HTR1B, IL3RA, KDR, LYL1, MAP3K3, NOTCH4, PNP, NPR1, TAL1, TBX1, TEK, TIE1, CLDN5, VWF, SCARF1, HYAL2, TNFSF18, TAOK2, PPM1F, RAMP3, SEMA6B, LYVE1, TFEC, ARHGEF15, MMRN1, CD93, ATF6, N4BP3, STAB1, |

| | | |
|---|---|---|
| | | CLEC1A, PCDH12, EMCN, SOX18, ROBO4, RASIP1, GIMAP4, ANO2, KIF17, CXorf36, MMRN2, FAM124B, MYCT1, KANK3, GIMAP6 |
| mv_Endothelial_cells_FANTOM_3___357 | Aran_et_al_2017 | ACTG1, ADRA1B, ANXA2, ARF1, RHOA, RHOC, ATP2B3, BMX, PTTG1IP, CAV1, CCKAR, CD34, AP2M1, CLTA, CPA1, CTNNA1, DAD1, DLST, DYNC1H1, EIF4G1, FOXC2, GJA4, GPR4, HSPA4, HSP90AB1, MAGEB1, MIF, MIP, MYL6, 44076, NEDD8, NOTCH4, OXA1L, P4HB, PLS3, POLR2J, PPP2R1A, PPP2R2A, MAPK3, PSMB7, PSMD1, PSMD10, RALA, RCN2, SLC6A7, TAF12, TJP1, CLDN5, DNAJC7, UFD1L, VWF, KCNAB1, USP5, CLPP, AKAP4, EIF2B2, TAOK2, MED20, SAE1, ABCF2, ACTR1A, B3GALT5, CLEC4M, PCGF3, SEMA6B, GLRX3, ARPC1A, YKT6, ACTL7A, COPS6, TMEM115, CAPN11, PWP1, CDC37, FAM107A, STRAP, LYPLA2, ECD, TUSC2, NCBP2, CD93, GANAB, PMPCA, NUP188, CLDN14, ARL2BP, PRND, PITPNB, MTCH1, TRPC4AP, NOC2L, FOXD3, TRAPPC3, SEC61A1, CHCHD2, PCDH12, SPTBN5, SNTG2, DDX56, TMED9, FNDC8, COMMD4, C1orf123, AURKAIP1, UNC45A, PCDHA6, C16orf62, LYZL6, MRPL17, MMRN2, FAM124B, LRRC3, DCTN5, YIF1B, POM121L2 |
| mv_Endothelial_cells_HPCA_1___358 | Aran_et_al_2017 | ACVRL1, CETP, RANGAP1, SELE, TIE1, CLDN5, VWF, HYAL2, EIF2B2, TNFSF18, LYVE1, ARHGEF15, CLEC1A, ROBO4, CXorf36, MMRN2, KANK3 |
| mv_Endothelial_cells_HPCA_2___359 | Aran_et_al_2017 | ACVRL1, CETP, FLT4, HCRTR1, KDR, NOVA2, RALA, RANGAP1, SELE, TEK, TIE1, CLDN5, TPM3, VWF, HYAL2, EIF2B2, TNFSF18, LYVE1, ARHGEF15, CLEC1A, PCDH12, SOX18, ROBO4, TUT1, FAM65A, CXorf36, MMRN2, MYCT1, KANK3 |
| mv_Endothelial_cells_HPCA_3___360 | Aran_et_al_2017 | ACVRL1, CETP, FLT4, RANGAP1, SELE, TIE1, CLDN5, VWF, HYAL2, EIF2B2, TNFSF18, LYVE1, ARHGEF15, CLEC1A, SOX18, ROBO4, TUT1, CXorf36, MMRN2, KANK3 |
| Myocytes_ENCODE_1___361 | Aran_et_al_2017 | EVC, SMAD5, MUSK, SGCA, SIM1, BAG2, SHQ1, IMPACT, EXOC1, KRTAP1-1 |
| Myocytes_ENCODE_2___362 | Aran_et_al_2017 | COPB1, EVC, SMAD5, MUSK, MYF5, SGCA, SIM1, MAP3K7, DENR, BAG2, SHQ1, IMPACT, EXOC1, XPNPEP3, KRTAP1-1, PRRC1 |
| Myocytes_ENCODE_3___363 | Aran_et_al_2017 | ACTG1, ALDOA, ANXA5, ARF4, CAD, CAST, CAV1, CAV3, CCNG1, CCT6A, CDH15, CHRNG, CLTC, COPB1, COX8A, CSNK1G3, DCTD, DDOST, TOR1A, EIF4EBP2, EPRS, ERCC4, EVC, GDI2, GLE1, GNAS, GRSF1, GTF2H1, HIF1A, HRC, HSPA8, IDUA, KIF2A, TNPO1, LAMP1, LGALS1, SMAD2, SMAD5, MIF, MKLN1, MMP11, MSH3, MUSK, MYBPH, MYF5, MYOD1, NDUFA10, 44076, NONO, PAFAH1B1, PARN, PCDHGC3, SLC25A3, PLXNB3, POLR2J, PSMB4, PTPN11, RAPSN, RCN2, RPN2, SGCA, FBXW4, SIM1, SPG7, SSR4, SS18, STAU1, MAP3K7, TIAL1, UBE3A, COL14A1, ZNF37A, ZNF221, ZNF214, RAB7A, DENR, WISP1, EIF2S2, USP14, BAG5, BAG2, PREPL, SNX17, UBAP2L, UBA2, RAD50, LRPPRC, MPHOSPH6, ZMPSTE24, BCKDK, YAP1, CDIPT, TMEM147, IPO7, PTGES3, FRS2, CPSF4, COPS8, SPIN1, AFG3L2, HNRNPA0, ERP29, METAP2, XPOT, HMGXB3, CLUAP1, POFUT2, DNAJC16, HARS2, POFUT1, ARL2BP, MKRN2, TXN2, IBTK, TCTN3, FBXL4, MYOF, RPS6KC1, RNF11, NPTN, SND1, HSPB7, TUBG2, VPS4A, NFU1, UTP20, MCTS1, MYLPF, EPN1, SEC61A1, MRPS16, IFT52, METTL9, MBTPS2, UCHL5, RWDD1, AMOTL2, ERGIC3, UBE2D4, CMPK1, NUDT9, NUP54, GPR173, RC3H2, CHCHD3, FAM120C, ZDHHC4, SHQ1, SLC38A7, ADI1, QRSL1, IMPACT, NOP10, HIF1AN, ZNF446, PEX26, IARS2, EXOC1, SPATA7, ACTR10, DNAH7, SPPL2B, PRX, EDA2R, CCDC90B, GUF1, PKNOX2, XPNPEP3, NUCKS1, FAM160B2, DCLRE1B, MRPS11, C2orf47, SPAG16, PALB2, TBC1D17, ZNF668, METTL8, C10orf88, MYO19, RUFY1, INTS5, KRTAP1-1, TM2D1, ACTR8, SLC38A10, PRRC1, MYL6B, YTHDF3 |
| Myocytes_FANTOM_1___364 | Aran_et_al_2017 | ALPL, CASQ2, CDH15, DES, HAS1, MYBPH, MYF5, MYH7, MYL1, MYL4, MYOG, RAPSN, SGCA, TNNI1, TNNT2, TTN, MYLPF |
| Myocytes_FANTOM_2___365 | Aran_et_al_2017 | ACTA1, ACTN2, ALPL, CDH15, CKM, MSTN, HAS1, HRC, MUSK, MYBPH, MYF5, MYH1, MYH2, MYH7, MYH8, MYL1, MYL4, MYOD1, MYOG, RAPSN, ROS1, SGCA, SGCG, SLN, TNNC2, TNNI1, TNNT2, ATP1B4, HEYL, ITGB1BP2, MYLPF, GSG1 |
| Myocytes_FANTOM_3___366 | Aran_et_al_2017 | CDH15, HAS1, MYF5, MYH7, MYL1, MYL4, MYOG, RAPSN, TNNI1, TNNT2 |
| naive_Bcells_BLUEPRINT_1___367 | Aran_et_al_2017 | BLK, CXCR5, CD19, MS4A1, CD72, SPIB, TCL1A, FCRL2 |
| naive_Bcells_BLUEPRINT_2___368 | Aran_et_al_2017 | BLK, CXCR5, CD19, MS4A1, CD22, CD72, CD79B, CCR6, GPR18, CD180, MGAT5, SPIB, TCL1A, MBD4, TSPAN13, UTP6, FCRL2, TREML2 |
| naive_Bcells_BLUEPRINT_3___369 | Aran_et_al_2017 | BLK, CXCR5, CD19, MS4A1, CD72, CD180, SPIB, TCL1A, FCRL2 |
| naive_Bcells_HPCA_1___370 | Aran_et_al_2017 | BLK, CD19, MS4A1, CD22, CD37, CD72, CD79A, CSNK1G3, DSP, FCER2, GMFB, PNOC, SNX2, GCM1, AP3B1, MBD4, STAG3, PRDM4, PWP1, RRAS2, GGA2, SIPA1L3, STAP1, P2RY10, DEF8, MFN1, FCRL2, EGOT |
| naive_Bcells_HPCA_2___371 | Aran_et_al_2017 | CD1A, CD19, MS4A1, CD22, CD37, CD72, CD79A, CSNK1G3, DSP, FCER2, GMFB, MGAT5, PNOC, SNX2, AP3B1, MBD4, STAG3, PRDM4, PWP1, RRAS2, GGA2, |

| | | |
|---|---|---|
| | | SIPA1L3, STAP1, P2RY10, VPREB3, DEF8, MFN1, FCRL2, EGOT |
| naive_Bcells_HPCA_3___372 | Aran_et_al_2017 | BLK, CAPN3, CD1A, CD19, MS4A1, CD22, CD37, CD72, CD79A, CD79B, CSNK1G3, DSP, FCER2, GMFB, HSPA4, MGAT5, PNOC, SNX2, GCM1, AP3B1, MBD4, STAG3, PRDM4, PWP1, SP140, RRAS2, GGA2, SIPA1L3, STAP1, P2RY10, VPREB3, DEF8, MFN1, FCRL2, C10orf76, SMC6, MCM9, EGOT |
| naive_Bcells_NOVERSHTERN_1___373 | Aran_et_al_2017 | CXCR5, BMP3, CACNA1F, CAPN3, CD19, MS4A1, CD22, CD72, COL19A1, CSNK1G3, DAZL, DSP, FCER2, GNG3, GPR18, MATN1, MAP3K9, MMP17, MYBPC2, PAX5, PHKG1, PYGM, ZNF154, PRDM2, USP7, TCL1A, SYN3, ADAM20, USP6, AKAP6, TCL1B, BCL2L10, FRS2, PRDM4, RRAS2, GGA2, SIPA1L3, TCL6, TSPAN13, P2RY10, MYO3A, SDK2, WDR74, UBE2O, RBM15, SMC6, KHDRBS2 |
| naive_Bcells_NOVERSHTERN_2___374 | Aran_et_al_2017 | CXCR5, BMP3, CACNA1F, CAPN3, CD19, MS4A1, CD22, CD72, COL19A1, CSNK1G3, DAZL, FCER2, GNG3, MAP3K9, MMP17, MYBPC2, PAX5, PHKG1, PRDM2, USP7, TCL1A, SYN3, ADAM20, USP6, AKAP6, TCL1B, FRS2, PRDM4, RRAS2, GGA2, SIPA1L3, TCL6, TSPAN13, P2RY10, WDR74, UBE2O, SMC6, KHDRBS2 |
| naive_Bcells_NOVERSHTERN_3___375 | Aran_et_al_2017 | RERE, CXCR5, BMP3, CACNA1F, CAPN3, CD1A, CD19, MS4A1, CD22, CD72, COL19A1, CSNK1G3, DAZL, DSP, FCER2, GH1, GNG3, HLA-DOA, LY9, MATN1, CIITA, MAP3K9, MMP17, MYBPC2, PAX5, PGAM2, PHKG1, POU2F1, PRKCB, PYGM, RB1, TRA2B, ZNF154, SLC30A4, PRDM2, USP7, CUBN, TCL1A, SYN3, CDK13, PTCH2, ADAM20, USP6, AKAP6, TCL1B, TBC1D5, BCL2L11, STAG3, FRS2, PRDM4, RRAS2, GGA2, SIPA1L3, N4BP3, TCL6, TSPAN13, P2RY10, SNTG2, SDK2, WDR74, UBE2O, NOC3L, RBM15, FCRL2, SMC6, TRAPPC9, PIKFYVE, KHDRBS2, 43898 |
| Neurons_ENCODE_1___376 | Aran_et_al_2017 | EPHA3, GNG3, INSM1, KCNQ2, NEUROD2, PCDH8, ACTL6B, CAMKV, STMN4 |
| Neurons_ENCODE_2___377 | Aran_et_al_2017 | ABCA3, ACVR2B, ADRA2A, CACNA1B, CACNB3, CHRNB2, CPE, CRABP1, CRMP1, NCAN, CTNNA1, EFNB3, CELSR3, ELAVL3, EPHA3, FOXG1, GAD2, GAP43, GEM, GNG3, GRIK3, GRM2, ID1, INSM1, IREB2, KCNN1, KCNQ2, KCNQ3, KLC1, MEIS2, MLF1, MLLT3, MN1, CD200, MVD, NEUROD2, NNAT, NPY, NPTX1, NTRK3, PCDH8, PLXNA2, POU3F1, POU3F2, POU3F3, PKIA, MAPK8, PTPRZ1, PTX3, RPE65, SCN3A, CXCL12, SFRP4, SH3GL2, SH3GL3, SOX4, SOX11, SPAST, ZNF354A, THRA, ZFP37, ZNF711, ZNF14, ZNF43, ZNF195, ZNF223, TUBA1A, ST8SIA4, PDHX, ST8SIA2, HIST1H3D, DCHS1, STX16, B3GALT2, CDK5R1, BSN, HAP1, CYTH2, LHX2, IPO13, NUP93, TSPAN2, CFDP1, SEMA6C, ZNF211, IFI44, DPYSL4, SCGN, PNMA2, ZBTB6, MLLT11, TMSB15A, STMN2, C14orf1, ZFP30, ZNF510, WDR47, KIF21B, ARC, KIAA1107, AGTPBP1, CUX2, SEZ6L, SULT4A1, SETBP1, LRRTM2, RANBP6, CACNG5, CACNG4, PDLIM3, PCSK1N, KCNMB4, SCG3, HUNK, PODXL2, PARD6A, ZNF117, ACTL6B, GPR173, FAM105A, LRRN3, FNBP1L, ENOX1, AGPAT5, AP1AR, ZNF821, ANKRD10, ZNF415, BEX1, PCDHB11, ZNF253, C21orf62, KCNK12, ZNF529, PTBP2, ZBED5, NEUROG2, NEUROD6, REEP1, RASL11B, GDAP1L1, CAMKV, NKAIN1, RNF219, ZMAT4, VASH2, ZNF669, PGAP1, ZNF606, ZNF614, ZNF430, ZSCAN16, ZNF34, STMN4, ZNF484, YIPF4, MUM1, C16orf45, ZNF682, C1orf216, EMID1, PAQR3, ZNF675, TET3, KCTD13, ZNF493 |
| Neurons_ENCODE_3___378 | Aran_et_al_2017 | ABCA3, ADRA2A, CACNA1B, CHRNB2, CPE, CRABP1, CRMP1, NCAN, CTNNA1, EFNB3, CELSR3, ELAVL3, EPHA3, FOXG1, GAD2, GAP43, GEM, GNG3, GRIK3, GRM2, INSM1, KCNQ2, KCNQ3, KLC1, MLLT3, CD200, NEUROD2, NNAT, NPY, NPTX1, PCDH8, PLXNA2, POU3F1, POU3F3, PKIA, MAPK8, PTPRZ1, SH3GL2, SH3GL3, SOX11, ZFP37, ZNF14, TUBA1A, ST8SIA4, PDHX, B3GALT2, BSN, HAP1, CYTH2, TSPAN2, SCGN, ZBTB6, MLLT11, TMSB15A, STMN2, WDR47, KIF21B, ARC, KIAA1107, CUX2, SEZ6L, SULT4A1, PCSK1N, SCG3, HUNK, PODXL2, ACTL6B, FAM105A, ENOX1, ZNF821, ZNF415, BEX1, KCNK12, PTBP2, ZBED5, NEUROG2, NEUROD6, REEP1, RASL11B, GDAP1L1, CAMKV, NKAIN1, RNF219, ZMAT4, PGAP1, ZNF614, ZNF34, STMN4, YIPF4, EMID1, ZNF675 |
| Neurons_FANTOM_1___379 | Aran_et_al_2017 | ALDOC, ATP1A3, ATP1B1, ATP2B2, ATP6V1G2, CA8, CACNB4, CALB1, CAMK2B, CDH18, COX7A1, COX7B, DGKB, DEFB1, DLG2, DPP6, FABP6, FGF9, FGF12, GABBR1, GABRA1, GABRB2, GABRG2, GAD1, GNAO1, GNG3, GPM6A, GRIA2, GRID2, GRIK1, GRM1, GRM7, ID2, ITPR1, KCNC1, KIF5C, LPL, MT3, NDUFA5, NEFM, NEFH, NELL1, NEFL, NPPC, NPTX1, OMG, PCDH9, PCP4, SERPINI1, PRKCG, PTPRR, PVALB, RGS16, RORA, RTN1, SH3GL2, SLC6A1, SNAP25, SNCG, SYP, TAC1, SEC62, TSPAN7, TRPC3, ZNF208, AP3B2, SPARCL1, CACNA1G, INA, NRXN1, ELMO1, SNAP91, C1orf61, STMN2, CNKSR2, FAIM2, ARHGAP26, FSTL4, KIAA1107, ACSL6, SEZ6L, SMPX, SLC24A2, SCG3, DDX25, SPOCK3, ST8SIA3, GPRC5B, GNG13, TM6SF1, FXYD7, IL20RA, LRRN3, SUSD4, BEX1, PRMT8, DNAJC12, ANKS1B, PLXDC1, SLC12A5, KLHL1, REEP1, GPR63, STMN4, TRIM9, TCEAL2, SHISA6 |

| | | |
|---|---|---|
| Neurons_FANTOM_2___380 | Aran_et_al_2017 | ACYP2, AGTR2, ALDOC, ATP1A3, ATP1B1, ATP1B2, ATP2A3, ATP2B2, ATP6V1G2, CA7, CA8, CACNA1A, CACNB2, CACNB4, CALB1, CAMK2B, CDH18, CHGB, COX7A1, COX7B, DAB1, DACH1, DGKB, DGKG, DEFB1, DLG2, DYNC1I1, DPP6, FABP3, FABP6, FGF9, FGF12, FGF14, GABBR1, GABRA1, GABRB2, GABRB3, GABRG2, GAD1, GAD2, GNAO1, GNG3, GNG4, GPM6A, GRIA2, GRIA3, GRID2, GRIK1, GRM1, GRM7, HSBP1, HTR5A, ID2, ITPR1, KCNA2, KCNC1, KCNK1, KIF5A, KIF5C, KPNA5, LPL, MT3, NAP1L2, NCAM1, NDUFA3, NDUFA4, NDUFA5, NEFM, NEFH, NELL1, NEFL, NELL2, NPPC, NPTX1, NRCAM, OMG, PCDH9, PCP4, PDE9A, PEG3, SERPINI1, PPP3CA, PRKCG, MAPK10, PTPRR, PVALB, RAB3A, RGS7, RGS16, RORA, RTN1, RYR2, SCN1A, SCN2A, SH3GL2, SLC1A6, SLC6A1, SLC8A1, SNAP25, SNCG, ABCC8, VAMP2, SYP, TAC1, SEC62, TSPAN7, TRPC3, ZNF208, KCNAB1, AP3B2, PIP5K1B, SPARCL1, PPFIA2, SYNJ1, CACNA1G, INA, CACNA2D2, CPNE6, RAB33A, NRXN3, NRXN1, AKAP7, GABBR2, GPRASP1, JAKMIP2, ELMO1, SNAP91, SAP18, CORO2B, C1orf61, ATP5L, STMN2, KIF3A, HHLA3, MGAT4A, GABARAPL2, CNKSR2, KIFAP3, SV2C, FAIM2, MYT1L, ARHGAP26, FSTL4, KIAA1107, ACSL6, SEZ6L, SMPX, SLC24A2, SULT4A1, DNM3, GALNT8, CLUL1, PCDH17, GOLIM4, KCNMB4, PCLO, NDUFAF4, SCG3, DDX25, NME7, SPOCK3, ST8SIA3, PIGP, GPRC5B, GNG13, BCL11A, TM6SF1, FXYD7, IL20RA, FAM134B, LRRN3, LRRC49, SUSD4, CISD1, BEX1, PRMT8, DNAJC12, ANKS1B, PLXDC1, TTYH1, SLC12A5, KLHL1, NDRG4, REEP1, FAM184A, ZNF385D, MAP9, CEP76, GPR63, STMN4, HOPX, TRIM9, IQCK, RALYL, TCEAL2, THSD7A, LPCAT4, FAM21A, SHISA6, |
| Neurons_FANTOM_3___381 | Aran_et_al_2017 | ACYP2, ALDOC, ANK2, ATP1A3, ATP1B1, ATP2A3, ATP2B2, ATP6V1G2, CA8, CACNA1A, CACNB2, CACNB4, CALB1, CAMK2B, CDH18, CHGB, CHN1, COX7A1, COX7B, CRMP1, DGKB, DGKG, DEFB1, DLG2, DYNC1I1, DPP6, FABP3, FABP6, FABP7, FGF9, FGF12, FGF14, GABBR1, GABRA1, GABRB2, GABRG2, GAD1, GNAO1, GNG3, GPM6A, GPM6B, GRIA2, GRID2, GRIK1, GRM1, GRM7, ID2, ITPR1, KCNC1, KIF5C, LHX1, LPL, MT3, NAP1L2, NCAM1, NDUFA3, NDUFA5, NEFM, NEFH, NELL1, NEFL, NELL2, NOVA1, NPPC, NPTX1, OMG, PCDH9, PCP4, PDE9A, PEG3, SERPINI1, PRKCG, PTPRN, PTPRR, PVALB, RAB3A, RGS16, RORA, RTN1, SCN1A, SH3GL2, SLC1A6, SLC6A1, SLC8A1, SNAP25, SNCG, SORL1, SYP, TAC1, SEC62, TSPAN7, TRPC3, ZNF208, KCNAB1, AP3B2, NME5, SPARCL1, CACNA1G, INA, RAB33A, NRXN3, NRXN1, GABBR2, SPOCK2, JAKMIP2, ELMO1, SNAP91, RCAN2, C1orf61, STMN2, KLK8, MGAT4A, GABARAPL2, CNKSR2, FAIM2, ARHGAP26, FSTL4, KIAA1107, ACSL6, SEZ6L, SMPX, SLC24A2, SOSTDC1, DNM3, PCDH17, NDUFAF4, SCG3, DDX25, SNX10, NME7, SPOCK3, ST8SIA3, PIGP, GPRC5B, GNG13, BCL11A, TM6SF1, FXYD7, IL20RA, FAM134B, LRRN3, LRRC49, SUSD4, BEX1, PRMT8, DNAJC12, ANKS1B, PLXDC1, SLC12A5, KLHL1, NDRG4, REEP1, ZNF385D, CEP76, GPR63, STMN4, HOPX, TRIM9, TCEAL2, SHISA6 |
| Neutrophils_BLUEPRINT_1___382 | Aran_et_al_2017 | CA4, CEACAM3, FCGR3B, CXCR1, CXCR2, PGLYRP1, MMP25, ZDHHC18, TRPM6 |
| Neutrophils_BLUEPRINT_2___383 | Aran_et_al_2017 | CA4, CEACAM3, FCGR3B, CXCR1, PGLYRP1, VNN3, MMP25, ZDHHC18 |
| Neutrophils_BLUEPRINT_3___384 | Aran_et_al_2017 | CA4, CEACAM3, CXCR1, CXCR2, PGLYRP1, P2RY13, MMP25, ZDHHC18 |
| Neutrophils_FANTOM_1___385 | Aran_et_al_2017 | APAF1, CBL, CEACAM3, CSF2RB, DDX3X, FCAR, FCGR3B, FPR2, CXCR2, NFYA, PAK2, PTEN, SLC19A1, UBE2B, BEST1, ELL, MTMR3, HERC3, PGLYRP1, SLC25A44, DHX34, TOX4, TECPR2, WWP2, LMTK2, WDFY3, MED13L, ACAP2, CLEC4E, UBN1, NRBF2, TREM1, HRH4, RMND5A, TMUB2, TMEM185B, GCC1, TREML2, BTNL8, NDEL1, ZDHHC18, UBXN2B |
| Neutrophils_FANTOM_2___386 | Aran_et_al_2017 | APAF1, CA4, CASP5, CBL, CEACAM3, CEACAM8, CSF2RB, DDX3X, FCAR, FCGR3B, FPR2, CXCR2, MAK, NFYA, PTEN, SDF2, SLC19A1, TGM3, TOP1, UBE2B, UBE2D1, BEST1, TRIM25, ELL, KSR1, MTMR3, HERC3, PGLYRP1, SPAG9, NMI, CIR1, TTLL4, SLC25A44, DHX34, IP6K1, TOX4, TECPR2, USP15, CAMKK2, LILRA1, WWP2, BTN2A1, LMTK2, WDFY3, MED13L, ACAP2, PADI4, CLEC4E, UBN1, NRBF2, TREM1, HRH4, TMUB2, GCC1, TREML2, BTNL8, FBXO38, NDEL1, ZDHHC18, UBXN2B |
| Neutrophils_FANTOM_3___387 | Aran_et_al_2017 | APAF1, CBL, CEACAM3, CSF2RB, DDX3X, FCGR3B, FPR2, CXCR2, NFYA, PTEN, SLC19A1, UBE2B, BEST1, ELL, MTMR3, HERC3, PGLYRP1, DHX34, TECPR2, WWP2, LMTK2, WDFY3, MED13L, ACAP2, CLEC4E, UBN1, NRBF2, TREM1, HRH4, TMUB2, GCC1, TREML2, BTNL8, NDEL1, ZDHHC18, UBXN2B |
| Neutrophils_HPCA_1___388 | Aran_et_al_2017 | CLC, CSF3R, FCGR3B, FPR2, HBB, CXCR2, S100A12, P2RY13, TREM1 |
| Neutrophils_HPCA_2___389 | Aran_et_al_2017 | CLC, CSF3R, FCGR3B, FPR2, HSPA6, CXCR2, S100A12, P2RY13, TREM1 |

| | | |
|---|---|---|
| Neutrophils_HPCA_3___390 | Aran_et_al_2017 | CLC, CSF3R, FCGR3B, FPR2, HBB, HSPA6, CXCR2, S100A12, LILRB2, LILRA2, P2RY13, TREM1 |
| Neutrophils_IRIS_1___391 | Aran_et_al_2017 | CEACAM3, FCGR3B, CXCR1, CXCR2, MEFV, VNN3, MMP25, BTNL8 |
| Neutrophils_IRIS_2___392 | Aran_et_al_2017 | BMX, CA4, CEACAM3, FCGR3B, FPR2, CXCR1, CXCR2, MEFV, AATK, P2RY13, TREM1, VNN3, MMP25, BTNL8, TRPM6 |
| Neutrophils_IRIS_3___393 | Aran_et_al_2017 | BMX, CA4, CEACAM3, FCGR3B, FPR2, GPR27, CXCR1, CXCR2, MEFV, IL18RAP, AATK, P2RY13, TREM1, VNN3, MMP25, BTNL8, TRPM6 |
| NK_cells_BLUEPRINT_1___394 | Aran_et_al_2017 | FASLG, KLRD1, PTGDR, PTPN4, XCL1, TKTL1, IL18RAP, NCR1, ZMYND11, SACM1L, CD244, AGK, DNAJB14 |
| NK_cells_BLUEPRINT_2___395 | Aran_et_al_2017 | FASLG, CX3CR1, GZMB, GZMM, IL2RB, KLRD1, KPNB1, MED1, PRF1, MAPK1, PTGDR, PTPN4, BRD2, XCL1, MAP3K7, TKTL1, TNFSF11, IL18RAP, ZNF264, NCR1, STX8, PJA2, HELZ, GNLY, STAG2, ZMYND11, ZBTB1, SACM1L, TBX21, RAB14, CD244, AGK, DNAJB14, FIP1L1, ARPC5L, HIPK1 |
| NK_cells_BLUEPRINT_3___396 | Aran_et_al_2017 | FASLG, CD247, CTSW, CX3CR1, GZMB, IL2RB, KLRD1, LTA, PRF1, PTGDR, PTPN4, XCL1, TKTL1, TNFSF11, IL18RAP, NCR1, GNLY, ZMYND11, SACM1L, TBX21, CD244, AGK, ZNF426, DNAJB14, HIPK1 |
| NK_cells_FANTOM_1___397 | Aran_et_al_2017 | FASLG, CD247, CX3CR1, GRIK4, GZMB, IL2RB, LIM2, PRF1, PTGDR, TKTL1, NCR1, NMUR1, GNLY, CD160, TBX21 |
| NK_cells_FANTOM_2___398 | Aran_et_al_2017 | IL2RB, PTGDR, XCL1, IL18RAP, NCR1, SACM1L, DNAJB14, ARPC5L |
| NK_cells_FANTOM_3___399 | Aran_et_al_2017 | GZMB, IL2RB, LIM2, PTGDR, NCR1, NMUR1, GNLY, CD160 |
| NK_cells_HPCA_1___400 | Aran_et_al_2017 | BAD, CD247, 44081, CHRNE, DR1, GIPR, GOLGA4, GZMH, GZMB, GZMM, HNRNPL, IFNG, LAG3, MGAT2, MLH1, NEK1, NFE2L2, NKG7, PPP2CA, PRF1, PRKAG1, PTGDR, XCL1, SON, SUPV3L1, TBCC, PRDM2, HIST1H3A, NCR1, PRDX6, YAF2, KLRG1, SF3B4, NMUR1, GNA13, GPATCH8, DNAJC2, ASTE1, ANKRD11, TBX21, AMZ2, THAP1, UBE2Q1, CDKN2AIP, RBM25, DNAJB14, OSBPL7, TSTD2, HIPK1, NCR3, C1orf174 |
| NK_cells_HPCA_2___401 | Aran_et_al_2017 | BAD, CHRNE, CTSW, DR1, GIPR, GOLGA4, GTF3C1, GZMH, IFNG, KLRD1, LAG3, MGAT2, NKG7, PPP2CA, PTGDR, XCL1, FBXW4, SUPV3L1, TSPYL1, HIST1H3A, RGS9, IL18RAP, NCR1, RBM39, NMUR1, GNLY, ZCCHC11, LEMD3, ASTE1, TBX21, IL21R, AMZ2, WBP11, RSRC2, ALG13, COQ10B, HIPK1 |
| NK_cells_HPCA_3___402 | Aran_et_al_2017 | BAD, CD247, 44081, CHRNE, DR1, GIPR, HNRNPL, IFNG, IL2RB, MLH1, NEK1, NKG7, PRF1, PRKAG1, PTGDR, SBF1, CCL4, XCL1, TBCC, HIST1H3A, NCR1, GGPS1, PRDX6, ZBTB39, YAF2, SF3B4, NMUR1, WDR45, GPATCH8, ASTE1, TBX21, IL21R, AMZ2, THAP1, UBE2Q1, CDKN2AIP, OSBPL7, TSTD2, NCR3 |
| NK_cells_IRIS_1___403 | Aran_et_al_2017 | FASLG, KLRD1, PTGDR, PTPN4, XCL1, TKTL1, IL18RAP, NCR1, ZMYND11, SACM1L, CD244, AGK, DNAJB14 |
| NK_cells_IRIS_2___404 | Aran_et_al_2017 | FASLG, CTSW, CX3CR1, GZMM, KLRD1, MED1, PTGDR, PTPN4, XCL1, TKTL1, TNFSF11, IL18RAP, ZNF264, NCR1, ZMYND11, SACM1L, CD244, AGK, DNAJB14 |
| NK_cells_IRIS_3___405 | Aran_et_al_2017 | KLRD1, PTGDR, PTPN4, XCL1, TKTL1, IL18RAP, NCR1, DNAJB14 |
| NK_cells_NOVERSHTERN_1___406 | Aran_et_al_2017 | CHRNE, GIPR, IFNG, HIST1H3A, NCR1, NMUR1, ASTE1, TBX21 |
| NK_cells_NOVERSHTERN_2___407 | Aran_et_al_2017 | CHRNE, GIPR, IFNG, PRKAG1, HIST1H3A, NCR1, NMUR1, ASTE1, AMZ2 |
| NK_cells_NOVERSHTERN_3___408 | Aran_et_al_2017 | GZMH, GZMB, KLRD1, PRF1, NCR1, NMUR1, GNLY, CD160, TBX21 |
| NKT_NOVERSHTERN_1___409 | Aran_et_al_2017 | CASP5, PHKG1, RARA, S100B, BEST1, DOLK, TP53TG5, GMIP, GSG1 |
| NKT_NOVERSHTERN_2___410 | Aran_et_al_2017 | AMBN, CASP5, PHKG1, RARA, S100B, SGCA, TCOF1, BEST1, IL17B, TP53TG5, GMIP, L1TD1 |
| NKT_NOVERSHTERN_3___411 | Aran_et_al_2017 | PHKG1, RARA, BEST1, ARPC1A, TP53TG5, L1TD1, KLHL26, GEMIN7, GSG1, PMFBP1 |
| Osteoblast_FANTOM_1___412 | Aran_et_al_2017 | BMPR1A, COMP, DCTD, DYNC1LI2, EPYC, GRSF1, IBSP, MFAP3, PRELP, SGCG, UBE2L3, COL14A1, ITGA8, EIF2S2, KIF3B, LAPTM4A, APPBP2, MAB21L2, ILVBL, XPOT, DOLK, ICMT, TMEM50A, CBY1, RPS6KC1, COPS7A, MYOZ2, NUDT9, RNF121, LIN7C, DHX32, IL26, STARD7, C5orf15, PAPPA2, EBF2, ATG9A, MUL1, TM2D3, UNC119B, LRRC15 |
| Osteoblast_FANTOM_2___413 | Aran_et_al_2017 | ARCN1, BMPR1A, DYNC1LI2, EPYC, GRSF1, HTR2A, IBSP, MFAP3, PSMB5, SGCG, UBE2L3, COL14A1, ITGA8, KIF3B, EIF4E2, FRMPD4, APPBP2, MAB21L2, YME1L1, |

| | | |
|---|---|---|
| | | ILVBL, XPOT, ICMT, CBY1, RPS6KC1, MYOZ2, RNF121, LIN7C, NPLOC4, IL26, STARD7, PAPPA2, EBF2, FTO, TM2D3, UNC119B, FAM168B, LRRC15, YIPF6 |
| Osteoblast_FANTOM_3___414 | Aran_et_al_2017 | ARCN1, BMPR1A, COMP, DCTD, DYNC1LI2, EPYC, GRSF1, HLCS, IBSP, MFAP3, PRELP, PSMB5, SGCD, SGCG, UBE2L3, COL14A1, ITGA8, EIF2S2, KIF3B, LAPTM4A, FRMPD4, YAP1, APPBP2, MAB21L2, YME1L1, ILVBL, XPOT, DOLK, TRIM32, ZNF629, ICMT, TMEM50A, MKRN2, CBY1, TRAF3IP1, RPS6KC1, COPS7A, AMOTL2, MYOZ2, NUDT9, RNF121, LIN7C, NPLOC4, DHX32, IL26, KCMF1, STARD7, C5orf15, CCDC47, PAPPA2, EBF2, ATG9A, FTO, TMEM185B, MUL1, TM2D3, UNC119B, FAM168B, LRRC15, DPY19L4 |
| Osteoblast_HPCA_1___415 | Aran_et_al_2017 | ACTG1, RHOA, CCKAR, EXTL1, FOXC2, GGCX, GLUD1, HIST1H1A, NFATC4, PGK1, SLC9A5, TUB, ZNF16, GDF5, SSNA1, EIF3G, FIBP, LONP1, ABCF2, RNF41, ARPC1A, MAB21L2, TXNL4A, CBY1, TRPC4AP, CHMP2A, ARL6IP4, TMED9, GLT8D1, PCDHGA11, IFT46, CUEDC2, FIP1L1, SFXN3, TMEM222 |
| Osteoblast_HPCA_2___416 | Aran_et_al_2017 | RHOA, EXTL1, GGCX, NFATC4, ZNF16, GDF5, FIBP, LONP1, ABCF2, MAB21L2, TXNL4A, CHMP2A, ARL6IP4, PCDHGA11, CUEDC2, TMEM222 |
| Osteoblast_HPCA_3___417 | Aran_et_al_2017 | ACTG1, RHOA, CCKAR, AP2S1, ATF6B, DPAGT1, DRG2, EXTL1, FOXC2, GGCX, GLUD1, HIST1H1A, HARS, IARS, EIF6, KIF22, NFATC4, PGK1, SLC9A5, SSR1, TUB, ZFPL1, ZNF16, GDF5, AKR7A2, SSNA1, EIF3G, FIBP, COX7A2L, LONP1, SNX17, ABCF2, SNUPN, RNF41, ARPC1A, MAB21L2, TXNL4A, PARK7, EXOC7, ABCB9, MTCH1, CBY1, LMOD1, TRPC4AP, HSPB7, CHMP2A, EPN1, ARL6IP4, TMED9, GLT8D1, PCDHGA11, IFT46, PRDM11, C12orf43, CUEDC2, ZNF768, CPSF7, FIP1L1, SFXN3, BRIP1, TMEM222 |
| pDC_FANTOM_1___418 | Aran_et_al_2017 | APOC3, CSHL1, DNASE1L3, GZMB, HIST1H2BB, HPD, KCNA5, SCT, P2RY14, LILRB4, CUX2, LRRC36 |
| pDC_FANTOM_2___419 | Aran_et_al_2017 | CSHL1, DNASE1L3, GZMB, HIST1H2BB, KCNA5, RPL3L, SCT, P2RY14, LILRB4, CUX2, LRRC36 |
| pDC_FANTOM_3___420 | Aran_et_al_2017 | CACNB1, DNASE1L3, FKBP2, GZMB, IL3RA, KCNA5, MYBPC1, SCT, SLC12A3, SPIB, ZNF221, MAPKAPK2, P2RY14, CXCL13, LILRB4, SLITRK3, CUX2, TSPAN13, SPCS1, TLR7, KCNK10, KCTD5, LRRC36, CELA2A, CCR2 |
| pDC_NOVERSHTERN_1___421 | Aran_et_al_2017 | FLT3, FUT7, GZMB, IDH3A, SCT, SPIB, CD2AP, TLR7, PTCRA |
| pDC_NOVERSHTERN_2___422 | Aran_et_al_2017 | FLT3, FUT7, GZMB, IDH3A, SCT, SPIB, CD2AP, TLR7, KCNK10, PTCRA |
| pDC_NOVERSHTERN_3___423 | Aran_et_al_2017 | RUNX2, FLT3, FUT7, CXCR3, GZMB, IDH3A, SCT, SPIB, TACR1, CD2AP, TLR7, KCNK10, PTCRA |
| Pericytes_ENCODE_1___424 | Aran_et_al_2017 | ADCY3, CDK4, GGCX, HIC1, MLLT1, MYBPC2, P4HB, MAPK3, PTGER1, SLC6A13, TAF15, DYNLL1, ASH2L, ZBTB22, S1PR2, BCKDK, COPS6, TMED1, CBX6, TXN2, TMEM184B, ERGIC3, MIER2, ZNF444, PRDM11 |
| Pericytes_ENCODE_2___425 | Aran_et_al_2017 | ADCY3, ARF5, ARNT, ATP5G1, ATP5J, CALR, CDK4, CTBP1, DMWD, ELK1, GGCX, HIC1, ISLR, KIF22, MLLT1, MMP11, MYBPC2, MYBPH, NDUFB1, NDUFB7, NNAT, P4HB, PFN1, PPP2CB, PPP2R5D, MAPK3, PSMD8, PTGER1, TRIM27, SLC6A13, TNXB, SLC35A2, ZFPL1, TAF15, LZTR1, AKR7A2, DYNLL1, AP3D1, ASH2L, KCNAB3, ZBTB22, S1PR2, MED16, CTDSP2, ARFRP1, RNF41, BCKDK, TAB1, TIMM44, TM9SF1, TRIM3, GIPC1, YIF1A, KDELR1, LMAN2, COPS6, TMED1, RAB35, TMEM115, LZTS1, WDR6, COPZ1, MLXIP, RALY, CIC, USP22, CBX6, LMOD1, TXN2, TMEM184B, ZNF500, IRF2BP1, ERAL1, VPS4A, AHDC1, C19orf53, MYLPF, GMPPA, ERGIC3, GAR1, MIER2, TMEM161A, ATP5SL, ZNF444, PCDHGA9, PRDM11, PKNOX2, C11orf95, C7orf26, TSEN34, TMEM185B, ZNF768, HPS6, ADAMTS12, IL17RC, TOM1L2, TXLNA |
| Pericytes_ENCODE_3___426 | Aran_et_al_2017 | ATP5J, BCS1L, CALR, DPAGT1, ELK1, GNAS, HIC1, DNAJC4, MMP11, MYBPC2, NDUFA8, NDUFB1, NDUFB7, NDUFS5, P4HB, PPIB, PPP2R5D, PSKH1, PSMD8, PTGER1, RAB5B, DPF2, TRIM27, RPN1, RXRB, SH3GL1, SLC6A4, SLC6A13, SSR2, SLC35A2, ZFPL1, USP5, LZTR1, CDK10, KHSRP, AKR7A2, DYNLL1, EDF1, AP3D1, ASH2L, KCNAB3, ZBTB22, SEC24C, SNX17, SEC16A, MED16, CTDSP2, ARFRP1, RNF41, TAB1, TM9SF1, 44083, EHMT2, KDELR1, LMAN2, TMEM115, WDR6, COPZ1, RALY, TCF25, GANAB, SMG5, SIRT3, CBX6, PES1, POFUT1, DNPEP, CBY1, TMEM184B, ZNF500, IRF2BP1, ERAL1, CHMP2A, SGSM3, REM1, C19orf53, MYLPF, GMPPA, ZNF771, CUTA, ERGIC3, GPR173, DDX49, TMEM161A, ATP5SL, PACS1, PCDHGB5, PRDM11, PKNOX2, TSEN34, HPS6, OPA3, UBXN6, TOM1L2 |
| Pericytes_FANTOM_1___427 | Aran_et_al_2017 | FLT1, GDNF, HTR2B, P2RX1, POU2F2, MASP1, KSR1, S1PR2, HAND2, ADAMTSL2, COLEC10, BTN2A1, KIF26B, ADAMTS12 |

| | | |
|---|---|---|
| Pericytes_FANTOM_2___428 | Aran_et_al_2017 | COL10A1, SLC31A1, FLT1, GDNF, HTR2B, MASP1, HAND2, ADAMTSL2, DSCR4, COLEC10, IGF2BP3, ABTB2, ADAMTS12 |
| Pericytes_FANTOM_3___429 | Aran_et_al_2017 | FLT1, GDNF, HTR2B, MASP1, COLEC10, BTN2A1, KIF26B, ADAMTS12 |
| Plasma_cells_BLUEPRINT_1___430 | Aran_et_al_2017 | AMPD1, TNFRSF17, FKBP2, PNOC, SSR4, RNGTT, SPATS2, UBA5, ZBP1 |
| Plasma_cells_BLUEPRINT_2___431 | Aran_et_al_2017 | ALPI, AMPD1, AVP, TNFRSF17, BMP8B, C21orf2, CAMP, CASP10, CCNC, CD27, ENTPD1, CD79A, CDH15, COX6A2, CRYBB1, CRYBB3, CRYGC, CYP11A1, EPO, FKBP2, GDF2, GOLGA3, GOLGA4, GRM4, GUCA1A, HIST1H2BB, HSF4, HSPA6, IDE, IRF4, KCNN3, MARS, MGAT2, NEUROG1, NOS2, NPAS1, NPPA, NPPC, PNOC, PRM1, PTPRS, RAB3A, RAD17, RGS1, RGS13, RPN2, CCL25, SLC5A2, SRP54, SSR4, SS18, SYT5, TG, SEC62, TNNT3, HSP90B1, UBE2G1, ZNF133, RNF113A, RNF103, MANF, ITGA8, B4GALT3, MBTPS1, RNGTT, MATN4, EBAG9, HAND2, TBX4, CLINT1, KNTC1, KIAA0125, DMTF1, PREB, SLC35B1, MRPS31, CNKSR1, SEMA6C, PRDX4, LBX1, AVIL, SEC24A, TMED10, ZPBP, WDR45, TREH, FNDC3A, ATF6, MAST1, MAPK8IP3, UBXN4, SCFD1, CUX2, AIPL1, GORASP2, TBL2, DKKL1, NDOR1, SERP1, ARHGEF16, MCTS1, ALG5, GMPPA, SHANK1, CELA2B, YIPF1, LAX1, CCDC40, TMEM39A, UFSP2, CCDC88A, LTB4R2, PRX, SRPRB, PRDM14, MAGEF1, CHST8, PDIA2, SPATS2, MIS12, CCDC121, UBA5, ADM2, CNTD2, CCDC33, TRABD, ZBP1, STMN4, SLCO5A1, GRWD1, USP48, MTDH, IMP4, SIX5 |
| Plasma_cells_BLUEPRINT_3___432 | Aran_et_al_2017 | AMPD1, APOA1, APOC3, ARL1, PHOX2A, ARSA, AUP1, TNFRSF17, BMP8B, CA7, CACNA1S, CASP10, CAV1, CCNC, CD19, CD79A, CD79B, CHRNA4, CHRNG, CLCNKB, CRYBA4, CRYBB1, CSHL1, CYBA, CYP11A1, DAD1, DDOST, DNASE1L2, DPAGT1, DRD4, DRD5, FGF6, FKBP2, GH2, GNB3, GOLGB1, GP9, GRIN1, HDLBP, DNAJC4, IGF1, KCNN3, KRT10, CD180, MYL2, NTRK1, P2RY4, PDE6A, PNOC, POMC, POU3F3, PPIB, PTGER1, RAD17, RFX2, RGS13, RPN1, RPN2, SHBG, SLC6A13, SMPD2, SNAPC4, SSR1, SSR4, SURF1, SYT5, T, TERT, LEFTY2, SEC62, TNNT3, TP73, HSP90B1, TSHR, VPREB1, ZNF37A, ZNF133, ZNF142, MANF, ELL, BFSP2, UTF1, ITGA8, B4GALT3, MYH13, PABPC4, FBP2, SLC13A2, KCNQ4, GPR37L1, PICK1, TMEM59, CUL7, KIAA0125, PREB, PDIA6, SLC35B1, TIMM17B, CNKSR1, CNPY2, TM9SF1, PRDX4, LBX1, SEC24A, FTCD, SEC61B, LMAN2, TMED10, KDELR2, WDR45, SEC63, FNDC3A, RALY, MAST1, SIPA1L3, DDN, ABCB9, ISCU, SEC61G, TNFRSF13B, TSSK2, PPIL2, GORASP2, HEYL, TBL2, NDOR1, SERP1, ARHGEF16, SPCS1, SAP30BP, VPREB3, ALG5, GMPPA, SEC61A1, RAX, VSX1, SLC35C2, THAP4, IFT52, ERGIC3, YIPF1, TMED9, DEF8, RASIP1, ZDHHC4, C19orf73, UFSP2, GPRC5D, NGLY1, GLT8D1, PCDHA5, IRGC, LTB4R2, UGGT1, SRPRB, MYL7, MAGEF1, ACBD3, KLC2, SPATS2, YIPF2, FCRL2, OGFOD2, LMAN1L, ALG9, CSPP1, UBA5, DOK3, CNTD2, C6orf25, CEACAM21, MTDH, SLC38A10, SIX5, R3HCC1, MIA3, RNF208 |
| Plasma_cells_HPCA_1___433 | Aran_et_al_2017 | TNFRSF17, CD79A, GABRR2, GPLD1, KCNN3, NPAS1, PNOC, RGS13, CCL25, SLC5A2, SSR4, TP73, TSHR, PREB, CNKSR1, KLF15, VPREB3, TMEM39A, FN3K, UBA5, ADM2, ZBP1 |
| Plasma_cells_HPCA_2___434 | Aran_et_al_2017 | TNFRSF17, BMP8B, C21orf2, CD27, CD79A, CNR1, FKBP2, GABRR2, GPLD1, HSF4, KCNN3, NPAS1, PNOC, RGS13, CCL25, SLC5A2, SSR4, TCF3, TP73, HSP90B1, TSHR, WNT1, PREB, CNKSR1, SERP1, KLF15, VPREB3, HOOK2, SEC61A1, TMEM39A, FN3K, C16orf58, SPATS2, ACBD4, UBA5, ADM2, ZBP1, SIX5 |
| Plasma_cells_HPCA_3___435 | Aran_et_al_2017 | TNFRSF17, CD79A, GABRR2, GPLD1, KCNN3, NPAS1, PNOC, RGS13, CCL25, SLC5A2, SSR4, TP73, CNKSR1, KLF15, VPREB3, ADM2, ZBP1 |
| Plasma_cells_IRIS_1___436 | Aran_et_al_2017 | AMPD1, TNFRSF17, KCNN3, PNOC, RGS13, SSR4, RNGTT, SEC24A, ZBP1 |
| Plasma_cells_IRIS_2___437 | Aran_et_al_2017 | AMPD1, TNFRSF17, CCNC, CD79A, FKBP2, GOLGA3, HSF4, KCNN3, MGAT2, PNOC, RGS13, SRP54, SSR4, HSP90B1, UBE2G1, RNF113A, MANF, RNGTT, EBAG9, PREB, MRPS31, SEC24A, FNDC3A, ATF6, MAST1, UBXN4, TBL2, SERP1, ALG5, GMPPA, YIPF1, LAX1, TMEM39A, LTB4R2, SPATS2, UBA5, ADM2, ZBP1, USP48, MTDH, IMP4 |
| Plasma_cells_IRIS_3___438 | Aran_et_al_2017 | AMPD1, TNFRSF17, CCNC, CD79A, FKBP2, KCNN3, PNOC, RGS13, SSR4, TSHR, UBE2G1, RNGTT, PREB, SEC24A, TMEM39A, SPATS2, UBA5, ZBP1, MTDH |
| Platelets_HPCA_1___439 | Aran_et_al_2017 | ADCY8, ALOX12, APOA1, ARHGAP6, AVPR1A, BMP8B, BNIP2, CASQ1, CETP, COL10A1, GNAS, GNB3, GP1BA, GP5, GPR3, GPR17, LPAR4, HPCA, HSD17B3, IL5, ITGA2B, KIF2A, LCN1, LIMS1, SMAD2, MAN2A2, MAX, MEA1, RAB8A, CIITA, MPL, MUC6, NPPA, SIX6, PDE6A, PDE6H, PF4V1, PHKB, PITX3, PLXNB3, PPM1A, PPY, PRKCG, PTPRA, RNF2, RPA1, SELP, SLC6A4, NEK4, TACR2, TAL1, TG, TNNC2, CRISP2, VCL, VDAC3, XPNPEP1, ZNF37A, ZNF214, ST7, NCOA4, CDK2AP1, HIST1H2BO, SNX3, MYOM1, KSR1, ASAP2, ENDOU, RNF8, CHD1L, RGS6, SEC14L5, SNPH, TLK1, TECPR2, RNF10, TRIM10, HTATIP2, AKAP3, LEFTY1, IGF2BP3, CD160, PARK7, |

| | | |
|---|---|---|
| | | ACSBG1, ARHGEF12, TMEM50A, FBXO9, RNF11, DNAI1, MLH3, EIF2AK1, MORC1, LSM1, RABGEF1, RAX, KLK14, DERA, PHF20L1, RDH11, GP6, CLEC1B, TAOK3, BIN2, NCKIPSD, UIMC1, CMPK1, TUBA8, RNF186, L1TD1, BEST2, TXNL4B, CCDC40, ADI1, WDR11, ACTR10, NXF3, PCDHGB5, PCDHGA1, CABP5, ANO2, LRTM1, NEUROD4, PCTP, CHST8, ARMC7, SYNPO2L, PANK2, RUFY1, ULBP1, TUBB1, ASB8, PTCRA |
| Platelets_HPCA_2___440 | Aran_et_al_2017 | ADCY8, ALOX12, AMD1, ANXA7, ARHGAP6, AVPR1A, BMP8B, BNIP2, CASQ1, CETN2, COL10A1, DPYS, EGR4, F13A1, GNAS, GNB3, GP1BA, GP5, GP9, GPR3, GPR17, LPAR4, GRIK1, HPCA, HSD17B3, IL5, ITGA2B, KIF2A, LCN1, LIMS1, SMAD2, MAN2A2, MAX, MEA1, RAB8A, FOXO4, MPL, MTR, NOS2, NPPA, ODC1, SIX6, PDE6A, PDE6H, PF4V1, PHKB, PITX3, PPM1A, PPY, PRKCG, PTGDR, PTPRA, RHCE, RIT2, RNF2, RPA1, SELP, SLC6A4, SLC18A2, NEK4, TACR2, TAL1, TG, TNNC2, CRISP2, VCL, VDAC3, XPNPEP1, ZNF37A, ZNF214, ST7, NCOA4, CDK2AP1, HIST1H2BO, NCK2, TCAP, SNX3, MYOM1, PEX11B, KSR1, ASAP2, ENDOU, RNF8, ADIPOQ, CHD1L, RGS6, ZNF592, SEC14L5, SNPH, TLK1, TECPR2, NR2E3, TRIM10, HTATIP2, AKAP3, MYL12A, LEFTY1, IGF2BP3, PGRMC1, CD160, PDCD10, PARK7, SNW1, ACSBG1, ARHGEF12, SLC16A8, TMEM50A, FAM32A, IPCEF1, LSM14A, FBXO9, RNF11, DNAI1, NPTN, MLH3, EIF2AK1, MORC1, LSM1, RABGEF1, RAX, KLK14, DERA, RDH11, GP6, CLEC1B, CEND1, TAOK3, BIN2, NCKIPSD, UIMC1, CMPK1, TUBA8, RNF186, L1TD1, BEST2, TXNL4B, CCDC40, ADI1, C7orf43, WDR11, ACTR10, NXF3, PCDHGB5, PCDHGA1, CABP5, KCMF1, ANO2, LRTM1, PRX, PCTP, NPFFR1, NPVF, ADIPOR2, ARMC7, ZMYM1, SYNPO2L, PANK2, RUFY1, ULBP1, TUBB1, TRAPPC9, ASB8, PTCRA, USP12, 43898 |
| Platelets_HPCA_3___441 | Aran_et_al_2017 | ALOX12, ARHGAP6, GP1BA, PDE6H, PF4V1, SELP, SEC14L5, ACSBG1, MLH3, CLEC1B, CABP5, TUBB1 |
| Preadipocytes_ENCODE_1___442 | Aran_et_al_2017 | ARCN1, ARL1, CDK7, FGF7, GLG1, HAS1, HDLBP, SSR1, TESK1, TMEM11, BUB3, HAND2, BAG3, MORF4L2, TUBA1B, NCKAP1, KDELR2, SEC63, FAM98A, TCTN3, MMADHC, GLT8D1, SRPRB, ALG8, SLC25A32 |
| Preadipocytes_ENCODE_2___443 | Aran_et_al_2017 | FGF7, HAS1, SSR1, TMEM11, BUB3, HAND2, BAG3, MORF4L2, NCKAP1, KDELR2, GLT8D1, ALG8, SLC25A32 |
| Preadipocytes_ENCODE_3___444 | Aran_et_al_2017 | ADH5, ALCAM, SLC25A6, ANXA1, ANXA5, ARCN1, RHOA, ARL1, ARSB, ATP5C1, ATP5G1, CANX, CAST, CDK7, COPA, COPB1, ATF6B, DPAGT1, TOR1A, ECT2, ELK1, EPRS, FGF7, GLG1, GRSF1, HAS1, HDLBP, HIF1A, HSPA4, IARS, ISLR, MOCS2, NACA, NDUFB4, OCRL, P4HB, PPIB, PTGIR, PTPN11, PURA, RARS, RNF2, RPN2, RRM1, SNAPC2, SSR1, TESK1, TNXB, HSP90B1, PTP4A2, GNPAT, EIF3H, MBTPS1, TMEM11, WISP1, EIF2S2, MTMR2, WASL, MPZL1, BUB3, COPB2, KIF3B, HAND2, FXR2, BAG3, BAG2, MINPP1, MORF4L2, LAPTM4A, RANBP9, RAD50, CNPY2, TUBA1B, YAP1, TMEM147, SYNCRIP, APPBP2, MYL12A, YKT6, DCTN6, NCKAP1, PGRMC1, YIF1A, COPS8, HNRNPA0, SEC61B, ASCC3, METAP2, KDELR2, DSTN, PWP1, MRPL3, SEC63, XPOT, SCRG1, DOLK, ICMT, MKRN2, LMOD1, FAM98A, TMEM87A, DHRS7B, TCTN3, MYOF, NPTN, MMADHC, SEC61A1, COQ6, TMED7, FCF1, COPS4, DDX47, MBTPS2, UCHL5, ATP6V1D, RWDD1, UFC1, ERGIC3, C14orf166, MRPL20, IMPACT, RIOK2, GLT8D1, IFT46, SRPRB, EDA2R, CCDC90B, RIC8A, RNF25, EBF2, ACBD3, C7orf25, C7orf26, ALG8, TBC1D17, METTL8, PODNL1, ADM2, SLC25A32, ADAMTS12, MAGT1, ADO, SLC38A10, LRRC15, PRRC1, SENP5, TIPRL, GTF2H5 |
| Preadipocytes_FANTOM_1___445 | Aran_et_al_2017 | ADCYAP1R1, AQP9, CYP19A1, HIST1H2BB, IRF4, IPO5, SLC1A2, TNXB, IPO7, ADAMTS8, ABCA6, FAM98A, SNED1, PIWIL2, EDA2R, EBF2, TRPM3, IL17RC, TBC1D16 |
| Preadipocytes_FANTOM_2___446 | Aran_et_al_2017 | ARHGAP6, ACE, HAS1, IRF4, TNXB, ADAMTS8, ABCA6, SNED1, EBF2 |
| Preadipocytes_FANTOM_3___447 | Aran_et_al_2017 | ADCYAP1R1, AQP9, ARHGAP6, CENPE, CYP19A1, ACE, GLUD1, HIST1H2BB, HAS1, IRF4, IPO5, LSP1, SLC1A2, TNXB, XPNPEP2, HIST1H2BM, S1PR2, IPO7, ADAMTS8, MRPS27, SMC5, MDN1, ABCA6, FAM98A, SNED1, TREM1, PIWIL2, EDA2R, EBF2, TRPM3, IL17RC, TBC1D16 |
| pro_Bcells_HPCA_1___448 | Aran_et_al_2017 | AFM, ALPL, CCR8, CRX, DCC, DNTT, GABRA6, GCK, GPR3, GPR4, GRIK3, GYS2, TLX2, KNG1, MEN1, MUSK, OMD, PARK2, POU3F1, RAG2, RPE65, SLC2A2, SLC12A1, SRY, VPREB1, CSRP3, HIST1H2BM, SOX14, CACNA1G, GREB1, CNKSR1, ARPP21, FRS3, LILRB1, CLDN14, PLA2G2D, OR7A5, AKAP8L, STRN4, GNL2, TNFRSF12A, LARP7, BTBD7, SPATA7, HAMP, C2orf49, CXorf36, LRRTM4, FIP1L1, SLC25A31, SCRT1, IMP4, HSPB6, ZNF81, ZNF674 |
| pro_Bcells_HPCA_2___449 | Aran_et_al_2017 | ADARB2, ADCY8, AGXT, ALOX15B, ANXA3, APOC3, ARG1, ART4, BMX, CA1, CCKAR, SIGLEC6, CETP, CLCN1, CNTFR, CRH, CSHL1, CTSG, CYP2A7, DCC, DLX4, DNTT, DPYS, DSP, EFNA2, FCAR, FCN2, FGF8, MSTN, GDF10, GPR3, GPR4, GRIK3, GYS2, HCRTR2, ONECUT1, PRMT1, HTR1B, HTR5A, IFNA1, IGLL1, IL12B, KCNJ9, KCNJ13, |

| | | |
|---|---|---|
| | | KRT12, KRT19, LLGL1, LTC4S, MAG, MKI67, TRPM1, MUC6, MYH4, MYH8, NEUROG1, NOTCH4, OMG, PARK2, PMP2, POU1F1, PPEF2, PRB4, PSG11, RAD23A, RAG2, RAPSN, RBP3, MRPL12, RRM2, CCL17, TACR3, TCF3, TESK1, TGM3, THPO, TNNT2, UCP3, VPREB1, ZNF155, CSRP3, HIST1H2BL, HIST1H2BM, SOX14, EDF1, ADAM21, KCNQ4, OTOF, KIF23, CNKSR1, NMUR1, PLK4, ARPP21, FRS3, LILRB1, STIP1, LILRB4, KERA, ADAMTS8, CLCA4, FSTL4, PMPCA, ATP1B4, CLDN14, PADI4, TSSK2, CA14, FBXO24, SLC13A4, OR7A5, AKAP8L, DKKL1, AHDC1, BMP10, VPREB3, ANAPC2, VSX1, CALY, HP1BP3, PDE11A, CLEC1A, GMIP, TNFRSF12A, SPTBN5, ZMYND10, LARP7, COQ3, CNGB3, ZNF407, IQCC, BTBD7, SPATA7, NXF3, PAPOLB, RPGRIP1, SLURP1, SPC25, HAMP, MYL7, KLHL12, FBRS, RNF25, TUT1, MRPS15, KRI1, NOL12, CXorf36, KRTAP1-3, PCDH11Y, IMP4, HSPB6, PPP4R2, KCNV2, PSORS1C2, R3HCC1, ASPM, NACA2 |
| pro_Bcells_HPCA_3___450 | Aran_et_al_2017 | ARG1, CLCN1, CNTFR, CSHL1, DCC, DNTT, FCAR, FGF8, GPR3, HTR1B, IGLL1, LTC4S, MEP1B, TRPM1, MUC6, PRB4, PSG11, RAG2, MRPL12, SGCA, TACR3, TCOF1, TESK1, TGM3, VPREB1, ZNF155, HIST1H2BL, SOX14, ADAM21, KCNQ4, ARPP21, FRS3, LILRB1, ADAMTS8, LAMB4, CLDN14, FBXO24, AKAP8L, AHDC1, BMP10, VSX1, PCDHA5, OTUD7B, SPC25, HAMP, MYL7, DPEP3, C2orf49, CXorf36, IMP4, HSPB6 |
| pro_Bcells_NOVERSHTERN_1___451 | Aran_et_al_2017 | AZU1, BLK, CD72, CD79B, CENPA, CETP, DNTT, FOXM1, FLT3, H2AFX, IGLL1, KIF11, LY6H, MKI67, MYBL2, PDE6D, POLA1, PRTN3, RAG2, RFC2, RFC5, RRM2, SMARCA4, SNRPD1, SPTA1, TCF3, TERT, TOP2B, TSSC1, VPREB1, XPNPEP2, PTTG1, TCL1B, ESPL1, KIF14, P2RY14, SMC4, TACC3, NOP56, SIVA1, ARPP21, HNRNPA0, UBE2C, KIF4A, OR7A5, VPREB3, TRA2A, SAC3D1, MRTO4, NUSAP1, AHSP, GTSE1, CEP55, QRSL1, SPC25, LSM2, CCDC81, SHCBP1, C16orf59, HPS4 |
| pro_Bcells_NOVERSHTERN_2___452 | Aran_et_al_2017 | BLK, CD72, CD79B, DNTT, FLT3, IGLL1, PRTN3, RAG2, VPREB1, ARPP21, VPREB3, QRSL1, CCDC81 |
| pro_Bcells_NOVERSHTERN_3___453 | Aran_et_al_2017 | BLK, CD72, CD79B, DNTT, FLT3, IGLL1, LY6H, MYBL2, PRTN3, RAG2, VPREB1, ARPP21, VPREB3, QRSL1, CCDC81 |
| Sebocytes_FANTOM_1___454 | Aran_et_al_2017 | CALML3, CSF2, CSTA, CTSK, DSG3, GJB5, SFN, IL1A, IRF6, KRT6B, LAD1, MMP3, PI3, PKP1, SOX15, SULT2B1, KRT75, FGFBP1, AP1M2, PDZK1IP1, FST, IL24, LPAR3, TFCP2L1, HES2, TMEM40, LTB4R2, RAB25, S100A14, C1orf116, ZNF750, KRT6C |
| Sebocytes_FANTOM_2___455 | Aran_et_al_2017 | CALML3, CSF2, CSTA, CTSK, DSG3, GJB5, SFN, IL1A, KRT6B, MMP3, PI3, PKP1, SOX15, SULT2B1, KRT75, FGFBP1, AP1M2, PDZK1IP1, HES2, TMEM40, RAB25, S100A14, ZNF750, KRT6C |
| Sebocytes_FANTOM_3___456 | Aran_et_al_2017 | CALML3, ENTPD3, CSF2, CSTA, CTSK, DSG3, GJB3, GJB5, SFN, IL1A, IRF6, KRT6B, LAD1, MMP3, PI3, PKP1, PTK6, SOX15, SULT2B1, KRT75, FGFBP1, AP1M2, PDZK1IP1, FST, IL24, LPAR3, TFCP2L1, HES2, TMEM40, LTB4R2, RAB25, S100A14, C1orf116, ZNF750, KRT6C |
| Skeletal_muscle_ENCODE_1___457 | Aran_et_al_2017 | CAV3, CDH15, CHRNG, MYBPH, MYF5, RAPSN, MYLPF, EBF2 |
| Skeletal_muscle_ENCODE_2___458 | Aran_et_al_2017 | CAV3, CDH15, CHRNG, MYBPH, MYF5, MYOD1, RAPSN, MYLPF, LSM2, EBF2 |
| Skeletal_muscle_ENCODE_3___459 | Aran_et_al_2017 | CDK1, CENPE, CHRNG, MYBPH, MYOG, RAPSN, MYLPF, EBF2 |
| Skeletal_muscle_FANTOM_1___460 | Aran_et_al_2017 | ACHE, ACTA1, ACTN2, AMPD1, ART1, ART3, ATP1A2, ATP2A1, CA3, CACNA1S, CACNB1, CACNG1, CASQ1, CDH15, CHRND, CHRNG, CKM, CNTFR, COX6A2, DES, RAPGEF1, HADHB, HRC, KCNN3, MUSK, MYBPC1, MYBPC2, MYBPH, MYF6, MYH1, MYH2, MYH6, MYH7, MYL1, MYL2, MYL3, MYOD1, MYOG, NRAP, PCNT, PGAM2, PGM1, PHKG1, PPP1R3A, PYGM, RAD23A, RAPSN, RPL3L, CLIP1, RYR3, MAPK12, SGCA, SGCG, SLC2A4, SLN, SPTB, TNNC2, TNNI1, TNNT3, TTN, UCP3, CSRP3, TCAP, MYOM1, FBP2, MYOT, BAG3, ABCC9, TRDN, UBAC1, SEMA6C, LBX1, APOBEC2, LDB3, SIRT2, IQSEC2, DDN, MAST2, KIAA0368, HSPB7, CTNNA3, MYLPF, UBE2D4, ASB4, FBXO40, MYOZ2, MIOS, CASZ1, MYOZ1, POPDC2, LONRF3, SYNPO2L, OBSCN, OSBPL11 |
| Skeletal_muscle_FANTOM_2___461 | Aran_et_al_2017 | ACTA1, ACTN2, AMPD1, ART1, ART3, ATP1A2, ATP2A1, CA3, CACNA1S, CACNB1, CACNG1, CASQ1, CDH15, CHRND, CHRNG, CKM, COX6A2, DES, HRC, KCNN3, MYBPC1, MYBPC2, MYBPH, MYF6, MYH1, MYH2, MYH6, MYH7, MYL1, MYL2, MYL3, MYOG, NRAP, PGAM2, PHKG1, PPP1R3A, PYGM, RAPSN, RPL3L, SGCA, SGCG, SLC2A4, SLN, SPTB, TNNC2, TNNI1, TNNT3, TTN, UCP3, CSRP3, TCAP, MYOM1, FBP2, MYOT, TRDN, SEMA6C, APOBEC2, LDB3, IQSEC2, DDN, CTNNA3, MYLPF, FBXO40, MYOZ2, MYOZ1, POPDC2, SYNPO2L, OBSCN, OSBPL11 |
| Skeletal_muscle_FANTOM_3___462 | Aran_et_al_2017 | ACTA1, ACTN2, AMPD1, ART1, ART3, ATP1A2, ATP2A1, CA3, CACNA1S, CACNB1, CACNG1, CASQ1, CDH15, CHRND, CHRNG, CKM, COX6A2, DES, HADHB, HRC, |

| | | |
|---|---|---|
| | | KCNN3, MUSK, MYBPC1, MYBPC2, MYBPH, MYF6, MYH1, MYH2, MYH6, MYH7, MYL1, MYL2, MYL3, MYOG, NRAP, PCNT, PGAM2, PHKG1, PPP1R3A, PYGM, RAPSN, RPL3L, CLIP1, SGCA, SGCG, SLC2A4, SLN, SPTB, TNNC2, TNNI1, TNNT3, TTN, UCP3, CSRP3, TCAP, MYOM1, FBP2, MYOT, TRDN, SEMA6C, LBX1, APOBEC2, LDB3, IQSEC2, DDN, HSPB7, CTNNA3, MYLPF, ASB4, FBXO40, MYOZ2, CASZ1, MYOZ1, POPDC2, SYNPO2L, OBSCN, OSBPL11 |
| Smooth_muscle_ENCODE_1___463 | Aran_et_al_2017 | COPA, COPB1, FKTN, HDLBP, PFN2, PRKAG1, PRKG1, USO1, ASAP2, BAG2, PREPL, VTI1B, HSPB6 |
| Smooth_muscle_ENCODE_2___464 | Aran_et_al_2017 | ARL1, COPA, COPB1, DCTN1, DYNC1LI2, FKTN, HDLBP, HOXA3, LGALS1, PFN2, PRKAG1, PRKG1, TCF21, GDF5, USO1, MBTPS1, ASAP2, COPS2, VTI1B, ANAPC13, MYOF, AMZ2, KLHL9, ACBD3, DYNLRB1, HSPB6 |
| Smooth_muscle_ENCODE_3___465 | Aran_et_al_2017 | ADH1B, ALCAM, ARCN1, ARL1, COPA, COPB1, DCTN1, DYNC1LI2, EIF4G2, FKTN, HDLBP, HOXA3, HPD, ISLR, IPO5, LGALS1, NBR1, OCRL, OGN, PFN2, PRKAG1, PRKG1, SDF2, TCF21, GDF5, USO1, MBTPS1, ASAP2, COPS2, KIF3B, SCAMP1, BAG2, PREPL, TTC37, CRYZL1, STAM2, VTI1B, ERLIN1, FAF2, EID1, ANAPC13, MYOF, INVS, COPS7A, DCTN4, AMZ2, UFSP2, KLHL9, THAP10, IFT46, EDA2R, PKNOX2, ACBD3, MRPL40, DYNLRB1, HSPB6 |
| Smooth_muscle_FANTOM_1___466 | Aran_et_al_2017 | ADD1, ADH1B, ADH5, ALCAM, ANXA5, ARCN1, ARF4, ARHGAP6, ARL1, BAD, CACNA1C, CAPNS1, CETN2, COPA, COPB1, CSNK1G3, CSNK2A2, CTNNA1, DCTN1, DDB1, DYNC1LI2, DPT, EIF4G2, ELN, FKTN, FGF7, FSHB, GDF10, GOLGA3, GRSF1, HDLBP, HLCS, HOXA3, HPD, HSP90AB1, ISLR, IPO5, KRT19, KTN1, LGALS1, LTC4S, NBR1, SMAD5, MEA1, 44076, NFATC4, OCRL, OGN, PCDHGC3, PEX12, PFN2, PHKG1, PPP2R1A, PRKAG1, PRKG1, PSMB7, PTPN11, RARS, RBMS1, RFX2, RING1, S100A6, S100A10, ATXN2, SDF2, SGCD, SNTB2, SSR1, MAP3K7, TCF21, CLEC3B, VCL, VIM, GDF5, USO1, MBTPS1, ASAP2, ZMYM4, COPS2, VAMP3, TXNL1, KIF3B, SCAMP1, BAG2, SPAG7, PREPL, TTC37, TOMM20, CUL7, CRYZL1, SNUPN, ABI2, STAM2, ZMPSTE24, SPEG, TFG, YAP1, VTI1B, ERLIN1, IGF2BP3, FAM189B, SPIN1, KDELR1, ASCC3, KDELR2, RER1, EMILIN1, SCRG1, EPN2, TRIM32, WDFY3, FBXO21, ERC1, SNX13, GANAB, SCFD1, EXOC7, DNAJC13, EID1, TMEM59L, CIZ1, LMOD1, TMEM184B, ANAPC13, FAM98A, GORASP2, MYOF, TBL2, MLH3, DKKL1, HSPB7, INVS, TUBG2, METTL5, TNPO2, COPS7A, CCDC53, DCTN4, AMZ2, NGRN, MBTPS2, ANKFY1, ERGIC3, MYOZ2, DHX29, TMED9, ASPN, FBXL12, DALRD3, UFSP2, EXOC1, SPATA7, GLT8D1, TMEM165, KLHL9, THAP10, IFT46, ZNF471, SRPRB, EDA2R, CCDC90B, MRPL17, PKNOX2, ACBD3, MRPL40, SPATS2, C11orf95, YIPF2, CUEDC2, DDA1, TSEN34, FTO, C2orf49, ZNF426, SPAG16, TBC1D17, CCDC102B, SLC35E1, SVEP1, NETO2, DYNLRB1, TM2D1, COG7, TEX261, HSPB6, PRRC1, MYL6B, ZNF358, SIX5, DPY19L4, C6orf120, GTF2H5 |
| Smooth_muscle_FANTOM_2___467 | Aran_et_al_2017 | ARL1, COPA, COPB1, DCTN1, DYNC1LI2, FKTN, HDLBP, HOXA3, LGALS1, PFN2, PRKAG1, PRKG1, TCF21, GDF5, USO1, MBTPS1, ASAP2, COPS2, VTI1B, ANAPC13, MYOF, AMZ2, KLHL9, ACBD3, DYNLRB1, HSPB6 |
| Smooth_muscle_FANTOM_3___468 | Aran_et_al_2017 | ADD1, ALCAM, ARF4, COPA, DCTN1, DPT, FKTN, FGF7, GDF10, HDLBP, HOXA3, HPD, ISLR, LGALS1, NBR1, SMAD5, NFATC4, OCRL, OGN, PCDHGC3, PEX12, PFN2, PRKG1, S100A6, S100A10, ATXN2, SSR1, TCF21, CLEC3B, GDF5, ZMYM4, COPS2, TXNL1, SCAMP1, BAG2, TTC37, CUL7, ABI2, SPIN1, KDELR1, EMILIN1, EXOC7, EID1, TMEM59L, CIZ1, FAM98A, HSPB7, AMZ2, TMED9, ASPN, SPATA7, KLHL9, THAP10, ZNF471, EDA2R, CCDC90B, ACBD3, C11orf95, FTO, SPAG16, CCDC102B, SVEP1, DYNLRB1, HSPB6, MYL6B, ZNF358 |
| Smooth_muscle_HPCA_1___469 | Aran_et_al_2017 | CDK4, LGALS1, PLS3, SOD1, TMED7, DDX47, UFC1, CISD1 |
| Smooth_muscle_HPCA_2___470 | Aran_et_al_2017 | CCNG1, CETN2, NDUFS4, PLS3, RPL10, RPS19, S1PR2, RNF7, MCTS1, DDX47, POMP, TMED9, ADAMTS12, GTF2H5 |
| Smooth_muscle_HPCA_3___471 | Aran_et_al_2017 | ACTG1, ADH5, ALDOA, ANXA1, ANXA5, RHOC, ATP5G1, ATP5J, BYSL, PTTG1IP, CALR, CANX, CAPN2, CCNG1, CDK4, CDK7, CETN2, CNN2, COPA, COPB1, COX8A, ECT2, EEF1D, ERCC1, FKTN, GOLGA3, HADHA, HDLBP, HIC1, HSPA8, EIF6, LAMP1, LGALS1, MIF, MYL6, NDUFA8, NDUFB4, NDUFS4, NDUFS5, 44076, NFATC4, P4HB, PLS3, POLR2F, PSMB1, PSMB4, PSMB5, PSMC5, PSMD10, PTGIR, PTPN11, PEX2, RARS, RPL4, RPL10, RPL35A, RPN2, RPS19, CCL8, SOD1, SSBP1, ST13, TPI1, TPM3, HSP90B1, UFD1L, VCP, MANF, SHFM1, PTP4A2, EIF3I, MBTPS1, MTMR2, ATP6V0E1, MPZL1, BUB3, COPB2, S1PR2, KIF3B, TMEM59, BAG3, C14orf2, MINPP1, RNF7, MORF4L2, LAPTM4A, RWDD2B, PSMD14, CDIPT, TIMM17A, ZNHIT1, AHSA1, NCKAP1, PGRMC1, SEC61B, TMED1, EMILIN1, POFUT2, ISCU, SEC61G, STX12, LMOD1, ATXN10, FAM98A, TMEM87A, FBXO22, UQCRQ, MMADHC, MCTS1, MRPL15, CNIH4, TMED7, DDX47, GMPR2, ZNF771, POMP, ATP6V1D, TRAPPC4, UFC1, |

| | | |
|---|---|---|
| | | CUTA, ERGIC3, MRPS33, RIN2, TMED9, C19orf24, NOP10, CISD1, PCDHGB5, SRPRB, C12orf10, EDA2R, FAM160B2, MRPS11, SPATS2, CUEDC2, TBC1D17, MYCT1, ADAMTS12, DYNLRB1, TM2D1, LRRC15, PRRC1, SENP5, GTF2H5 |
| Tgd_cells_HPCA_1___472 | Aran_et_al_2017 | AK2, ABCD2, FASLG, BUB1, CCNF, CD2, CD247, CD40LG, CDC25C, CENPA, CHEK1, CLIC1, CCR3, CCR5, DAXX, DR1, ECT2, GLE1, GLO1, GPI, CXCR3, GPR15, GYG1, GZMH, GZMA, GZMB, GZMK, H2AFX, HMOX2, IL2RA, IL2RB, IL4, IL5, IL12RB1, IL13, ITGAL, ITGB7, KIF2A, KIF11, KIF22, LAG3, LAIR2, LCP2, LIM2, LTA, MKI67, MSH3, NEK2, NKG7, PDE4A, PDE6D, SLC26A4, PFN1, PPID, PPP1CA, PRF1, PSMA3, PSMB2, PSMC4, PSMD13, PTPN4, PTPN7, PTPN9, PEX2, RBL1, RPA1, RRM1, RRM2, SOS1, AURKA, TMPO, ZBTB16, ZNF174, PTP4A2, GPR68, CLPP, DGCR14, RANBP3, HAT1, RGS9, IL18RAP, TOP3B, TAF1B, PSTPIP1, CD101, GRAP2, ATG5, DLGAP5, KIF14, ARPC2, RAD50, KLRG1, CD96, SF3B4, HMGN4, GNLY, SMC2, CXCR6, GMEB1, PLK4, POP4, DBF4, STIP1, KIF2C, HNRNPUL1, GLMN, CD300A, RALY, TPX2, NCDN, FAM120A, NCAPD3, RNF167, TOR1AIP1, TINF2, ZBTB32, GPKOW, CHMP4A, RACGAP1, SAC3D1, GPR171, TBX21, NUSAP1, UCHL5, FAM96B, HSPB11, WBP11, CD244, ARMC1, CEP55, ASXL2, UEVLD, HJURP, CHST12, IL26, CENPJ, CENPN, ZMAT5, KLHL7, SPC25, IKZF4, MMP25, ACD, PVRIG, ZNF668, ATP8B4, VANGL1, ARPC5L, MTDH, ACTR8, SFXN1, NCR3, CCR2 |
| Tgd_cells_HPCA_2___473 | Aran_et_al_2017 | ABCD2, FASLG, BUB1, CCNA2, CCNF, CD2, CD247, CDK1, CDC5L, CDC25C, CENPA, CLIC1, CCR3, CCR5, COX8A, CSTF1, DAXX, DR1, DRG2, TOR1A, ECT2, GLE1, GLO1, GPI, CXCR3, GPR15, GYG1, GZMH, GZMA, GZMB, GZMK, H2AFX, HIC1, HMOX2, IFNG, IL2RA, IL2RB, IL4, IL5, IL12RB1, IL13, INPP4A, ITGAL, ITGB7, LAG3, LAIR2, LCP2, LIM2, LTA, MKI67, MNAT1, MSH3, NEK2, NFKBIB, NMT1, PDE4A, PDE6D, SLC26A4, PPID, PPP1CA, PRF1, PSMA1, PSMA3, PSMB2, PSMC4, PSMD13, PTPN4, PTPN7, PTPN9, PEX2, RBL1, RNF6, RRM1, RRM2, SOS1, SRF, AURKA, TMPO, TTK, ZBTB16, ZNF174, PTP4A2, GPR68, CLPP, DGCR14, RANBP3, RGS9, CDC123, TOP3B, TAF1B, PSTPIP1, CIAO1, CD101, GRAP2, ATG5, STX8, DLGAP5, ARPC2, PSMD14, KLRG1, CD96, DCAF7, SF3B4, GNLY, SMC2, CXCR6, GMEB1, PLK4, POP4, ARPP19, DBF4, STIP1, KIF2C, FAF1, GLMN, PUF60, TPX2, NCDN, FAM120A, RNF167, TOR1AIP1, ZBTB32, GPKOW, ZCCHC4, CHMP4A, GPR171, TBX21, ASCC1, DBR1, GMIP, UCHL5, PIAS4, FAM96B, HSPB11, WBP11, SASH3, COMMD8, ARMC1, CEP55, ASXL2, UEVLD, HJURP, TDP1, IL26, CENPN, KLHL7, SPC25, IKZF4, MMP25, MRPS15, ACD, PVRIG, ZNF668, ATP8B4, SLC25A32, VANGL1, ARPC5L, MTDH, ACTR8, SFXN1, NCR3, TIPRL, TMEM110, CCR2 |
| Tgd_cells_HPCA_3___474 | Aran_et_al_2017 | ABCD2, FASLG, BARD1, BUB1, CCNA2, CCNF, CD2, CD247, CD40LG, CDK1, CDC25C, CENPA, CLIC1, CCR3, CCR5, COX8A, CTLA4, DAXX, DR1, DRG2, ECT2, GLE1, GLO1, GPI, CXCR3, GPR15, GYG1, GZMH, GZMA, GZMB, GZMK, H2AFX, HIC1, HMOX2, HNRNPF, IFNG, IL2RA, IL2RB, IL4, IL5, IL12RB1, IL13, INPP4A, ITGAL, ITGB7, KIF2A, KIF22, LAG3, LAIR2, LCP2, LIM2, LTA, SH2D1A, MKI67, MNAT1, MSH3, NEK2, NFKBIB, NKG7, NMT1, PDE4A, PDE6D, SLC26A4, PPID, PPP1CA, PRF1, PSMA3, PSMB2, PSMC4, PSMD4, PSMD7, PSMD13, PTPN4, PTPN7, PTPN9, PEX2, RAD21, RB1, RBL1, RRM1, CCL1, SHMT2, SLAMF1, SOS1, AURKA, TMPO, USP1, ZBTB16, ZNF174, PTP4A2, GPR68, CLPP, DGCR14, COLQ, RANBP3, RGS9, CDC123, TOP3B, TAF1B, PSTPIP1, CIAO1, CD101, GRAP2, ATG5, STX8, DLGAP5, MELK, G3BP2, KIF14, DCLRE1A, SCAMP2, ARPC2, RAD50, PSMD14, KLRG1, CD96, SF3B4, TACC3, GNLY, SMC2, CXCR6, GMEB1, PLK4, POP4, ARPP19, DBF4, STIP1, KIF2C, GLMN, CD300A, TPX2, TAB2, NCDN, FAM120A, RNF167, TOR1AIP1, ZBTB32, GPKOW, ZCCHC4, CHMP4A, GPR171, TBX21, ASCC1, DBR1, GMIP, AMZ2, FZR1, UCHL5, FAM96B, HSPB11, WBP11, CD244, CSNK1G1, SASH3, RC3H2, COMMD8, AGGF1, CEP55, ASXL2, UEVLD, HJURP, IL26, CENPN, PBK, KLHL7, SPC25, HIVEP3, IKZF4, MMP25, ACD, PVRIG, ZNF668, ATP8B4, SLC25A32, VANGL1, MFSD5, MTDH, ACTR8, SFXN1, NCR3, TIPRL, TMEM110, CCR2 |
| Th1_cells_IRIS_1___475 | Aran_et_al_2017 | CHD4, CSTF1, IFNG, LAG3, MNAT1, POLD2, PPM1G, SLAMF1, SNRPC, THOP1, CDC123, EIF2B2, FIBP, CHD1L, MDC1, TRIM28, GNLY, RUVBL2, NCAPD3, R3HDM1, TACO1, TMEM39B, UBAP2, CUEDC2 |
| Th1_cells_IRIS_2___476 | Aran_et_al_2017 | IFNG, LAG3, SNRPC, GNLY, RUVBL2, NCAPD3, TACO1, TMEM39B, UBAP2, CUEDC2 |
| Th1_cells_IRIS_3___477 | Aran_et_al_2017 | COX10, IFNG, LAG3, PSMD3, SNRPC, EIF2B2, PKMYT1, PTTG1, KIF20A, RNPS1, TTLL5, NUP205, ZBTB32, HTRA2, WRAP53, WDR18, CUEDC2 |
| Th2_cells_IRIS_1___478 | Aran_et_al_2017 | GZMK, IL5, IL13, MAD2L1, RRM2, BAG2, CXCR6, CEP55 |
| Th2_cells_IRIS_2___479 | Aran_et_al_2017 | IL5, IL13, MAD2L1, BAG2, CXCR6, RRAS2, CEP55, NUP37, NPHP4 |

| | | |
|---|---|---|
| Th2_cells_IRIS_3___480 | Aran_et_al_2017 | GPR15, GZMA, IL5, IL13, SMAD2, CDK2AP1, RGS9, BAG2, SLC25A44, RAD50, CXCR6, TMEM39B, UBAP2, THADA, RNF34, NPHP4 |
| Tregs_BLUEPRINT_1___481 | Aran_et_al_2017 | CCR3, CTLA4, IL2RA, PLCL1, PPM1B, TTN, ZNF236, STAM, UBE4A, CXCR6, IPCEF1, ICOS, VPS54, LAX1, BANP, ATG2B, ZCCHC8, TULP4, IKZF4, ZMYM1, ZFC3H1, MCM9 |
| Tregs_BLUEPRINT_2___482 | Aran_et_al_2017 | CTLA4, IL2RA, PLCL1, ZNF236, STAM, IPCEF1, ICOS, BANP, IKZF4 |
| Tregs_BLUEPRINT_3___483 | Aran_et_al_2017 | CTLA4, IL2RA, PLCL1, PPM1B, ZNF236, STAM, IPCEF1, ICOS, VPS54, BANP, ATG2B, ZCCHC8, TULP4, IKZF4, ZFC3H1 |
| Tregs_FANTOM_1___484 | Aran_et_al_2017 | CTLA4, IL2RA, PLCL1, ZNF236, STAM, IPCEF1, ICOS, BANP, IKZF4 |
| Tregs_FANTOM_2___485 | Aran_et_al_2017 | CTLA4, IL2RA, PLCL1, PPM1B, ZNF236, STAM, IPCEF1, ICOS, VPS54, BANP, ATG2B, ZCCHC8, TULP4, IKZF4, ZFC3H1 |
| Tregs_FANTOM_3___486 | Aran_et_al_2017 | CD5, CD28, CCR4, CCR8, CTLA4, GPR25, IL2RA, IL10RA, ITGB7, KCNA2, PLCL1, RGS1, SPTAN1, HS3ST3B1, MCF2L2, GALNT8, SIT1, ICOS, FOXP3, LRP2BP, TULP4 |
| Tregs_HPCA_1___487 | Aran_et_al_2017 | CCR4, CCR8, CTLA4, GPR25, IL2RA, KCNA2, LAIR2, RGS1, HS3ST3B1, MCF2L2, ICOS, FOXP3 |
| Tregs_HPCA_2___488 | Aran_et_al_2017 | CCR4, CCR8, CTLA4, GPR25, IL2RA, KCNA2, LAIR2, HS3ST3B1, MCF2L2, FOXP3 |
| Tregs_HPCA_3___489 | Aran_et_al_2017 | CCR4, CCR8, CTLA4, GPR25, IL2RA, KCNA2, LAIR2, RGS1, HS3ST3B1, MCF2L2, FOXP3 |
| Bcells___490 | Bindea_et_al_2013 | ABCB4, BACH2, BCL11A, BLK, BLNK, CCR9, CD19, CD72, COCH, CR2, DTNB, FCRL2, GLDC, GNG7, HLA-DOB, HLA-DQA1, IGHA1, IGHG1, IGHM, IGKC, IGL, KIAA0125, MEF2C, MICAL3, MS4A1, OSBPL10, PNOC, QRSL1, SCN3A, SLC15A2, SPIB, TCL1A, TNFRSF17 |
| CD4_Tcm___491 | Bindea_et_al_2013 | AQP3, ATF7IP, ATM, CASP8, CDC14A, CEP68, CLUAP1, CREBZF, CYLD, DOCK9, FAM153B, FOXP1, FYB, HNRPH1, INPP4B, KLF12, LOC441155, MAP3K1, MLL, N4BP2L2-IT2, NEFL, NFATC3, PCM1, PCNX, PDXDC2, PHC3, POLR2J2, PSPC1, REPS1, RPP38, SLC7A6, SNRPN, ST3GAL1, STX16, TIMM8A, TRAF3IP3, TXK, TXLNGY, USP9Y |
| CD4_Tem___492 | Bindea_et_al_2013 | AKT3, C7orf54, CCR2, DDX17, EWSR1, FLI1, GDPD5, LTK, MEFV, NFATC4, PRKY, TBC1D5, TBCD, TRA, VIL2 |
| CD8_Tcells___493 | Bindea_et_al_2013 | ABT1, AES, APBA2, ARHGAP8, C12orf47, C19orf6, C4orf15, CAMLG, CD8A, CD8B, CDKN2AIP, DNAJB1, FLT3LG, GADD45A, GZMM, KLF9, LEPROTL1, LIME1, MYST3, PF4, PPP1R2, PRF1, PRR5, RBM3, SF1, SFRS7, SLC16A7, TBCC, THUMPD1, TMC6, TSC22D3, VAMP2, ZEB1, ZFP36L2, ZNF22, ZNF609, ZNF91 |
| DC___494 | Bindea_et_al_2013 | CCL13, CCL17, CCL22, CD209, HSD11B1, NPR1, PPFIBP2 |
| Eosinophils___495 | Bindea_et_al_2013 | ABHD2, ACACB, C9orf156, CAT, CCR3, CLC, CYSLTR2, EMR1, EPN2, GALC, GPR44, HES1, HIST1H1C, HRH4, IGSF2, IL5RA, KBTBD11, KCNH2, LRP5L, MYO15B, RCOR3, RNASE2, RRP12, SIAH1, SMPD3, SYNJ1, TGIF1, THBS1, THBS4, TIPARP, TKTL1 |
| Macrophages___496 | Bindea_et_al_2013 | APOE, ATG7, BCAT1, CCL7, CD163, CD68, CD84, CHI3L1, CHIT1, CLEC5A, COL8A2, COLEC12, CTSK, CXCL5, CYBB, DNASE2B, EMP1, FDX1, FN1, GM2A, GPC4, KAL1, MARCO, ME1, MS4A4A, MSR1, PCOLCE2, PTGDS, RAI14, SCARB2, SCG5, SGMS1, SULT1C2 |
| Mast_cells___497 | Bindea_et_al_2013 | ABCC4, ADCYAP1, CALB2, CEACAM8, CMA1, CPA3, CTSG, ELA2, GATA2, HDC, HPGD, HPGDS, KIT, LINC01140, MAOB, MLPH, MPO, MS4A2, NR0B1, PPM1H, PRG2, PTGS1, SCG2, SIGLEC6, SLC18A2, SLC24A3, TAL1, TPSAB1, TPSB2, VWA5A |
| Neutrophils___498 | Bindea_et_al_2013 | ADARB1, AF107846, ALDH1B1, APBB2, ATL2, BCL2, CDC5L, FGF18, FUT5, FZR1, GAGE2A, IGFBP5, KANK2, LDB3, MAPRE3, MCM3AP, MRC2, NCR1, PDLIM4, PRX, PSMD4, RP5-886K2.1, SGMS1, SLC30A5, SMEK1, SPN, TBXA2R, TCTN2, TINAGL1, TRPV6, XCL1, XCL2, ZNF205, ZNF528, ZNF747 |
| NK_cells___499 | Bindea_et_al_2013 | ALPL, BST1, CD93, CEACAM3, CREB5, CRISPLD2, CSF3R, CYP4F3, DYSF, FCAR, FCGR3B, FLJ11151, FPR1, FPRL1, G0S2, HIST1H2BC, HPSE, IL8RA, IL8RB, KCNJ15, LILRB2, MGAM, MME, PDE4B, S100A12, SIGLEC5, SLC22A4, SLC25A37, TECPR2, TNFRSF10C, VNN3 |
| Tgd_cells___500 | Bindea_et_al_2013 | C1orf61, CD160, FEZ1, TARP, TRD, TRGV9 |
| Th1_cells___501 | Bindea_et_al_2013 | APBB2, APOD, ATP9A, BST2, BTG3, CCL4, CD38, CD70, CMAH, CSF2, CTLA4, DGKI, DOK5, DPP4, DUSP5, EGFL6, GGT1, HBEGF, IFNG, IL12RB2, IL22, LRP8, LRRN3, |

| | | |
|---|---|---|
| | | LTA, SGCB, SYNGR3, ZBTB32 |
| Th2_cells___502 | Bindea_et_al_2013 | ADCY1, AHI1, ANK1, BIRC5, CDC25C, CDC7, CENPF, CXCR6, DHFR, EVI5, GATA3, GSTA4, HELLS, IL26, LAIR2, LIMA1, MB, MICAL2, NEIL3, PHEX, PMCH, PTGIS, SLC39A14, SMAD2, SNRPD1, WDHD1 |
| Tregs___503 | Bindea_et_al_2013 | FOXP3 |
| aDC___504 | Bindea_et_al_2013 | CCL1, EBI3, INDO, LAMP3, OAS3 |
| iDC___505 | Bindea_et_al_2013 | ABCG2, BLVRB, CARD9, CD1A, CD1B, CD1C, CD1E, CH25H, CLEC10A, CSF1R, CTNS, F13A1, FABP4, FZD2, GSTT1, GUCA1A, HS3ST2, LMAN2L, MMP12, MS4A6A, NUDT9, PDXK, PPARG, PREP, RAP1GAP, SLC26A6, SLC7A8, SYT17, TACSTD2, TM7SF4, VASH1 |
| pDC___506 | Bindea_et_al_2013 | IL3RA |
| Bcells___507 | Charoentong_et_al_2017 | CD180, CD79B, BLK, CD19, MS4A1, TNFRSF17, IGHM, GNG7, MICAL3, SPIB, HLA-DOB, IGKC, PNOC, FCRL2, BACH2, CR2, TCL1A, AKNA, ARHGAP25, CCL21, CD27, CD38, CLEC17A, CLEC9A, CLECL1 |
| CD4_Tcells___508 | Charoentong_et_al_2017 | AIM2, BIRC3, BRIP1, CCL20, CCL4, CCL5, CCNB1, CCR7, DUSP2, ESCO2, ETS1, EXO1, EXOC6, IARS, ITK, KIF11, KNTC1, NUF2, PRC1, PSAT1, RGS1, RTKN2, SAMSN1, SELL, TRAT1 |
| CD4_Tcm___509 | Charoentong_et_al_2017 | ABHD3, AHNAK, ANXA2P2, AQP3, ATHL1, BMI1, BZW2, CD63, COL4A1, CYLD, ELMO2, FYN, GLIPR1, GSS, IFITM2, ITGB1, ITGB2, KLF5, LSP1, NDUFB9, PKM2, SFXN3, SIRPG, SMAD4, STX4, TRADD, VIM, XRCC6 |
| CD4_Tem___510 | Charoentong_et_al_2017 | ATM, CASP3, CASQ1, CD300E, DARS, DOCK9, EXOSC9, EZH2, GDE1, IL34, NCOA4, NEFL, PDGFRL, PTGS1, REPS1, SCG2, SDPR, SIGLEC14, SIGLEC6, TAL1, TFEC, TIPIN, TPK1, UQCRB, USP9Y, WIPF1, ZCRB1 |
| CD8_Tcells___511 | Charoentong_et_al_2017 | ADRM1, AHSA1, C1GALT1C1, CCT6B, CD37, CD3D, CD3E, CD3G, CD69, CD8A, CETN3, CSE1L, GEMIN6, GNLY, GPT2, GZMA, GZMH, GZMK, IL2RB, LCK, MPZL1, NKG7, PIK3IP1, PTRH2, TIMM13, ZAP70 |
| CD8_Tcm___512 | Charoentong_et_al_2017 | ACTN4, ADAM12, ADCY9, F13A1, FCER1G, FCGR3B, FGF7, FKBP4, GLUD1, GM2A, GUSB, IL1RN, NOL11, NTRK1, RARA, RNF128, SIGLEC1, TNFRSF11A, TOX4, UBA52, ULBP1 |
| CD8_Tem___513 | Charoentong_et_al_2017 | ACAP1, APOL3, ARHGAP10, ATP10D, C3AR1, CCR5, CD160, CD55, CFLAR, CMKLR1, DAPP1, FCRL6, FLT3LG, GZMM, HAPLN3, HLA-DMB, HLA-DPA1, HLA-DPB1, IFI16, LIME1, LTK, NFKBIA, SETD7, SIK1, TRIB2 |
| Eosinophils___514 | Charoentong_et_al_2017 | GIPR, KRT18P50, LRMP, FOSB, RRP12, GPR183, NR4A3, ST3GAL6, DEPDC5, PDE6C, PKD2L2, GPR65, IL5RA, P2RY14, DACH1, DAPK2, EMR3 |
| Macrophages___515 | Charoentong_et_al_2017 | AIF1, CCL1, CCL14, CCL23, CCL26, CD300LB, CNR1, CNR2, EIF1, EIF4A1, FPR1, FPR2, FRAT2, GPR27, GPR77, RNASE2, MS4A2, BASP1, IGSF6, HK3, VNN1, FES, NPL, FZD2, FAM198B, HNMT, SLC15A3, CD4, TXNDC3, FRMD4A, CRYBB1, HRH1, WNT5B |
| Mast_cells___516 | Charoentong_et_al_2017 | ADAMTS3, CPA3, CMA1, CTSG, ARHGAP15, CPM, FCN1, FTL, HSPA6, ITGA9, RNASE3, S100A4, SIGLEC8, SLC6A4, PTGS2, EGR3, PILRA |
| Memory_Bcells___517 | Charoentong_et_al_2017 | AICDA, CCNA2, CDKN3, CLCN5, ENPP1, FCER1A, FCRL4, MYC, RUNX2, SORL1, SOX5, STAT5A, STAT5B, TLR9 |
| Monocytes___518 | Charoentong_et_al_2017 | ASGR2, CFP, ASGR1, CD1D, UPK3A, ACTG1, ANXA5, ATP6V1B2, CFL1, DAZAP2, CTBS, EMR4P, HIVEP2, MARCKSL1, MBP, MMP15, PNPLA6, TMBIM6, PQBP1, TEX264, IKZF1 |
| NK_cells___519 | Charoentong_et_al_2017 | AKT3, AXL, BST2, CDH2, CRTAM, CSF2RA, CTSZ, CXCL1, CYTH1, DAXX, DGKH, DLL4, DPYD, ERBB3, F11R, FAM27A, FAM49A, FASLG, FCGR1A, FN1, FSTL1, FUCA1, GBP3, GLS2, GRB2, LST1, BCL2, CDC5L, FGF18, FUT5, FZR1, GAGE2, IGFBP5, KANK2, LDB3 |
| NKT___520 | Charoentong_et_al_2017 | BTN2A2, CD101, CD109, CNPY3, CNPY4, CREB1, CRTC2, CRTC3, CSF2, KLRC1, FUT4, ICAM2, IL32, LAMP2, LILRB5, KLRG1, HSPA4, HSPB6, ISM2, ITIH2, KDM4C, |

| | | |
|---|---|---|
| | | KIR2DS4, KIRREL3, SDCBP, NFATC2IP, MICB, KIR2DL1, KIR2DL3, KIR3DL1, KIR3DL2, NCR1, FOSL1, TSLP, SLC7A7, SPP1, TREM2, UBASH3A, YBX2, CCDC88A, CLEC1A, THBD, PDPN, VCAM1, EMR1 |
| Neutrophils___521 | Charoentong_et_al_2017 | CREB5, CDA, CHST15, S100A12, APOBEC3A, CASP5, MMP25, HAL, C1orf183, FFAR2, MAK, CXCR1, STEAP4, MGAM, BTNL8, CXCR2, TNFRSF10C, VNN3 |
| Tgd_cells___522 | Charoentong_et_al_2017 | ACP5, AQP9, BTN3A2, C1orf54, CARD8, CCL18, CD209, CD33, CD36, CDK5, IL10RB, KLRF1, LGALS1, MAPK7, KLHL7, KRT80, LAMC1, LCORL, LMNB1, MEIS3P1, MPL, FABP1, FABP5, FADD, MFAP3L, MINPP1, RPS24, RPS7, RPS9, ABP1, CCL13 |
| Th1_cells___523 | Charoentong_et_al_2017 | CD70, TBX21, ADAM8, AHCYL2, ALCAM, B3GALNT1, BBS12, BST1, CD151, CD47, CD48, CD52, CD53, CD59, CD6, CD68, CD7, CD96, CFHR3, CHRM3, CLEC7A, COL23A1, COL4A4, COL5A3, DAB1, DLEU7, DOC2B, EMP1, F12, FURIN, GAB3, GATM, GFPT2, GPR25, GREM2, HAVCR1, HSD11B1, HUNK, IGF2, RCSD1, RYR1, SAV1, SELE, SELP, SH3KBP1, SIT1, SLC35B3, SIGLEC10, SKAP1, THUMPD2, TIGIT, ZEB2, ENC1, FAM134B, FBXO30, FCGR2C, STAC, LTC4S, MAN1B1, MDH1, MMD, RGS16, IL12A, P2RX5, CD97, ITGB4, ICAM3, METRNL, TNFRSF1A, IRF1, HTR2B, CALD1, MOCOS, TRAF3IP2, TLR8, TRAF1, DUSP14 |
| Th2_cells___524 | Charoentong_et_al_2017 | ASB2, CSRP2, DAPK1, DLC1, DNAJC12, DUSP6, GNAI1, LAMP3, NRP2, OSBPL1A, PDE4B, PHLDA1, PLA2G4A, RAB27B, RBMS3, RNF125, TMPRSS3, GATA3 |
| Tregs___525 | Charoentong_et_al_2017 | CCL3L1, CD72, CLEC5A, FOXP3, ITGA4, L1CAM, LIPA, LRP1, LRRC42, MARCO, MMP12, MNDA, MRC1, MS4A6A, PELO, PLEK, PRSS23, PTGIR, ST8SIA4, STAB1 |
| aDC___526 | Charoentong_et_al_2017 | ABCD1, C1QC, CAPG, CCL3L3, CD207, CD302, ATP5B, ATP5L, ATP6V1A, BCL2L1, C1QB, SNURF, SPCS3, CCNA1, CEACAM8, NOS2, SRA1, TNFRSF6B, TREM1, TREML1, RHOA, SLC25A37, TNFSF14, TREML4, VNN2, XPO6, CLEC4C, TNFAIP2, UBD, ACTR3, RAB1A, SLA, HLA-DQA2, SIGLEC5, SLAMF9 |
| iDC___527 | Charoentong_et_al_2017 | ACADM, AHCYL1, ALDH1A2, ALDH3A2, ALDH9A1, ALOX15, AMT, ARL1, ATIC, ATP5A1, CAPZA1, LILRA5, RDX, RRAGD, TACSTD2, INPP5F, RAB38, PLAU, CSF3R, SLC18A2, AMPD2, CLTB, C1orf162 |
| pDC___528 | Charoentong_et_al_2017 | CBX6, DAB2, DDX17, HIGD1A, IDH3A, IL3RA, MAGED1, NUCB2, OFD1, OGT, PDIA4, SERTAD2, SIRPA, TMED2, ENG, FCAR, IGF1, ITGA2B, GABARAP, GPX1, KRT23, PROK2, RALB, RETNLB, RNF141, SEC14L1, SEPX1, EMP3, CD300LF, ABTB1, KLHL21, PHRF1 |
| Bcells___529 | Rooney_et_al_2015 | CD79B, BTLA, FCRL3, BANK1, CD79A, BLK, RALGPS2, FCRL1, HVCN1, BACH2 |
| CD4_Tcells___530 | Rooney_et_al_2015 | FOXP3, C15orf53, IL5, CTLA4, IL32, GPR15, IL4 |
| CD8_Tcells___531 | Rooney_et_al_2015 | CD8A |
| DC___532 | Rooney_et_al_2015 | LILRA4, CLEC4C, PLD4, PHEX, IL3RA, PTCRA, IRF8, IRF7, GZMB, CXCR3 |
| Macrophages___533 | Rooney_et_al_2015 | FUCA1, MMP9, LGMN, HS3ST2, TM4SF19, CLEC5A, GPNMB, C11orf45, CD68, CYBB |
| NK_cells___534 | Rooney_et_al_2015 | KLRF1, KLRC1 |
| Neutrophils___535 | Rooney_et_al_2015 | KDM6B, HSD17B11, EVI2B, MNDA, MEGF9, SELL, NLRP12, PADI4, TRANK1, VNN3 |
| Bcells___536 | Tirosh_et_al_2016b | CD19, CD79A, CD79B, BLK, MS4A1, BANK1, IGLL3P, FCRL1, PAX5, CLEC17A, CD22, BCL11A, VPREB3, HLA-DOB, STAP1, FAM129C, TLR10, RALGPS2, AFF3, POU2AF1, CXCR5, PLCG2, HVCN1, CCR6, P2RX5, BLNK, KIAA0226L, POU2F2, IRF8, FCRLA, CD37 |
| Endothelial_cells___537 | Tirosh_et_al_2016b | PECAM1, VWF, CDH5, CLDN5, PLVAP, ECSCR, SLCO2A1, CCL14, MMRN1, MYCT1, KDR, TM4SF18, TIE1, ERG, FABP4, SDPR, HYAL2, FLT4, EGFL7, ESAM, CXorf36, TEK, TSPAN18, EMCN, MMRN2, ELTD1, PDE2A, NOS3, ROBO4, APOLD1, PTPRB, RHOJ, RAMP2, GPR116, F2RL3, JUP, CCBP2, GPR146, RGS16, TSPAN7, RAMP3, PLA2G4C, TGM2, LDB2, PRCP, ID1, SMAD1, AFAP1L1, ELK3, ANGPT2, LYVE1, ARHGAP29, IL3RA, ADCY4, TFPI, TNFAIP1, SYT15, DYSF, PODXL, SEMA3A, DOCK9, F8, NPDC1, TSPAN15, CD34, THBD, ITGB4, RASA4, COL4A1, ECE1, GFOD2, EFNA1, PVRL2, GNG11, HERC2P2, MALL, HERC2P9, PPM1F, PKP4, LIMS3, CD9, RAI14, ZNF521, RGL2, HSPG2, TGFBR2, RBP1, FXYD6, MATN2, S1PR1, PIEZO1, PDGFA, ADAM15, HAPLN3, APP |
| Fibroblasts___538 | Tirosh_et_al_2016b | FAP, THY1, DCN, COL1A1, COL1A2, COL6A1, COL6A2, COL6A3, CXCL14, LUM, COL3A1, DPT, ISLR, PODN, CD248, FGF7, MXRA8, PDGFRL, COL14A1, MFAP5, MEG3, |

| | | |
|---|---|---|
| | | SULF1, AOX1, SVEP1, LPAR1, PDGFRB, TAGLN, IGFBP6, FBLN1, CA12, SPOCK1, TPM2, THBS2, FBLN5, TMEM119, ADAM33, PRRX1, PCOLCE, IGF2, GFPT2, PDGFRA, CRISPLD2, CPE, F3, MFAP4, C1S, PTGIS, LOX, CYP1B1, CLDN11, SERPINF1, OLFML3, COL5A2, ACTA2, MSC, VASN, ABI3BP, C1R, ANTXR1, MGST1, C3, PALLD, FBN1, CPXM1, CYBRD1, IGFBP5, PRELP, PAPSS2, MMP2, CKAP4, CCDC80, ADAMTS2, TPM1, PCSK5, ELN, CXCL12, OLFML2B, PLAC9, RCN3, LTBP2, NID2, SCARA3, AMOTL2, TPST1, MIR100HG, CTGF, RARRES2, FHL2 |
| Macrophages___539 | Tirosh_et_al_2016b | CD163, CD14, CSF1R, C1QC, VSIG4, C1QA, FCER1G, F13A1, TYROBP, MSR1, C1QB, MS4A4A, FPR1, S100A9, IGSF6, LILRB4, FPR3, SIGLEC1, LILRA1, LYZ, HK3, SLC11A1, CSF3R, CD300E, PILRA, FCGR3A, AIF1, SIGLEC9, FCGR1C, OLR1, TLR2, LILRB2, C5AR1, FCGR1A, MS4A6A, C3AR1, HCK, IL4I1, LST1, LILRA5, CSTA, IFI30, CD68, TBXAS1, FCGR1B, LILRA6, CXCL16, NCF2, RAB20, MS4A7, NLRP3, LRRC25, ADAP2, SPP1, CCR1, TNFSF13, RASSF4, SERPINA1, MAFB, IL18, FGL2, SIRPB1, CLEC4A, MNDA, FCGR2A, CLEC7A, SLAMF8, SLC7A7, ITGAX, BCL2A1, PLAUR, SLCO2B1, PLBD1, APOC1, RNF144B, SLC31A2, PTAFR, NINJ1, ITGAM, CPVL, PLIN2, C1orf162, FTL, LIPA, CD86, GLUL, FGR, GK, TYMP, GPX1, NPL, ACSL1 |
| Melanocytes___540 | Tirosh_et_al_2016b | MIA, TYR, SLC45A2, CDH19, PMEL, SLC24A5, MAGEA6, GJB1, PLP1, PRAME, CAPN3, ERBB3, GPM6B, S100B, FXYD3, PAX3, S100A1, MLANA, SLC26A2, GPR143, CSPG4, SOX10, MLPH, LOXL4, PLEKHB1, RAB38, QPCT, BIRC7, MFI2, LINC00473, SEMA3B, SERPINA3, PIR, MITF, ST6GALNAC2, ROPN1B, CDH1, ABCB5, QDPR, SERPINE2, ATP1A1, ST3GAL4, CDK2, ACSL3, NT5DC3, IGSF8, MBP |

**Supplementary Data 2A. The grouping of GO terms in Cohort 1.**

| GO_term | Enrichment in | Type 1 | Type 2 | Type 3 | Type 4 |
|---|---|---|---|---|---|
| GO_REGULATION_OF_VASCULAR_ENDOTHELIAL_GROWTH_FACTOR_PRODUCTION | NIR | angiogenesis | | | |
| GO_REGULATION_OF_VASCULATURE_DEVELOPMENT | NIR | angiogenesis | | | |
| GO_POSITIVE_REGULATION_OF_VASCULATURE_DEVELOPMENT | NIR | angiogenesis | | | |
| GO_POSITIVE_REGULATION_OF_VASODILATION | NIR | angiogenesis | | | |
| GO_CELL_CELL_ADHESION_VIA_PLASMA_MEMBRANE_ADHESION_MOLECULES | PIR | Cell migration | cell adhesion | | |
| GO_HOMOPHILIC_CELL_ADHESION_VIA_PLASMA_MEMBRANE_ADHESION_MOLECULES | PIR | Cell migration | cell adhesion | | |
| GO_NEURON_CELL_CELL_ADHESION | PIR | immune system | cell adhesion | | |
| GO_SYNAPSE_ASSEMBLY | PIR | immune system | cell adhesion | | |
| GO_REGULATION_OF_CELL_CELL_ADHESION | NIR | immune system | cell adhesion | | |
| GO_PROTEIN_BINDING_INVOLVED_IN_CELL_ADHESION | NIR | immune system | cell adhesion | | |
| GO_DESMOSOME | NIR | immune system | cell adhesion | | |
| GO_REGULATION_OF_CELL_ADHESION_MEDIATED_BY_INTEGRIN | NIR | immune system | cell adhesion | | |
| GO_REGULATION_OF_HOMOTYPIC_CELL_CELL_ADHESION | NIR | immune system | cell adhesion | | |
| GO_POSITIVE_REGULATION_OF_CELL_ADHESION_MEDIATED_BY_INTEGRIN | NIR | immune system | cell adhesion | | |
| GO_HETEROTYPIC_CELL_CELL_ADHESION | NIR | immune system | cell adhesion | | |
| GO_NEGATIVE_REGULATION_OF_CELL_CELL_ADHESION | NIR | immune system | cell adhesion | | |
| GO_NEGATIVE_REGULATION_OF_HOMOTYPIC_CELL_CELL_ADHESION | NIR | immune system | cell adhesion | | |
| GO_POSITIVE_REGULATION_OF_CELL_ADHESION | NIR | immune system | cell adhesion | | |
| GO_SINGLE_ORGANISM_CELL_ADHESION | NIR | immune system | cell adhesion | | |
| GO_POSITIVE_REGULATION_OF_CELL_CELL_ADHESION | NIR | immune system | cell adhesion | | |
| GO_LEUKOCYTE_CELL_CELL_ADHESION | NIR | immune system | cell adhesion | | |
| GO_REGULATION_OF_LYMPHOCYTE_APOPTOTIC_PROCESS | NIR | immune system | cell death | apoptosis | |
| GO_REGULATION_OF_T_CELL_APOPTOTIC_PROCESS | NIR | immune system | cell death | apoptosis | |
| GO_NEGATIVE_REGULATION_OF_T_CELL_APOPTOTIC_PROCESS | NIR | immune system | cell death | apoptosis | |
| GO_NEGATIVE_REGULATION_OF_LYMPHOCYTE_APOPTOTIC_PROCESS | NIR | immune system | cell death | apoptosis | |
| GO_NEGATIVE_REGULATION_OF_LEUKOCYTE_APOPTOTIC_PROCESS | NIR | immune system | cell death | apoptosis | |
| GO_REGULATION_OF_LEUKOCYTE_APOPTOTIC_PROCESS | NIR | immune system | cell death | apoptosis | |
| GO_NEGATIVE_REGULATION_OF_CELL_KILLING | NIR | immune system | cell death | cell kill | |
| GO_REGULATION_OF_CELL_KILLING | NIR | immune system | cell death | cell kill | |
| GO_CELL_KILLING | NIR | immune system | cell death | cell kill | |
| GO_NECROTIC_CELL_DEATH | NIR | immune system | cell death | necrosis | |
| GO_NECROPTOTIC_PROCESS | NIR | immune system | cell death | necrosis | |
| GO_REGULATION_OF_LYMPHOCYTE_CHEMOTAXIS | NIR | immune system | cell migration | | |
| GO_LEUKOCYTE_CHEMOTAXIS | NIR | immune system | cell migration | | |
| GO_DENDRITIC_CELL_CHEMOTAXIS | NIR | immune system | cell migration | | |
| GO_POSITIVE_REGULATION_OF_CHEMOTAXIS | NIR | immune system | cell migration | | |
| GO_REGULATION_OF_CELLULAR_EXTRAVASATION | NIR | immune system | cell migration | | |
| GO_REGULATION_OF_LEUKOCYTE_CHEMOTAXIS | NIR | immune system | cell migration | | |
| GO_GRANULOCYTE_MIGRATION | NIR | immune system | cell migration | | |
| GO_REGULATION_OF_NEUTROPHIL_MIGRATION | NIR | immune system | cell migration | | |
| GO_REGULATION_OF_CHEMOTAXIS | NIR | immune system | cell migration | | |
| GO_REGULATION_OF_MACROPHAGE_CHEMOTAXIS | NIR | immune system | cell migration | | |
| GO_REGULATION_OF_LEUKOCYTE_MIGRATION | NIR | immune system | cell migration | | |
| GO_CELL_CHEMOTAXIS | NIR | immune system | cell migration | | |
| GO_MYELOID_LEUKOCYTE_MIGRATION | NIR | immune system | cell migration | | |
| GO_POSITIVE_REGULATION_OF_LEUKOCYTE_CHEMOTAXIS | NIR | immune system | cell migration | | |
| GO_REGULATION_OF_GRANULOCYTE_CHEMOTAXIS | NIR | immune system | cell migration | | |
| GO_POSITIVE_REGULATION_OF_LYMPHOCYTE_MIGRATION | NIR | immune system | cell migration | | |
| GO_DENDRITIC_CELL_MIGRATION | NIR | immune system | cell migration | | |
| GO_POSITIVE_REGULATION_OF_NEUTROPHIL_MIGRATION | NIR | immune system | cell migration | | |
| GO_REGULATION_OF_T_CELL_MIGRATION | NIR | immune system | cell migration | | |
| GO_REGULATION_OF_PEPTIDE_TRANSPORT | NIR | immune system | cytokine | | |
| GO_REGULATION_OF_TUMOR_NECROSIS_FACTOR_SUPERFAMILY_CYTOKINE_PRODUCTION | NIR | immune system | cytokine | | |
| GO_REGULATION_OF_INTERLEUKIN_4_PRODUCTION | NIR | immune system | cytokine | | |
| GO_POSITIVE_REGULATION_OF_INTERLEUKIN_1_SECRETION | NIR | immune system | cytokine | | |
| GO_REGULATION_OF_INTERLEUKIN_5_PRODUCTION | NIR | immune system | cytokine | | |
| GO_REGULATION_OF_INTERLEUKIN_2_PRODUCTION | NIR | immune system | cytokine | | |
| GO_POSITIVE_REGULATION_OF_INTERLEUKIN_6_PRODUCTION | NIR | immune system | cytokine | | |
| GO_REGULATION_OF_INTERLEUKIN_6_PRODUCTION | NIR | immune system | cytokine | | |
| GO_POSITIVE_REGULATION_OF_INTERLEUKIN_1_BETA_PRODUCTION | NIR | immune system | cytokine | | |
| GO_REGULATION_OF_IMMUNOGLOBULIN_PRODUCTION | NIR | immune system | cytokine | | |
| GO_NEGATIVE_REGULATION_OF_CYTOKINE_SECRETION | NIR | immune system | cytokine | | |
| GO_POSITIVE_REGULATION_OF_TRANSCRIPTION_FACTOR_IMPORT_INTO_NUCLEUS | NIR | immune system | cytokine | | |
| GO_REGULATION_OF_INTERLEUKIN_12_PRODUCTION | NIR | immune system | cytokine | | |
| GO_POSITIVE_REGULATION_OF_NF_KAPPAB_IMPORT_INTO_NUCLEUS | NIR | immune system | cytokine | | |
| GO_POSITIVE_REGULATION_OF_INTERLEUKIN_1_PRODUCTION | NIR | immune system | cytokine | | |
| GO_NEGATIVE_REGULATION_OF_INTERFERON_GAMMA_PRODUCTION | NIR | immune system | cytokine | | |
| GO_PROTEIN_ACTIVATION_CASCADE | NIR | immune system | cytokine | | |
| GO_POSITIVE_REGULATION_OF_CHEMOKINE_PRODUCTION | NIR | immune system | cytokine | | |
| GO_NEGATIVE_REGULATION_OF_PROTEIN_SECRETION | NIR | immune system | cytokine | | |
| GO_POSITIVE_REGULATION_OF_INTERLEUKIN_8_PRODUCTION | NIR | immune system | cytokine | | |
| GO_REGULATION_OF_INTERLEUKIN_2_BIOSYNTHETIC_PROCESS | NIR | immune system | cytokine | | |
| O_NEGATIVE_REGULATION_OF_TUMOR_NECROSIS_FACTOR_SUPERFAMILY_CYTOKINE_PRODUCTIC | NIR | immune system | cytokine | | |
| GO_POSITIVE_REGULATION_OF_HORMONE_SECRETION | NIR | immune system | cytokine | | |
| GO_POSITIVE_REGULATION_OF_CYTOKINE_PRODUCTION | NIR | immune system | cytokine | | |
| GO_POSITIVE_REGULATION_OF_CYTOKINE_PRODUCTION_INVOLVED_IN_IMMUNE_RESPONSE | NIR | immune system | cytokine | | |
| GO_REGULATION_OF_INTERLEUKIN_1_BETA_PRODUCTION | NIR | immune system | cytokine | | |
| GO_CYTOKINE_PRODUCTION | NIR | immune system | cytokine | | |
| iO_POSITIVE_REGULATION_OF_TUMOR_NECROSIS_FACTOR_SUPERFAMILY_CYTOKINE_PRODUCTIO | NIR | immune system | cytokine | | |
| GO_POSITIVE_REGULATION_OF_INSULIN_SECRETION | NIR | immune system | cytokine | | |
| GO_NEGATIVE_REGULATION_OF_CYTOKINE_BIOSYNTHETIC_PROCESS | NIR | immune system | cytokine | | |
| GO_REGULATION_OF_PROTEIN_SECRETION | NIR | immune system | cytokine | | |
| GO_CYTOKINE_SECRETION | NIR | immune system | cytokine | | |
| GO_REGULATION_OF_INTERLEUKIN_8_PRODUCTION | NIR | immune system | cytokine | | |
| GO_POSITIVE_REGULATION_OF_PROTEIN_MATURATION | NIR | immune system | cytokine | | |

| | | | |
|---|---|---|---|
| GO_POSITIVE_REGULATION_OF_INTERLEUKIN_12_PRODUCTION | NIR | immune system | cytokine |
| GO_REGULATION_OF_PRODUCTION_OF_MOLECULAR_MEDIATOR_OF_IMMUNE_RESPONSE | NIR | immune system | cytokine |
| GO_REGULATION_OF_INTERLEUKIN_1_SECRETION | NIR | immune system | cytokine |
| GO_POSITIVE_REGULATION_OF_INTERFERON_GAMMA_PRODUCTION | NIR | immune system | cytokine |
| GO_REGULATION_OF_INTERLEUKIN_10_PRODUCTION | NIR | immune system | cytokine |
| GO_POSITIVE_REGULATION_OF_PROTEIN_SECRETION | NIR | immune system | cytokine |
| GO_NEGATIVE_REGULATION_OF_INTERLEUKIN_2_PRODUCTION | NIR | immune system | cytokine |
| ITIVE_REGULATION_OF_INSULIN_SECRETION_INVOLVED_IN_CELLULAR_RESPONSE_TO_GLUCOSE_S | NIR | immune system | cytokine |
| O_POSITIVE_REGULATION_OF_PRODUCTION_OF_MOLECULAR_MEDIATOR_OF_IMMUNE_RESPONS | NIR | immune system | cytokine |
| GO_POSITIVE_REGULATION_OF_CYTOKINE_SECRETION | NIR | immune system | cytokine |
| GO_POSITIVE_REGULATION_OF_CYTOKINE_BIOSYNTHETIC_PROCESS | NIR | immune system | cytokine |
| GO_REGULATION_OF_CYTOKINE_SECRETION | NIR | immune system | cytokine |
| GO_NEGATIVE_REGULATION_OF_CYTOKINE_PRODUCTION | NIR | immune system | cytokine |
| GO_POSITIVE_REGULATION_OF_INTERLEUKIN_4_PRODUCTION | NIR | immune system | cytokine |
| GO_POSITIVE_REGULATION_OF_SECRETION | NIR | immune system | cytokine |
| GO_POSITIVE_REGULATION_OF_INTERLEUKIN_10_PRODUCTION | NIR | immune system | cytokine |
| GO_INTERLEUKIN_1_PRODUCTION | NIR | immune system | cytokine |
| GO_REGULATION_OF_INTERFERON_GAMMA_PRODUCTION | NIR | immune system | cytokine |
| GO_REGULATION_OF_INTERLEUKIN_1_PRODUCTION | NIR | immune system | cytokine |
| GO_REGULATION_OF_CYTOKINE_BIOSYNTHETIC_PROCESS | NIR | immune system | cytokine |
| GO_NEGATIVE_REGULATION_OF_INTERLEUKIN_6_PRODUCTION | NIR | immune system | cytokine |
| GO_REGULATION_OF_CYTOKINE_PRODUCTION_INVOLVED_IN_IMMUNE_RESPONSE | NIR | immune system | cytokine |
| GO_NEGATIVE_REGULATION_OF_CYTOKINE_PRODUCTION_INVOLVED_IN_IMMUNE_RESPONSE | NIR | immune system | cytokine |
| GO_REGULATION_OF_PROTEIN_MATURATION | NIR | immune system | cytokine |
| GO_NEGATIVE_REGULATION_OF_CHEMOKINE_PRODUCTION | NIR | immune system | cytokine |
| GO_REGULATION_OF_CHEMOKINE_PRODUCTION | NIR | immune system | cytokine |
| GO_CYTOKINE_ACTIVITY | NIR | immune system | cytokine |
| GO_POSITIVE_REGULATION_OF_IMMUNOGLOBULIN_PRODUCTION | NIR | immune system | cytokine |
| GO_REGULATION_OF_NF_KAPPAB_IMPORT_INTO_NUCLEUS | NIR | immune system | cytokine |
| GO_COMPLEMENT_ACTIVATION | NIR | immune system | cytokine |
| GO_REGULATION_OF_TUMOR_NECROSIS_FACTOR_BIOSYNTHETIC_PROCESS | NIR | immune system | cytokine |
| GO_NEGATIVE_REGULATION_OF_TYPE_I_INTERFERON_PRODUCTION | NIR | immune system | cytokine |
| GO_POSITIVE_REGULATION_OF_PEPTIDE_SECRETION | NIR | immune system | cytokine |
| GO_REGULATION_OF_INTERLEUKIN_8_SECRETION | NIR | immune system | cytokine |
| O_NEGATIVE_REGULATION_OF_PRODUCTION_OF_MOLECULAR_MEDIATOR_OF_IMMUNE_RESPON | NIR | immune system | cytokine |
| GO_PRODUCTION_OF_MOLECULAR_MEDIATOR_OF_IMMUNE_RESPONSE | NIR | immune system | cytokine |
| GO_CHEMOKINE_ACTIVITY | NIR | immune system | cytokine |
| GO_MHC_CLASS_I_PROTEIN_BINDING | NIR | immune system | cytokine |
| GO_MHC_PROTEIN_BINDING | NIR | immune system | cytokine |
| GO_NEGATIVE_REGULATION_OF_MULTI_ORGANISM_PROCESS | NIR | immune system | defense response |
| GO_RESPONSE_TO_PROTOZOAN | NIR | immune system | defense response |
| GO_DEFENSE_RESPONSE_TO_GRAM_NEGATIVE_BACTERIUM | NIR | immune system | defense response |
| GO_DETECTION_OF_OTHER_ORGANISM | NIR | immune system | defense response |
| GO_MODULATION_OF_GROWTH_OF_SYMBIONT_INVOLVED_IN_INTERACTION_WITH_HOST | NIR | immune system | defense response |
| GO_RESPONSE_TO_VIRUS | NIR | immune system | defense response |
| GO_DETECTION_OF_BIOTIC_STIMULUS | NIR | immune system | defense response |
| GO_NEGATIVE_REGULATION_OF_VIRAL_ENTRY_INTO_HOST_CELL | NIR | immune system | defense response |
| GO_POSITIVE_REGULATION_OF_RESPONSE_TO_EXTERNAL_STIMULUS | NIR | immune system | defense response |
| GO_DEFENSE_RESPONSE_TO_BACTERIUM | NIR | immune system | defense response |
| GO_RESPONSE_TO_BACTERIUM | NIR | immune system | defense response |
| GO_KILLING_OF_CELLS_OF_OTHER_ORGANISM | NIR | immune system | defense response |
| GO_DISRUPTION_OF_CELLS_OF_OTHER_ORGANISM | NIR | immune system | defense response |
| GO_RESPONSE_TO_FUNGUS | NIR | immune system | defense response |
| GO_INFLAMMATORY_RESPONSE_TO_ANTIGENIC_STIMULUS | NIR | immune system | defense response |
| GO_REGULATION_OF_VIRAL_ENTRY_INTO_HOST_CELL | NIR | immune system | defense response |
| GO_DEFENSE_RESPONSE_TO_OTHER_ORGANISM | NIR | immune system | defense response |
| GO_CELLULAR_RESPONSE_TO_INTERFERON_GAMMA | NIR | immune system | immune response |
| ISE_BASED_ON_SOMATIC_RECOMBINATION_OF_IMMUNE_RECEPTORS_BUILT_FROM_IMMUNOGL | NIR | immune system | immune response |
| GO_ACTIVATION_OF_INNATE_IMMUNE_RESPONSE | NIR | immune system | immune response |
| GO_REGULATION_OF_CELL_ACTIVATION | NIR | immune system | immune response |
| GO_ACUTE_INFLAMMATORY_RESPONSE | NIR | immune system | immune response |
| GO_POSITIVE_REGULATION_OF_INFLAMMATORY_RESPONSE | NIR | immune system | immune response |
| GO_POSITIVE_REGULATION_OF_RESPONSE_TO_WOUNDING | NIR | immune system | immune response |
| GO_REGULATION_OF_PROTEIN_ACTIVATION_CASCADE | NIR | immune system | immune response |
| GO_REGULATION_OF_ACUTE_INFLAMMATORY_RESPONSE | NIR | immune system | immune response |
| GO_NEGATIVE_REGULATION_OF_IMMUNE_RESPONSE | NIR | immune system | immune response |
| GO_POSITIVE_REGULATION_OF_ADAPTIVE_IMMUNE_RESPONSE | NIR | immune system | immune response |
| GO_INFLAMMATORY_RESPONSE | NIR | immune system | immune response |
| GO_POSITIVE_REGULATION_OF_INNATE_IMMUNE_RESPONSE | NIR | immune system | immune response |
| GO_T_HELPER_1_TYPE_IMMUNE_RESPONSE | NIR | immune system | immune response |
| GO_REGULATION_OF_RESPONSE_TO_WOUNDING | NIR | immune system | immune response |
| GO_REGULATION_OF_HUMORAL_IMMUNE_RESPONSE | NIR | immune system | immune response |
| GO_ACUTE_PHASE_RESPONSE | NIR | immune system | immune response |
| GO_NEGATIVE_REGULATION_OF_IMMUNE_EFFECTOR_PROCESS | NIR | immune system | immune response |
| GO_POSITIVE_REGULATION_OF_HUMORAL_IMMUNE_RESPONSE | NIR | immune system | immune response |
| GO_RESPONSE_TO_INTERFERON_GAMMA | NIR | immune system | immune response |
| GO_POSITIVE_REGULATION_OF_IMMUNE_EFFECTOR_PROCESS | NIR | immune system | immune response |
| GO_RESPONSE_TO_TUMOR_NECROSIS_FACTOR | NIR | immune system | immune response |
| GO_CELL_ACTIVATION_INVOLVED_IN_IMMUNE_RESPONSE | NIR | immune system | immune response |
| GO_NEGATIVE_REGULATION_OF_CELL_ACTIVATION | NIR | immune system | immune response |
| GO_REGULATION_OF_TYPE_2_IMMUNE_RESPONSE | NIR | immune system | immune response |
| GO_REGULATION_OF_IMMUNE_EFFECTOR_PROCESS | NIR | immune system | immune response |
| GO_POSITIVE_REGULATION_OF_DEFENSE_RESPONSE | NIR | immune system | immune response |
| GO_HUMORAL_IMMUNE_RESPONSE | NIR | immune system | immune response |
| GO_NEGATIVE_REGULATION_OF_DEFENSE_RESPONSE | NIR | immune system | immune response |
| GO_ACTIVATION_OF_IMMUNE_RESPONSE | NIR | immune system | immune response |
| GO_REGULATION_OF_ADAPTIVE_IMMUNE_RESPONSE | NIR | immune system | immune response |

| | | | | | |
|---|---|---|---|---|---|
| GO_IMMUNE_EFFECTOR_PROCESS | NIR | immune system | immune response | | |
| GO_REGULATION_OF_INFLAMMATORY_RESPONSE | NIR | immune system | immune response | | |
| GO_RESPIRATORY_BURST | NIR | immune system | immune response | | |
| GO_POSITIVE_REGULATION_OF_CELL_ACTIVATION | NIR | immune system | immune response | | |
| GO_NEGATIVE_REGULATION_OF_ADAPTIVE_IMMUNE_RESPONSE | NIR | immune system | immune response | | |
| GO_NEGATIVE_REGULATION_OF_INNATE_IMMUNE_RESPONSE | NIR | immune system | immune response | | |
| GO_ADAPTIVE_IMMUNE_RESPONSE | NIR | immune system | immune response | | |
| GO_NEGATIVE_REGULATION_OF_IMMUNE_SYSTEM_PROCESS | NIR | immune system | immune response | | |
| GO_REGULATION_OF_INNATE_IMMUNE_RESPONSE | NIR | immune system | immune response | | |
| NSE_BASED_ON_SOMATIC_RECOMBINATION_OF_IMMUNE_RECEPTORS_BUILT_FROM_IMMUNOGL | NIR | immune system | immune response | | |
| GO_HUMORAL_IMMUNE_RESPONSE | NIR | immune system | lymphoid lineage | B cell | immune response |
| GO_REGULATION_OF_HUMORAL_IMMUNE_RESPONSE | NIR | immune system | lymphoid lineage | B cell | immune response |
| GO_POSITIVE_REGULATION_OF_HUMORAL_IMMUNE_RESPONSE | NIR | immune system | lymphoid lineage | B cell | immune response |
| GO_REGULATION_OF_IMMUNOGLOBULIN_SECRETION | NIR | immune system | lymphoid lineage | B cell | immunoglobulin |
| GO_HUMORAL_IMMUNE_RESPONSE_MEDIATED_BY_CIRCULATING_IMMUNOGLOBULIN | NIR | immune system | lymphoid lineage | B cell | immunoglobulin |
| GO_REGULATION_OF_B_CELL_MEDIATED_IMMUNITY | NIR | immune system | lymphoid lineage | B cell | |
| GO_B_CELL_PROLIFERATION | NIR | immune system | lymphoid lineage | B cell | |
| GO_POSITIVE_REGULATION_OF_B_CELL_ACTIVATION | NIR | immune system | lymphoid lineage | B cell | |
| GO_POSITIVE_REGULATION_OF_B_CELL_MEDIATED_IMMUNITY | NIR | immune system | lymphoid lineage | B cell | |
| GO_REGULATION_OF_B_CELL_DIFFERENTIATION | NIR | immune system | lymphoid lineage | B cell | |
| GO_REGULATION_OF_B_CELL_ACTIVATION | NIR | immune system | lymphoid lineage | B cell | |
| GO_B_CELL_ACTIVATION | NIR | immune system | lymphoid lineage | B cell | |
| GO_B_CELL_MEDIATED_IMMUNITY | NIR | immune system | lymphoid lineage | B cell | |
| GO_NEGATIVE_REGULATION_OF_B_CELL_ACTIVATION | NIR | immune system | lymphoid lineage | B cell | |
| GO_MATURE_B_CELL_DIFFERENTIATION | NIR | immune system | lymphoid lineage | B cell | |
| GO_CD4_POSITIVE_ALPHA_BETA_T_CELL_ACTIVATION | NIR | immune system | lymphoid lineage | T cell | |
| GO_REGULATION_OF_ALPHA_BETA_T_CELL_DIFFERENTIATION | NIR | immune system | lymphoid lineage | T cell | |
| GO_POSITIVE_T_CELL_SELECTION | NIR | immune system | lymphoid lineage | T cell | |
| GO_REGULATION_OF_T_CELL_DIFFERENTIATION | NIR | immune system | lymphoid lineage | T cell | |
| GO_ALPHA_BETA_T_CELL_DIFFERENTIATION | NIR | immune system | lymphoid lineage | T cell | |
| GO_THYMIC_T_CELL_SELECTION | NIR | immune system | lymphoid lineage | T cell | |
| GO_POSITIVE_REGULATION_OF_ALPHA_BETA_T_CELL_ACTIVATION | NIR | immune system | lymphoid lineage | T cell | |
| GO_T_CELL_RECEPTOR_COMPLEX | NIR | immune system | lymphoid lineage | T cell | |
| GO_ALPHA_BETA_T_CELL_ACTIVATION | NIR | immune system | lymphoid lineage | T cell | |
| GO_T_CELL_MEDIATED_IMMUNITY | NIR | immune system | lymphoid lineage | T cell | |
| GO_NEGATIVE_REGULATION_OF_T_CELL_PROLIFERATION | NIR | immune system | lymphoid lineage | T cell | |
| GO_NEGATIVE_REGULATION_OF_ALPHA_BETA_T_CELL_ACTIVATION | NIR | immune system | lymphoid lineage | T cell | |
| GO_NEGATIVE_REGULATION_OF_T_CELL_MEDIATED_IMMUNITY | NIR | immune system | lymphoid lineage | T cell | |
| GO_REGULATION_OF_T_CELL_PROLIFERATION | NIR | immune system | lymphoid lineage | T cell | |
| GO_T_CELL_DIFFERENTIATION_IN_THYMUS | NIR | immune system | lymphoid lineage | T cell | |
| GO_T_CELL_ACTIVATION_INVOLVED_IN_IMMUNE_RESPONSE | NIR | immune system | lymphoid lineage | T cell | |
| GO_REGULATION_OF_T_HELPER_CELL_DIFFERENTIATION | NIR | immune system | lymphoid lineage | T cell | |
| GO_REGULATION_OF_ALPHA_BETA_T_CELL_PROLIFERATION | NIR | immune system | lymphoid lineage | T cell | |
| GO_POSITIVE_REGULATION_OF_CD4_POSITIVE_ALPHA_BETA_T_CELL_ACTIVATION | NIR | immune system | lymphoid lineage | T cell | |
| GO_T_CELL_SELECTION | NIR | immune system | lymphoid lineage | T cell | |
| GO_REGULATION_OF_CD4_POSITIVE_ALPHA_BETA_T_CELL_ACTIVATION | NIR | immune system | lymphoid lineage | T cell | |
| GO_T_CELL_DIFFERENTIATION | NIR | immune system | lymphoid lineage | T cell | |
| GO_POSITIVE_REGULATION_OF_T_CELL_PROLIFERATION | NIR | immune system | lymphoid lineage | T cell | |
| GO_T_CELL_LINEAGE_COMMITMENT | NIR | immune system | lymphoid lineage | T cell | |
| GO_THYMOCYTE_AGGREGATION | NIR | immune system | lymphoid lineage | T cell | |
| GO_NEGATIVE_REGULATION_OF_T_CELL_DIFFERENTIATION | NIR | immune system | lymphoid lineage | T cell | |
| GO_POSITIVE_REGULATION_OF_ALPHA_BETA_T_CELL_DIFFERENTIATION | NIR | immune system | lymphoid lineage | T cell | |
| GO_REGULATION_OF_ALPHA_BETA_T_CELL_ACTIVATION | NIR | immune system | lymphoid lineage | T cell | |
| GO_REGULATION_OF_T_CELL_MEDIATED_IMMUNITY | NIR | immune system | lymphoid lineage | T cell | |
| GO_POSITIVE_REGULATION_OF_ALPHA_BETA_T_CELL_PROLIFERATION | NIR | immune system | lymphoid lineage | T cell | |
| GO_T_CELL_DIFFERENTIATION_INVOLVED_IN_IMMUNE_RESPONSE | NIR | immune system | lymphoid lineage | T cell | |
| GO_NEGATIVE_REGULATION_OF_CD4_POSITIVE_ALPHA_BETA_T_CELL_ACTIVATION | NIR | immune system | lymphoid lineage | T cell | |
| GO_NEGATIVE_REGULATION_OF_T_CELL_RECEPTOR_SIGNALING_PATHWAY | NIR | immune system | lymphoid lineage | T cell | |
| NSE_BASED_ON_SOMATIC_RECOMBINATION_OF_IMMUNE_RECEPTORS_BUILT_FROM_IMMUNOGL | NIR | immune system | lymphoid lineage | | immune response |
| GO_REGULATION_OF_ADAPTIVE_IMMUNE_RESPONSE | NIR | immune system | lymphoid lineage | | immune response |
| GO_NEGATIVE_REGULATION_OF_ADAPTIVE_IMMUNE_RESPONSE | NIR | immune system | lymphoid lineage | | immune response |
| GO_LYMPHOCYTE_ACTIVATION | NIR | immune system | lymphoid lineage | | |
| GO_NATURAL_KILLER_CELL_MEDIATED_IMMUNITY | NIR | immune system | lymphoid lineage | | |
| GO_NEGATIVE_REGULATION_OF_LYMPHOCYTE_DIFFERENTIATION | NIR | immune system | lymphoid lineage | | |
| GO_LYMPHOCYTE_DIFFERENTIATION | NIR | immune system | lymphoid lineage | | |
| GO_REGULATION_OF_LYMPHOCYTE_DIFFERENTIATION | NIR | immune system | lymphoid lineage | | |
| GO_REGULATION_OF_NATURAL_KILLER_CELL_MEDIATED_IMMUNITY | NIR | immune system | lymphoid lineage | | |
| GO_LYMPHOCYTE_MEDIATED_IMMUNITY | NIR | immune system | lymphoid lineage | | |
| GO_POSITIVE_REGULATION_OF_LYMPHOCYTE_MEDIATED_IMMUNITY | NIR | immune system | lymphoid lineage | | |
| GO_NATURAL_KILLER_CELL_ACTIVATION | NIR | immune system | lymphoid lineage | | |
| GO_POSITIVE_REGULATION_OF_LYMPHOCYTE_DIFFERENTIATION | NIR | immune system | lymphoid lineage | | |
| GO_REGULATION_OF_COAGULATION | NIR | immune system | Myeloid lineage | | |
| GO_REGULATION_OF_PLATELET_ACTIVATION | NIR | immune system | Myeloid lineage | | |
| GO_POSITIVE_REGULATION_OF_HEMOPOIESIS | NIR | immune system | Myeloid lineage | | |
| GO_REGULATION_OF_MAST_CELL_ACTIVATION_INVOLVED_IN_IMMUNE_RESPONSE | NIR | immune system | Myeloid lineage | | |
| GO_MYELOID_LEUKOCYTE_ACTIVATION | NIR | immune system | Myeloid lineage | | |
| GO_POSITIVE_REGULATION_OF_LEUKOCYTE_MEDIATED_IMMUNITY | NIR | immune system | Myeloid lineage | | |
| GO_NEGATIVE_REGULATION_OF_LEUKOCYTE_PROLIFERATION | NIR | immune system | Myeloid lineage | | |
| GO_LEUKOCYTE_PROLIFERATION | NIR | immune system | Myeloid lineage | | |
| GO_REGULATION_OF_LEUKOCYTE_DIFFERENTIATION | NIR | immune system | Myeloid lineage | | |
| GO_LEUKOCYTE_DEGRANULATION | NIR | immune system | Myeloid lineage | | |
| GO_REGULATION_OF_LEUKOCYTE_DEGRANULATION | NIR | immune system | Myeloid lineage | | |
| GO_LEUKOCYTE_DIFFERENTIATION | NIR | immune system | Myeloid lineage | | |
| GO_MYELOID_CELL_ACTIVATION_INVOLVED_IN_IMMUNE_RESPONSE | NIR | immune system | Myeloid lineage | | |
| GO_LEUKOCYTE_MEDIATED_CYTOTOXICITY | NIR | immune system | Myeloid lineage | | |
| GO_POSITIVE_REGULATION_OF_MYELOID_LEUKOCYTE_MEDIATED_IMMUNITY | NIR | immune system | Myeloid lineage | | |
| GO_MAST_CELL_MEDIATED_IMMUNITY | NIR | immune system | Myeloid lineage | | |

| | | | |
|---|---|---|---|
| GO_MYELOID_LEUKOCYTE_MEDIATED_IMMUNITY | NIR | immune system | Myeloid lineage |
| GO_POSITIVE_REGULATION_OF_MAST_CELL_ACTIVATION | NIR | immune system | Myeloid lineage |
| GO_MAST_CELL_ACTIVATION | NIR | immune system | Myeloid lineage |
| GO_REGULATION_OF_LEUKOCYTE_MEDIATED_CYTOTOXICITY | NIR | immune system | Myeloid lineage |
| GO_MYELOID_CELL_DIFFERENTIATION | NIR | immune system | Myeloid lineage |
| GO_MACROPHAGE_ACTIVATION | NIR | immune system | Myeloid lineage |
| GO_REGULATION_OF_LEUKOCYTE_MEDIATED_IMMUNITY | NIR | immune system | Myeloid lineage |
| GO_NEGATIVE_REGULATION_OF_LYMPHOCYTE_MEDIATED_IMMUNITY | NIR | immune system | Myeloid lineage |
| GO_MAST_CELL_GRANULE | NIR | immune system | Myeloid lineage |
| GO_DENDRITIC_CELL_DIFFERENTIATION | NIR | immune system | Myeloid lineage |
| GO_MYELOID_LEUKOCYTE_DIFFERENTIATION | NIR | immune system | Myeloid lineage |
| GO_MACROPHAGE_DIFFERENTIATION | NIR | immune system | Myeloid lineage |
| GO_LEUKOCYTE_ACTIVATION | NIR | immune system | Myeloid lineage |
| GO_POSITIVE_REGULATION_OF_LEUKOCYTE_PROLIFERATION | NIR | immune system | Myeloid lineage |
| GO_LYSOSOME_LOCALIZATION | NIR | immune system | Myeloid lineage |
| GO_NEGATIVE_REGULATION_OF_LEUKOCYTE_MEDIATED_IMMUNITY | NIR | immune system | Myeloid lineage |
| GO_LYMPHOCYTE_ACTIVATION_INVOLVED_IN_IMMUNE_RESPONSE | NIR | immune system | Myeloid lineage |
| GO_POSITIVE_REGULATION_OF_LEUKOCYTE_DEGRANULATION | NIR | immune system | Myeloid lineage |
| GO_REGULATION_OF_MAST_CELL_DEGRANULATION | NIR | immune system | Myeloid lineage |
| GO_POSITIVE_REGULATION_OF_LEUKOCYTE_DIFFERENTIATION | NIR | immune system | Myeloid lineage |
| GO_REGULATION_OF_LYMPHOCYTE_MEDIATED_IMMUNITY | NIR | immune system | Myeloid lineage |
| GO_GRANULOCYTE_ACTIVATION | NIR | immune system | Myeloid lineage |
| GO_GRANULOCYTE_DIFFERENTIATION | NIR | immune system | Myeloid lineage |
| GO_MYELOID_DENDRITIC_CELL_ACTIVATION | NIR | immune system | Myeloid lineage |
| GO_LEUKOCYTE_MEDIATED_IMMUNITY | NIR | immune system | Myeloid lineage |
| GO_REGULATION_OF_LEUKOCYTE_PROLIFERATION | NIR | immune system | Myeloid lineage |
| GO_TOLL_LIKE_RECEPTOR_SIGNALING_PATHWAY | NIR | immune system | signal transduction |
| GO_CHEMOKINE_RECEPTOR_BINDING | NIR | immune system | signal transduction |
| GO_POSITIVE_REGULATION_OF_TOLL_LIKE_RECEPTOR_SIGNALING_PATHWAY | NIR | immune system | signal transduction |
| GO_REGULATION_OF_T_CELL_RECEPTOR_SIGNALING_PATHWAY | NIR | immune system | signal transduction |
| GO_IMMUNE_RESPONSE_REGULATING_CELL_SURFACE_RECEPTOR_SIGNALING_PATHWAY | NIR | immune system | signal transduction |
| GO_POSITIVE_REGULATION_OF_ANTIGEN_PROCESSING_AND_PRESENTATION | NIR | immune system | signal transduction |
| GO_PATTERN_RECOGNITION_RECEPTOR_SIGNALING_PATHWAY | NIR | immune system | signal transduction |
| GO_CYTOKINE_BINDING | NIR | immune system | signal transduction |
| GO_DIVALENT_INORGANIC_CATION_HOMEOSTASIS | NIR | immune system | signal transduction |
| GO_MHC_PROTEIN_COMPLEX | NIR | immune system | signal transduction |
| GO_SIDE_OF_MEMBRANE | NIR | immune system | signal transduction |
| GO_CELL_CELL_RECOGNITION | NIR | immune system | signal transduction |
| GO_REGULATION_OF_ANTIGEN_RECEPTOR_MEDIATED_SIGNALING_PATHWAY | NIR | immune system | signal transduction |
| GO_REGULATION_OF_ANTIGEN_PROCESSING_AND_PRESENTATION | NIR | immune system | signal transduction |
| GO_IMMUNOLOGICAL_SYNAPSE | NIR | immune system | signal transduction |
| GO_TUMOR_NECROSIS_FACTOR_MEDIATED_SIGNALING_PATHWAY | NIR | immune system | signal transduction |
| GO_REGULATION_OF_ERK1_AND_ERK2_CASCADE | NIR | immune system | signal transduction |
| GO_REGULATION_OF_CYTOSOLIC_CALCIUM_ION_CONCENTRATION | NIR | immune system | signal transduction |
| GO_CYTOKINE_MEDIATED_SIGNALING_PATHWAY | NIR | immune system | signal transduction |
| GO_CLATHRIN_COATED_ENDOCYTIC_VESICLE | NIR | immune system | signal transduction |
| GO_LYMPHOCYTE_COSTIMULATION | NIR | immune system | signal transduction |
| GO_B_CELL_RECEPTOR_SIGNALING_PATHWAY | NIR | immune system | signal transduction |
| GO_MHC_CLASS_II_PROTEIN_COMPLEX_BINDING | NIR | immune system | signal transduction |
| GO_CYTOKINE_RECEPTOR_ACTIVITY | NIR | immune system | signal transduction |
| GO_CLATHRIN_COATED_ENDOCYTIC_VESICLE_MEMBRANE | NIR | immune system | signal transduction |
| GO_CHEMOKINE_MEDIATED_SIGNALING_PATHWAY | NIR | immune system | signal transduction |
| GO_CCR_CHEMOKINE_RECEPTOR_BINDING | NIR | immune system | signal transduction |
| GO_PHAGOCYTOSIS_RECOGNITION | NIR | immune system | signal transduction |
| GO_NEGATIVE_REGULATION_OF_ANTIGEN_RECEPTOR_MEDIATED_SIGNALING_PATHWAY | NIR | immune system | signal transduction |
| GO_CYTOKINE_RECEPTOR_BINDING | NIR | immune system | signal transduction |
| GO_PEPTIDYL_TYROSINE_AUTOPHOSPHORYLATION | NIR | immune system | signal transduction |
| GO_T_CELL_RECEPTOR_SIGNALING_PATHWAY | NIR | immune system | signal transduction |
| GO_MHC_PROTEIN_COMPLEX_BINDING | NIR | immune system | signal transduction |
| GO_INTERFERON_GAMMA_MEDIATED_SIGNALING_PATHWAY | NIR | immune system | signal transduction |
| GO_ANTIGEN_RECEPTOR_MEDIATED_SIGNALING_PATHWAY | NIR | immune system | signal transduction |
| GO_EXTERNAL_SIDE_OF_PLASMA_MEMBRANE | NIR | immune system | signal transduction |
| GO_POSITIVE_REGULATION_OF_ERK1_AND_ERK2_CASCADE | NIR | immune system | signal transduction |
| GO_ANTIGEN_BINDING | NIR | immune system | signal transduction |
| GO_MHC_CLASS_II_PROTEIN_COMPLEX | NIR | immune system | signal transduction |
| GO_PLASMA_MEMBRANE_RECEPTOR_COMPLEX | NIR | immune system | signal transduction |
| GO_CELL_RECOGNITION | NIR | immune system | signal transduction |
| GO_TUMOR_NECROSIS_FACTOR_RECEPTOR_BINDING | NIR | immune system | signal transduction |
| GO_CHEMOKINE_BINDING | NIR | immune system | signal transduction |
| GO_AROMATIC_AMINO_ACID_FAMILY_METABOLIC_PROCESS | NIR | metabolism | amino-acid |
| GO_AMINE_CATABOLIC_PROCESS | NIR | metabolism | amino-acid |
| GO_INDOLALKYLAMINE_METABOLIC_PROCESS | NIR | metabolism | amino-acid |
| GO_CELLULAR_MODIFIED_AMINO_ACID_METABOLIC_PROCESS | NIR | metabolism | amino-acid |
| GO_GLUTATHIONE_DERIVATIVE_METABOLIC_PROCESS | NIR | metabolism | amino-acid |
| GO_AROMATIC_AMINO_ACID_FAMILY_CATABOLIC_PROCESS | NIR | metabolism | amino-acid |
| GO_GLUTATHIONE_TRANSFERASE_ACTIVITY | NIR | metabolism | amino-acid |
| GO_CELLULAR_BIOGENIC_AMINE_CATABOLIC_PROCESS | NIR | metabolism | amino-acid |
| GO_AMINE_METABOLIC_PROCESS | NIR | metabolism | amino-acid |
| GO_GLUTATHIONE_DERIVATIVE_BIOSYNTHETIC_PROCESS | NIR | metabolism | amino-acid |
| GO_BENZENE_CONTAINING_COMPOUND_METABOLIC_PROCESS | NIR | metabolism | amino-acid |
| GO_SMALL_MOLECULE_BIOSYNTHETIC_PROCESS | NIR | metabolism | amino-acid |
| GO_ARACHIDONIC_ACID_METABOLIC_PROCESS | NIR | metabolism | amino-acid |
| GO_CELLULAR_AMINO_ACID_BIOSYNTHETIC_PROCESS | NIR | metabolism | amino-acid |
| [illegible]_REGULATION_OF_INTRINSIC_APOPTOTIC_SIGNALING_PATHWAY_IN_RESPONSE_TO_DNA_DAMA | NIR | metabolism | apoptosis |
| GO_POSITIVE_REGULATION_OF_INTRINSIC_APOPTOTIC_SIGNALING_PATHWAY | NIR | metabolism | apoptosis |
| [illegible]GATIVE_REGULATION_OF_INTRINSIC_APOPTOTIC_SIGNALING_PATHWAY_IN_RESPONSE_TO_DNA_[illegible] | NIR | metabolism | apoptosis |
| GO_REGULATION_OF_INTRINSIC_APOPTOTIC_SIGNALING_PATHWAY | NIR | metabolism | apoptosis |

| | | | |
|---|---|---|---|
| GO_REGULATION_OF_PHAGOCYTOSIS | NIR | metabolism | cytosis |
| GO_POSITIVE_REGULATION_OF_ENDOCYTOSIS | NIR | metabolism | cytosis |
| GO_POSITIVE_REGULATION_OF_PHAGOCYTOSIS | NIR | metabolism | cytosis |
| GO_MEMBRANE_INVAGINATION | NIR | metabolism | cytosis |
| GO_PHAGOCYTOSIS_ENGULFMENT | NIR | metabolism | cytosis |
| GO_ACTIVATION_OF_CYSTEINE_TYPE_ENDOPEPTIDASE_ACTIVITY | NIR | metabolism | decomposition |
| GO_METALLOEXOPEPTIDASE_ACTIVITY | NIR | metabolism | decomposition |
| GO_SERINE_TYPE_EXOPEPTIDASE_ACTIVITY | NIR | metabolism | decomposition |
| GO_POSITIVE_REGULATION_OF_PEPTIDASE_ACTIVITY | NIR | metabolism | decomposition |
| _REGULATION_OF_CYSTEINE_TYPE_ENDOPEPTIDASE_ACTIVITY_INVOLVED_IN_APOPTOTIC_SIGNALI | NIR | metabolism | decomposition |
| GO_CARBOXYPEPTIDASE_ACTIVITY | NIR | metabolism | decomposition |
| GO_NEGATIVE_REGULATION_OF_PROTEOLYSIS | NIR | metabolism | decomposition |
| ;ATIVE_REGULATION_OF_EXTRINSIC_APOPTOTIC_SIGNALING_PATHWAY_VIA_DEATH_DOMAIN_RE( | NIR | metabolism | decomposition |
| GO_PEPTIDASE_INHIBITOR_ACTIVITY | NIR | metabolism | decomposition |
| GO_PEPTIDASE_ACTIVATOR_ACTIVITY_INVOLVED_IN_APOPTOTIC_PROCESS | NIR | metabolism | decomposition |
| GO_CALCIUM_DEPENDENT_CYSTEINE_TYPE_ENDOPEPTIDASE_ACTIVITY | NIR | metabolism | decomposition |
| ULATION_OF_CYSTEINE_TYPE_ENDOPEPTIDASE_ACTIVITY_INVOLVED_IN_APOPTOTIC_SIGNALING_P | NIR | metabolism | decomposition |
| GO_CYSTEINE_TYPE_ENDOPEPTIDASE_INHIBITOR_ACTIVITY | NIR | metabolism | decomposition |
| GO_EXOPEPTIDASE_ACTIVITY | NIR | metabolism | decomposition |
| GO_CYSTEINE_TYPE_ENDOPEPTIDASE_ACTIVITY | NIR | metabolism | decomposition |
| GO_REGULATION_OF_PEPTIDASE_ACTIVITY | NIR | metabolism | decomposition |
| GO_NEGATIVE_REGULATION_OF_HYDROLASE_ACTIVITY | NIR | metabolism | decomposition |
| GO_PEPTIDASE_REGULATOR_ACTIVITY | NIR | metabolism | decomposition |
| GO_METALLOCARBOXYPEPTIDASE_ACTIVITY | NIR | metabolism | decomposition |
| GO_CYSTEINE_TYPE_ENDOPEPTIDASE_INHIBITOR_ACTIVITY_INVOLVED_IN_APOPTOTIC_PROCESS | NIR | metabolism | decomposition |
| GO_SERINE_TYPE_ENDOPEPTIDASE_INHIBITOR_ACTIVITY | NIR | metabolism | decomposition |
| GO_CYSTEINE_TYPE_ENDOPEPTIDASE_REGULATOR_ACTIVITY_INVOLVED_IN_APOPTOTIC_PROCESS | NIR | metabolism | decomposition |
| GO_ENDOPEPTIDASE_ACTIVITY | NIR | metabolism | decomposition |
| GO_PROTEIN_MATURATION | NIR | metabolism | decomposition |
| GO_PEPTIDASE_ACTIVATOR_ACTIVITY | NIR | metabolism | decomposition |
| GO_ZYMOGEN_ACTIVATION | NIR | metabolism | decomposition |
| GO_SERINE_HYDROLASE_ACTIVITY | NIR | metabolism | decomposition |
| GO_NEGATIVE_REGULATION_OF_PEPTIDASE_ACTIVITY | NIR | metabolism | decomposition |
| GO_CARBON_CARBON_LYASE_ACTIVITY | NIR | metabolism | decomposition |
| GO_HYDRO_LYASE_ACTIVITY | NIR | metabolism | decomposition |
| GO_NUCLEOSIDE_BISPHOSPHATE_METABOLIC_PROCESS | NIR | metabolism | decomposition |
| GO_HYDROLASE_ACTIVITY_ACTING_ON_CARBON_NITROGEN_BUT_NOT_PEPTIDE_BONDS | NIR | metabolism | decomposition |
| GO_RIBONUCLEOSIDE_BISPHOSPHATE_METABOLIC_PROCESS | NIR | metabolism | decomposition |
| GO_PURINE_RIBONUCLEOSIDE_BISPHOSPHATE_METABOLIC_PROCESS | NIR | metabolism | decomposition |
| GO_PURINE_NUCLEOSIDE_BISPHOSPHATE_METABOLIC_PROCESS | NIR | metabolism | decomposition |
| GO_LYASE_ACTIVITY | NIR | metabolism | decomposition |
| IYDROLASE_ACTIVITY_ACTING_ON_CARBON_NITROGEN_BUT_NOT_PEPTIDE_BONDS_IN_LINEAR_AI | NIR | metabolism | decomposition |
| GO_DEAMINASE_ACTIVITY | NIR | metabolism | decomposition |
| GO_PYRIDOXAL_PHOSPHATE_BINDING | NIR | metabolism | decomposition |
| 'DROLASE_ACTIVITY_ACTING_ON_CARBON_NITROGEN_BUT_NOT_PEPTIDE_BONDS_IN_CYCLIC_AM | NIR | metabolism | decomposition |
| GO_PIR_CHAIN_FATTY_ACID_METABOLIC_PROCESS | NIR | metabolism | fatty-acid |
| GO_LIGASE_ACTIVITY_FORMING_CARBON_SULFUR_BONDS | NIR | metabolism | fatty-acid |
| GO_FATTY_ACID_BIOSYNTHETIC_PROCESS | NIR | metabolism | fatty-acid |
| GO_PROSTAGLANDIN_METABOLIC_PROCESS | NIR | metabolism | fatty-acid |
| GO_PROSTAGLANDIN_BIOSYNTHETIC_PROCESS | NIR | metabolism | fatty-acid |
| GO_ICOSANOID_BIOSYNTHETIC_PROCESS | NIR | metabolism | fatty-acid |
| GO_FATTY_ACID_DERIVATIVE_BIOSYNTHETIC_PROCESS | NIR | metabolism | fatty-acid |
| GO_LEUKOTRIENE_METABOLIC_PROCESS | NIR | metabolism | fatty-acid |
| GO_FATTY_ACID_LIGASE_ACTIVITY | NIR | metabolism | fatty-acid |
| GO_UNSATURATED_FATTY_ACID_METABOLIC_PROCESS | NIR | metabolism | fatty-acid |
| GO_ICOSANOID_METABOLIC_PROCESS | NIR | metabolism | fatty-acid |
| GO_ACID_THIOL_LIGASE_ACTIVITY | NIR | metabolism | fatty-acid |
| GO_PROSTANOID_METABOLIC_PROCESS | NIR | metabolism | fatty-acid |
| GO_CARBOXYLIC_ACID_BIOSYNTHETIC_PROCESS | NIR | metabolism | fatty-acid |
| GO_ORGANIC_ACID_BIOSYNTHETIC_PROCESS | NIR | metabolism | fatty-acid |
| GO_FATTY_ACID_DERIVATIVE_METABOLIC_PROCESS | NIR | metabolism | fatty-acid |
| GO_LEUKOTRIENE_BIOSYNTHETIC_PROCESS | NIR | metabolism | fatty-acid |
| GO_PROSTANOID_BIOSYNTHETIC_PROCESS | NIR | metabolism | fatty-acid |
| GO_UNSATURATED_FATTY_ACID_BIOSYNTHETIC_PROCESS | NIR | metabolism | fatty-acid |
| GO_DNA_DEALKYLATION | PIR | metabolism | organonitrogen |
| GO_DNA_DEMETHYLATION | PIR | metabolism | organonitrogen |
| GO_ORGANONITROGEN_COMPOUND_CATABOLIC_PROCESS | NIR | metabolism | organonitrogen |
| GO_CELLULAR_METABOLIC_COMPOUND_SALVAGE | NIR | metabolism | organonitrogen |
| GO_GLYCOSYL_COMPOUND_CATABOLIC_PROCESS | NIR | metabolism | organonitrogen |
| GO_COFACTOR_CATABOLIC_PROCESS | NIR | metabolism | organonitrogen |
| GO_NUCLEOSIDE_PHOSPHATE_CATABOLIC_PROCESS | NIR | metabolism | organonitrogen |
| GO_CARBOHYDRATE_DERIVATIVE_CATABOLIC_PROCESS | NIR | metabolism | organonitrogen |
| GO_PURINE_CONTAINING_COMPOUND_SALVAGE | NIR | metabolism | organonitrogen |
| GO_PURINE_CONTAINING_COMPOUND_CATABOLIC_PROCESS | NIR | metabolism | organonitrogen |
| GO_RIBONUCLEOSIDE_CATABOLIC_PROCESS | NIR | metabolism | organonitrogen |
| GO_POSITIVE_REGULATION_OF_REACTIVE_OXYGEN_SPECIES_METABOLIC_PROCESS | NIR | metabolism | oxidoreduction |
| GO_MONOOXYGENASE_ACTIVITY | NIR | metabolism | oxidoreduction |
| GO_IRON_ION_BINDING | NIR | metabolism | oxidoreduction |
| GO_REGULATION_OF_RESPONSE_TO_OXIDATIVE_STRESS | NIR | metabolism | oxidoreduction |
| GO_REGULATION_OF_REACTIVE_OXYGEN_SPECIES_METABOLIC_PROCESS | NIR | metabolism | oxidoreduction |
| NORS_WITH_INCORPORATION_OR_REDUCTION_OF_MOLECULAR_OXYGEN_NAD_P_H_AS_ONE_DC | NIR | metabolism | oxidoreduction |
| GO_HYDROGEN_PEROXIDE_METABOLIC_PROCESS | NIR | metabolism | oxidoreduction |
| GO_TETRAPYRROLE_BINDING | NIR | metabolism | oxidoreduction |
| GO_OXIDOREDUCTASE_ACTIVITY_ACTING_ON_PEROXIDE_AS_ACCEPTOR | NIR | metabolism | oxidoreduction |
| GO_REGULATION_OF_NITRIC_OXIDE_BIOSYNTHETIC_PROCESS | NIR | metabolism | oxidoreduction |
| GO_RESPONSE_TO_HYDROPEROXIDE | NIR | metabolism | oxidoreduction |
| )OREDUCTASE_ACTIVITY_ACTING_ON_SINGLE_DONORS_WITH_INCORPORATION_OF_MOLECULAR_ | NIR | metabolism | oxidoreduction |

| | | | |
|---|---|---|---|
| GO_ANTIOXIDANT_ACTIVITY | NIR | metabolism | oxidoreduction |
| GO_TERPENOID_METABOLIC_PROCESS | NIR | metabolism | sterols/alcohols |
| GO_ALCOHOL_DEHYDROGENASE_NADP_ACTIVITY | NIR | metabolism | sterols/alcohols |
| GO_ORGANIC_HYDROXY_COMPOUND_TRANSPORT | NIR | metabolism | sterols/alcohols |
| GO_PHOSPHATIDYLCHOLINE_METABOLIC_PROCESS | NIR | metabolism | sterols/alcohols |
| GO_REGULATION_OF_STEROID_METABOLIC_PROCESS | NIR | metabolism | sterols/alcohols |
| GO_REGULATION_OF_HORMONE_LEVELS | NIR | metabolism | sterols/alcohols |
| GO_ALDO_KETO_REDUCTASE_NADP_ACTIVITY | NIR | metabolism | sterols/alcohols |
| GO_REGULATION_OF_ALCOHOL_BIOSYNTHETIC_PROCESS | NIR | metabolism | sterols/alcohols |
| GO_PRIMARY_ALCOHOL_METABOLIC_PROCESS | NIR | metabolism | sterols/alcohols |
| GO_HORMONE_METABOLIC_PROCESS | NIR | metabolism | sterols/alcohols |
| GO_ALCOHOL_BINDING | NIR | metabolism | sterols/alcohols |
| GO_POSITIVE_REGULATION_OF_FATTY_ACID_BIOSYNTHETIC_PROCESS | NIR | metabolism | sterols/alcohols |
| GO_DIGESTIVE_SYSTEM_PROCESS | NIR | metabolism | sterols/alcohols |
| GO_REGULATION_OF_STEROID_BIOSYNTHETIC_PROCESS | NIR | metabolism | sterols/alcohols |
| GO_PHOSPHOLIPASE_C_ACTIVITY | NIR | metabolism | sterols/alcohols |
| GO_REGULATION_OF_LIPID_BIOSYNTHETIC_PROCESS | NIR | metabolism | sterols/alcohols |
| GO_LIPID_DIGESTION | NIR | metabolism | sterols/alcohols |
| GO_VITAMIN_METABOLIC_PROCESS | NIR | metabolism | sterols/alcohols |
| GO_CELLULAR_HORMONE_METABOLIC_PROCESS | NIR | metabolism | sterols/alcohols |
| GO_ALCOHOL_BIOSYNTHETIC_PROCESS | NIR | metabolism | sterols/alcohols |
| GO_ORGANIC_HYDROXY_COMPOUND_TRANSMEMBRANE_TRANSPORTER_ACTIVITY | NIR | metabolism | sterols/alcohols |
| GO_RETINOL_DEHYDROGENASE_ACTIVITY | NIR | metabolism | sterols/alcohols |
| GO_ETHANOLAMINE_CONTAINING_COMPOUND_METABOLIC_PROCESS | NIR | metabolism | sterols/alcohols |
| GO_ISOPRENOID_METABOLIC_PROCESS | NIR | metabolism | sterols/alcohols |
| GO_FAT_SOLUBLE_VITAMIN_METABOLIC_PROCESS | NIR | metabolism | sterols/alcohols |
| GO_SECONDARY_METABOLIC_PROCESS | NIR | metabolism | sterols/alcohols |
| GO_REGULATION_OF_PLASMA_LIPOPROTEIN_PARTICLE_LEVELS | NIR | metabolism | sterols/alcohols |
| GO_AZOLE_TRANSPORT | NIR | metabolism | sterols/alcohols |
| GO_PHOSPHOLIPASE_A2_ACTIVITY | NIR | metabolism | sterols/alcohols |
| GO_PHOSPHORIC_DIESTER_HYDROLASE_ACTIVITY | NIR | metabolism | sterols/alcohols |
| GO_STEROID_BIOSYNTHETIC_PROCESS | NIR | metabolism | sterols/alcohols |
| GO_C21_STEROID_HORMONE_METABOLIC_PROCESS | NIR | metabolism | sterols/alcohols |
| GO_STEROID_DEHYDROGENASE_ACTIVITY | NIR | metabolism | sterols/alcohols |
| GO_ALCOHOL_METABOLIC_PROCESS | NIR | metabolism | sterols/alcohols |
| GO_POSITIVE_REGULATION_OF_LIPID_BIOSYNTHETIC_PROCESS | NIR | metabolism | sterols/alcohols |
| GO_OXIDOREDUCTASE_ACTIVITY_ACTING_ON_CH_OH_GROUP_OF_DONORS | NIR | metabolism | sterols/alcohols |
| GO_ORGANIC_HYDROXY_COMPOUND_BIOSYNTHETIC_PROCESS | NIR | metabolism | sterols/alcohols |
| GO_CARBOXYLIC_ESTER_HYDROLASE_ACTIVITY | NIR | metabolism | sterols/alcohols |
| GO_MONOAMINE_TRANSPORT | NIR | metabolism | sterols/alcohols |
| GO_STEROL_TRANSPORT | NIR | metabolism | sterols/alcohols |
| GO_HIGH_DENSITY_LIPOPROTEIN_PARTICLE_REMODELING | NIR | metabolism | sterols/alcohols |
| GO_DIGESTION | NIR | metabolism | sterols/alcohols |
| GO_GLYCOSIDE_METABOLIC_PROCESS | NIR | metabolism | sterols/alcohols |
| GO_PHOSPHOLIPASE_ACTIVITY | NIR | metabolism | sterols/alcohols |
| GO_STEROID_BINDING | NIR | metabolism | sterols/alcohols |
| GO_REGULATION_OF_CHOLESTEROL_METABOLIC_PROCESS | NIR | metabolism | sterols/alcohols |
| GO_AMMONIUM_TRANSPORT | NIR | metabolism | sterols/alcohols |
| GO_LIPID_LOCALIZATION | NIR | metabolism | sterols/alcohols |
| GO_ORGANIC_HYDROXY_COMPOUND_METABOLIC_PROCESS | NIR | metabolism | sterols/alcohols |
| GO_POSITIVE_REGULATION_OF_STEROID_METABOLIC_PROCESS | NIR | metabolism | sterols/alcohols |
| GO_LIPASE_ACTIVITY | NIR | metabolism | sterols/alcohols |
| GO_STEROID_METABOLIC_PROCESS | NIR | metabolism | sterols/alcohols |
| GO_NEGATIVE_REGULATION_OF_LIPID_METABOLIC_PROCESS | NIR | metabolism | sterols/alcohols |
| GO_PROTEIN_LIPID_COMPLEX_SUBUNIT_ORGANIZATION | NIR | metabolism | sterols/alcohols |
| GO_RETINOL_METABOLIC_PROCESS | NIR | metabolism | sterols/alcohols |
| GO_RESPONSE_TO_TOXIC_SUBSTANCE | NIR | metabolism | toxicity |
| GO_PROTEIN_DNA_COMPLEX | PIR | Nucleus activity | |
| GO_NON_RECOMBINATIONAL_REPAIR | PIR | Nucleus activity | |
| GO_NEGATIVE_REGULATION_OF_HEMATOPOIETIC_PROGENITOR_CELL_DIFFERENTIATION | PIR | Nucleus activity | |
| GO_BETA_CATENIN_TCF_COMPLEX_ASSEMBLY | PIR | Nucleus activity | |
| GO_GENE_SILENCING_BY_RNA | PIR | Nucleus activity | |
| GO_DNA_PACKAGING | PIR | Nucleus activity | |
| GO_CHROMATIN_SILENCING_AT_RDNA | PIR | Nucleus activity | |
| GO_CHROMATIN_ASSEMBLY_OR_DISASSEMBLY | PIR | Nucleus activity | |
| GO_NUCLEAR_CHROMOSOME_TELOMERIC_REGION | PIR | Nucleus activity | |
| GO_TELOMERE_CAPPING | PIR | Nucleus activity | |
| GO_PROTEIN_HETEROTETRAMERIZATION | PIR | Nucleus activity | |
| GO_REGULATION_OF_HEMATOPOIETIC_PROGENITOR_CELL_DIFFERENTIATION | PIR | Nucleus activity | |
| GO_REGULATION_OF_GENE_EXPRESSION_EPIGENETIC | PIR | Nucleus activity | |
| GO_POSITIVE_REGULATION_OF_GENE_EXPRESSION_EPIGENETIC | PIR | Nucleus activity | |
| GO_NEGATIVE_REGULATION_OF_GENE_EXPRESSION_EPIGENETIC | PIR | Nucleus activity | |
| GO_DNA_PACKAGING_COMPLEX | PIR | Nucleus activity | |
| GO_DNA_REPLICATION_DEPENDENT_NUCLEOSOME_ASSEMBLY | PIR | Nucleus activity | |
| GO_TELOMERE_ORGANIZATION | PIR | Nucleus activity | |
| GO_REGULATION_OF_GENE_SILENCING | PIR | Nucleus activity | |
| GO_GENE_SILENCING | PIR | Nucleus activity | |
| GO_DNA_REPLICATION_DEPENDENT_NUCLEOSOME_ORGANIZATION | PIR | Nucleus activity | |
| GO_PYRIDINE_CONTAINING_COMPOUND_BIOSYNTHETIC_PROCESS | NIR | Nucleus activity | |
| GO_NICOTINAMIDE_NUCLEOTIDE_BIOSYNTHETIC_PROCESS | NIR | Nucleus activity | |
| GO_PYRIDINE_NUCLEOTIDE_BIOSYNTHETIC_PROCESS | NIR | Nucleus activity | |
| GO_POSITIVE_REGULATION_OF_EPIDERMIS_DEVELOPMENT | NIR | organogenesis | epidermis development |
| GO_REGULATION_OF_HAIR_FOLLICLE_DEVELOPMENT | NIR | organogenesis | epidermis development |
| GO_MUSCULOSKELETAL_MOVEMENT | NIR | physiological function | muslce movement |
| GO_SKELETAL_MUSCLE_CONTRACTION | NIR | physiological function | muslce movement |
| GO_MULTICELLULAR_ORGANISMAL_MOVEMENT | NIR | physiological function | muslce movement |
| GO_MUSCLE_FILAMENT_SLIDING | NIR | physiological function | muslce movement |

| | | | |
|---|---|---|---|
| GO_ACTIN_MYOSIN_FILAMENT_SLIDING | NIR | physiological function | muslce movement |
| GO_STRIATED_MUSCLE_CONTRACTION | NIR | physiological function | muslce movement |
| GO_ACTIN_FILAMENT_POLYMERIZATION | NIR | organogenesis | muscle development |
| GO_ACTIN_POLYMERIZATION_OR_DEPOLYMERIZATION | NIR | organogenesis | muscle development |
| GO_CEREBRAL_CORTEX_RADIAL_GLIA_GUIDED_MIGRATION | PIR | organogenesis | neural development |
| GO_CEREBRAL_CORTEX_RADIALLY_ORIENTED_CELL_MIGRATION | PIR | organogenesis | neural development |
| GO_TELENCEPHALON_GLIAL_CELL_MIGRATION | PIR | organogenesis | neural development |
| GO_ORGAN_REGENERATION | NIR | organogenesis | organ regeneration |
| GO_REGENERATION | NIR | organogenesis | organ regeneration |
| GO_LIVER_REGENERATION | NIR | organogenesis | organ regeneration |
| GO_TISSUE_REGENERATION | NIR | organogenesis | organ regeneration |
| GO_PLATELET_ALPHA_GRANULE_LUMEN | NIR | organogenesis | Secretory granules |
| GO_PLATELET_DEGRANULATION | NIR | organogenesis | Secretory granules |
| GO_PRIMARY_LYSOSOME | NIR | organogenesis | Secretory granules |
| GO_SECRETORY_GRANULE | NIR | organogenesis | Secretory granules |
| GO_VESICLE_LUMEN | NIR | organogenesis | Secretory granules |
| GO_SECRETORY_GRANULE_LUMEN | NIR | organogenesis | Secretory granules |
| GO_PLATELET_ALPHA_GRANULE | NIR | organogenesis | Secretory granules |
| GO_SECRETORY_GRANULE_MEMBRANE | NIR | organogenesis | Secretory granules |
| GO_MAINTENANCE_OF_GASTROINTESTINAL_EPITHELIUM | NIR | organogenesis | Tissue homeostasis |
| GO_EPITHELIAL_STRUCTURE_MAINTENANCE | NIR | organogenesis | Tissue homeostasis |
| GO_TISSUE_HOMEOSTASIS | NIR | organogenesis | Tissue homeostasis |
| GO_RETINA_HOMEOSTASIS | NIR | organogenesis | Tissue homeostasis |
| GO_MULTICELLULAR_ORGANISMAL_HOMEOSTASIS | NIR | organogenesis | Tissue homeostasis |
| GO_CELL_MATURATION | NIR | physiological function | fertility |
| GO_SPERM_CAPACITATION | NIR | physiological function | fertility |
| GO_SPERM_PART | NIR | physiological function | fertility |
| GO_NEGATIVE_REGULATION_OF_STAT_CASCADE | PIR | signal transduction | |
| GO_NEGATIVE_REGULATION_OF_JAK_STAT_CASCADE | PIR | signal transduction | |
| GO_POSITIVE_REGULATION_OF_TYROSINE_PHOSPHORYLATION_OF_STAT3_PROTEIN | NIR | signal transduction | |
| GO_POSITIVE_REGULATION_OF_STAT_CASCADE | NIR | signal transduction | |
| GO_POSITIVE_REGULATION_OF_JAK_STAT_CASCADE | NIR | signal transduction | |
| GO_REGULATION_OF_PEPTIDYL_TYROSINE_PHOSPHORYLATION | NIR | signal transduction | |
| GO_REGULATION_OF_TYROSINE_PHOSPHORYLATION_OF_STAT_PROTEIN | NIR | signal transduction | |
| GO_POSITIVE_REGULATION_OF_PEPTIDYL_TYROSINE_PHOSPHORYLATION | NIR | signal transduction | |
| GO_POSITIVE_REGULATION_OF_SODIUM_ION_TRANSMEMBRANE_TRANSPORT | NIR | transport | |
| GO_POSITIVE_REGULATION_OF_TRANSMEMBRANE_TRANSPORT | NIR | transport | |
| GO_POSITIVE_REGULATION_OF_CALCIUM_MEDIATED_SIGNALING | NIR | transport | |
| GO_POSITIVE_REGULATION_OF_CALCIUM_ION_TRANSPORT | NIR | transport | |
| GO_POSITIVE_REGULATION_OF_CATION_TRANSMEMBRANE_TRANSPORT | NIR | transport | |
| GO_REGULATION_OF_SYNAPTIC_TRANSMISSION_DOPAMINERGIC | NIR | transport | |
| GO_POSITIVE_REGULATION_OF_ION_TRANSPORT | NIR | transport | |
| GO_POSITIVE_REGULATION_OF_CALCIUM_ION_TRANSMEMBRANE_TRANSPORT | NIR | transport | |
| GO_POSITIVE_REGULATION_OF_CALCIUM_ION_TRANSPORT_INTO_CYTOSOL | NIR | transport | |
| GO_BASOLATERAL_PLASMA_MEMBRANE | NIR | transport | |
| GO_POSITIVE_REGULATION_OF_SODIUM_ION_TRANSPORT | NIR | transport | |
| GO_POSITIVE_REGULATION_OF_TRANSPORTER_ACTIVITY | NIR | transport | |
| GO_REGULATION_OF_CALCIUM_MEDIATED_SIGNALING | NIR | transport | |
| GO_CHLORIDE_CHANNEL_REGULATOR_ACTIVITY | NIR | transport | |
| GO_BICARBONATE_TRANSMEMBRANE_TRANSPORTER_ACTIVITY | NIR | transport | |
| GO_BASAL_PLASMA_MEMBRANE | NIR | transport | |
| GO_REGULATION_OF_CALCIUM_ION_TRANSPORT | NIR | transport | |
| GO_POSITIVE_REGULATION_OF_LIPID_TRANSPORT | NIR | transport | |
| GO_CALCIUM_MEDIATED_SIGNALING | NIR | transport | |
| GO_CALCIUM_MEDIATED_SIGNALING_USING_INTRACELLULAR_CALCIUM_SOURCE | NIR | transport | |

**Supplementary Data 2B. The grouping of GO terms in Cohort 2.**

| GO_term | Enrichment in | Type 1 | Type 2 | Type 3 | Type 4 |
|---|---|---|---|---|---|
| GO_CELLULAR_RESPONSE_TO_GLUCAGON_STIMULUS | PIR | Cellular response | hormone | glucagon | |
| GO_RESPONSE_TO_GLUCAGON | PIR | Cellular response | hormone | glucagon | |
| GO_CELLULAR_RESPONSE_TO_PROSTAGLANDIN_E_STIMULUS | NIR | Cellular response | hormone | | |
| GO_RESPONSE_TO_PROSTAGLANDIN | NIR | Cellular response | hormone | | |
| GO_RESPONSE_TO_PROSTAGLANDIN_E | NIR | Cellular response | hormone | | |
| GO_CELLULAR_RESPONSE_TO_CALCIUM_ION | PIR | Cellular response | metal ion | | |
| GO_CELLULAR_RESPONSE_TO_INORGANIC_SUBSTANCE | PIR | Cellular response | metal ion | | |
| GO_RESPONSE_TO_CALCIUM_ION | PIR | Cellular response | metal ion | | |
| GO_RESPONSE_TO_MANGANESE_ION | PIR | Cellular response | metal ion | | |
| GO_RESPONSE_TO_METAL_ION | PIR | Cellular response | metal ion | | |
| GO_CELLULAR_RESPONSE_TO_NUTRIENT | NIR | Cellular response | nutrient | | |
| GO_CELLULAR_RESPONSE_TO_VITAMIN | NIR | Cellular response | nutrient | | |
| GO_CELLULAR_RESPONSE_TO_RADIATION | NIR | Cellular response | radiation | | |
| GO_CELLULAR_RESPONSE_TO_IONIZING_RADIATION | NIR | Cellular response | radiation | | |
| GO_RESPONSE_TO_GAMMA_RADIATION | NIR | Cellular response | radiation | | |
| GO_RESPONSE_TO_IONIZING_RADIATION | NIR | Cellular response | radiation | | |
| GO_RESPONSE_TO_UV | NIR | Cellular response | radiation | | |
| GO_RESPONSE_TO_X_RAY | NIR | Cellular response | radiation | | |
| GO_FEAR_RESPONSE | PIR | Cellular response | stress | | |
| GO_MULTICELLULAR_ORGANISMAL_RESPONSE_TO_STRESS | PIR | Cellular response | stress | | |
| GO_LEUKOCYTE_CELL_CELL_ADHESION | NIR | immune system | cell adhesion | | |
| GO_NEGATIVE_REGULATION_OF_CELL_ADHESION | NIR | immune system | cell adhesion | | |
| GO_NEGATIVE_REGULATION_OF_CELL_CELL_ADHESION | NIR | immune system | cell adhesion | | |
| GO_NEGATIVE_REGULATION_OF_HOMOTYPIC_CELL_CELL_ADHESION | NIR | immune system | cell adhesion | | |
| GO_POSITIVE_REGULATION_OF_CELL_ADHESION | NIR | immune system | cell adhesion | | |
| GO_POSITIVE_REGULATION_OF_CELL_CELL_ADHESION | NIR | immune system | cell adhesion | | |
| GO_REGULATION_OF_CELL_CELL_ADHESION | NIR | immune system | cell adhesion | | |
| GO_REGULATION_OF_HOMOTYPIC_CELL_CELL_ADHESION | NIR | immune system | cell adhesion | | |
| GO_LYMPHOCYTE_APOPTOTIC_PROCESS | NIR | immune system | cell death | apoptosis | |
| GO_NEGATIVE_REGULATION_OF_LEUKOCYTE_APOPTOTIC_PROCESS | NIR | immune system | cell death | apoptosis | |
| GO_NEGATIVE_REGULATION_OF_LYMPHOCYTE_APOPTOTIC_PROCESS | NIR | immune system | cell death | apoptosis | |
| GO_NEGATIVE_REGULATION_OF_T_CELL_APOPTOTIC_PROCESS | NIR | immune system | cell death | apoptosis | |
| GO_POSITIVE_REGULATION_OF_LYMPHOCYTE_APOPTOTIC_PROCESS | NIR | immune system | cell death | apoptosis | |
| GO_REGULATION_OF_B_CELL_APOPTOTIC_PROCESS | NIR | immune system | cell death | apoptosis | |
| GO_REGULATION_OF_LEUKOCYTE_APOPTOTIC_PROCESS | NIR | immune system | cell death | apoptosis | |
| GO_REGULATION_OF_LYMPHOCYTE_APOPTOTIC_PROCESS | NIR | immune system | cell death | apoptosis | |
| GO_REGULATION_OF_T_CELL_APOPTOTIC_PROCESS | NIR | immune system | cell death | apoptosis | |
| GO_NEGATIVE_REGULATION_OF_CELL_KILLING | NIR | immune system | cell death | cell kill | |
| GO_REGULATION_OF_CELL_KILLING | NIR | immune system | cell death | cell kill | |
| GO_LEUKOCYTE_HOMEOSTASIS | NIR | immune system | cell death | HEMOPOIESIS | |
| GO_LYMPHOCYTE_HOMEOSTASIS | NIR | immune system | cell death | HEMOPOIESIS | |
| GO_NEGATIVE_REGULATION_OF_HEMOPOIESIS | NIR | immune system | cell death | HEMOPOIESIS | |
| GO_REGULATION_OF_HEMOPOIESIS | NIR | immune system | cell death | HEMOPOIESIS | |
| GO_DENDRITIC_CELL_CHEMOTAXIS | NIR | immune system | cell migration | | |
| GO_DENDRITIC_CELL_MIGRATION | NIR | immune system | cell migration | | |
| GO_POSITIVE_REGULATION_OF_CELL_ADHESION_MEDIATED_BY_INTEGRIN | NIR | immune system | cell migration | | |
| GO_REGULATION_OF_CELLULAR_EXTRAVASATION | NIR | immune system | cell migration | | |
| GO_CYTOKINE_PRODUCTION | NIR | immune system | cytokine | | |
| GO_CYTOKINE_SECRETION | NIR | immune system | cytokine | | |
| GO_LYMPHOCYTE_COSTIMULATION | NIR | immune system | cytokine | | |
| GO_NEGATIVE_REGULATION_OF_CYTOKINE_PRODUCTION | NIR | immune system | cytokine | | |
| GO_NEGATIVE_REGULATION_OF_INTERFERON_GAMMA_PRODUCTION | NIR | immune system | cytokine | | |
| GO_NEGATIVE_REGULATION_OF_INTERLEUKIN_12_PRODUCTION | NIR | immune system | cytokine | | |
| GATIVE_REGULATION_OF_TUMOR_NECROSIS_FACTOR_SUPERFAMILY_CYTOKINE_PRODU | NIR | immune system | cytokine | | |
| GO_NEGATIVE_REGULATION_OF_TYPE_I_INTERFERON_PRODUCTION | NIR | immune system | cytokine | | |
| GO_POSITIVE_REGULATION_OF_CYTOKINE_BIOSYNTHETIC_PROCESS | NIR | immune system | cytokine | | |
| GO_POSITIVE_REGULATION_OF_INTERFERON_ALPHA_PRODUCTION | NIR | immune system | cytokine | | |
| GO_POSITIVE_REGULATION_OF_INTERFERON_BETA_PRODUCTION | NIR | immune system | cytokine | | |
| GO_POSITIVE_REGULATION_OF_INTERFERON_GAMMA_PRODUCTION | NIR | immune system | cytokine | | |
| GO_POSITIVE_REGULATION_OF_INTERLEUKIN_1_BETA_PRODUCTION | NIR | immune system | cytokine | | |
| GO_POSITIVE_REGULATION_OF_INTERLEUKIN_1_PRODUCTION | NIR | immune system | cytokine | | |
| GO_POSITIVE_REGULATION_OF_INTERLEUKIN_12_PRODUCTION | NIR | immune system | cytokine | | |
| OSITIVE_REGULATION_OF_TUMOR_NECROSIS_FACTOR_SUPERFAMILY_CYTOKINE_PRODU | NIR | immune system | cytokine | | |
| GO_POSITIVE_REGULATION_OF_TYPE_I_INTERFERON_PRODUCTION | NIR | immune system | cytokine | | |
| GO_REGULATION_OF_CYTOKINE_BIOSYNTHETIC_PROCESS | NIR | immune system | cytokine | | |
| GO_REGULATION_OF_INTERFERON_ALPHA_PRODUCTION | NIR | immune system | cytokine | | |
| GO_REGULATION_OF_INTERFERON_BETA_PRODUCTION | NIR | immune system | cytokine | | |
| GO_REGULATION_OF_INTERFERON_GAMMA_PRODUCTION | NIR | immune system | cytokine | | |
| GO_REGULATION_OF_INTERLEUKIN_1_BETA_PRODUCTION | NIR | immune system | cytokine | | |
| GO_REGULATION_OF_INTERLEUKIN_1_PRODUCTION | NIR | immune system | cytokine | | |
| GO_REGULATION_OF_INTERLEUKIN_1_SECRETION | NIR | immune system | cytokine | | |
| GO_REGULATION_OF_INTERLEUKIN_10_PRODUCTION | NIR | immune system | cytokine | | |
| GO_REGULATION_OF_INTERLEUKIN_12_PRODUCTION | NIR | immune system | cytokine | | |
| GO_REGULATION_OF_INTERLEUKIN_2_BIOSYNTHETIC_PROCESS | NIR | immune system | cytokine | | |
| GO_REGULATION_OF_INTERLEUKIN_8_SECRETION | NIR | immune system | cytokine | | |
| O_REGULATION_OF_TUMOR_NECROSIS_FACTOR_SUPERFAMILY_CYTOKINE_PRODUCTIO | NIR | immune system | cytokine | | |
| GO_REGULATION_OF_TYPE_I_INTERFERON_PRODUCTION | NIR | immune system | cytokine | | |
| GO_MHC_PROTEIN_BINDING | NIR | immune system | cytokine | | |
| GO_MHC_CLASS_I_PROTEIN_BINDING | NIR | immune system | cytokine | | |
| ATIVE_REGULATION_OF_SEQUENCE_SPECIFIC_DNA_BINDING_TRANSCRIPTION_FACTOR_ | NIR | immune system | cytokine | | |
| GO_NEGATIVE_REGULATION_OF_NF_KAPPAB_TRANSCRIPTION_FACTOR_ACTIVITY | NIR | immune system | cytokine | | |
| GO_DEFENSE_RESPONSE_TO_OTHER_ORGANISM | NIR | immune system | defense response | | |
| GO_DEFENSE_RESPONSE_TO_VIRUS | NIR | immune system | defense response | | |
| GO_DETECTION_OF_BIOTIC_STIMULUS | NIR | immune system | defense response | | |
| GO_NEGATIVE_REGULATION_OF_DEFENSE_RESPONSE | NIR | immune system | defense response | | |
| GO_NEGATIVE_REGULATION_OF_DEFENSE_RESPONSE_TO_VIRUS | NIR | immune system | defense response | | |
| GO_NEGATIVE_REGULATION_OF_MULTI_ORGANISM_PROCESS | NIR | immune system | defense response | | |
| GO_NEGATIVE_REGULATION_OF_RESPONSE_TO_BIOTIC_STIMULUS | NIR | immune system | defense response | | |
| GO_NEGATIVE_REGULATION_OF_VIRAL_PROCESS | NIR | immune system | defense response | | |
| GO_POSITIVE_REGULATION_OF_MULTI_ORGANISM_PROCESS | NIR | immune system | defense response | | |
| GO_POSITIVE_REGULATION_OF_VIRAL_GENOME_REPLICATION | NIR | immune system | defense response | | |
| GO_POSITIVE_REGULATION_OF_VIRAL_PROCESS | NIR | immune system | defense response | | |
| GO_REGULATION_OF_DEFENSE_RESPONSE_TO_VIRUS | NIR | immune system | defense response | | |
| GO_REGULATION_OF_DEFENSE_RESPONSE_TO_VIRUS_BY_HOST | NIR | immune system | defense response | | |
| GO_REGULATION_OF_DEFENSE_RESPONSE_TO_VIRUS_BY_VIRUS | NIR | immune system | defense response | | |
| GO_REGULATION_OF_MULTI_ORGANISM_PROCESS | NIR | immune system | defense response | | |
| GO_REGULATION_OF_RESPONSE_TO_BIOTIC_STIMULUS | NIR | immune system | defense response | | |
| O_REGULATION_OF_SYMBIOSIS_ENCOMPASSING_MUTUALISM_THROUGH_PARASITISM | NIR | immune system | defense response | | |
| GO_REGULATION_OF_VIRAL_GENOME_REPLICATION | NIR | immune system | defense response | | |
| GO_REGULATION_OF_VIRAL_TRANSCRIPTION | NIR | immune system | defense response | | |
| GO_RESPONSE_TO_VIRUS | NIR | immune system | defense response | | |

| | | | | | |
|---|---|---|---|---|---|
| GO_TRANSPORT_VESICLE | NIR | immune system | defense response | | |
| GO_ACTIVATION_OF_IMMUNE_RESPONSE | NIR | immune system | immune response | | |
| GO_ACTIVATION_OF_INNATE_IMMUNE_RESPONSE | NIR | immune system | immune response | | |
| GO_ADAPTIVE_IMMUNE_RESPONSE | NIR | immune system | immune response | | |
| GO_INNATE_IMMUNE_RESPONSE | NIR | immune system | immune response | | |
| GO_NEGATIVE_REGULATION_OF_ADAPTIVE_IMMUNE_RESPONSE | NIR | immune system | immune response | | |
| GO_NEGATIVE_REGULATION_OF_CELL_ACTIVATION | NIR | immune system | immune response | | |
| GO_NEGATIVE_REGULATION_OF_IMMUNE_EFFECTOR_PROCESS | NIR | immune system | immune response | | |
| GO_NEGATIVE_REGULATION_OF_IMMUNE_RESPONSE | NIR | immune system | immune response | | |
| GO_NEGATIVE_REGULATION_OF_IMMUNE_SYSTEM_PROCESS | NIR | immune system | immune response | | |
| GO_NEGATIVE_REGULATION_OF_INNATE_IMMUNE_RESPONSE | NIR | immune system | immune response | | |
| GO_NEGATIVE_REGULATION_OF_TOLL_LIKE_RECEPTOR_SIGNALING_PATHWAY | NIR | immune system | immune response | | |
| GO_POSITIVE_REGULATION_OF_ADAPTIVE_IMMUNE_RESPONSE | NIR | immune system | immune response | | |
| GO_POSITIVE_REGULATION_OF_CELL_ACTIVATION | NIR | immune system | immune response | | |
| GO_POSITIVE_REGULATION_OF_IMMUNE_RESPONSE | NIR | immune system | immune response | | |
| GO_POSITIVE_REGULATION_OF_INNATE_IMMUNE_RESPONSE | NIR | immune system | immune response | | |
| GO_REGULATION_OF_ADAPTIVE_IMMUNE_RESPONSE | NIR | immune system | immune response | | |
| GO_REGULATION_OF_CELL_ACTIVATION | NIR | immune system | immune response | | |
| GO_REGULATION_OF_IMMUNE_EFFECTOR_PROCESS | NIR | immune system | immune response | | |
| GO_REGULATION_OF_INNATE_IMMUNE_RESPONSE | NIR | immune system | immune response | | |
| GO_REGULATION_OF_TOLERANCE_INDUCTION | NIR | immune system | immune response | | |
| GO_RESPONSE_TO_INTERFERON_ALPHA | NIR | immune system | immune response | | |
| GO_RESPONSE_TO_INTERFERON_BETA | NIR | immune system | immune response | | |
| GO_RESPONSE_TO_TYPE_I_INTERFERON | NIR | immune system | immune response | | |
| ASED_ON_SOMATIC_RECOMBINATION_OF_IMMUNE_RECEPTORS_BUILT_FROM_IMMUI | NIR | immune system | lymphoid lineage | Bcell | immunoglobulin |
| GO_B_CELL_ACTIVATION | NIR | immune system | lymphoid lineage | Bcell | |
| GO_B_CELL_ACTIVATION_INVOLVED_IN_IMMUNE_RESPONSE | NIR | immune system | lymphoid lineage | Bcell | |
| GO_B_CELL_DIFFERENTIATION | NIR | immune system | lymphoid lineage | Bcell | |
| GO_B_CELL_MEDIATED_IMMUNITY | NIR | immune system | lymphoid lineage | Bcell | |
| GO_B_CELL_PROLIFERATION | NIR | immune system | lymphoid lineage | Bcell | |
| GO_IMMUNOGLOBULIN_PRODUCTION | NIR | immune system | lymphoid lineage | Bcell | immunoglobulin |
| JNOGLOBULIN_PRODUCTION_INVOLVED_IN_IMMUNOGLOBULIN_MEDIATED_IMMUNE_ | NIR | immune system | lymphoid lineage | Bcell | immunoglobulin |
| GO_ISOTYPE_SWITCHING | NIR | immune system | lymphoid lineage | Bcell | immunoglobulin |
| GO_NEGATIVE_REGULATION_OF_B_CELL_ACTIVATION | NIR | immune system | lymphoid lineage | Bcell | |
| GO_POSITIVE_REGULATION_OF_B_CELL_ACTIVATION | NIR | immune system | lymphoid lineage | Bcell | |
| GO_PRODUCTION_OF_MOLECULAR_MEDIATOR_OF_IMMUNE_RESPONSE | NIR | immune system | lymphoid lineage | Bcell | |
| GO_REGULATION_OF_B_CELL_ACTIVATION | NIR | immune system | lymphoid lineage | Bcell | |
| GO_REGULATION_OF_B_CELL_DIFFERENTIATION | NIR | immune system | lymphoid lineage | Bcell | |
| GO_REGULATION_OF_ISOTYPE_SWITCHING | NIR | immune system | lymphoid lineage | Bcell | |
| GO_SOMATIC_DIVERSIFICATION_OF_IMMUNOGLOBULINS | NIR | immune system | lymphoid lineage | Bcell | immunoglobulin |
| _SOMATIC_DIVERSIFICATION_OF_IMMUNOGLOBULINS_INVOLVED_IN_IMMUNE_RESPOI | NIR | immune system | lymphoid lineage | Bcell | immunoglobulin |
| GO_SOMATIC_RECOMBINATION_OF_IMMUNOGLOBULIN_GENE_SEGMENTS | NIR | immune system | lymphoid lineage | Bcell | immunoglobulin |
| MATIC_RECOMBINATION_OF_IMMUNOGLOBULIN_GENES_INVOLVED_IN_IMMUNE_RES | NIR | immune system | lymphoid lineage | Bcell | immunoglobulin |
| GO_LYMPHOCYTE_ACTIVATION | NIR | immune system | lymphoid lineage | lymphocyte | |
| GO_LYMPHOCYTE_ACTIVATION_INVOLVED_IN_IMMUNE_RESPONSE | NIR | immune system | lymphoid lineage | lymphocyte | |
| GO_LYMPHOCYTE_DIFFERENTIATION | NIR | immune system | lymphoid lineage | lymphocyte | |
| GO_LYMPHOCYTE_MEDIATED_IMMUNITY | NIR | immune system | lymphoid lineage | lymphocyte | |
| GO_NEGATIVE_REGULATION_OF_LYMPHOCYTE_DIFFERENTIATION | NIR | immune system | lymphoid lineage | lymphocyte | |
| GO_NEGATIVE_REGULATION_OF_LYMPHOCYTE_MEDIATED_IMMUNITY | NIR | immune system | lymphoid lineage | lymphocyte | |
| GO_REGULATION_OF_LYMPHOCYTE_MEDIATED_IMMUNITY | NIR | immune system | lymphoid lineage | lymphocyte | |
| GO_REGULATION_OF_NATURAL_KILLER_CELL_MEDIATED_IMMUNITY | NIR | immune system | lymphoid lineage | NK cell | |
| GO_SOMATIC_CELL_DNA_RECOMBINATION | NIR | immune system | lymphoid lineage | RECOMBINATION | |
| GO_SOMATIC_DIVERSIFICATION_OF_IMMUNE_RECEPTORS | NIR | immune system | lymphoid lineage | RECOMBINATION | |
| IVERSIFICATION_OF_IMMUNE_RECEPTORS_VIA_GERMLINE_RECOMBINATION_WITHIN_A | NIR | immune system | lymphoid lineage | RECOMBINATION | |
| GO_V_D_J_RECOMBINATION | NIR | immune system | lymphoid lineage | RECOMBINATION | |
| GO_ALPHA_BETA_T_CELL_ACTIVATION | NIR | immune system | lymphoid lineage | Tcell | |
| GO_ALPHA_BETA_T_CELL_DIFFERENTIATION | NIR | immune system | lymphoid lineage | Tcell | |
| GO_CD4_POSITIVE_ALPHA_BETA_T_CELL_ACTIVATION | NIR | immune system | lymphoid lineage | Tcell | |
| GO_NEGATIVE_REGULATION_OF_ALPHA_BETA_T_CELL_ACTIVATION | NIR | immune system | lymphoid lineage | Tcell | |
| GO_NEGATIVE_REGULATION_OF_T_CELL_DIFFERENTIATION | NIR | immune system | lymphoid lineage | Tcell | |
| GO_NEGATIVE_REGULATION_OF_T_CELL_PROLIFERATION | NIR | immune system | lymphoid lineage | Tcell | |
| GO_POSITIVE_REGULATION_OF_ALPHA_BETA_T_CELL_ACTIVATION | NIR | immune system | lymphoid lineage | Tcell | |
| GO_POSITIVE_REGULATION_OF_ALPHA_BETA_T_CELL_DIFFERENTIATION | NIR | immune system | lymphoid lineage | Tcell | |
| GO_POSITIVE_T_CELL_SELECTION | NIR | immune system | lymphoid lineage | Tcell | |
| GO_REGULATION_OF_ACTIVATED_T_CELL_PROLIFERATION | NIR | immune system | lymphoid lineage | Tcell | |
| GO_REGULATION_OF_ALPHA_BETA_T_CELL_ACTIVATION | NIR | immune system | lymphoid lineage | Tcell | |
| GO_REGULATION_OF_ALPHA_BETA_T_CELL_DIFFERENTIATION | NIR | immune system | lymphoid lineage | Tcell | |
| GO_REGULATION_OF_ALPHA_BETA_T_CELL_PROLIFERATION | NIR | immune system | lymphoid lineage | Tcell | |
| GO_REGULATION_OF_T_CELL_DIFFERENTIATION | NIR | immune system | lymphoid lineage | Tcell | |
| GO_REGULATION_OF_T_CELL_PROLIFERATION | NIR | immune system | lymphoid lineage | Tcell | |
| GO_T_CELL_ACTIVATION_INVOLVED_IN_IMMUNE_RESPONSE | NIR | immune system | lymphoid lineage | Tcell | |
| GO_T_CELL_DIFFERENTIATION | NIR | immune system | lymphoid lineage | Tcell | |
| GO_T_CELL_DIFFERENTIATION_INVOLVED_IN_IMMUNE_RESPONSE | NIR | immune system | lymphoid lineage | Tcell | |
| GO_T_CELL_MEDIATED_IMMUNITY | NIR | immune system | lymphoid lineage | Tcell | |
| GO_T_CELL_SELECTION | NIR | immune system | lymphoid lineage | Tcell | |
| GO_T_HELPER_1_TYPE_IMMUNE_RESPONSE | NIR | immune system | lymphoid lineage | Tcell | |
| GO_LYMPH_NODE_DEVELOPMENT | NIR | immune system | lymphoid lineage | | |
| GO_POSITIVE_REGULATION_OF_ERYTHROCYTE_DIFFERENTIATION | NIR | immune system | Myeloid lineage | MEP | |
| GO_REGULATION_OF_ERYTHROCYTE_DIFFERENTIATION | NIR | immune system | Myeloid lineage | MEP | |
| GO_DENDRITIC_CELL_DIFFERENTIATION | NIR | immune system | Myeloid lineage | | |
| GO_LEUKOCYTE_ACTIVATION | NIR | immune system | Myeloid lineage | | |
| GO_LEUKOCYTE_DIFFERENTIATION | NIR | immune system | Myeloid lineage | | |
| GO_LEUKOCYTE_PROLIFERATION | NIR | immune system | Myeloid lineage | | |
| GO_MYELOID_CELL_DIFFERENTIATION | NIR | immune system | Myeloid lineage | | |
| GO_MYELOID_DENDRITIC_CELL_DIFFERENTIATION | NIR | immune system | Myeloid lineage | | |
| GO_MYELOID_LEUKOCYTE_DIFFERENTIATION | NIR | immune system | Myeloid lineage | | |
| GO_NEGATIVE_REGULATION_OF_LEUKOCYTE_DIFFERENTIATION | NIR | immune system | Myeloid lineage | | |
| GO_NEGATIVE_REGULATION_OF_LEUKOCYTE_MEDIATED_IMMUNITY | NIR | immune system | Myeloid lineage | | |
| GO_NEGATIVE_REGULATION_OF_LEUKOCYTE_PROLIFERATION | NIR | immune system | Myeloid lineage | | |
| GO_NEGATIVE_REGULATION_OF_MYELOID_CELL_DIFFERENTIATION | NIR | immune system | Myeloid lineage | | |
| GO_NEGATIVE_REGULATION_OF_MYELOID_LEUKOCYTE_DIFFERENTIATION | NIR | immune system | Myeloid lineage | | |
| GO_POSITIVE_REGULATION_OF_LEUKOCYTE_DEGRANULATION | NIR | immune system | Myeloid lineage | | |
| GO_POSITIVE_REGULATION_OF_LEUKOCYTE_MEDIATED_IMMUNITY | NIR | immune system | Myeloid lineage | | |
| GO_POSITIVE_REGULATION_OF_MYELOID_LEUKOCYTE_MEDIATED_IMMUNITY | NIR | immune system | Myeloid lineage | | |
| GO_REGULATION_OF_LEUKOCYTE_DEGRANULATION | NIR | immune system | Myeloid lineage | | |
| GO_REGULATION_OF_LEUKOCYTE_DIFFERENTIATION | NIR | immune system | Myeloid lineage | | |
| GO_REGULATION_OF_LEUKOCYTE_MEDIATED_CYTOTOXICITY | NIR | immune system | Myeloid lineage | | |
| GO_REGULATION_OF_LEUKOCYTE_MEDIATED_IMMUNITY | NIR | immune system | Myeloid lineage | | |
| GO_REGULATION_OF_LEUKOCYTE_PROLIFERATION | NIR | immune system | Myeloid lineage | | |
| GO_REGULATION_OF_LYMPHOCYTE_DIFFERENTIATION | NIR | immune system | Myeloid lineage | | |
| GO_REGULATION_OF_MAST_CELL_ACTIVATION | NIR | immune system | Myeloid lineage | | |
| GO_REGULATION_OF_MAST_CELL_ACTIVATION_INVOLVED_IN_IMMUNE_RESPONSE | NIR | immune system | Myeloid lineage | | |
| GO_REGULATION_OF_MAST_CELL_DEGRANULATION | NIR | immune system | Myeloid lineage | | |

| | | | | |
|---|---|---|---|---|
| GO_REGULATION_OF_MYELOID_CELL_DIFFERENTIATION | NIR | immune system | Myeloid lineage | |
| GO_ANTIGEN_PROCESSING_AND_PRESENTATION | NIR | immune system | signal transduction | |
| GEN_PROCESSING_AND_PRESENTATION_OF_EXOGENOUS_PEPTIDE_ANTIGEN_VIA_MHC | NIR | immune system | signal transduction | |
| GO_ANTIGEN_PROCESSING_AND_PRESENTATION_OF_PEPTIDE_ANTIGEN | NIR | immune system | signal transduction | |
| )_ANTIGEN_PROCESSING_AND_PRESENTATION_OF_PEPTIDE_ANTIGEN_VIA_MHC_CLASS | NIR | immune system | signal transduction | |
| _PROCESSING_AND_PRESENTATION_OF_PEPTIDE_OR_POLYSACCHARIDE_ANTIGEN_VIA_ | NIR | immune system | signal transduction | |
| GO_ANTIGEN_RECEPTOR_MEDIATED_SIGNALING_PATHWAY | NIR | immune system | signal transduction | |
| GO_B_CELL_RECEPTOR_SIGNALING_PATHWAY | NIR | immune system | signal transduction | |
| GO_CELLULAR_RESPONSE_TO_GROWTH_HORMONE_STIMULUS | NIR | immune system | signal transduction | |
| GO_CHEMOKINE_BINDING | NIR | immune system | signal transduction | |
| GO_CYTOKINE_MEDIATED_SIGNALING_PATHWAY | NIR | immune system | signal transduction | |
| GO_CYTOKINE_RECEPTOR_ACTIVITY | NIR | immune system | signal transduction | |
| GO_ENDOLYSOSOME | NIR | immune system | signal transduction | |
| GO_ER_TO_GOLGI_TRANSPORT_VESICLE | NIR | immune system | signal transduction | |
| GO_FC_EPSILON_RECEPTOR_SIGNALING_PATHWAY | NIR | immune system | signal transduction | |
| GO_FC_GAMMA_RECEPTOR_SIGNALING_PATHWAY | NIR | immune system | signal transduction | |
| GO_FC_RECEPTOR_SIGNALING_PATHWAY | NIR | immune system | signal transduction | |
| GO_G_PROTEIN_COUPLED_CHEMOATTRACTANT_RECEPTOR_ACTIVITY | NIR | immune system | signal transduction | |
| ;O_IMMUNE_RESPONSE_REGULATING_CELL_SURFACE_RECEPTOR_SIGNALING_PATHWA | NIR | immune system | signal transduction | |
| NNATE_IMMUNE_RESPONSE_ACTIVATING_CELL_SURFACE_RECEPTOR_SIGNALING_PATH | NIR | immune system | signal transduction | |
| GO_LIPOPOLYSACCHARIDE_BINDING | NIR | immune system | signal transduction | |
| GO_LUMENAL_SIDE_OF_MEMBRANE | NIR | immune system | signal transduction | |
| GO_MHC_PROTEIN_COMPLEX | NIR | immune system | signal transduction | |
| GO_MYD88_DEPENDENT_TOLL_LIKE_RECEPTOR_SIGNALING_PATHWAY | NIR | immune system | signal transduction | |
| GO_MYD88_INDEPENDENT_TOLL_LIKE_RECEPTOR_SIGNALING_PATHWAY | NIR | immune system | signal transduction | |
| GO_NON_MEMBRANE_SPANNING_PROTEIN_TYROSINE_KINASE_ACTIVITY | NIR | immune system | signal transduction | |
| GO_PATTERN_RECOGNITION_RECEPTOR_SIGNALING_PATHWAY | NIR | immune system | signal transduction | |
| GO_PEPTIDYL_TYROSINE_AUTOPHOSPHORYLATION | NIR | immune system | signal transduction | |
| GO_PEPTIDYL_TYROSINE_MODIFICATION | NIR | immune system | signal transduction | |
| GO_POSITIVE_REGULATION_OF_TOLL_LIKE_RECEPTOR_SIGNALING_PATHWAY | NIR | immune system | signal transduction | |
| GO_PROTEIN_AUTOPHOSPHORYLATION | NIR | immune system | signal transduction | |
| GO_PROTEIN_POLYUBIQUITINATION | NIR | immune system | signal transduction | |
| GO_REGULATION_OF_ANTIGEN_PROCESSING_AND_PRESENTATION | NIR | immune system | signal transduction | |
| GO_REGULATION_OF_TOLL_LIKE_RECEPTOR_4_SIGNALING_PATHWAY | NIR | immune system | signal transduction | |
| GO_REGULATION_OF_TOLL_LIKE_RECEPTOR_SIGNALING_PATHWAY | NIR | immune system | signal transduction | |
| GO_T_CELL_RECEPTOR_COMPLEX | NIR | immune system | signal transduction | |
| GO_T_CELL_RECEPTOR_SIGNALING_PATHWAY | NIR | immune system | signal transduction | |
| GO_TOLL_LIKE_RECEPTOR_4_SIGNALING_PATHWAY | NIR | immune system | signal transduction | |
| GO_TOLL_LIKE_RECEPTOR_SIGNALING_PATHWAY | NIR | immune system | signal transduction | |
| GO_TUMOR_NECROSIS_FACTOR_MEDIATED_SIGNALING_PATHWAY | NIR | immune system | signal transduction | |
| GO_TUMOR_NECROSIS_FACTOR_RECEPTOR_BINDING | NIR | immune system | signal transduction | |
| GO_TUMOR_NECROSIS_FACTOR_RECEPTOR_SUPERFAMILY_BINDING | NIR | immune system | signal transduction | |
| GO_IMMUNE_EFFECTOR_PROCESS | NIR | immune system | | |
| GO_POLYUBIQUITIN_BINDING | NIR | metabolism | Apoptosis | ubiqutin |
| GO_UBIQUITIN_LIKE_PROTEIN_BINDING | NIR | metabolism | Apoptosis | ubiqutin |
| GO_DNA_DAMAGE_RESPONSE_SIGNAL_TRANSDUCTION_RESULTING_IN_TRANSCRIPTION | NIR | metabolism | Apoptosis | |
| IC_APOPTOTIC_SIGNALING_PATHWAY_IN_RESPONSE_TO_DNA_DAMAGE_BY_P53_CLASS | NIR | metabolism | Apoptosis | |
| _OF_MITOCHONDRIAL_OUTER_MEMBRANE_PERMEABILIZATION_INVOLVED_IN_APOPT | NIR | metabolism | Apoptosis | |
| I_OF_PROTEIN_INSERTION_INTO_MITOCHONDRIAL_MEMBRANE_INVOLVED_IN_APOPT | NIR | metabolism | Apoptosis | |
| MITOCHONDRIAL_OUTER_MEMBRANE_PERMEABILIZATION_INVOLVED_IN_APOPTOTIC_ | NIR | metabolism | Apoptosis | |
| _PROTEIN_INSERTION_INTO_MITOCHONDRIAL_MEMBRANE_INVOLVED_IN_APOPTOTIC_ | NIR | metabolism | Apoptosis | |
| GO_SIGNAL_TRANSDUCTION_BY_P53_CLASS_MEDIATOR | NIR | metabolism | Apoptosis | |
| ON_OF_CYSTEINE_TYPE_ENDOPEPTIDASE_ACTIVITY_INVOLVED_IN_APOPTOTIC_SIGNALI | NIR | metabolism | Apoptosis | |
| ULATION_OF_CYSTEINE_TYPE_ENDOPEPTIDASE_ACTIVITY_INVOLVED_IN_APOPTOTIC_SIG | NIR | metabolism | Apoptosis | |
| GO_REGULATION_OF_MICROTUBULE_POLYMERIZATION | PIR | metabolism | Cell proliferation | |
| GO_ACTIN_NUCLEATION | NIR | metabolism | Cell proliferation | |
| GO_MICROTUBULE_CYTOSKELETON_ORGANIZATION_INVOLVED_IN_MITOSIS | NIR | metabolism | Cell proliferation | |
| GO_MITOTIC_SPINDLE_ASSEMBLY | NIR | metabolism | Cell proliferation | |
| GO_POSITIVE_REGULATION_OF_PROTEIN_COMPLEX_ASSEMBLY | NIR | metabolism | Cell proliferation | |
| GO_SPINDLE_ASSEMBLY | NIR | metabolism | Cell proliferation | |
| GO_ATPASE_ACTIVATOR_ACTIVITY | PIR | metabolism | cellular respiration | |
| GO_ATPASE_REGULATOR_ACTIVITY | PIR | metabolism | cellular respiration | |
| GO_POSITIVE_REGULATION_OF_ATPASE_ACTIVITY | PIR | metabolism | cellular respiration | |
| GO_REGULATION_OF_ATPASE_ACTIVITY | PIR | metabolism | cellular respiration | |
| GO_BETA_CATENIN_TCF_COMPLEX_ASSEMBLY | NIR | Nucleus activity | function | |
| GO_CHROMATIN | NIR | Nucleus activity | function | |
| GO_CHROMATIN_SILENCING | NIR | Nucleus activity | function | |
| GO_CHROMATIN_SILENCING_AT_RDNA | NIR | Nucleus activity | function | |
| GO_CHROMOSOME_TELOMERIC_REGION | NIR | Nucleus activity | function | |
| GO_DNA_DEPENDENT_DNA_REPLICATION_MAINTENANCE_OF_FIDELITY | NIR | Nucleus activity | function | |
| GO_DNA_REPLICATION_DEPENDENT_NUCLEOSOME_ASSEMBLY | NIR | Nucleus activity | function | |
| GO_DNA_REPLICATION_DEPENDENT_NUCLEOSOME_ORGANIZATION | NIR | Nucleus activity | function | |
| GO_GENE_SILENCING | NIR | Nucleus activity | function | |
| GO_GENE_SILENCING_BY_RNA | NIR | Nucleus activity | function | |
| GO_NEGATIVE_REGULATION_OF_DNA_METABOLIC_PROCESS | NIR | Nucleus activity | function | |
| GO_NEGATIVE_REGULATION_OF_DNA_RECOMBINATION | NIR | Nucleus activity | function | |
| GO_NEGATIVE_REGULATION_OF_DNA_REPLICATION | NIR | Nucleus activity | function | |
| GO_NEGATIVE_REGULATION_OF_GENE_EXPRESSION_EPIGENETIC | NIR | Nucleus activity | function | |
| GO_NEGATIVE_REGULATION_OF_GENE_SILENCING | NIR | Nucleus activity | function | |
| GO_NUCLEAR_CHROMATIN | NIR | Nucleus activity | function | |
| GO_NUCLEAR_CHROMOSOME | NIR | Nucleus activity | function | |
| GO_NUCLEAR_CHROMOSOME_TELOMERIC_REGION | NIR | Nucleus activity | function | |
| GO_POSITIVE_REGULATION_OF_GENE_EXPRESSION_EPIGENETIC | NIR | Nucleus activity | function | |
| GO_PROTEIN_HETEROTETRAMERIZATION | NIR | Nucleus activity | function | |
| GO_PROTEIN_HOMOTETRAMERIZATION | NIR | Nucleus activity | function | |
| GO_PROTEIN_TETRAMERIZATION | NIR | Nucleus activity | function | |
| GO_REGULATION_OF_DNA_METABOLIC_PROCESS | NIR | Nucleus activity | function | |
| GO_REGULATION_OF_DNA_RECOMBINATION | NIR | Nucleus activity | function | |
| GO_REGULATION_OF_DOUBLE_STRAND_BREAK_REPAIR | NIR | Nucleus activity | function | |
| GO_REGULATION_OF_GENE_EXPRESSION_EPIGENETIC | NIR | Nucleus activity | function | |
| GO_REGULATION_OF_GENE_SILENCING | NIR | Nucleus activity | function | |
| GO_REGULATION_OF_RESPONSE_TO_DNA_DAMAGE_STIMULUS | NIR | Nucleus activity | function | |
| GO_TELOMERE_ORGANIZATION | NIR | Nucleus activity | function | |
| GO_DNA_DEALKYLATION | NIR | Nucleus activity | function | |
| GO_DNA_MODIFICATION | NIR | Nucleus activity | function | |
| GO_ENDONUCLEASE_ACTIVITY | NIR | Nucleus activity | nucleotide decomposition | |
| GO_ENDORIBONUCLEASE_ACTIVITY | NIR | Nucleus activity | nucleotide decomposition | |
| GO_NUCLEIC_ACID_PHOSPHODIESTER_BOND_HYDROLYSIS | NIR | Nucleus activity | nucleotide decomposition | |
| GO_RNA_PHOSPHODIESTER_BOND_HYDROLYSIS_ENDONUCLEOLYTIC | NIR | Nucleus activity | nucleotide decomposition | |
| GO_GLYCOSYL_COMPOUND_CATABOLIC_PROCESS | NIR | Nucleus activity | nucleotide synthesis | |
| GO_PYRIMIDINE_CONTAINING_COMPOUND_CATABOLIC_PROCESS | NIR | Nucleus activity | nucleotide synthesis | |
| GO_PYRIMIDINE_NUCLEOSIDE_CATABOLIC_PROCESS | NIR | Nucleus activity | nucleotide synthesis | |
| GO_PYRIMIDINE_NUCLEOSIDE_METABOLIC_PROCESS | NIR | Nucleus activity | nucleotide synthesis | |

| | | | | |
|---|---|---|---|---|
| GO_PYRIMIDINE_RIBONUCLEOSIDE_METABOLIC_PROCESS | NIR | Nucleus activity | nucleotide synthesis | |
| GO_RIBONUCLEOSIDE_CATABOLIC_PROCESS | NIR | Nucleus activity | nucleotide synthesis | |
| GO_NEGATIVE_REGULATION_OF_RNA_SPLICING | PIR | Nucleus activity | RNA regulation | |
| GO_REGULATION_OF_ALTERNATIVE_MRNA_SPLICING_VIA_SPLICEOSOME | PIR | Nucleus activity | RNA regulation | |
| GO_REGULATION_OF_MRNA_SPLICING_VIA_SPLICEOSOME | PIR | Nucleus activity | RNA regulation | |
| GO_MAIN_AXON | PIR | organogenesis | Neural development | neural cells |
| GO_NEGATIVE_REGULATION_OF_DENDRITE_DEVELOPMENT | PIR | organogenesis | Neural development | neural cells |
| GO_NEGATIVE_REGULATION_OF_DENDRITE_MORPHOGENESIS | PIR | organogenesis | Neural development | neural cells |
| GO_POSITIVE_REGULATION_OF_CELL_PROJECTION_ORGANIZATION | PIR | organogenesis | Neural development | neural cells |
| GO_POSITIVE_REGULATION_OF_DENDRITE_EXTENSION | PIR | organogenesis | Neural development | neural cells |
| GO_POSITIVE_REGULATION_OF_LAMELLIPODIUM_ASSEMBLY | PIR | organogenesis | Neural development | neural cells |
| GO_POSITIVE_REGULATION_OF_NEURON_PROJECTION_DEVELOPMENT | PIR | organogenesis | Neural development | neural cells |
| GO_REGULATION_OF_DENDRITE_DEVELOPMENT | PIR | organogenesis | Neural development | neural cells |
| GO_REGULATION_OF_DENDRITE_EXTENSION | PIR | organogenesis | Neural development | neural cells |
| GO_REGULATION_OF_DENDRITE_MORPHOGENESIS | PIR | organogenesis | Neural development | neural cells |
| GO_REGULATION_OF_NEURON_PROJECTION_DEVELOPMENT | PIR | organogenesis | Neural development | neural cells |
| GO_SITE_OF_POLARIZED_GROWTH | PIR | organogenesis | Neural development | neural cells |
| GO_NEURON_PROJECTION_REGENERATION | PIR | organogenesis | Neural development | neural regeneration |
| GO_AXON_REGENERATION | PIR | organogenesis | Neural development | neural regeneration |
| GO_NEGATIVE_REGULATION_OF_MACROAUTOPHAGY | PIR | organogenesis | Neural development | nutrient |
| GO_NEGATIVE_REGULATION_OF_RESPONSE_TO_EXTRACELLULAR_STIMULUS | PIR | organogenesis | Neural development | nutrient |
| GO_NEGATIVE_REGULATION_OF_RESPONSE_TO_NUTRIENT_LEVELS | PIR | organogenesis | Neural development | nutrient |
| GO_REGULATION_OF_APPETITE | PIR | organogenesis | Neural development | nutrient |
| GO_REGULATION_OF_RESPONSE_TO_EXTRACELLULAR_STIMULUS | PIR | organogenesis | Neural development | nutrient |
| GO_REGULATION_OF_RESPONSE_TO_FOOD | PIR | organogenesis | Neural development | nutrient |
| GO_REGULATION_OF_RESPONSE_TO_NUTRIENT_LEVELS | PIR | organogenesis | Neural development | nutrient |
| GO_NEURAL_RETINA_DEVELOPMENT | PIR | organogenesis | Neural development | retine |
| GO_RETINA_DEVELOPMENT_IN_CAMERA_TYPE_EYE | PIR | organogenesis | Neural development | retine |
| GO_RETINA_LAYER_FORMATION | PIR | organogenesis | Neural development | retine |
| GO_RETINA_MORPHOGENESIS_IN_CAMERA_TYPE_EYE | PIR | organogenesis | Neural development | retine |
| GO_AUDITORY_RECEPTOR_CELL_DEVELOPMENT | PIR | organogenesis | Neural development | sensory of mechanical stimulus |
| GO_AUDITORY_RECEPTOR_CELL_DIFFERENTIATION | PIR | organogenesis | Neural development | sensory of mechanical stimulus |
| GO_DETECTION_OF_MECHANICAL_STIMULUS_INVOLVED_IN_SENSORY_PERCEPTION | PIR | organogenesis | Neural development | sensory of mechanical stimulus |
| GO_SENSORY_PERCEPTION | PIR | organogenesis | Neural development | sensory of mechanical stimulus |
| GO_SENSORY_PERCEPTION_OF_MECHANICAL_STIMULUS | PIR | organogenesis | Neural development | sensory of mechanical stimulus |
| GO_PROSTATE_GLAND_DEVELOPMENT | NIR | organogenesis | secretory gland | prostate gland |
| GO_PROSTATE_GLAND_MORPHOGENESIS | NIR | organogenesis | secretory gland | prostate gland |
| GO_MAMMARY_GLAND_EPITHELIAL_CELL_DIFFERENTIATION | NIR | organogenesis | secretory gland | |
| GO_MAMMARY_GLAND_EPITHELIUM_DEVELOPMENT | NIR | organogenesis | secretory gland | |
| GO_ANATOMICAL_STRUCTURE_HOMEOSTASIS | NIR | organogenesis | Tissue homeostasis | |
| GO_BONE_RESORPTION | NIR | organogenesis | Tissue homeostasis | |
| GO_HOMEOSTASIS_OF_NUMBER_OF_CELLS | NIR | organogenesis | Tissue homeostasis | |
| GO_HOMEOSTASIS_OF_NUMBER_OF_CELLS_WITHIN_A_TISSUE | NIR | organogenesis | Tissue homeostasis | |
| GO_TISSUE_HOMEOSTASIS | NIR | organogenesis | Tissue homeostasis | |
| GO_POSITIVE_REGULATION_OF_MYOTUBE_DIFFERENTIATION | NIR | organogenesis | muscle development | |
| GO_REGULATION_OF_SYNCYTIUM_FORMATION_BY_PLASMA_MEMBRANE_FUSION | NIR | organogenesis | muscle development | |
| GO_ACTIN_FILAMENT_BASED_MOVEMENT | PIR | physiological function | Cardiovascular function | |
| GO_ACTIN_MEDIATED_CELL_CONTRACTION | PIR | physiological function | Cardiovascular function | |
| GO_ACTIN_MYOSIN_FILAMENT_SLIDING | PIR | physiological function | Cardiovascular function | |
| GO_ACTION_POTENTIAL | PIR | physiological function | Cardiovascular function | |
| GO_CARDIAC_MUSCLE_CELL_CONTRACTION | PIR | physiological function | Cardiovascular function | |
| GO_CELL_CELL_SIGNALING_INVOLVED_IN_CARDIAC_CONDUCTION | PIR | physiological function | Cardiovascular function | |
| GO_CELL_COMMUNICATION_BY_ELECTRICAL_COUPLING | PIR | physiological function | Cardiovascular function | |
| GO_HEART_PROCESS | PIR | physiological function | Cardiovascular function | |
| GO_MEMBRANE_DEPOLARIZATION_DURING_ACTION_POTENTIAL | PIR | physiological function | Cardiovascular function | |
| GO_MUSCLE_CONTRACTION | PIR | physiological function | Cardiovascular function | |
| GO_MUSCLE_FILAMENT_SLIDING | PIR | physiological function | Cardiovascular function | |
| GO_MUSCLE_SYSTEM_PROCESS | PIR | physiological function | Cardiovascular function | |
| GO_REGULATION_OF_BLOOD_CIRCULATION | PIR | physiological function | Cardiovascular function | |
| GO_REGULATION_OF_HEART_CONTRACTION | PIR | physiological function | Cardiovascular function | |
| GO_REGULATION_OF_HEART_RATE | PIR | physiological function | Cardiovascular function | |
| GO_REGULATION_OF_STRIATED_MUSCLE_CONTRACTION | PIR | physiological function | Cardiovascular function | |
| GO_REGULATION_OF_THE_FORCE_OF_HEART_CONTRACTION | PIR | physiological function | Cardiovascular function | |
| GO_STRIATED_MUSCLE_CONTRACTION | PIR | physiological function | Cardiovascular function | |
| GO_VOLTAGE_GATED_ION_CHANNEL_ACTIVITY | PIR | physiological function | Cardiovascular function | |
| GO_ACETYLCHOLINE_RECEPTOR_ACTIVITY | PIR | physiological function | neural function | synapse |
| GO_ADULT_BEHAVIOR | PIR | physiological function | neural function | synapse |
| GO_AMMONIUM_ION_BINDING | PIR | physiological function | neural function | synapse |
| GO_ANTEROGRADE_AXONAL_TRANSPORT | PIR | physiological function | neural function | synapse |
| GO_AXON | PIR | physiological function | neural function | synapse |
| GO_AXON_CYTOPLASM | PIR | physiological function | neural function | synapse |
| GO_AXON_PART | PIR | physiological function | neural function | synapse |
| GO_BLOC_1_COMPLEX | PIR | physiological function | neural function | synapse |
| GO_BLOC_COMPLEX | PIR | physiological function | neural function | synapse |
| GO_CALCIUM_ION_REGULATED_EXOCYTOSIS | PIR | physiological function | neural function | synapse |
| GO_CALCIUM_ION_REGULATED_EXOCYTOSIS_OF_NEUROTRANSMITTER | PIR | physiological function | neural function | synapse |
| GO_CELL_BODY | PIR | physiological function | neural function | synapse |
| GO_CELL_PROJECTION_CYTOPLASM | PIR | physiological function | neural function | synapse |
| D_CELL_SURFACE_RECEPTOR_SIGNALING_PATHWAY_INVOLVED_IN_CELL_CELL_SIGNALII | PIR | physiological function | neural function | synapse |
| GO_CLATHRIN_BINDING | PIR | physiological function | neural function | synapse |
| GO_CLATHRIN_MEDIATED_ENDOCYTOSIS | PIR | physiological function | neural function | synapse |
| GO_COPI_COATED_VESICLE | PIR | physiological function | neural function | synapse |
| GO_CYTOSOLIC_TRANSPORT | PIR | physiological function | neural function | synapse |
| GO_DENDRITE | PIR | physiological function | neural function | synapse |
| GO_DENDRITE_MEMBRANE | PIR | physiological function | neural function | synapse |
| GO_DENDRITIC_SHAFT | PIR | physiological function | neural function | synapse |
| GO_DNA_DAMAGE_RESPONSE_SIGNAL_TRANSDUCTION_RESULTING_IN_TRANSCRIPTION | PIR | physiological function | neural function | synapse |
| GO_DOPAMINE_METABOLIC_PROCESS | PIR | physiological function | neural function | synapse |
| GO_ESTABLISHMENT_OF_MITOCHONDRION_LOCALIZATION | PIR | physiological function | neural function | synapse |
| GO_EXCITATORY_EXTRACELLULAR_LIGAND_GATED_ION_CHANNEL_ACTIVITY | PIR | physiological function | neural function | synapse |
| GO_EXCITATORY_POSTSYNAPTIC_POTENTIAL | PIR | physiological function | neural function | synapse |
| GO_EXCITATORY_SYNAPSE | PIR | physiological function | neural function | synapse |
| GO_EXOCYTIC_VESICLE | PIR | physiological function | neural function | synapse |
| GO_EXOCYTIC_VESICLE_MEMBRANE | PIR | physiological function | neural function | synapse |
| GO_EXTRACELLULAR_LIGAND_GATED_ION_CHANNEL_ACTIVITY | PIR | physiological function | neural function | synapse |
| GO_GLUTAMATE_SECRETION | PIR | physiological function | neural function | synapse |
| GO_INTRASPECIES_INTERACTION_BETWEEN_ORGANISMS | PIR | physiological function | neural function | synapse |
| GO_LIGAND_GATED_CHANNEL_ACTIVITY | PIR | physiological function | neural function | synapse |
| GO_PIR_TERM_MEMORY | PIR | physiological function | neural function | synapse |
| GO_PIR_TERM_SYNAPTIC_DEPRESSION | PIR | physiological function | neural function | synapse |
| GO_MITOCHONDRION_LOCALIZATION | PIR | physiological function | neural function | synapse |
| GO_MODULATION_OF_EXCITATORY_POSTSYNAPTIC_POTENTIAL | PIR | physiological function | neural function | synapse |
| GO_MODULATION_OF_SYNAPTIC_TRANSMISSION | PIR | physiological function | neural function | synapse |

| | | | | |
|---|---|---|---|---|
| GO_NEGATIVE_REGULATION_OF_AMINE_TRANSPORT | PIR | physiological function | neural function | synapse |
| GO_NEGATIVE_REGULATION_OF_CATECHOLAMINE_SECRETION | PIR | physiological function | neural function | synapse |
| GO_NEUROMUSCULAR_SYNAPTIC_TRANSMISSION | PIR | physiological function | neural function | synapse |
| GO_NEURON_NEURON_SYNAPTIC_TRANSMISSION | PIR | physiological function | neural function | synapse |
| GO_NEURON_PROJECTION_MEMBRANE | PIR | physiological function | neural function | synapse |
| GO_NEURON_PROJECTION_TERMINUS | PIR | physiological function | neural function | synapse |
| GO_NEURONAL_POSTSYNAPTIC_DENSITY | PIR | physiological function | neural function | synapse |
| GO_NEUROTRANSMITTER_BINDING | PIR | physiological function | neural function | synapse |
| GO_NEUROTRANSMITTER_METABOLIC_PROCESS | PIR | physiological function | neural function | synapse |
| GO_NEUROTRANSMITTER_RECEPTOR_ACTIVITY | PIR | physiological function | neural function | synapse |
| GO_NEUROTRANSMITTER_TRANSPORT | PIR | physiological function | neural function | synapse |
| GO_ORGANELLE_FUSION | PIR | physiological function | neural function | synapse |
| GO_ORGANELLE_LOCALIZATION | PIR | physiological function | neural function | synapse |
| GO_ORGANELLE_MEMBRANE_FUSION | PIR | physiological function | neural function | synapse |
| GO_ORGANELLE_TRANSPORT_APIR_MICROTUBULE | PIR | physiological function | neural function | synapse |
| GO_PERIKARYON | PIR | physiological function | neural function | synapse |
| GO_POSITIVE_REGULATION_OF_EXCITATORY_POSTSYNAPTIC_POTENTIAL | PIR | physiological function | neural function | synapse |
| GO_POSITIVE_REGULATION_OF_SYNAPSE_ASSEMBLY | PIR | physiological function | neural function | synapse |
| GO_POSITIVE_REGULATION_OF_SYNAPTIC_TRANSMISSION | PIR | physiological function | neural function | synapse |
| GO_POSTSYNAPSE | PIR | physiological function | neural function | synapse |
| GO_POSTSYNAPTIC_MEMBRANE | PIR | physiological function | neural function | synapse |
| GO_PRESYNAPSE | PIR | physiological function | neural function | synapse |
| GO_PRESYNAPTIC_ACTIVE_ZONE | PIR | physiological function | neural function | synapse |
| GO_PRESYNAPTIC_MEMBRANE | PIR | physiological function | neural function | synapse |
| GO_PRESYNAPTIC_PROCESS_INVOLVED_IN_SYNAPTIC_TRANSMISSION | PIR | physiological function | neural function | synapse |
| GO_REGULATION_OF_CALCIUM_ION_DEPENDENT_EXOCYTOSIS | PIR | physiological function | neural function | synapse |
| GO_REGULATION_OF_CATECHOLAMINE_METABOLIC_PROCESS | PIR | physiological function | neural function | synapse |
| GO_REGULATION_OF_DOPAMINE_METABOLIC_PROCESS | PIR | physiological function | neural function | synapse |
| GO_REGULATION_OF_PIR_TERM_NEURONAL_SYNAPTIC_PLASTICITY | PIR | physiological function | neural function | synapse |
| GO_REGULATION_OF_MEMBRANE_POTENTIAL | PIR | physiological function | neural function | synapse |
| GO_REGULATION_OF_NEURONAL_SYNAPTIC_PLASTICITY | PIR | physiological function | neural function | synapse |
| GO_REGULATION_OF_NEUROTRANSMITTER_LEVELS | PIR | physiological function | neural function | synapse |
| GO_REGULATION_OF_NEUROTRANSMITTER_SECRETION | PIR | physiological function | neural function | synapse |
| GO_REGULATION_OF_NOREPINEPHRINE_SECRETION | PIR | physiological function | neural function | synapse |
| GO_REGULATION_OF_POSTSYNAPTIC_MEMBRANE_POTENTIAL | PIR | physiological function | neural function | synapse |
| GO_REGULATION_OF_SYNAPSE_ASSEMBLY | PIR | physiological function | neural function | synapse |
| GO_REGULATION_OF_SYNAPSE_ORGANIZATION | PIR | physiological function | neural function | synapse |
| GO_REGULATION_OF_SYNAPSE_STRUCTURE_OR_ACTIVITY | PIR | physiological function | neural function | synapse |
| GO_REGULATION_OF_SYNAPTIC_PLASTICITY | PIR | physiological function | neural function | synapse |
| GO_REGULATION_OF_SYNAPTIC_VESICLE_EXOCYTOSIS | PIR | physiological function | neural function | synapse |
| GO_REGULATION_OF_SYNAPTIC_VESICLE_TRANSPORT | PIR | physiological function | neural function | synapse |
| GO_RESPONSE_TO_ALKALOID | PIR | physiological function | neural function | synapse |
| GO_RESPONSE_TO_ISOQUINOLINE_ALKALOID | PIR | physiological function | neural function | synapse |
| GO_RESPONSE_TO_MORPHINE | PIR | physiological function | neural function | synapse |
| GO_SIGNAL_RELEASE | PIR | physiological function | neural function | synapse |
| GO_SINGLE_ORGANISM_MEMBRANE_FUSION | PIR | physiological function | neural function | synapse |
| GO_SNAP_RECEPTOR_ACTIVITY | PIR | physiological function | neural function | synapse |
| GO_SNARE_BINDING | PIR | physiological function | neural function | synapse |
| GO_SNARE_COMPLEX | PIR | physiological function | neural function | synapse |
| GO_SOCIAL_BEHAVIOR | PIR | physiological function | neural function | synapse |
| GO_STARTLE_RESPONSE | PIR | physiological function | neural function | synapse |
| GO_SYNAPTIC_MEMBRANE | PIR | physiological function | neural function | synapse |
| GO_SYNAPTIC_SIGNALING | PIR | physiological function | neural function | synapse |
| GO_SYNAPTIC_TRANSMISSION_CHOLINERGIC | PIR | physiological function | neural function | synapse |
| GO_SYNAPTIC_TRANSMISSION_DOPAMINERGIC | PIR | physiological function | neural function | synapse |
| GO_SYNAPTIC_TRANSMISSION_GLUTAMATERGIC | PIR | physiological function | neural function | synapse |
| GO_SYNAPTIC_VESICLE_CYCLE | PIR | physiological function | neural function | synapse |
| GO_SYNAPTIC_VESICLE_ENDOCYTOSIS | PIR | physiological function | neural function | synapse |
| GO_SYNAPTIC_VESICLE_LOCALIZATION | PIR | physiological function | neural function | synapse |
| GO_SYNAPTIC_VESICLE_RECYCLING | PIR | physiological function | neural function | synapse |
| GO_SYNTAXIN_1_BINDING | PIR | physiological function | neural function | synapse |
| GO_SYNTAXIN_BINDING | PIR | physiological function | neural function | synapse |
| GO_TERMINAL_BOUTON | PIR | physiological function | neural function | synapse |
| GO_TRANSPORT_VESICLE | PIR | physiological function | neural function | synapse |
| GO_VESICLE_DOCKING | PIR | physiological function | neural function | synapse |
| GO_VESICLE_LOCALIZATION | PIR | physiological function | neural function | synapse |
| GO_VESICLE_ORGANIZATION | PIR | physiological function | neural function | synapse |
| GO_INOSITOL_LIPID_MEDIATED_SIGNALING | NIR | Signal transduction | lipid | |
| GO_LIPID_PHOSPHORYLATION | NIR | Signal transduction | lipid | |
| GO_PHOSPHATIDYLINOSITOL_3_KINASE_ACTIVITY | NIR | Signal transduction | lipid | |
| GO_PHOSPHATIDYLINOSITOL_3_KINASE_COMPLEX | NIR | Signal transduction | lipid | |
| GO_REGULATION_OF_PHOSPHATIDYLINOSITOL_3_KINASE_SIGNALING | NIR | Signal transduction | lipid | |
| GO_JNK_CASCADE | NIR | Signal transduction | protein | |
| GO_POSITIVE_REGULATION_OF_JUN_KINASE_ACTIVITY | NIR | Signal transduction | protein | |
| )_POSITIVE_REGULATION_OF_STRESS_ACTIVATED_PROTEIN_KINASE_SIGNALING_CASCA | NIR | Signal transduction | protein | |
| GO_REGULATION_OF_JNK_CASCADE | NIR | Signal transduction | protein | |
| GO_REGULATION_OF_JUN_KINASE_ACTIVITY | NIR | Signal transduction | protein | |
| GO_REGULATION_OF_STRESS_ACTIVATED_PROTEIN_KINASE_SIGNALING_CASCADE | NIR | Signal transduction | protein | |
| GO_STRESS_ACTIVATED_PROTEIN_KINASE_SIGNALING_CASCADE | NIR | Signal transduction | protein | |
| GO_SIGNALING_ADAPTOR_ACTIVITY | NIR | Signal transduction | protein | |
| GO_SH3_SH2_ADAPTOR_ACTIVITY | NIR | Signal transduction | protein | |
| GO_ANDROGEN_RECEPTOR_SIGNALING_PATHWAY | NIR | Signal transduction | sterols | |
| GO_BHLH_TRANSCRIPTION_FACTOR_BINDING | NIR | Signal transduction | sterols | |
| GO_DNA_BINDING_BENDING | NIR | Signal transduction | sterols | |
| GO_DNA_TEMPLATED_TRANSCRIPTION_INITIATION | NIR | Signal transduction | sterols | |
| GO_ESTROGEN_RECEPTOR_BINDING | NIR | Signal transduction | sterols | |
| GO_INTRACELLULAR_RECEPTOR_SIGNALING_PATHWAY | NIR | Signal transduction | sterols | |
| GO_REPRESSING_TRANSCRIPTION_FACTOR_BINDING | NIR | Signal transduction | sterols | |
| GO_RETINOIC_ACID_RECEPTOR_BINDING | NIR | Signal transduction | sterols | |
| GO_STEROID_HORMONE_MEDIATED_SIGNALING_PATHWAY | NIR | Signal transduction | sterols | |
| GO_STEROID_HORMONE_RECEPTOR_ACTIVITY | NIR | Signal transduction | sterols | |
| GO_STEROID_HORMONE_RECEPTOR_BINDING | NIR | Signal transduction | sterols | |
| RIPTION_FACTOR_ACTIVITY_RNA_POLYMERASE_II_DISTAL_ENHANCER_SEQUENCE_SPECI | NIR | Signal transduction | sterols | |
| GO_TRANSCRIPTION_FACTOR_BINDING | NIR | Signal transduction | sterols | |
| _REGULATION_OF_INTRACELLULAR_STEROID_HORMONE_RECEPTOR_SIGNALING_PATHV | NIR | Signal transduction | sterols | |
| ATIVE_REGULATION_OF_INTRACELLULAR_STEROID_HORMONE_RECEPTOR_SIGNALING_I | NIR | Signal transduction | sterols | |
| GO_POSITIVE_REGULATION_OF_POTASSIUM_ION_TRANSMEMBRANE_TRANSPORT | PIR | transport | synaptic | |
| _POSITIVE_REGULATION_OF_POTASSIUM_ION_TRANSMEMBRANE_TRANSPORTER_ACTIV | PIR | transport | synaptic | |
| GO_POSITIVE_REGULATION_OF_POTASSIUM_ION_TRANSPORT | PIR | transport | synaptic | |
| GO_POSITIVE_REGULATION_OF_TRANSPORTER_ACTIVITY | PIR | transport | synaptic | |
| GO_REGULATION_OF_CALCIUM_ION_TRANSMEMBRANE_TRANSPORTER_ACTIVITY | PIR | transport | synaptic | |
| GO_REGULATION_OF_CATION_CHANNEL_ACTIVITY | PIR | transport | synaptic | |
| GO_REGULATION_OF_POTASSIUM_ION_TRANSMEMBRANE_TRANSPORTER_ACTIVITY | PIR | transport | synaptic | |

| | | | |
|---|---|---|---|
| GO_REGULATION_OF_VOLTAGE_GATED_CALCIUM_CHANNEL_ACTIVITY | PIR | transport | synaptic |
| GO_NEGATIVE_REGULATION_OF_CALCIUM_MEDIATED_SIGNALING | NIR | transport | synaptic |
| GO_ACTIVE_ION_TRANSMEMBRANE_TRANSPORTER_ACTIVITY | PIR | transport | |
| GO_ACTIVE_TRANSMEMBRANE_TRANSPORTER_ACTIVITY | PIR | transport | |
| GO_AMINO_ACID_TRANSMEMBRANE_TRANSPORT | PIR | transport | |
| GO_AMINO_ACID_TRANSMEMBRANE_TRANSPORTER_ACTIVITY | PIR | transport | |
| GO_AMINO_ACID_TRANSPORT | PIR | transport | |
| GO_AMMONIUM_TRANSMEMBRANE_TRANSPORT | PIR | transport | |
| GO_AMMONIUM_TRANSPORT | PIR | transport | |
| GO_ANION_CATION_SYMPORTER_ACTIVITY | PIR | transport | |
| GO_ANION_TRANSMEMBRANE_TRANSPORT | PIR | transport | |
| GO_ANION_TRANSMEMBRANE_TRANSPORTER_ACTIVITY | PIR | transport | |
| GO_ANION_TRANSPORT | PIR | transport | |
| GO_CATION_CHANNEL_ACTIVITY | PIR | transport | |
| GO_CATION_CHANNEL_COMPLEX | PIR | transport | |
| GO_CHLORIDE_CHANNEL_COMPLEX | PIR | transport | |
| GO_CHLORIDE_TRANSPORT | PIR | transport | |
| GO_DELAYED_RECTIFIER_POTASSIUM_CHANNEL_ACTIVITY | PIR | transport | |
| GO_GATED_CHANNEL_ACTIVITY | PIR | transport | |
| GO_INORGANIC_ANION_TRANSMEMBRANE_TRANSPORTER_ACTIVITY | PIR | transport | |
| GO_INORGANIC_ANION_TRANSPORT | PIR | transport | |
| GO_INORGANIC_CATION_TRANSMEMBRANE_TRANSPORTER_ACTIVITY | PIR | transport | |
| GO_L_AMINO_ACID_TRANSMEMBRANE_TRANSPORTER_ACTIVITY | PIR | transport | |
| GO_L_AMINO_ACID_TRANSPORT | PIR | transport | |
| GO_METAL_ION_TRANSMEMBRANE_TRANSPORTER_ACTIVITY | PIR | transport | |
| GO_MONOVALENT_INORGANIC_CATION_TRANSMEMBRANE_TRANSPORTER_ACTIVITY | PIR | transport | |
| GO_MONOVALENT_INORGANIC_CATION_TRANSPORT | PIR | transport | |
| GO_NEUROTRANSMITTER_SODIUM_SYMPORTER_ACTIVITY | PIR | transport | |
| GO_NEUROTRANSMITTER_TRANSPORTER_ACTIVITY | PIR | transport | |
| GO_NEUTRAL_AMINO_ACID_TRANSMEMBRANE_TRANSPORTER_ACTIVITY | PIR | transport | |
| GO_NEUTRAL_AMINO_ACID_TRANSPORT | PIR | transport | |
| GO_NITROGEN_COMPOUND_TRANSPORT | PIR | transport | |
| GO_NUCLEOTIDE_TRANSPORT | PIR | transport | |
| GO_ORGANIC_ACID_SODIUM_SYMPORTER_ACTIVITY | PIR | transport | |
| GO_ORGANIC_ACID_TRANSMEMBRANE_TRANSPORT | PIR | transport | |
| GO_ORGANIC_ACID_TRANSMEMBRANE_TRANSPORTER_ACTIVITY | PIR | transport | |
| GO_ORGANIC_ACID_TRANSPORT | PIR | transport | |
| GO_PASSIVE_TRANSMEMBRANE_TRANSPORTER_ACTIVITY | PIR | transport | |
| GO_POTASSIUM_CHANNEL_COMPLEX | PIR | transport | |
| GO_SODIUM_CHANNEL_ACTIVITY | PIR | transport | |
| GO_SODIUM_CHANNEL_COMPLEX | PIR | transport | |
| GO_SODIUM_ION_TRANSMEMBRANE_TRANSPORT | PIR | transport | |
| GO_SODIUM_ION_TRANSMEMBRANE_TRANSPORTER_ACTIVITY | PIR | transport | |
| GO_SODIUM_ION_TRANSPORT | PIR | transport | |
| GO_SOLUTE_CATION_SYMPORTER_ACTIVITY | PIR | transport | |
| GO_SOLUTE_SODIUM_SYMPORTER_ACTIVITY | PIR | transport | |
| GO_SYMPORTER_ACTIVITY | PIR | transport | |
| GO_TRANSPORTER_COMPLEX | PIR | transport | |
| GO_VOLTAGE_GATED_CATION_CHANNEL_ACTIVITY | PIR | transport | |
| GO_VOLTAGE_GATED_SODIUM_CHANNEL_ACTIVITY | PIR | transport | |

**Supplementary Data 2C. The grouping of GO terms in Cohort 3.**

| GO_term | Enrichment in | Type 1 | Type 2 |
|---|---|---|---|
| GO_POSITIVE_REGULATION_OF_ENDOTHELIAL_CELL_APOPTOTIC_PROCESS | NIR | angiogenesis | apototic |
| GO_POSITIVE_REGULATION_OF_EPITHELIAL_CELL_APOPTOTIC_PROCESS | NIR | angiogenesis | apototic |
| GO_REGULATION_OF_ENDOTHELIAL_CELL_APOPTOTIC_PROCESS | NIR | angiogenesis | apototic |
| GO_REGULATION_OF_EPITHELIAL_CELL_APOPTOTIC_PROCESS | NIR | angiogenesis | apototic |
| GO_ENDOTHELIUM_DEVELOPMENT | NIR | angiogenesis | |
| GO_MORPHOGENESIS_OF_AN_ENDOTHELIUM | NIR | angiogenesis | |
| GO_NEGATIVE_REGULATION_OF_BLOOD_VESSEL_ENDOTHELIAL_CELL_MIGRATION | NIR | angiogenesis | |
| GO_NEGATIVE_REGULATION_OF_ENDOTHELIAL_CELL_MIGRATION | NIR | angiogenesis | |
| GO_NEGATIVE_REGULATION_OF_ENDOTHELIAL_CELL_PROLIFERATION | NIR | angiogenesis | |
| GO_NEGATIVE_REGULATION_OF_EPITHELIAL_CELL_MIGRATION | NIR | angiogenesis | |
| GO_POSITIVE_REGULATION_OF_ENDOTHELIAL_CELL_MIGRATION | NIR | angiogenesis | |
| GO_POSITIVE_REGULATION_OF_EPITHELIAL_CELL_MIGRATION | NIR | angiogenesis | |
| GO_REGULATION_OF_BLOOD_VESSEL_ENDOTHELIAL_CELL_MIGRATION | NIR | angiogenesis | |
| GO_REGULATION_OF_ENDOTHELIAL_CELL_CHEMOTAXIS | NIR | angiogenesis | |
| GO_REGULATION_OF_ENDOTHELIAL_CELL_MIGRATION | NIR | angiogenesis | |
| GO_REGULATION_OF_ENDOTHELIAL_CELL_PROLIFERATION | NIR | angiogenesis | |
| GO_REGULATION_OF_EPITHELIAL_CELL_MIGRATION | NIR | angiogenesis | |
| GO_REGULATION_OF_VASCULAR_ENDOTHELIAL_GROWTH_FACTOR_RECEPTOR_SIGNALING_PATHWAY | NIR | angiogenesis | |
| GO_NEGATIVE_REGULATION_OF_SMOOTH_MUSCLE_CELL_PROLIFERATION | NIR | angiogenesis | |
| GO_ARTERY_MORPHOGENESIS | NIR | angiogenesis | |
| GO_INTRACILIARY_TRANSPORT | PIR | cell structure | intracellular transport |
| GO_INTRACILIARY_TRANSPORT_PARTICLE | PIR | cell structure | intracellular transport |
| GO_INTRACILIARY_TRANSPORT_PARTICLE_B | PIR | cell structure | intracellular transport |
| GO_ANCHORED_COMPONENT_OF_EXTERNAL_SIDE_OF_PLASMA_MEMBRANE | NIR | cell structure | plasma membrane |
| GO_ANCHORED_COMPONENT_OF_PLASMA_MEMBRANE | NIR | cell structure | plasma membrane |
| GO_INTRINSIC_COMPONENT_OF_EXTERNAL_SIDE_OF_PLASMA_MEMBRANE | NIR | cell structure | plasma membrane |
| GO_AXONEMAL_DYNEIN_COMPLEX_ASSEMBLY | PIR | cell structure | |
| GO_AXONEME_ASSEMBLY | PIR | cell structure | |
| GO_AXONEME_PART | PIR | cell structure | |
| GO_CELLULAR_COMPONENT_ASSEMBLY_INVOLVED_IN_MORPHOGENESIS | PIR | cell structure | |
| GO_CELL_PROJECTION_ASSEMBLY | PIR | cell structure | |
| GO_CENTRIOLAR_SATELLITE | PIR | cell structure | |
| GO_CILIARY_BASAL_BODY | PIR | cell structure | |
| GO_CILIARY_PART | PIR | cell structure | |
| GO_CILIARY_PLASM | PIR | cell structure | |
| GO_CILIARY_TIP | PIR | cell structure | |
| GO_CILIARY_TRANSITION_ZONE | PIR | cell structure | |
| GO_CILIUM | PIR | cell structure | |
| GO_CILIUM_MORPHOGENESIS | PIR | cell structure | |
| GO_CILIUM_MOVEMENT | PIR | cell structure | |
| GO_CILIUM_ORGANIZATION | PIR | cell structure | |
| GO_DYNEIN_COMPLEX | PIR | cell structure | |
| GO_EPITHELIAL_CILIUM_MOVEMENT | PIR | cell structure | |
| GO_MICROTUBULE_BASED_MOVEMENT | PIR | cell structure | |
| GO_MICROTUBULE_BUNDLE_FORMATION | PIR | cell structure | |
| GO_MOTILE_CILIUM | PIR | cell structure | |
| GO_NONMOTILE_PRIMARY_CILIUM | PIR | cell structure | |
| GO_NONMOTILE_PRIMARY_CILIUM_ASSEMBLY | PIR | cell structure | |
| GO_PHOTORECEPTOR_CONNECTING_CILIUM | PIR | cell structure | |
| GO_PRIMARY_CILIUM | PIR | cell structure | |
| GO_PROTEIN_COMPLEX_LOCALIZATION | PIR | cell structure | |
| GO_PROTEIN_TRANSPORT_APIR_MICROTUBULE | PIR | cell structure | |
| GO_NEGATIVE_REGULATION_OF_NF_KAPPAB_IMPORT_INTO_NUCLEUS | NIR | immune system | cytokine |
| GO_NEGATIVE_REGULATION_OF_INTERLEUKIN_12_PRODUCTION | NIR | immune system | cytokine |
| GO_CYTOKINE_BINDING | NIR | immune system | cytokine |
| GO_MODULATION_BY_HOST_OF_VIRAL_PROCESS | NIR | immune system | defense response |
| GO_INFLAMMATORY_RESPONSE_TO_ANTIGENIC_STIMULUS | NIR | immune system | defense response |
| GO_POSITIVE_REGULATION_OF_NATURAL_KILLER_CELL_MEDIATED_IMMUNITY | NIR | immune system | lymphoid lineage |
| GO_T_CELL_DIFFERENTIATION_INVOLVED_IN_IMMUNE_RESPONSE | NIR | immune system | lymphoid lineage |
| GO_ANTIGEN_PROCESSING_AND_PRESENTATION_VIA_MHC_CLASS_IB | NIR | immune system | lymphoid lineage |
| GO_MAST_CELL_GRANULE | NIR | immune system | myeloid lineage |
| GO_REGULATION_OF_MACROPHAGE_ACTIVATION | NIR | immune system | myeloid lineage |
| GO_NEGATIVE_REGULATION_OF_LYMPHOCYTE_APOPTOTIC_PROCESS | NIR | immune system | lymphoid lineage |
| GO_NEGATIVE_REGULATION_OF_T_CELL_APOPTOTIC_PROCESS | NIR | immune system | lymphoid lineage |
| GO_POSITIVE_REGULATION_OF_LYMPHOCYTE_APOPTOTIC_PROCESS | NIR | immune system | lymphoid lineage |
| GO_REGULATION_OF_B_CELL_APOPTOTIC_PROCESS | NIR | immune system | lymphoid lineage |
| GO_REGULATION_OF_LYMPHOCYTE_APOPTOTIC_PROCESS | NIR | immune system | lymphoid lineage |
| GO_REGULATION_OF_T_CELL_APOPTOTIC_PROCESS | NIR | immune system | lymphoid lineage |
| GO_AMMONIUM_ION_BINDING | NIR | metabolism | ammonium |
| GO_PHOSPHATIDYLCHOLINE_BINDING | NIR | metabolism | ammonium |
| GO_QUATERNARY_AMMONIUM_GROUP_BINDING | NIR | metabolism | ammonium |
| GO_POSITIVE_REGULATION_OF_APOPTOTIC_SIGNALING_PATHWAY | NIR | metabolism | apotosis |
| GO_POSITIVE_REGULATION_OF_EXTRINSIC_APOPTOTIC_SIGNALING_PATHWAY_VIA_DEATH_DOMAIN_RECEPTORS | NIR | metabolism | apotosis |
| GO_POSITIVE_REGULATION_OF_INTRINSIC_APOPTOTIC_SIGNALING_PATHWAY | NIR | metabolism | apotosis |
| GO_ATP_HYDROLYSIS_COUPLED_TRANSMEMBRANE_TRANSPORT | NIR | metabolism | cell respiration |
| GO_HYDROGEN_EXPORTING_ATPASE_ACTIVITY | NIR | metabolism | cell respiration |
| GO_PH_REDUCTION | NIR | metabolism | cell respiration |
| GO_PROTON_TRANSPORTING_TWO_SECTOR_ATPASE_COMPLEX_CATALYTIC_DOMAIN | NIR | metabolism | cell respiration |
| GO_PROTON_TRANSPORTING_V_TYPE_ATPASE_COMPLEX | NIR | metabolism | cell respiration |
| GO_REGULATION_OF_CELLULAR_PH | NIR | metabolism | cell respiration |
| GO_REGULATION_OF_PH | NIR | metabolism | cell respiration |
| GO_COATED_PIT | NIR | metabolism | cytosis |
| GO_PROTEIN_LIPID_COMPLEX_BINDING | NIR | metabolism | cytosis |
| GO_RECEPTOR_MEDIATED_ENDOCYTOSIS | NIR | metabolism | cytosis |
| GO_EMBRYONIC_HEMOPOIESIS | NIR | metabolism | homeostasis |
| GO_MACROMOLECULE_METHYLATION | PIR | metabolism | methylation |
| GO_S_ADENOSYLMETHIONINE_DEPENDENT_METHYLTRANSFERASE_ACTIVITY | PIR | metabolism | methylation |
| GO_TRNA_METHYLATION | PIR | metabolism | methylation |
| GO_NEGATIVE_REGULATION_OF_ERBB_SIGNALING_PATHWAY | NIR | metabolism | signal transport |

| | | | |
|---|---|---|---|
| GO_NEGATIVE_REGULATION_OF_PROTEIN_TYROSINE_KINASE_ACTIVITY | NIR | metabolism | signal transport |
| GO_REGULATION_OF_PROTEIN_TYROSINE_KINASE_ACTIVITY | NIR | metabolism | signal transport |
| GO_CHOLESTEROL_HOMEOSTASIS | NIR | metabolism | sterol |
| GO_HIGH_DENSITY_LIPOPROTEIN_PARTICLE | NIR | metabolism | sterol |
| GO_PLASMA_LIPOPROTEIN_PARTICLE_CLEARANCE | NIR | metabolism | sterol |
| GO_STEROL_HOMEOSTASIS | NIR | metabolism | sterol |
| GO_TRIGLYCERIDE_RICH_LIPOPROTEIN_PARTICLE | NIR | metabolism | sterol |
| GO_VERY_LOW_DENSITY_LIPOPROTEIN_PARTICLE | NIR | metabolism | sterol |
| GO_FILAMENTOUS_ACTIN | NIR | organogenesis | muscle |
| GO_ACTIN_CYTOSKELETON_REORGANIZATION | NIR | organogenesis | muscle |
| GO_ACTOMYOSIN | NIR | organogenesis | muscle |
| GO_POSITIVE_REGULATION_OF_HEART_CONTRACTION | NIR | physiological function | Cardiovascular function |
| GO_POSITIVE_REGULATION_OF_HEART_RATE | NIR | physiological function | Cardiovascular function |
| GO_POSITIVE_REGULATION_OF_STRIATED_MUSCLE_CONTRACTION | NIR | physiological function | Cardiovascular function |
| GO_REGULATION_OF_MUSCLE_CONTRACTION | NIR | physiological function | Cardiovascular function |
| GO_SPECIFICATION_OF_SYMMETRY | PIR | physiological function | fertility |
| GO_SPERM_FLAGELLUM | PIR | physiological function | fertility |
| GO_SPERM_MOTILITY | PIR | physiological function | fertility |
| GO_FERTILIZATION | PIR | physiological function | fertility |
| GO_SINGLE_FERTILIZATION | PIR | physiological function | fertility |
| GO_SPERM_EGG_RECOGNITION | PIR | physiological function | fertility |
| GO_REGULATION_OF_RESPIRATORY_SYSTEM_PROCESS | PIR | physiological function | respiratoration |
| GO_REGULATION_OF_RESPIRATORY_GASEOUS_EXCHANGE | PIR | physiological function | respiratoration |

**Supplementary Data 2D. The grouping of GO terms in Cohort 4.**

| GO_term | Enrichment in | Type 1 | Type 2 | Type 3 |
|---|---|---|---|---|
| GO_REGULATION_OF_CELL_SUBSTRATE_ADHESION | NIR | cell migration | | |
| GO_POSITIVE_REGULATION_OF_CELL_JUNCTION_ASSEMBLY | NIR | cell migration | | |
| GO_EXTRACELLULAR_MATRIX | NIR | cell structure | Extracellular components | |
| GO_EXTRACELLULAR_MATRIX_COMPONENT | NIR | cell structure | Extracellular components | |
| GO_COLLAGEN_TRIMER | NIR | cell structure | Extracellular components | |
| GO_FRIZZLED_BINDING | NIR | cell structure | Extracellular components | |
| GO_PROTEINACEOUS_EXTRACELLULAR_MATRIX | NIR | cell structure | Extracellular components | |
| GO_RECEPTOR_AGONIST_ACTIVITY | NIR | cell structure | Extracellular components | |
| GO_COLLAGEN_FIBRIL_ORGANIZATION | NIR | cell structure | Extracellular components | |
| GO_RECEPTOR_ACTIVATOR_ACTIVITY | NIR | cell structure | Extracellular components | |
| GO_RECEPTOR_REGULATOR_ACTIVITY | NIR | cell structure | Extracellular components | |
| GO_ER_TO_GOLGI_TRANSPORT_VESICLE | NIR | cell structure | Intracellular components | |
| GO_ER_TO_GOLGI_TRANSPORT_VESICLE_MEMBRANE | NIR | cell structure | Intracellular components | |
| GO_REGULATION_OF_MEMBRANE_LIPID_DISTRIBUTION | NIR | cell structure | trans-membrane | |
| GO_LIPID_TRANSLOCATION | NIR | cell structure | trans-membrane | |
| GO_TUMOR_NECROSIS_FACTOR_RECEPTOR_SUPERFAMILY_BINDING | NIR | immune system | cytokine | |
| GO_TUMOR_NECROSIS_FACTOR_RECEPTOR_BINDING | NIR | immune system | cytokine | |
| GO_REGULATION_OF_I_KAPPAB_KINASE_NF_KAPPAB_SIGNALING | NIR | immune system | cytokine | |
| GO_DEATH_RECEPTOR_BINDING | NIR | immune system | cytokine | |
| GO_NEGATIVE_REGULATION_OF_I_KAPPAB_KINASE_NF_KAPPAB_SIGNALING | NIR | immune system | cytokine | |
| GO_NEGATIVE_REGULATION_OF_CYTOKINE_BIOSYNTHETIC_PROCESS | NIR | immune system | cytokine | |
| GO_NEGATIVE_REGULATION_OF_CYTOKINE_PRODUCTION | NIR | immune system | cytokine | |
| GO_INTERACTION_WITH_SYMBIONT | NIR | immune system | defense response | |
| GO_NEGATIVE_REGULATION_OF_RESPONSE_TO_BIOTIC_STIMULUS | NIR | immune system | defense response | |
| GO_NEGATIVE_REGULATION_OF_MULTI_ORGANISM_PROCESS | NIR | immune system | defense response | |
| GO_REGULATION_OF_MULTI_ORGANISM_PROCESS | NIR | immune system | defense response | |
| GO_REGULATION_OF_SYMBIOSIS_ENCOMPASSING_MUTUALISM_THROUGH_PARASITISM | NIR | immune system | defense response | |
| GO_REGULATION_OF_DEFENSE_RESPONSE_TO_VIRUS | NIR | immune system | defense response | |
| GO_NEGATIVE_REGULATION_OF_VIRAL_TRANSCRIPTION | NIR | immune system | defense response | |
| GO_REGULATION_OF_LIPOPOLYSACCHARIDE_MEDIATED_SIGNALING_PATHWAY | NIR | immune system | defense response | |
| GO_NEGATIVE_REGULATION_OF_VIRAL_PROCESS | NIR | immune system | defense response | |
| GO_NEGATIVE_REGULATION_OF_DEFENSE_RESPONSE_TO_VIRUS | NIR | immune system | defense response | |
| GO_MODIFICATION_OF_MORPHOLOGY_OR_PHYSIOLOGY_OF_OTHER_ORGANISM | NIR | immune system | defense response | |
| GO_REGULATION_OF_MACROPHAGE_CHEMOTAXIS | NIR | immune system | myeloid lineage | |
| GO_REGULATION_OF_MONONUCLEAR_CELL_MIGRATION | NIR | immune system | myeloid lineage | |
| GO_VITAMIN_BINDING | NIR | metablolism | oxidoreduction | |
| GO_L_ASCORBIC_ACID_BINDING | NIR | metablolism | oxidoreduction | |
| GO_DIOXYGENASE_ACTIVITY | NIR | metablolism | oxidoreduction | |
| GO_PEPTIDYL_PROLINE_MODIFICATION | NIR | metablolism | oxidoreduction | |
| CORPORATION_OR_REDUCTION_OF_MOLECULAR_OXYGEN_2_OXOGLUTARATE_AS_ONE_DONOR_AND_IN | NIR | metablolism | oxidoreduction | |
| GO_POSITIVE_REGULATION_OF_CARBOHYDRATE_METABOLIC_PROCESS | NIR | metabolism | carbonhydrate | |
| GO_REGULATION_OF_CARBOHYDRATE_METABOLIC_PROCESS | NIR | metabolism | carbonhydrate | |
| GO_OXIDOREDUCTASE_ACTIVITY_ACTING_ON_A_HEME_GROUP_OF_DONORS | PIR | metabolism | cellular respiration | |
| GO_MITOCHONDRIAL_ELECTRON_TRANSPORT_CYTOCHROME_C_TO_OXYGEN | PIR | metabolism | cellular respiration | |
| GO_HYDROGEN_ION_TRANSMEMBRANE_TRANSPORT | PIR | metabolism | cellular respiration | |
| GO_RESPIRATORY_CHAIN | PIR | metabolism | cellular respiration | |
| GO_CYTOCHROME_COMPLEX | PIR | metabolism | cellular respiration | |
| GO_HYDROGEN_ION_TRANSMEMBRANE_TRANSPORTER_ACTIVITY | PIR | metabolism | cellular respiration | |
| GO_REGULATION_OF_CALCIUM_ION_DEPENDENT_EXOCYTOSIS | PIR | metabolism | cytosis | exo- |
| GO_REGULATION_OF_SYNAPTIC_VESICLE_EXOCYTOSIS | PIR | metabolism | cytosis | exo- |
| GO_MEMBRANE_INVAGINATION | NIR | metabolism | cytosis | phago- |
| GO_PHAGOCYTOSIS_ENGULFMENT | NIR | metabolism | cytosis | phago- |
| GO_GLUCURONATE_METABOLIC_PROCESS | PIR | metabolism | others | |
| GO_URONIC_ACID_METABOLIC_PROCESS | PIR | metabolism | others | |
| GO_SULFURIC_ESTER_HYDROLASE_ACTIVITY | NIR | metabolism | others | |
| GO_ENDOPLASMIC_RETICULUM_LUMEN | NIR | metabolism | others | |
| GO_RESPONSE_TO_TOPOLOGICALLY_INCORRECT_PROTEIN | NIR | metabolism | protein | |
| GO_RETROGRADE_PROTEIN_TRANSPORT_ER_TO_CYTOSOL | NIR | metabolism | protein | |
| GO_RESPONSE_TO_ENDOPLASMIC_RETICULUM_STRESS | NIR | metabolism | protein | |
| GO_ERAD_PATHWAY | NIR | metabolism | protein | |
| GO_REGULATION_OF_ENDOPLASMIC_RETICULUM_UNFOLDED_PROTEIN_RESPONSE | NIR | metabolism | protein | |
| GO_ENDOPLASMIC_RETICULUM_TO_CYTOSOL_TRANSPORT | NIR | metabolism | protein | |
| GO_PROTEIN_EXIT_FROM_ENDOPLASMIC_RETICULUM | NIR | metabolism | protein | |
| GO_CELLULAR_RESPONSE_TO_TOPOLOGICALLY_INCORRECT_PROTEIN | NIR | metabolism | protein | |
| GO_ER_ASSOCIATED_UBIQUITIN_DEPENDENT_PROTEIN_CATABOLIC_PROCESS | NIR | metabolism | protein | |
| GO_AMINOGLYCAN_METABOLIC_PROCESS | NIR | metabolism | proteoglycan | |
| GO_HEPARAN_SULFATE_PROTEOGLYCAN_BIOSYNTHETIC_PROCESS | NIR | metabolism | proteoglycan | |
| GO_PROTEOGLYCAN_METABOLIC_PROCESS | NIR | metabolism | proteoglycan | |
| GO_HEPARAN_SULFATE_PROTEOGLYCAN_METABOLIC_PROCESS | NIR | metabolism | proteoglycan | |
| GO_AMINOGLYCAN_BIOSYNTHETIC_PROCESS | NIR | metabolism | proteoglycan | |
| GO_REGULATION_OF_ALCOHOL_BIOSYNTHETIC_PROCESS | NIR | metabolism | sterols | |
| GO_NEGATIVE_REGULATION_OF_STEROID_METABOLIC_PROCESS | NIR | metabolism | sterols | |
| GO_NEGATIVE_REGULATION_OF_LIPID_BIOSYNTHETIC_PROCESS | NIR | metabolism | sterols | |
| GO_REGULATION_OF_STEROID_METABOLIC_PROCESS | NIR | metabolism | sterols | |
| GO_NEGATIVE_REGULATION_OF_ALCOHOL_BIOSYNTHETIC_PROCESS | NIR | metabolism | sterols | |
| GO_POSITIVE_REGULATION_OF_CARTILAGE_DEVELOPMENT | NIR | organogenesis | bone development | cartilage development |
| GO_CHONDROCYTE_DEVELOPMENT | NIR | organogenesis | bone development | cartilage development |
| GO_CONNECTIVE_TISSUE_DEVELOPMENT | NIR | organogenesis | bone development | cartilage development |
| GO_POSITIVE_REGULATION_OF_CHONDROCYTE_DIFFERENTIATION | NIR | organogenesis | bone development | cartilage development |
| GO_BONE_MORPHOGENESIS | NIR | organogenesis | bone development | cartilage development |
| GO_REGULATION_OF_CHONDROCYTE_DIFFERENTIATION | NIR | organogenesis | bone development | cartilage development |
| GO_ENDOCHONDRAL_BONE_MORPHOGENESIS | NIR | organogenesis | bone development | cartilage development |
| GO_CHONDROCYTE_DIFFERENTIATION | NIR | organogenesis | bone development | cartilage development |
| GO_CARTILAGE_DEVELOPMENT_INVOLVED_IN_ENDOCHONDRAL_BONE_MORPHOGENESIS | NIR | organogenesis | bone development | cartilage development |
| GO_CARTILAGE_DEVELOPMENT | NIR | organogenesis | bone development | cartilage development |
| GO_REGULATION_OF_CARTILAGE_DEVELOPMENT | NIR | organogenesis | bone development | cartilage development |
| GO_REGULATION_OF_OSTEOBLAST_DIFFERENTIATION | NIR | organogenesis | bone development | ossfication |
| GO_POSITIVE_REGULATION_OF_OSSIFICATION | NIR | organogenesis | bone development | ossfication |
| GO_POSITIVE_REGULATION_OF_OSTEOBLAST_DIFFERENTIATION | NIR | organogenesis | bone development | ossfication |
| GO_SYNCYTIUM_FORMATION | NIR | organogenesis | muscle development | |
| GO_MYOBLAST_FUSION | NIR | organogenesis | muscle development | |
| GO_STRIATED_MUSCLE_CELL_DIFFERENTIATION | NIR | organogenesis | muscle development | |
| GO_CARDIAC_MUSCLE_CELL_DIFFERENTIATION | NIR | organogenesis | muscle development | |
| GO_SUBSTANTIA_NIGRA_DEVELOPMENT | PIR | organogenesis | neural development | |
| GO_NEURAL_NUCLEUS_DEVELOPMENT | PIR | organogenesis | neural development | |
| GO_NEURON_PROJECTION_REGENERATION | NIR | organogenesis | regeneration | |
| GO_NEURON_PROJECTION_REGENERATION | NIR | organogenesis | regeneration | |
| GO_ORGAN_REGENERATION | NIR | organogenesis | regeneration | |
| GO_REGENERATION | NIR | organogenesis | regeneration | |

| | | | |
|---|---|---|---|
| GO_ORGAN_REGENERATION | NIR | organogenesis | regeneration |
| GO_SERTOLI_CELL_DEVELOPMENT | NIR | physiological function | fertility |
| GO_SERTOLI_CELL_DIFFERENTIATION | NIR | physiological function | fertility |

**Supplementary Data 3. Classification of all eight samples with scRNA-seq data available into NIR or PIR clusters.**

| | | | | Cohort 1 | | | Cohort 2 | | | Cohort 3 | | | Cohort 4 | | |
|---|---|---|---|---|---|---|---|---|---|---|---|---|---|---|---|
| | **total foldchange** | **Cluster** | **macrophage proportion** | **PIR** | **NIR** | **foldchange** | **PIR** | **NIR** | **foldchange** | **PIR** | **NIR** | **foldchange** | **PIR** | **NIR** | **foldchange** |
| **PJ016** | 0.957350139 | NIR | 0.06% | 0.8858871 | 0.8665323 | 1.02233597 | 0.8816182 | 0.9370079 | 0.9408867 | 0.9133958 | 0.917739 | 0.995267526 | 1 | 1 | 1 |
| **PJ017** | 0.622907489 | NIR | 46.63% | 0.0846774 | 0.0302419 | 2.8 | 0.0045707 | 0.0205453 | 0.22246696 | 1 | 1 | 1 | 1 | 1 | 1 |
| **PJ018** | 3.506670298 | PIR | 2.28% | 0.3677419 | 0.2423387 | 1.517470882 | 1 | 1 | 1 | 0.1408291 | 0.1198263 | 1.175277252 | 0.1920917 | 0.0976955 | 1.9662296 |
| **PJ025** | 4.050609406 | PIR | 1.70% | 0.0697581 | 0.0177419 | 3.931818182 | 1 | 1 | 1 | 0.4597608 | 0.4462775 | 1.030212797 | 1 | 1 | 1 |
| **PJ030** | 1.629268176 | PIR | 8.33% | 0.3112903 | 0.275 | 1.131964809 | 0.1394696 | 0.1247172 | 1.118287373 | 0.2588727 | 0.250705 | 1.032579186 | 0.1355575 | 0.1087528 | 1.246473 |
| **PJ032** | 0.900833333 | NIR | 55.12% | 1 | 1 | 1 | 0.1956738 | 0.2172142 | 0.900833333 | 1 | 1 | 1 | 1 | 1 | 1 |
| **PJ035** | 1.161568474 | PIR | 8.12% | 0.2616935 | 0.2516129 | 1.040064103 | 0.1916916 | 0.1477057 | 1.297794118 | 1 | 0.0161735 | 1 | 0.2324331 | 0.2700966 | 0.8605555 |
| **PJ048** | 0.27266869 | NIR | 0% | 1 | 1 | 1 | 1 | 1 | 1 | 0.0043756 | 0.0160438 | 0.272727273 | 0.2125374 | 0.2125831 | 0.9997852 |

Negative coefficient value is not taken into consideration. To simplify the caculation, those values were replaced with 1.

**Supplementary Data 4. Clinical information and IHC staning results for 12 patients.**

| ID | OS | OS_Censor | PFS | PFS_Censor | MS4A4A | Grade |
|---|---|---|---|---|---|---|
| CGGA_1481 | 4.37 | 1 | 0.933333333 | 1 | 27.579 | 4 |
| CGGA_P25 | 4.90 | 1 | 4.9 | 1 | 35.905 | 4 |
| CGGA_1422 | 6.80 | 1 | 6.7 | 1 | 29.481 | 4 |
| CGGA_1521 | 6.83 | 1 | 5.833333333 | 1 | 21.728 | 4 |
| CGGA_P143 | 8.70 | 1 | 8.7 | 1 | 41.402 | 4 |
| CGGA_1494 | 8.97 | 1 | 8.966666667 | 1 | 21.35 | 4 |
| CGGA_P178 | 25.87 | 0 | 25.86666667 | 0 | 0.853 | 4 |
| CGGA_1735 | 27.10 | 1 | 22 | 1 | 1.516 | 4 |
| CGGA_1467 | 28.87 | 1 | 20.76666667 | 1 | 1.596 | 4 |
| CGGA_1282 | 37.20 | 1 | 20 | 1 | 1.241 | 4 |
| CGGA_1780 | 37.43 | 0 | 35.56666667 | 1 | 0.818 | 4 |
| CGGA_1086 | 65.9 | 1 | 42.23333333 | 1 | 2.013 | 4 |
